# Curated and harmonised transcriptomics datasets of interstitial lung diseases

**DOI:** 10.1101/2025.04.08.647776

**Authors:** Simo Inkala, Antonio Federico, Angela Serra, Dario Greco

**Author notes:** These authors contributed equally (Simo Inkala, Antonio Federico). Corresponding author’s email address and Twitter handle.

## Abstract

This study provides manually curated and homogenised transcriptomics data of interstitial lung disease (ILD) patients retrieved from the NCBI Gene Expression Omnibus and European Nucleotide Archive repositories. The compendium includes 30 transcriptomics datasets generated with DNA microarrays and RNA sequencing technologies for a total of 1,371 samples. All the datasets underwent metadata curation and harmonisation, data quality check, and preprocessing with standardised procedures. Furthermore, a robust data model was developed to standardise phenotypic data, thereby enhancing comparability across heterogeneous datasets. Gene expression data and lists of differentially expressed genes computed between ILD and healthy samples are provided. Among the ILDs included in this study, idiopathic pulmonary fibrosis (IPF) is the most represented worldwide. Co-expression networks of IPF and healthy samples were inferred, which are also included in this study. This work significantly improves the Findability, Accessibility, Interoperability, and Reusability (FAIR) of publicly available transcriptomics data of ILDs, providing a platform to implement and validate integrated systems biology and pharmacology approaches for novel interstitial lung disease diagnostics and therapeutics.

## SPECIFICATIONS TABLE

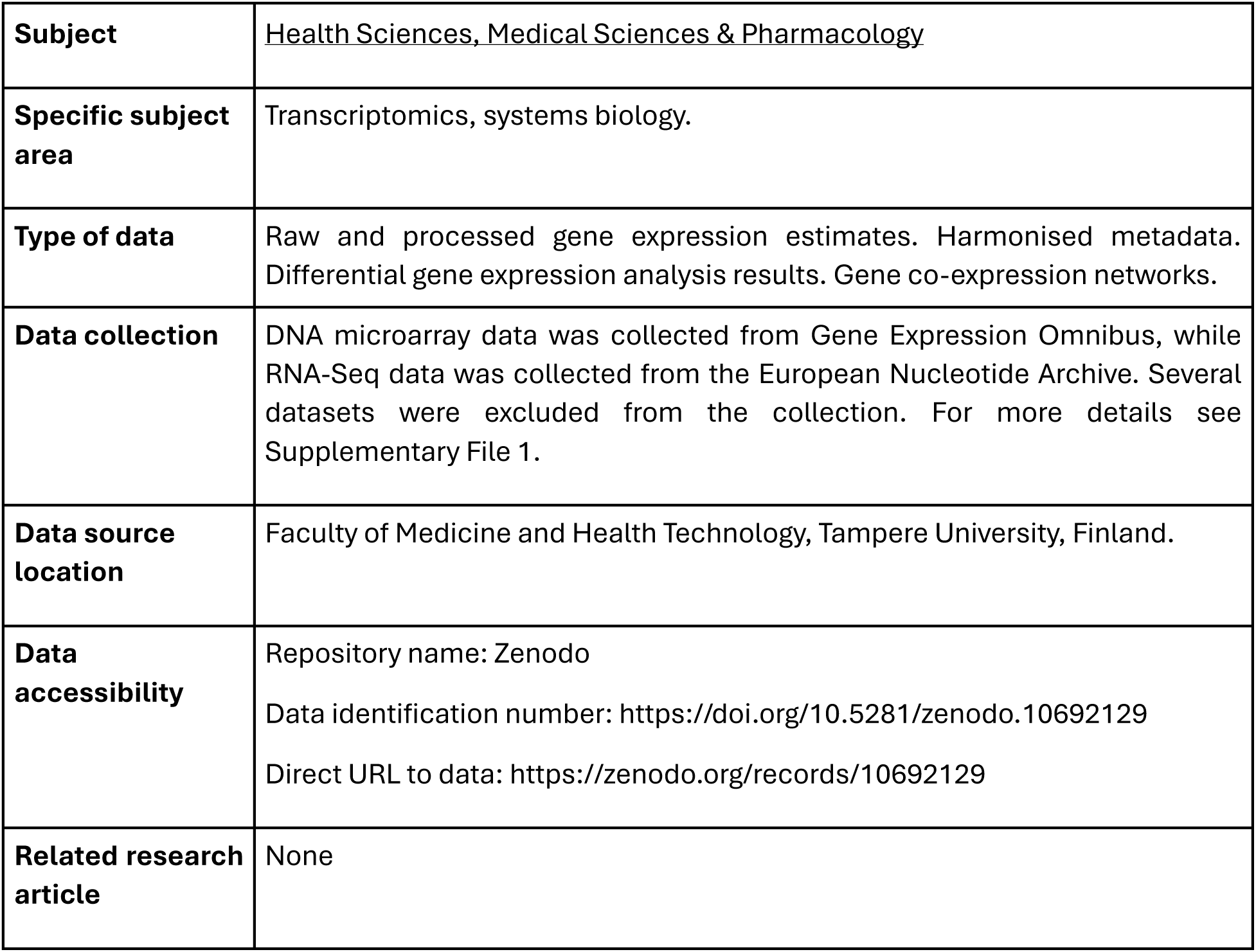

|  |  |
| --- | --- |
| **Subject** | Health Sciences, Medical Sciences & Pharmacology |
| **Specific subject area** | Transcriptomics, systems biology. |
| **Type of data** | Raw and processed gene expression estimates. Harmonised metadata. Differential gene expression analysis results. Gene co-expression networks. |
| **Data collection** | DNA microarray data was collected from Gene Expression Omnibus, while RNA-Seq data was collected from the European Nucleotide Archive. Several datasets were excluded from the collection. For more details see Supplementary File 1. |
| **Data source location** | Faculty of Medicine and Health Technology, Tampere University, Finland. |
| **Data accessibility** | Repository name: Zenodo<br>Data identification number: https://doi.org/10.5281/zenodo.10692129<br>Direct URL to data: https://zenodo.org/records/10692129 |
| **Related research article** | None |

## VALUE OF THE DATA

In this article, a comprehensive, harmonised, and “ready-to-use” FAIR collection of transcriptomics data of IPF patients is showcased. The data allow to elucidate the molecular mechanisms underlying ILDs through a wide range of possible applications. The scientific community can benefit from using these data by applying integrative methodologies (i.e. gene expression meta-analysis) across studies, technologies, and platforms to extrapolate robust gene expression signatures that underlie the clinical manifestations of the disease and its heterogeneity across patients.

In a previous effort [1], we presented a curated collection of transcriptomics datasets of patients affected by inflammatory dermatological diseases, such as psoriasis and atopic dermatitis. Our work posed the roots for a thorough exploitation of the data that resulted in high-impact scientific investigations of these two skin diseases. For instance, by analysing the collection of transcriptomics data of psoriatic patients, we were able to build a disease map through the representation of the lesional skin through network models [2]. This map allows to shed light on pivotal mechanisms underlying the lesional phenotype, pinpointing molecular effectors that can enrich the pool of current therapeutic targets. In addition, the integration of the psoriasis network model with single cell RNA-seq (scRNA-seq) data highlighted immune- and skin-specific transcriptional signatures and pharmacological properties of the psoriatic lesion. In another study [3], the same data collection was exploited by analysing transcriptomics data of atopic dermatitis patients. A network model of atopic dermatitis was built by integrating curated data with *a priori* knowledge (i.e. genomic variants, disease associated genes, known biomarkers, and drug targets). Given the high phenotypic heterogeneity of atopic dermatitis across patients, the main aim of the study was to facilitate tailored drug discovery efforts to improve the current pharmacological applications. Through the investigation of topological characteristics of the model drugs with potential activity towards atopic dermatitis were prioritised. Therefore, the application of cheminformatic methods (i.e. pharmacophore extrapolation followed by virtual screening) served to compile a compound library that will facilitate the development of more specific, efficient and safe therapeutic solutions for atopic dermatitis patients [3].

Data presented in this study offer the opportunity not only to carry out similar kinds of exploration in the field of ILDs, but also to expand towards the identification of disease subtypes or endotypes. In fact, ILDs include several phenotypes that, although very similar from the clinical point of view, arise from heterogeneous molecular makeups. For this reason, our data poses the basis for a classification of ILD patients into subgroups thoroughly characterised at a molecular resolution. Such an approach would bring the necessary innovation beyond the current diagnostic routines, merely based on phenotypical observation. In this context, this data could be an excellent reservoir to extract specific disease/subtype biomarkers. Currently, biomarker discovery is the driving force in profiling multi-factorial diseases, such as IPF. Although several biomarkers have already been identified for IPF, and more in general for ILDs, identifying novel molecular vulnerabilities of disease would improve current diagnostic capabilities and expand the repertoire of drug targets. In this regard, systems pharmacology is a relatively new field of research that aims to describe multi-factorial diseases as complex systems. In systems pharmacology, the intricate relations between molecular entities (i.e. genes or proteins) are disentangled and exploited to identify novel drugs with improved efficacy and safety for a given disease. In this article, gene co-expression networks of IPF patients are provided, which can be exploited to model the complexity of IPF and extrapolate candidate compounds that could improve existing pharmacological treatments.

## BACKGROUND

Interstitial lung diseases (ILD) encompass a spectrum of disorders characterised by chronic inflammation and scarring of the lung tissue, leading to a progressive impairment of the respiratory function. The most represented ILD in the global population is idiopathic pulmonary fibrosis (IPF). IPF is a chronic and progressive lung disease that has unknown origin. IPF is irreversible and usually lethal. To date, only a few treatment options with limited efficacy are available [4].

Transcriptomic technologies are established tools for elucidating the molecular intricacies underlying complex diseases, including ILDs. In addition, the analysis of molecular networks represents a well-established approach that leverages systems biology to investigate transcriptional relationships, define functionally related gene communities and pinpointing pivotal regulatory elements governing disease phenotypes [5]. Despite the wealth of transcriptomics data available for ILDs, these datasets are scattered across repositories, hindering comprehensive analysis and interpretation.

Here, we provide curated and preprocessed transcriptomics data of ILD patients and healthy counterparts obtained from public repositories, along with harmonised metadata.

By consolidating and standardising ILD datasets, this work aims to catalyse future research endeavours, foster collaboration, and expedite the discovery of novel insights and therapeutic strategies. The analytical pipeline of the study is shown in Fig. 1.

**Figure 1.**
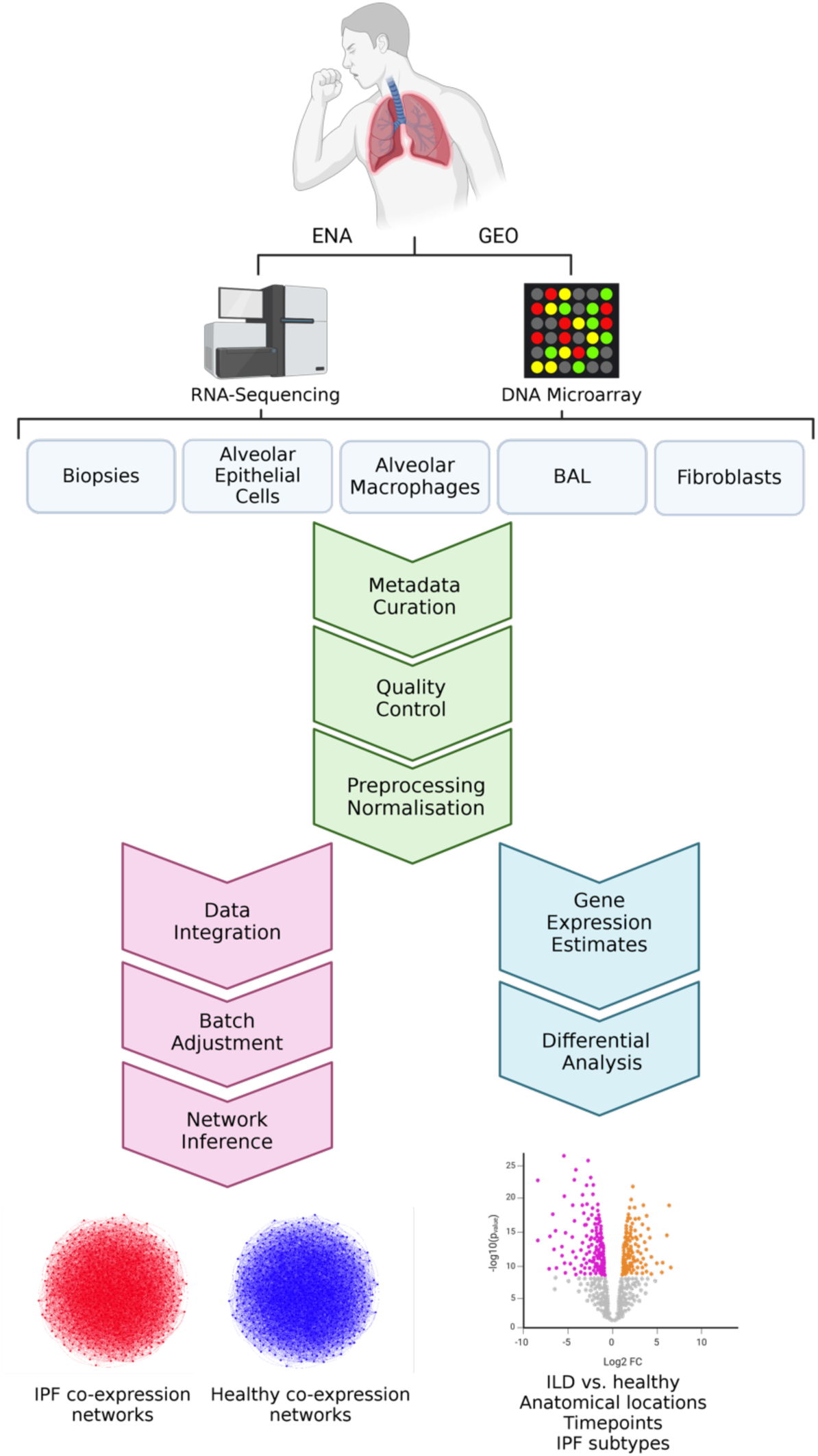
Analytical pipeline of the study.

## DATA DESCRIPTION

The preprocessed microarray datasets provided in this study were collected from the NCBI Gene Expression Omnibus (GEO) repository, while the RNA sequencing (RNA-seq) datasets were retrieved from the European Nucleotide Archive (ENA) (https://www.ebi.ac.uk/ena/browser/). Overall, 14 microarray datasets were collected, for a total of 471 samples, 297 of which were from patients affected by ILD and 174 samples from healthy individuals. These datasets were generated with commercially available Affymetrix and Agilent platforms (Table 1). 16 RNA-seq datasets were also retrieved, for a total of 900 samples, 589 of which were from patients affected by ILD and 311 samples from healthy individuals (Table 2). RNA-seq data were produced through either Illumina or Ion Torrent platforms.

**Table 1.** . **Microarray datasets included in the study.**

| **GEO dataset ID** | **Platform** | **Citation** | **Number of disease samples included** | **Number of healthy samples included** | **Cell type** |
| --- | --- | --- | --- | --- | --- |
| GSE53845 | GPL6480 | 82 | IPF: 40 | 8 | Biopsy |
| GSE110147 | GPL6244 | 83 | IPF: 22<br>NSIP: 10<br>Mixed IPF-NSIP: 5 | 11 | Biopsy |
| GSE21369 | GPL570 | 84 | UIP/IPF: 11<br>COP: 2<br>HP: 2<br>NSIP: 5<br>RB-ILD: 2<br>UF: 1 | 6 | Biopsy |
| GSE24206 | GPL570 | 85 | IPF: 17 | 6 | Biopsy |
| GSE72073 | GPL17586 | 86 | IPF: 5 | 3 | Biopsy |
| GSE10667 | GPL4133 | 87, 88, 89, 90 | UIP: 31 | 15 | Biopsy |
| GSE11196 | GPL6732 | 61 | IPF: 12 | 12 | Fibroblast |
| GSE129164 | GPL17586 | 64 | IPF: 10 | 10 | Fibroblast |
| GSE44723 | GPL570 | 91 | IPF: 10 | 4 | Fibroblast |
| GSE144338 | GPL22321 | 92, 93 | IPF: 4<br>Lung<br>Adenocarcinoma: 4 | 4 | Fibroblast |
| GSE40839 | GPL96 | 94 | UIP: 3<br>Sc-ILD: 8 | 10 | Fibroblast |
| GSE70866 | GPL14550<br>GPL17077 | 60 | IPF: 62 | 20 | BAL |
| GSE49072 | GPL96 | 63 | IPF: 15<br>Familial IPF: 8 | 61 | Macrophage |
| GSE90010 | GPL17692 | 62 | IPF: 4<br>RB-ILD: 4 | MDM: 4 | Macrophage |

*Abbreviations; IPF: Idiopathic Pulmonary Fibrosis, NSIP: Nonspecific Interstitial Pneumonia, Mixed IPF-NSIP: Mixed Idiopathic Pulmonary Fibrosis Nonspecific Interstitial Pneumonia, COP: Cryptogenic Organizing Pneumonia, HP: Hypersensitivity Pneumonitis, RB-ILD: Respiratory Bronchiolitis-Interstitial Lung Disease, UF: Uncharacterized Fibrosis, UIP: Usual Interstitial Pneumonia, Sc-ILD: Scleroderma-Associated Interstitial Lung Disease, MDM: Monocyte Derived Macrophage.*

**Table 2.**
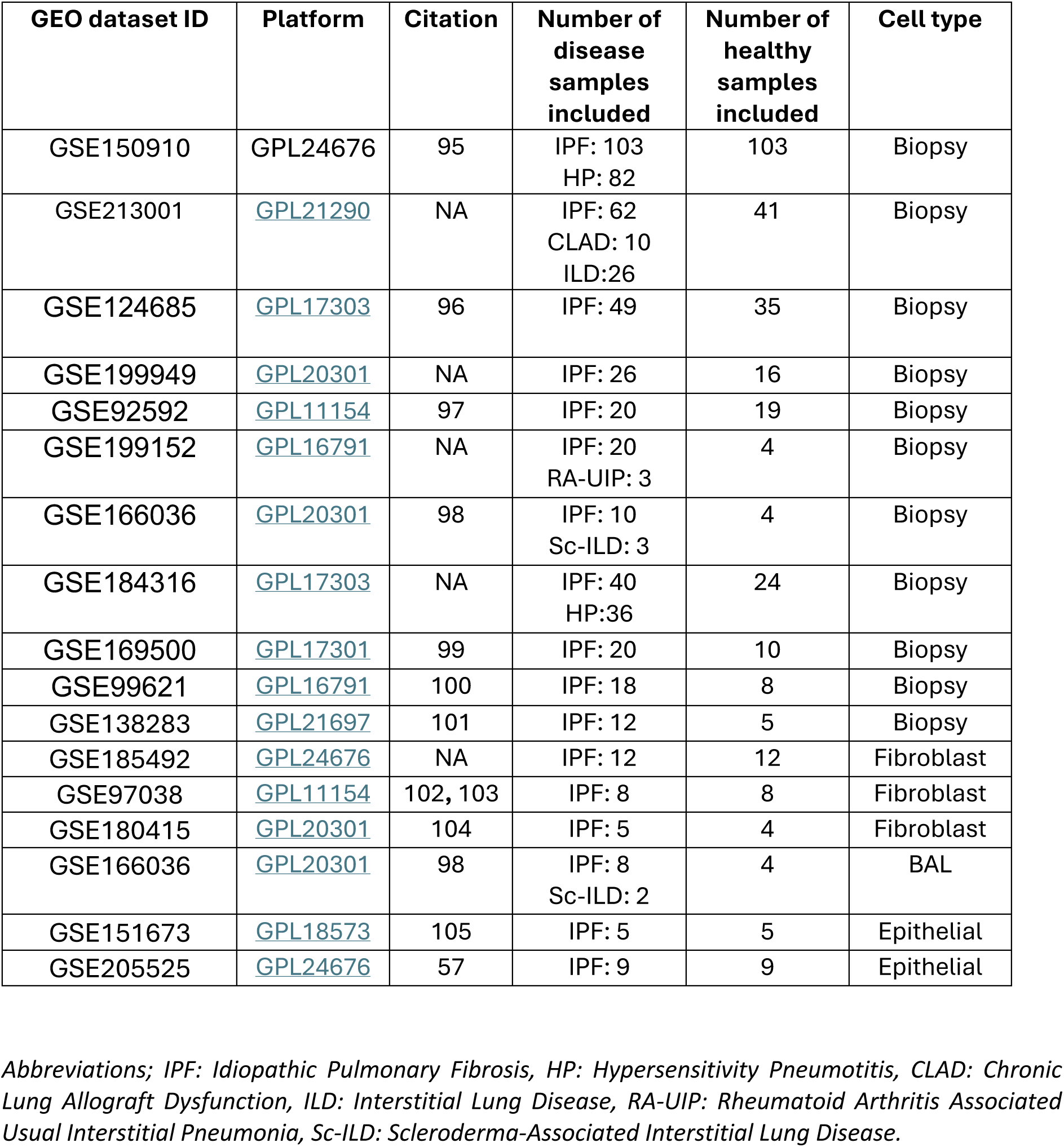
RNA-seq dataset included in the study.

The distribution of samples across the DNA microarray and RNA-Seq technologies are showed in Fig. 2A. IPF was the most represented disease across all the collected samples (49.1%), followed by hypersensitivity pneumonitis (8.75%, Fig. 2B). Collected datasets encompass lung biopsy, fibroblast, macrophage, alveolar epithelial, and bronchoalveolar lavage (BAL) samples.

**Figure 2.**
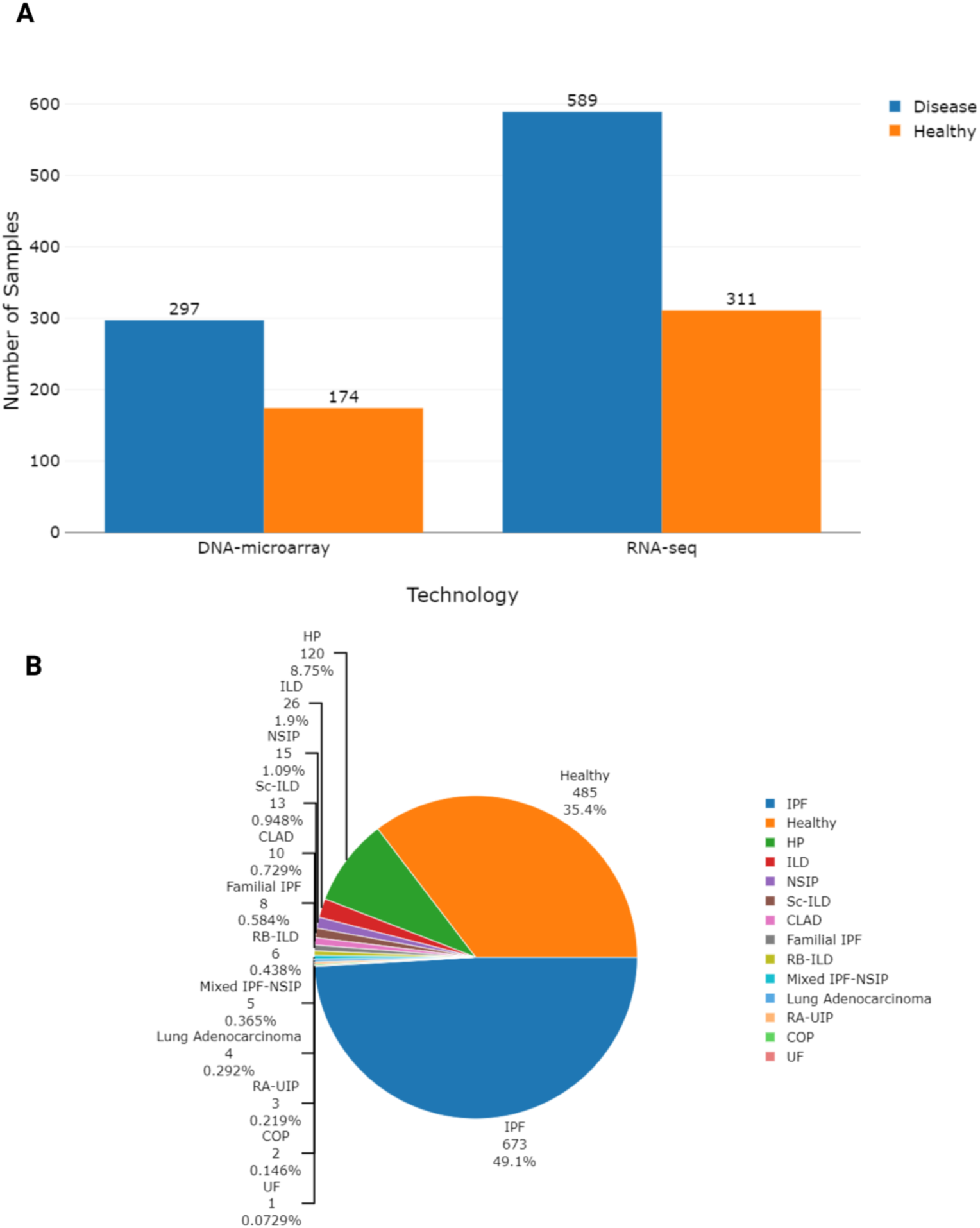
Distribution of samples across different conditions and technologies.

The progression of IPF is tightly correlated with the sex, with males experiencing higher mortality and earlier hospitalization and being more frequently represented in the IPF population compared to females. Advanced age has also been found to be associated with poor prognosis in IPF. Fig. 3 illustrates the distribution of samples among sexes across various ILDs and healthy counterparts when data was accessible. The figure highlights a predominant representation of males in the dataset, particularly evident in IPF cases.

**Figure 3.**
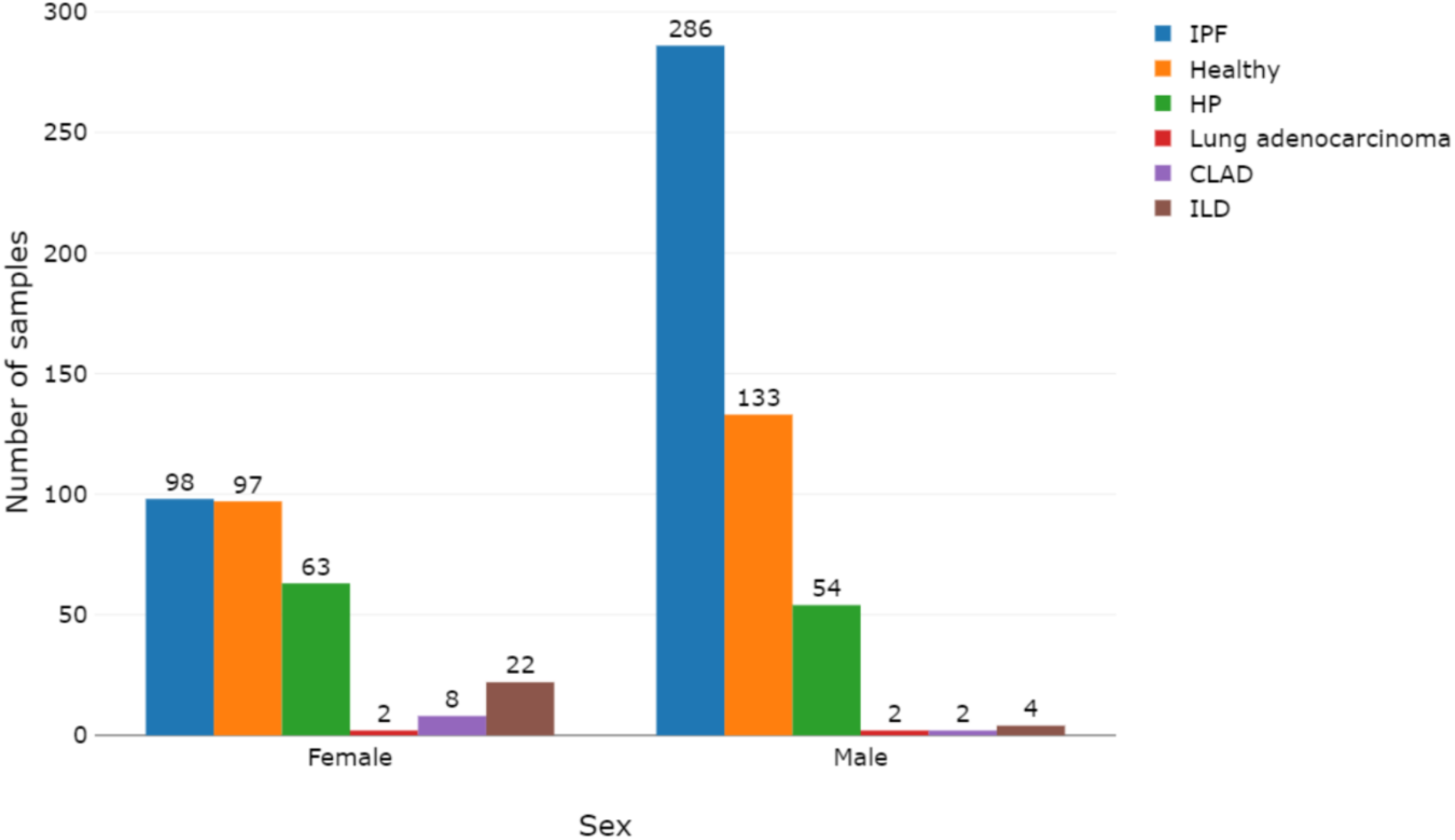
Sample distribution across sexes and conditions when the information was available.

Based on the available data, Fig. 4 displays the distribution of samples among age groups across different ILDs. The figure emphasizes that while the prevalence of IPF increases with age, higher mortality rates result in lower prevalence among the most advanced age groups.

**Figure 4.**
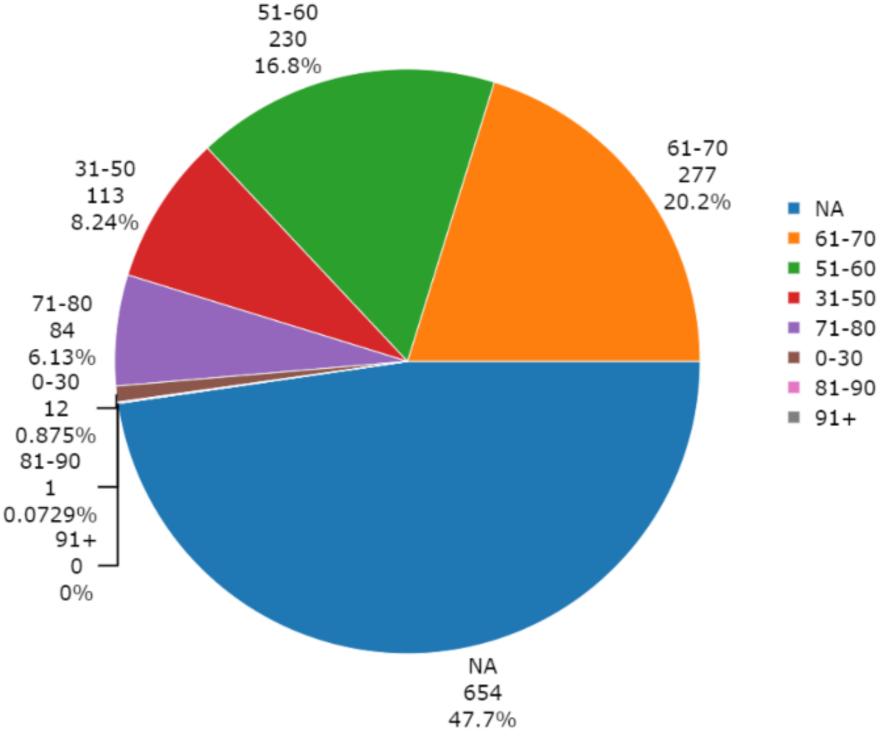
Distribution of samples across age groups. NA indicates samples where the information was not available.

The complexity of the disease makes IPF well-suited for network-based modeling by systems biology and pharmacology approaches that comprehend the intricate interactions of biomolecules. By integrating large-scale biological data and considering the interconnectedness of various signaling cascades, these methodologies offer a holistic understanding of disease pathogenesis, facilitating the identification of potential therapeutic targets and the development of more effective treatment strategies. In this regard, co-expression network models are built by using the curated gene expression estimates of each dataset to support future research in the fields of systems biology. The co-expression network models developed in this study present a robust tool for investigating the molecular landscape of IPF, enabling the identification of candidate biomarkers for disease diagnosis, progression, and treatment response. Additionally, they shed light on the intricate regulatory circuits and offer promising avenues for therapeutic intervention and dysregulated pathways underlying IPF pathogenesis, facilitating the discovery of novel therapeutic targets and the development of targeted therapies.

## EXPERIMENTAL DESIGN, MATERIALS AND METHODS

### Metadata harmonsation

The curation and harmonisation of metadata were performed with the ESPERANTO software [6]. ESPERANTO is designed for efficient semi-supervised curation tasks on omics metadata. The user actively participates in the decision process while harmonising data within a consistent framework and enhancing data FAIRness. This approach combines the benefits of both automated and manual curation methods. The data models, to which the raw metadata were mapped, are reported in the data dictionary file (enclosed with the preprocessed data). The data dictionary describes all the allowed variables, variable synonyms, allowed features, and feature synonyms reported in the final metadata tables. At the same time, this work aims to homogenise the preprocessing procedures to improve the comparability of gene expression data across different studies and platforms. Metadata visualization in Figs. 2-4 was created using the plotly 4.10.1 R-package [7].

### DNA microarray data

#### Data collection and homogenization

DNA microarray generated transcriptomics data were retrieved from NCBI GEO 21 (GEO - https://www.ncbi.nlm.nih.gov/geo/) repository by using the GEOquery [8] R package. For each dataset, a metadata table specifying the phenotype (i.e. IPF, healthy) and the biological system (i.e. biopsy, fibroblast, macrophage, epithelial, BAL), along with other phenotypic information, was also retrieved. The DNA microarray datasets with the GEO identifiers are reported in Table 1. The samples from patients with usual interstitial pneumonia (UIP) were included in the IPF group, as these two diseases are often used synonymously.

#### Data quality check

The retrieved datasets were evaluated by visual inspection of the quality check reports and multi-dimensional scaling (MDS) plots using the eUTOPIA software [9]. Furthermore, in the case of Affymetrix datasets, identification of outlier samples were identified by using the Normalized Unscaled Standard Error (NUSE) and the Relative Log Expression using the affyPLM v1.78.0 [10] R package and the RNA degradation curves (RNADeg) through the affy v1.76.0 R-package [11]. The Agilent quality control report was generated by using arrayQualityMetrics v3.54.0 [12].

#### Normalisation

Data normalisation was carried out by using the eUTOPIA software. The normalisation for Affymetrix-based studies was performed by using the *justRMA* from the R affy v1.76.0 package [13]. Normalisation of Agilent-based studies was performed with the *normalizeQuantiles* function from the limma v3.54.0 package [14].

#### Surrogate variable analysis

To analyse investigate the effect of unknown technical variables that might conceal biological variability, Surrogate Variable Analysis (SVA) was performed with the eUTOPIA software, which implements the sva v3.46.0 R package [15]. The analysis was performed by using the disease as a variable of interest. The other biological variables were used as covariates if present and not confounded with the variable of interest.

#### Differential gene expression analysis

The differential gene expression analysis was performed with eUTOPIA, which applies linear model implementation from the R-package limma. The *lmFit* function from the limma R-package fits gene-wise linear models to the microarray data. The variable of interest in the model was the diagnosis (disease vs. healthy), and other relevant biological and technical variables (if present and if not confounded with the variable of interest) were used as covariates. Also, comparisons between different lung locations, disease severity states and disease subtypes were performed if data was available. eUTOPIA applies *eBayes* function to assess differential expression by using the fitted model with the contrast coefficients. The p-value adjustment was carried out with the “Benjamini & Hochberg” method. The differential gene expression results were filtered by considering adjusted p-values less than or equal to 0.01 and absolute log-fold changes greater than or equal to 0.58.

#### Probe annotation

In eUTOPIA, the analysis was performed with the raw data for Agilent datasets and the expression matrix was aggregated by computing the median of the expression of the Agilent probes mapping to the same probe name. For Affymetrix-based microarrays custom annotation files (CDF files) were downloaded from Brainarray (http://brainarray.mbni.med.umich.edu/Brainarray/Database/CustomCDF/25.0.0/ensg.asp). Probe annotation was performed using biomaRt v.2.40.1. The probe names in expression matrices and differential gene expression tables were annotated to the Ensembl gene IDs and Gene Symbols using R. Gene names were grouped based on unique values with dplyr 1.0.2 package. Subsequently, the median of the expression value for each sample was calculated within each gene. This resulted in the creation of a new data frame, where each row represents a unique gene, and the columns contain the computed median values for the respective expression value associated with each gene. The analytical steps for processing Affymetrix DNA microarrays are illustrated in Supplementary Fig. 4, while the analytical processing steps for Agilent DNA microarrays are represented in Supplementary Fig. 5. Supplementary Figs. Figs. 4-7 are created with Biorender.com.

### RNA sequencing data

#### Data collection and homogenization

Raw files in “*.fastq*” format were obtained from the European Nucleotide Archive (ENA - https://www.ebi.ac.uk/ena/browser/home). In case the raw data was unavailable, we retrieved the normalised expression matrix from GEO, ensuring that the read alignment and normalisation were compatible with our pipeline. In addition to the raw data files, the metadata tables containing the sample-wise clinical features for each dataset were also collected using the GEOquery R package. RNA-seq datasets with their GEO identifiers are reported in Table 2. As with the DNA microarray data, the samples from patients with usual UIP were included in the IPF group, as these two diseases are often used synonymously.

#### Quality control

Quality assessments were conducted on RNA-Seq datasets utilizing FastQC v0.11.7 (https://www.bioinformatics.babraham.ac.uk/projects/fastqc/). Trimming of reads for low-quality ends in addition to adapter removal was performed by Cutadapt version 3.7. In particular, the Phred score threshold for trimming was set to 20, and the minimum read length for 60 nucleotides. The adapter-clipped trimmed raw files were further quality-checked with FastQC v0.11.7.

#### Read alignment

Subsequently, RNA-seq reads were aligned against the human reference genome assembly GRCh38. The alignment was performed using the HISAT2 algorithm [16] utilising the genome indexes built for usage with HISAT2 (retrieved from https://ccb.jhu.edu/software/hisat2/manual.shtml).

#### Read counts extraction

Transcript quantification was performed by using the *featureCounts* function from the Rsubread v1.34.4 R package [17]. To accomplish this task, the Ensembl version 108 annotation was downloaded from http://www.ensembl.org and then utilised for read count extraction.

#### Low counts filtering

To exclude the transcripts with low expression levels across all samples within each dataset, the proportion test strategy was employed as implemented in the function *filtered.data* of the R package NOISeq [18].

#### Gene annotation

The Ensembl gene IDs were annotated to gene symbols using biomaRt v.2.40.1. Gene names were grouped based on unique values with dplyr 1.0.2 package. As with the DNA microarray datasets, the median of the expression value was calculated for each sample within each gene. This process resulted in a new data frame, where each row represents a unique gene, and the columns contain the computed median values for the respective expression value associated with each gene.

#### Normalisation and differential gene expression analysis

Normalisation and differential gene expression analysis of RNA-seq data was carried out by using DESeq2 1.24.0 [19]. The variable of interest in the model was the diagnosis (disease vs. healthy) and other relevant biological and technical variables were used as covariates when available and not confounded with the variable of interest. Moreover, comparisons between distinct lung locations and disease severity states as well as ILD subtypes were conducted when the data was present. The differential gene expression results underwent a similar filtering process as the microarray results, involving the consideration of adjusted p-values less than or equal to 0.01 and absolute log-fold changes greater than or equal to 0.58. The preprocessing pipeline of the RNA-seq data is illustrated in Supplementary Fig. 6.

### Gene co-expression networks

#### Dataset integration and batch effect mitigation

In order to infer co-expression networks representing each of the IPF and healthy biological systems (biopsies, BAL, fibroblasts, macrophages, epithelial cells), datasets were aggregated by selecting common genes across the platforms. Batch effect deriving from the different origin of the datasets was mitigated through the pamR package.

#### Network inference

To infer the networks from the batch-adjusted expression matrices, INfORM functions were utilised for the network inference [20]. The "*get_ranked_consensus_matrix*" function was applied to determine the correlations between genes, based on their expression levels, for both RNA-seq and microarray and for disease and healthy samples independently. The CLR algorithm was carried out for network inference and Pearson correlation was employed as the correlation method. To set a threshold on the edges to be included in the networks, the "*parse_edge_rank_matrix*" function. This function ranks the edges based on their weight (represented by Pearson correlation measures) and then systematically adds edges from the top of the rank until all the nodes within the network are connected. The networks were then converted to igraph objects using the *“get_iGraph”* function. For each network, centrality measures were calculated, including degree, betweenness, closeness, clustering coefficient and eigenvector. Subsequently, such centrality measures were aggregated through the Borda function from the TopKLists package and a global centrality-based gene rank was computed. The analytical steps of network inference is represented in Supplementary Fig. 7.

## LIMITATIONS

All the datasets included in this study encompass metadata files, reporting sample-wise technical and clinical information such as technology, platform, sampling site, gender, age or disease state. Metadata were remarkably heterogeneous across the datasets. This is a common problem when dealing with high-dimensional omics data, that hampers their integrability. In this study, the compliance of the data with FAIR principles was increased, facilitating their usage in future efforts of the scientific community. Moreover, batch effect can extensively affect the results of omics data, leading to artifacts in the results and their interpretation. This effect can derive from protocols, reagents, equipment, laboratory conditions, sample collection, storage, preparation methods, and time. Since gene expression datasets presented in this manuscript derive from different studies, it is essential to mitigate the batch effect to accurately compute differentially expressed genes and infer network relationships. Therefore, a rigorous mitigation of batch effect was carried out by considering the diverse sources of gene expression datasets as a batch variable, with the aim to integrate datasets produced with the same technology and platform to ultimately infer network relationships. PCA plots illustrating sample distribution pre- and post-batch effect mitigation are provided in supplementary materials.

## ETHICS STATEMENT

The authors have read and follow the ethical requirements for publication in Data in Brief and confirm that the current work does not involve human subjects, animal experiments, or any data collected from social media platforms.

## CRediT AUTHOR STATEMENT

**SI**: data curation, writing – original draft, writing – review & editing; **AF**: supervision, conceptualization, writing – original draft, writing – review & editing; **AS**: methodology, writing – original draft, writing – review & editing; **DG**: supervision, conceptualization, funding acquisition, writing – original draft, writing – review & editing.

## ACKNOWLEDGEMENTS

This work was supported by the Academy of Finland [grant agreement number 322761], and the European Research Council (ERC) programme, Consolidator project “ARCHIMEDES” [grant agreement number 101043848]. AF and AS were supported by Tampere Institute for Advanced Study (IAS). AF was supported by the Health Data Science (HDS) program of Tampere University.

---

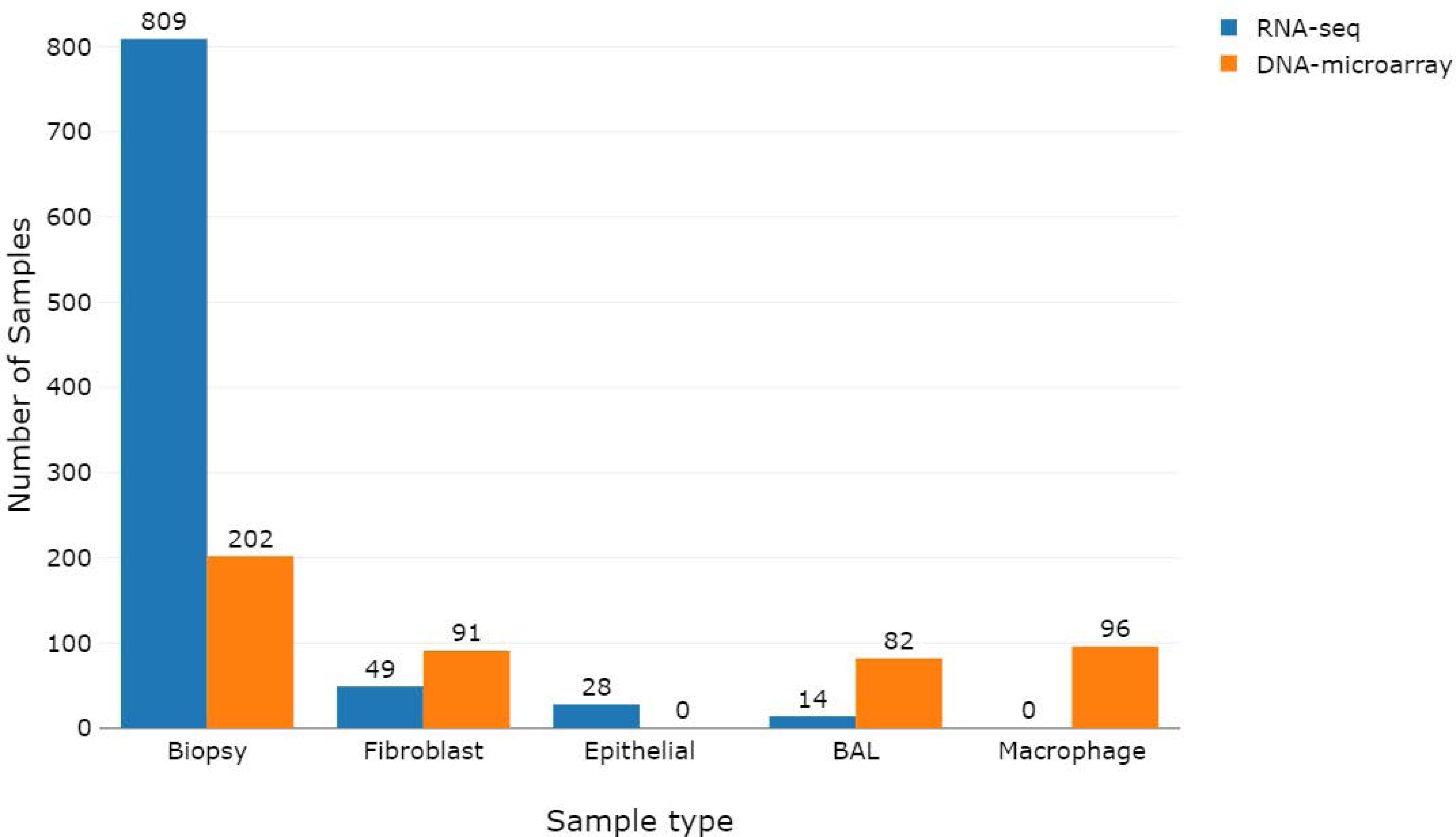

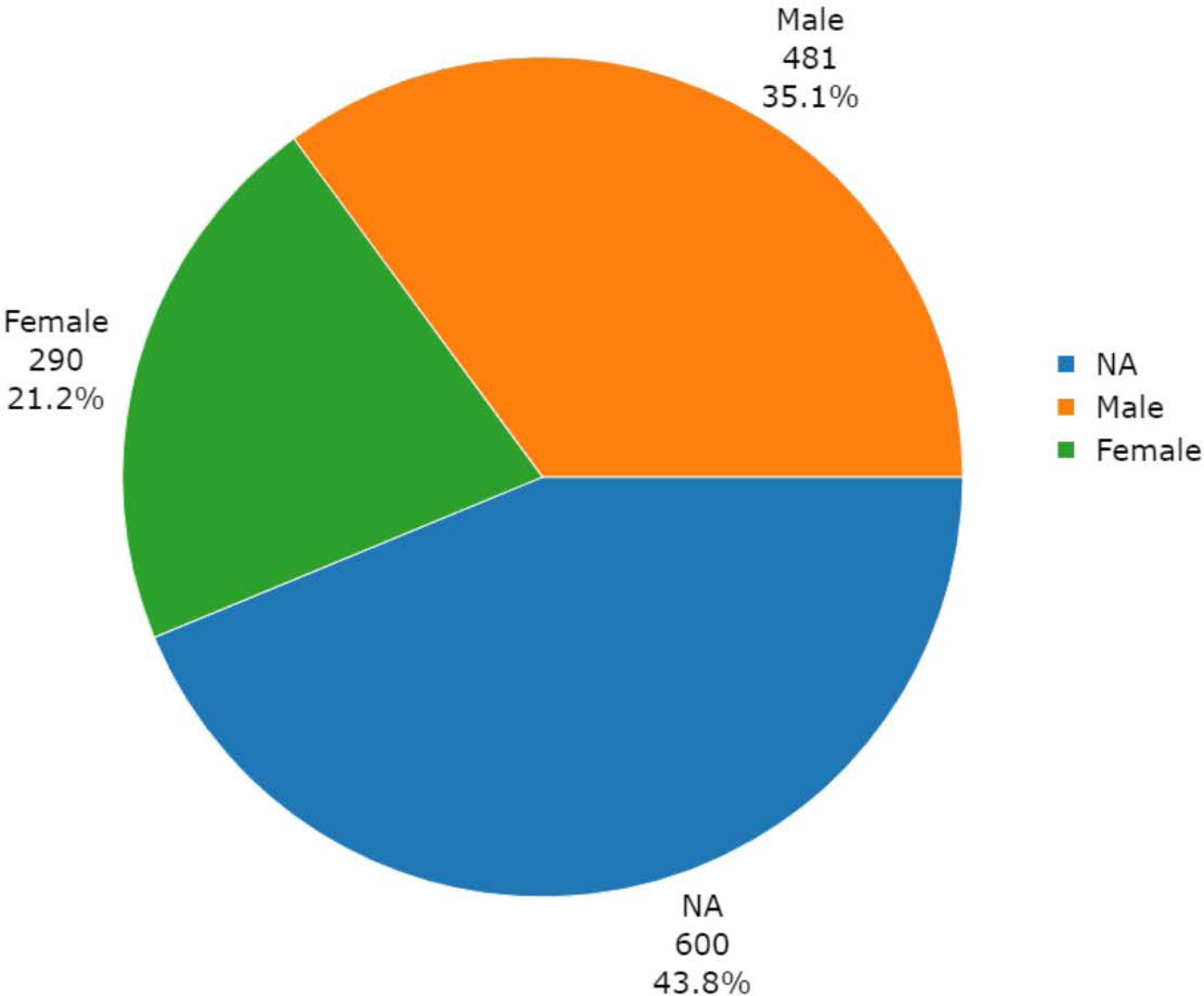

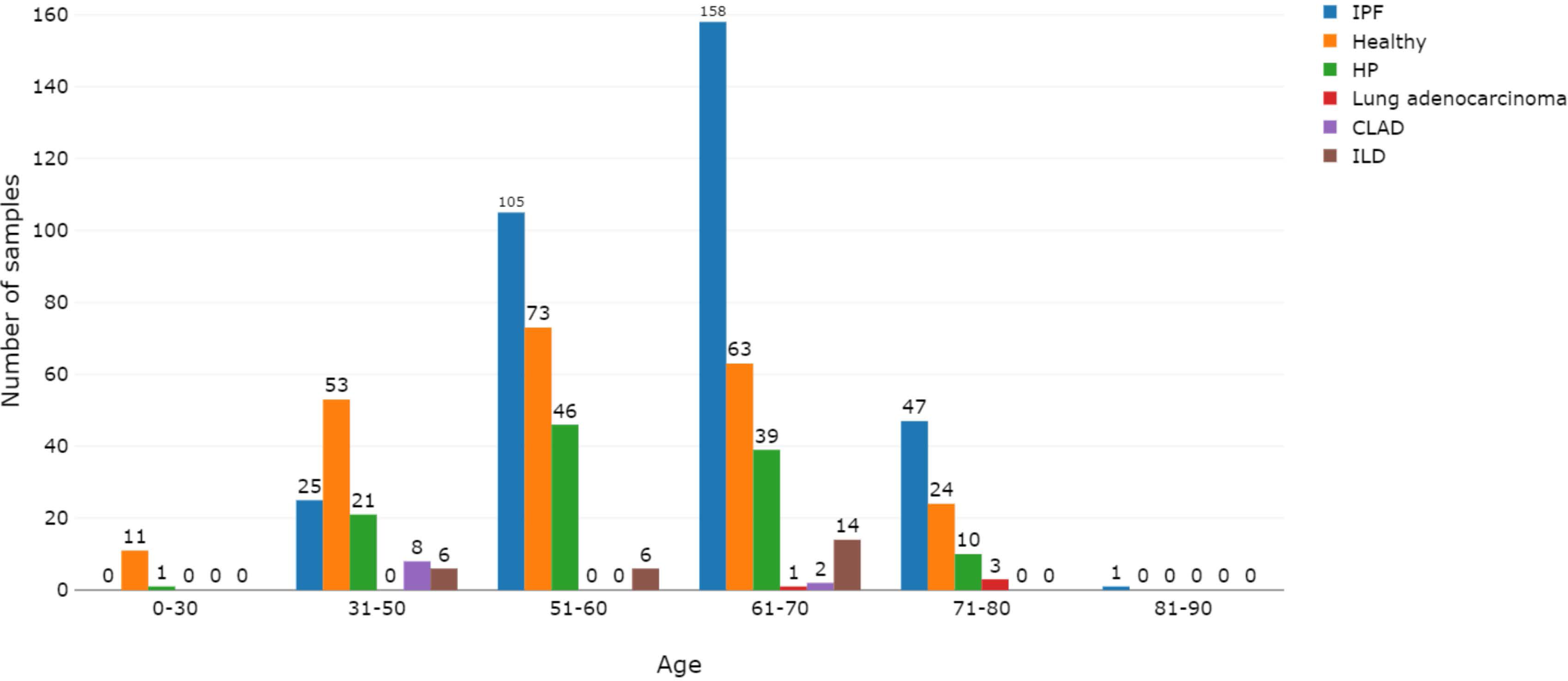

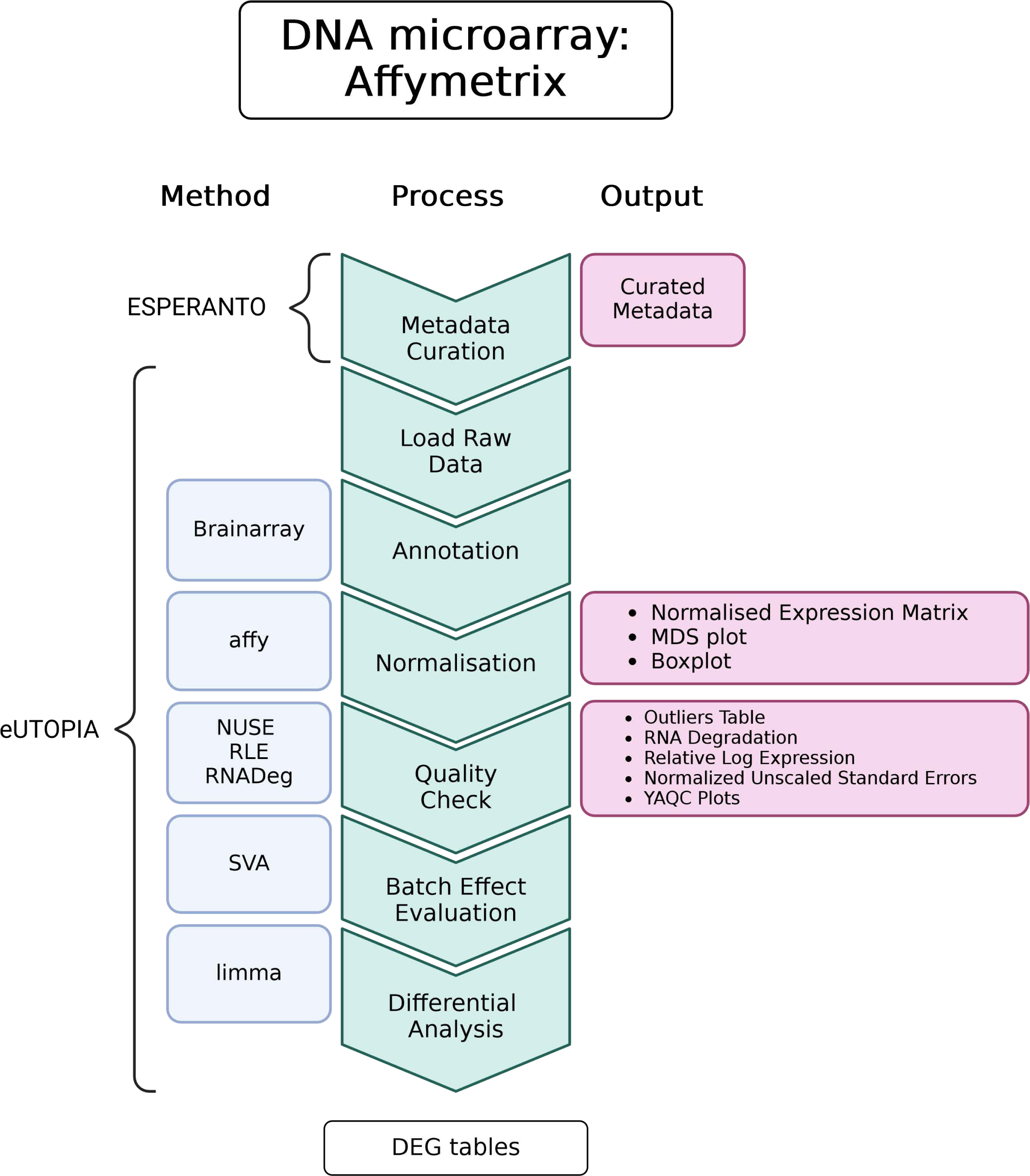

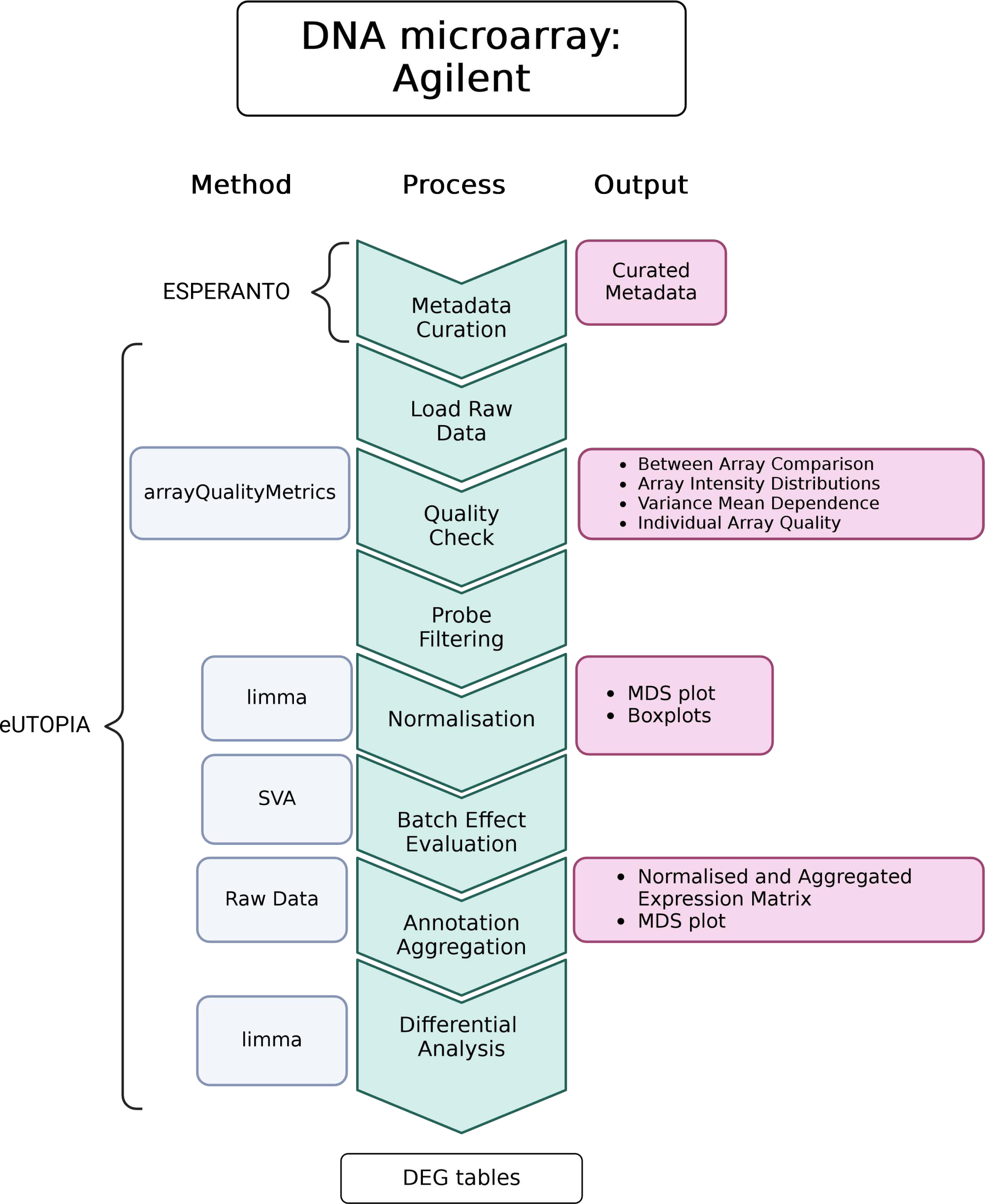

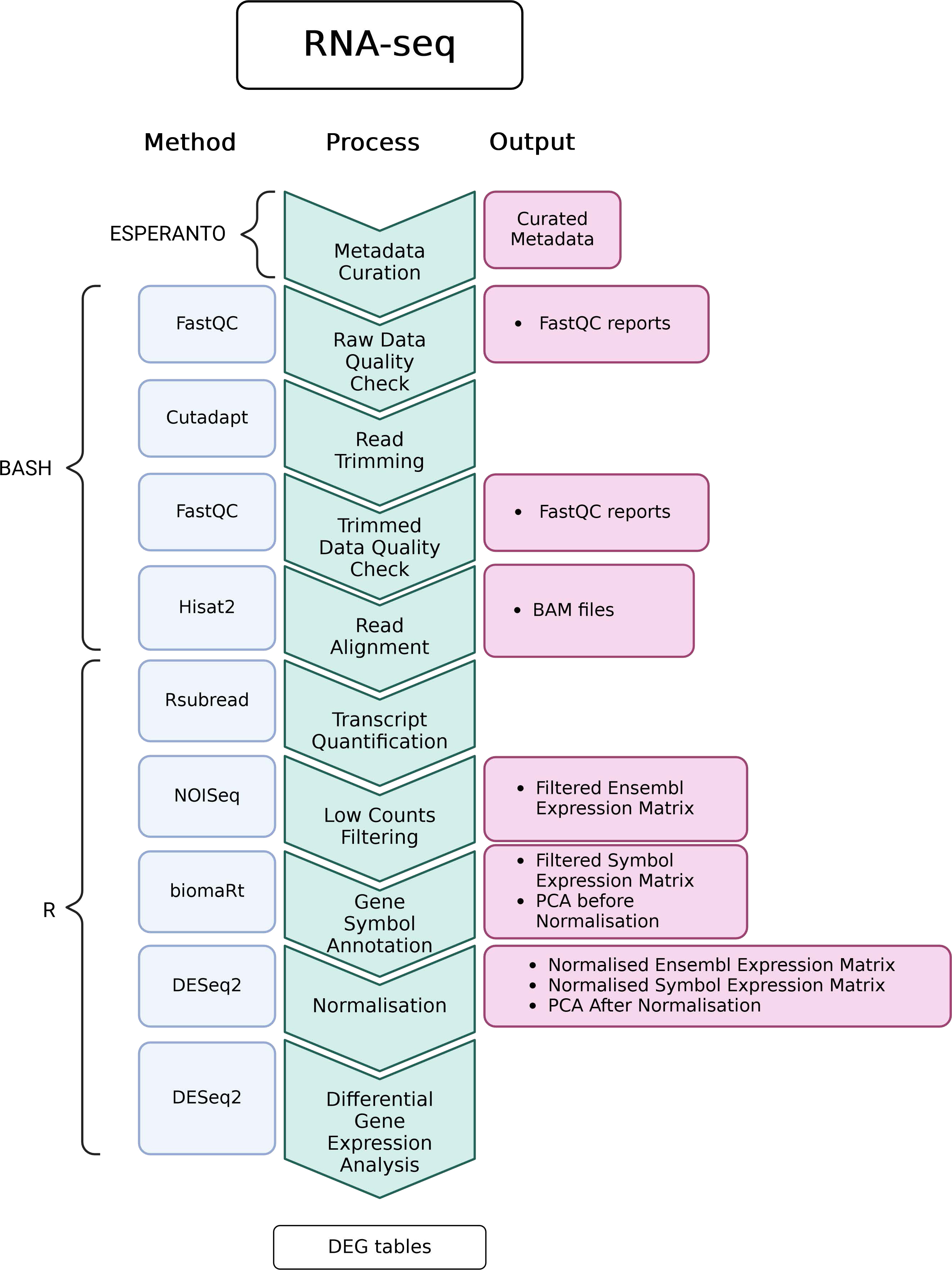

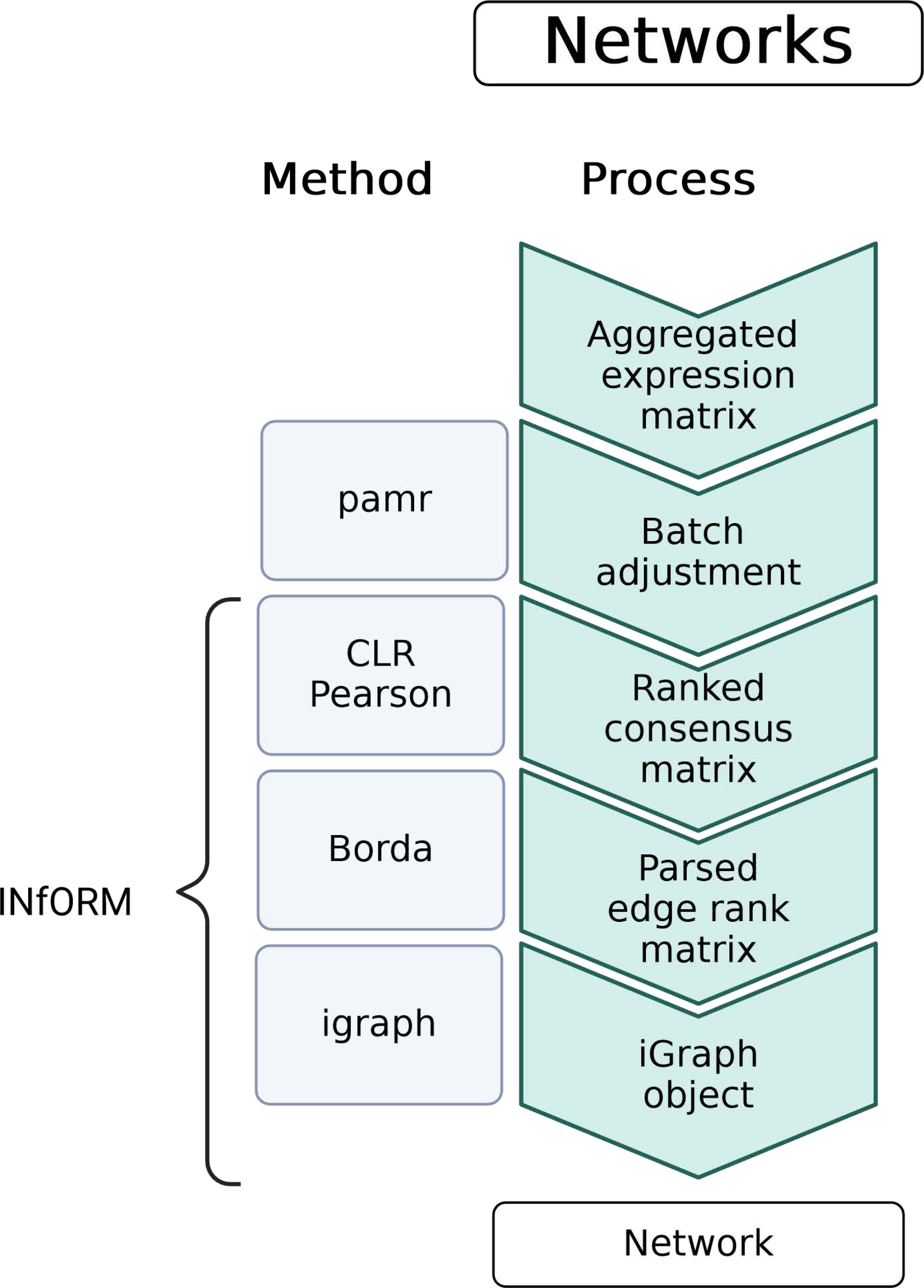

