## Supplementary material for "Curated and harmonised transcriptomics datasets of interstitial lung diseases": GSE11196_eUTOPIA_Affymetrix_QC_Report_2024-02-12.pdf

*eUTOPIA*

**5 YAQC Plots** **8**

### 1 Outliers Table

| | RLE | NUSE | DEG | SUM |
|---|---|---|---|---|
| Control_(Control128)_Contractile_Heavy_polyribosomal_RNA | 0 | 0 | 1 | 1 |
| IPF_(IPF129)_Contractile_Heavy_polyribosomal_RNA | 0 | 0 | 1 | 1 |
| IPF_(IPF75)_Contractile_Heavy_polyribosomal_RNA | 0 | 0 | 1 | 1 |
| Control_(Control128)_Non_contractile_Heavy_polyribosomal_RNA | 0 | 0 | 1 | 1 |
| Control_(Control106)_Contractile_Total_RNA | 0 | 0 | 1 | 1 |
| Control_(Control128)_Contractile_Total_RNA | 0 | 0 | 1 | 1 |
| IPF_(IPF75)_Contractile_Total_RNA | 0 | 0 | 1 | 1 |
| Control_(Control106)_Non_contractile_Total_RNA | 0 | 0 | 1 | 1 |
| Control_(Control128)_Non_contractile_Total_RNA | 0 | 0 | 1 | 1 |
| Control_(Control54)_Non_contractile_Total_RNA | 0 | 0 | 1 | 1 |
| IPF_(IPF129)_Non_contractile_Total_RNA | 0 | 0 | 1 | 1 |
| IPF_(IPF75)_Non_contractile_Total_RNA | 0 | 0 | 1 | 1 |
| IPF_(IPF12)_Contractile_Heavy_polyribosomal_RNA | 1 | 0 | 0 | 1 |
| IPF_(IPF14)_Contractile_Heavy_polyribosomal_RNA | 1 | 0 | 0 | 1 |
| Control_(Control54)_Non_contractile_Heavy_polyribosomal_RNA | 1 | 0 | 0 | 1 |
| IPF_(IPF12)_Non_contractile_Heavy_polyribosomal_RNA | 0 | 1 | 0 | 1 |
| Control_(Control89)_Contractile_Total_RNA | 0 | 1 | 0 | 1 |
| IPF_(IPF12)_Contractile_Total_RNA | 0 | 1 | 0 | 1 |

#### 1.1 Outliers (All Methods)

| Outliers overall |
| --- |
| NA |

##### 1.2 Outliers (At Least One Method)

| Outliers at least 1 |
| --- |
| Control_(Control128)_Contractile_Heavy_polyribosomal_RNA |
| IPF_(IPF129)_Contractile_Heavy_polyribosomal_RNA |
| IPF_(IPF75)_Contractile_Heavy_polyribosomal_RNA |
| Control_(Control128)_Non_contractile_Heavy_polyribosomal_RNA |
| Control_(Control106)_Contractile_Total_RNA |
| Control_(Control128)_Contractile_Total_RNA |
| IPF_(IPF75)_Contractile_Total_RNA |
| Control_(Control106)_Non_contractile_Total_RNA |
| Control_(Control128)_Non_contractile_Total_RNA |
| Control_(Control54)_Non_contractile_Total_RNA |
| IPF_(IPF129)_Non_contractile_Total_RNA |
| IPF_(IPF75)_Non_contractile_Total_RNA |
| IPF_(IPF12)_Contractile_Heavy_polyribosomal_RNA |
| IPF_(IPF14)_Contractile_Heavy_polyribosomal_RNA |
| Control_(Control54)_Non_contractile_Heavy_polyribosomal_RNA |
| IPF_(IPF12)_Non_contractile_Heavy_polyribosomal_RNA |
| Control_(Control89)_Contractile_Total_RNA |
| IPF_(IPF12)_Contractile_Total_RNA |

#### 2 RNA Degradation

##### 2.1 Summarized Mean QC

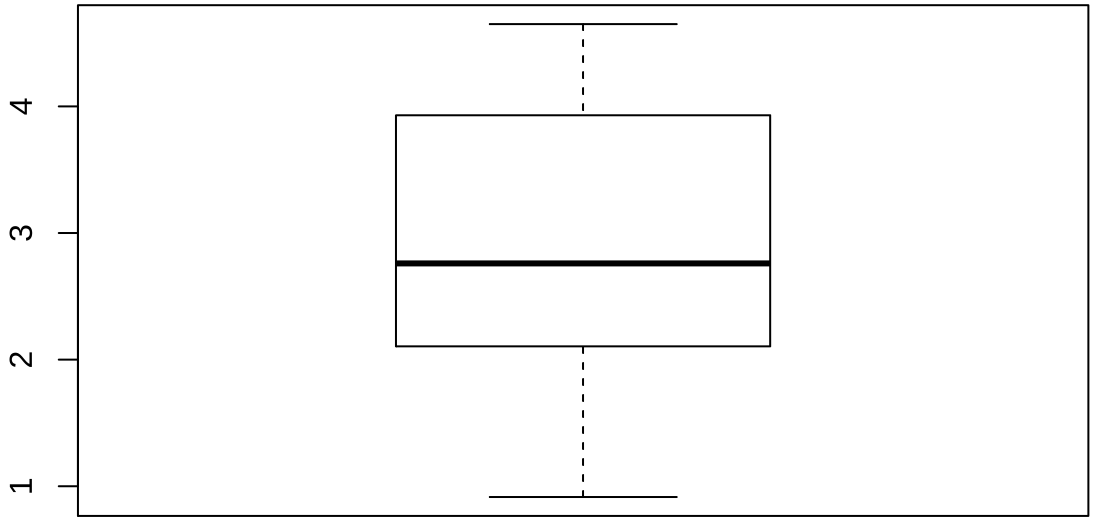

#### 2.2 Discrete QC Plots

Sample Group [1]

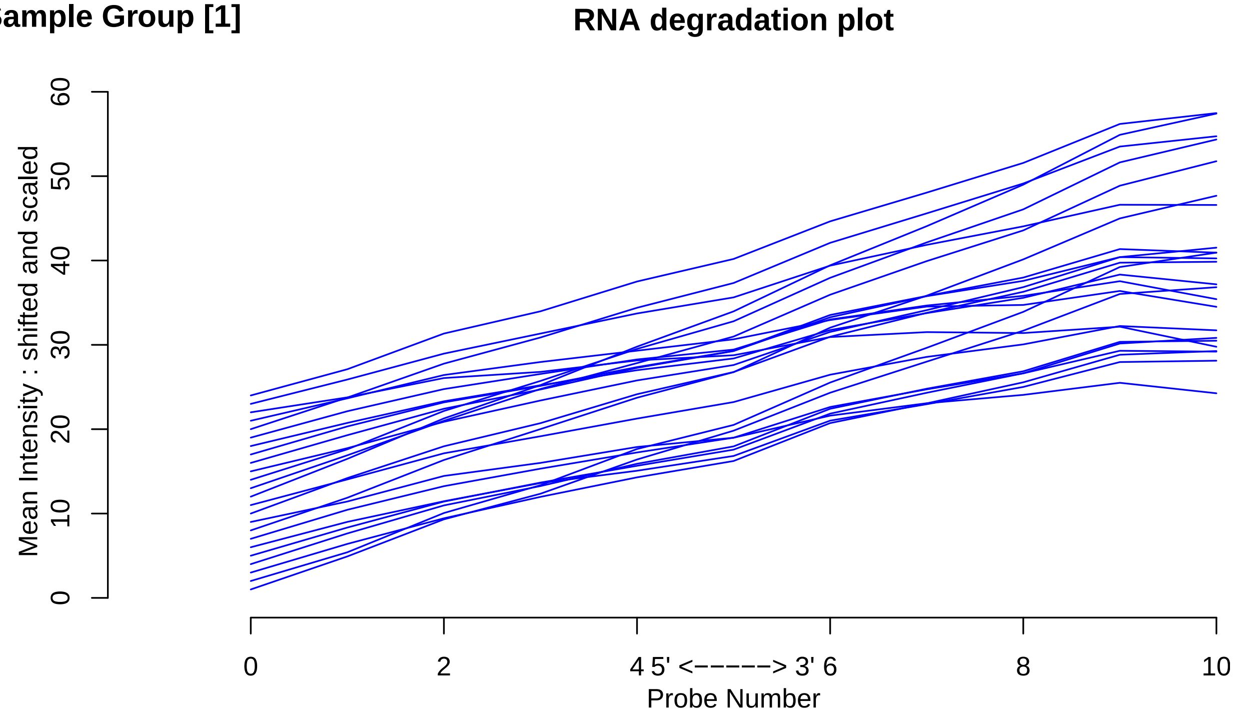

Sample Group [2]

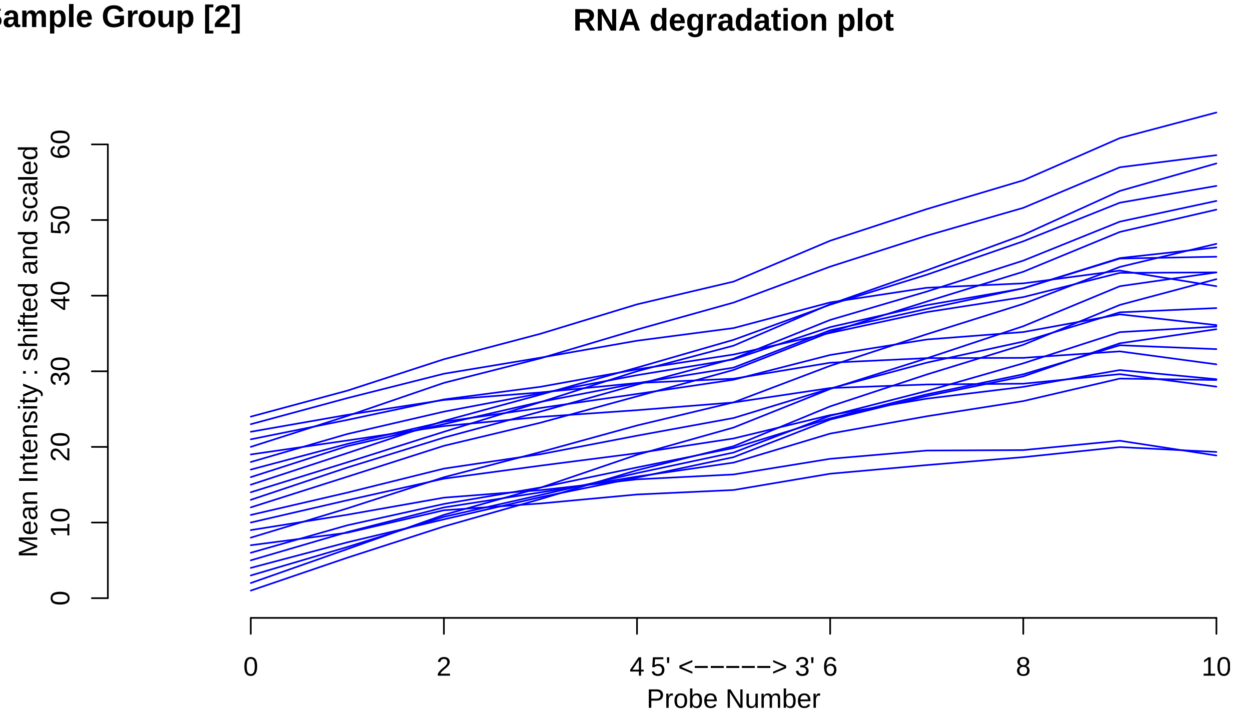

### 3 Relative Log Expression

#### 3.1 Summarized Median QC

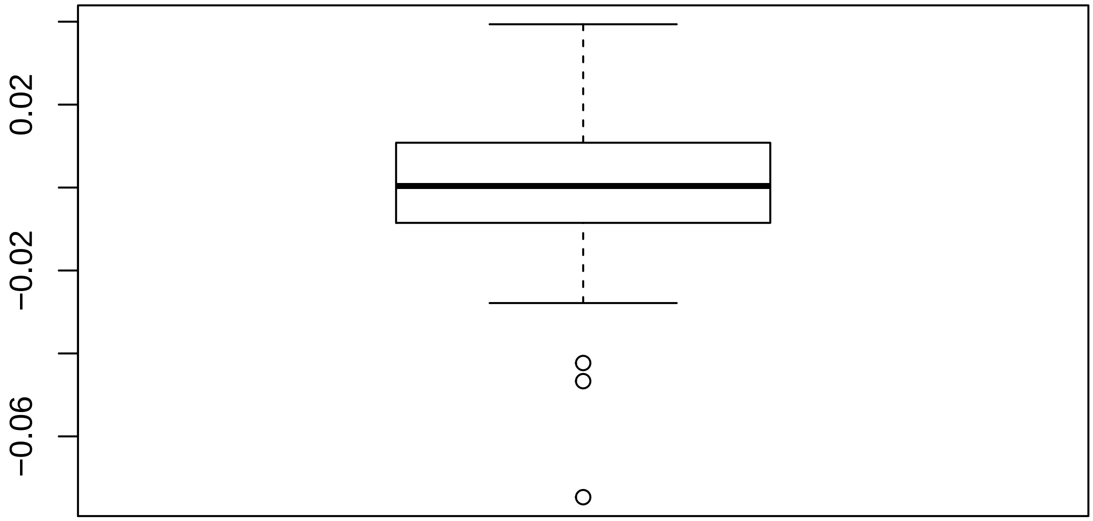

#### 3.2 Discrete QC Plots

Sample Group [1]

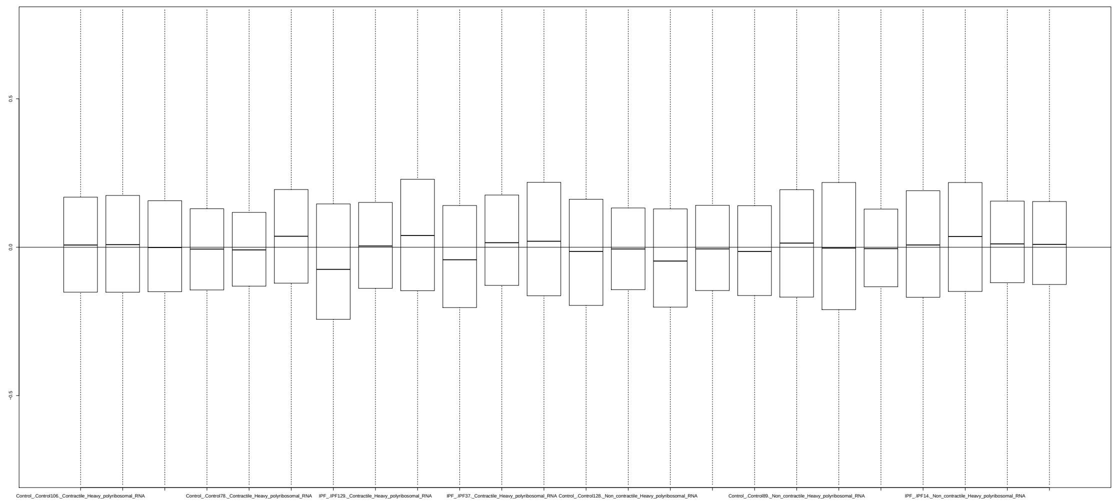

Sample Group [2]

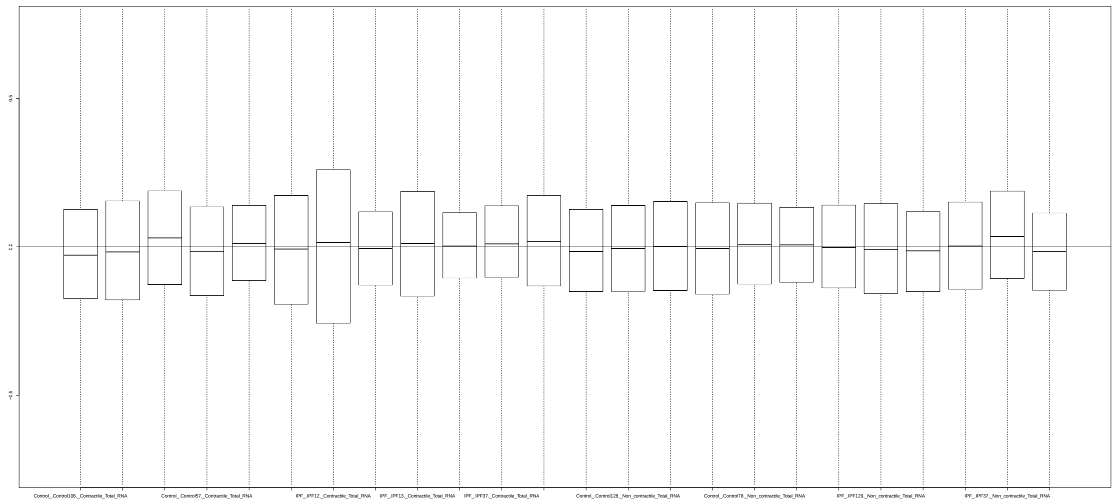

### 4 Normalized Unscaled Standard Errors

#### 4.1 Summarized Median QC

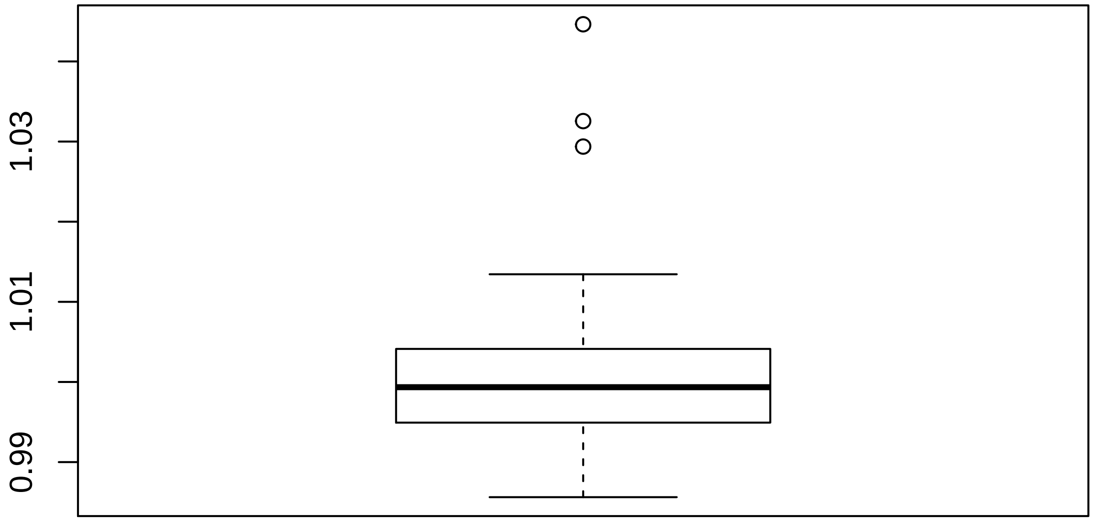

#### 4.2 Discrete QC Plots

Sample Group [1]

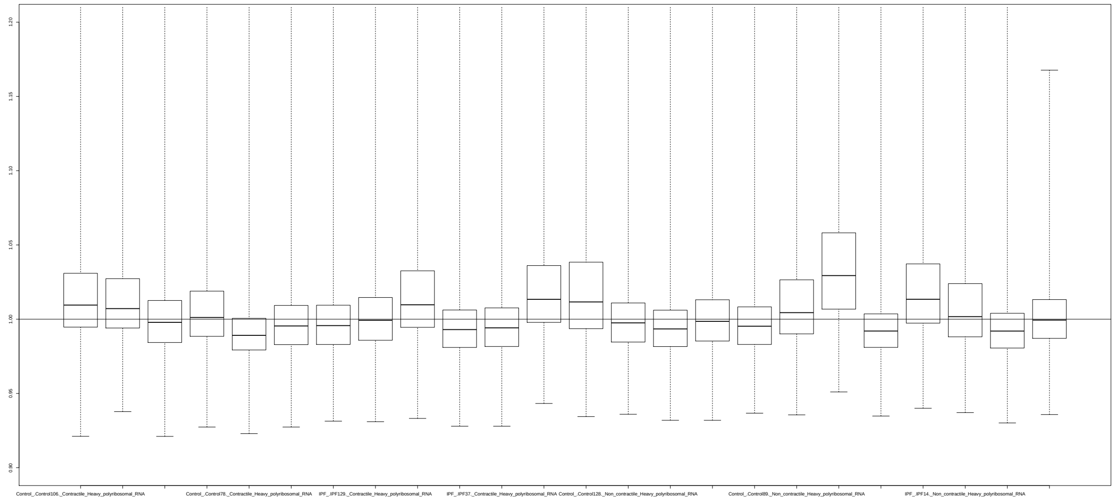

Sample Group [2]

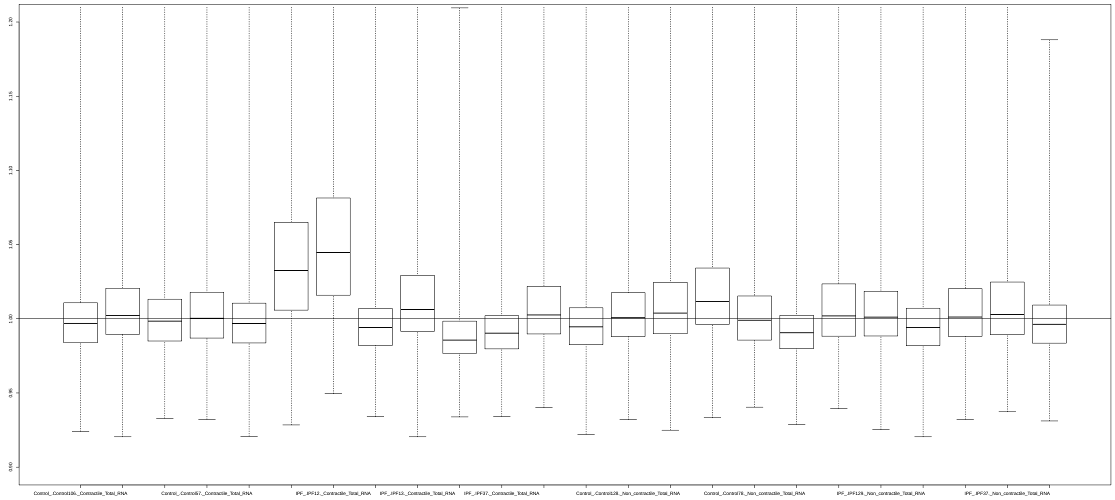

### 5 YAQC Plots

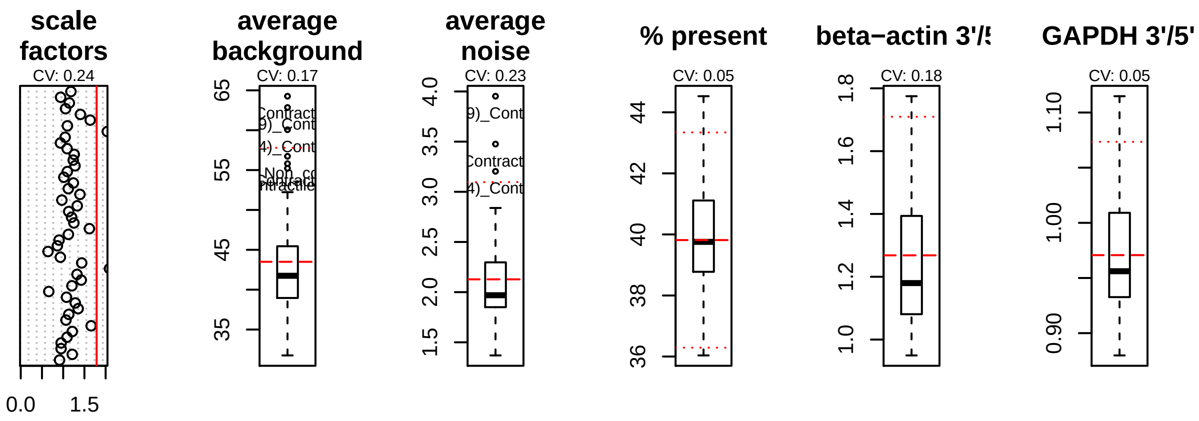

scale factors
CV: 0.24
0.0
1.5
average background
CV: 0.17
65
55
45
35
average noise
CV: 0.23
4.0
3.5
3.0
2.5
2.0
1.5
% present
CV: 0.05
44
42
40
38
36
beta−actin 3'/5'
CV: 0.18
1.8
1.6
1.4
1.2
1.0
GAPDH 3'/5'
CV: 0.05
1.10
1.00
0.90

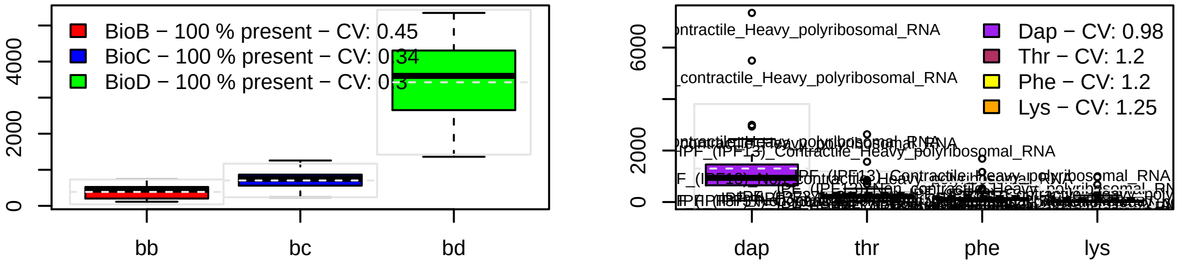

BioB − 100 % present − CV: 0.45
BioC − 100 % present − CV: 0.34
BioD − 100 % present − CV: 0.3
4000
2000
0
bb
bc
bd
Dap − CV: 0.98
Thr − CV: 1.2
Phe − CV: 1.2
Lys − CV: 1.25
6000
2000
0
dap
thr
phe
lys
