## Supplementary material for "Curated and harmonised transcriptomics datasets of interstitial lung diseases": GSE21369_eUTOPIA_Affymetrix_QC_Report_2024-02-09.pdf

*eUTOPIA*

**5 YAQC Plots** **8**

### 1 Outliers Table

| | RLE | NUSE | DEG | SUM |
|---|---|---|---|---|
| ILD_4 | 0 | 0 | 1 | 1 |
| ILD_5 | 0 | 0 | 1 | 1 |
| ILD_13 | 0 | 0 | 1 | 1 |
| ILD_16 | 0 | 0 | 1 | 1 |
| ILD_17 | 0 | 0 | 1 | 1 |
| ILD_18 | 0 | 0 | 1 | 1 |
| ILD_23 | 0 | 0 | 1 | 1 |
| Control_5 | 1 | 0 | 0 | 1 |
| ILD_1 | 1 | 1 | 0 | 2 |
| ILD_2 | 1 | 0 | 0 | 1 |
| ILD_7 | 1 | 1 | 0 | 2 |
| ILD_9 | 1 | 1 | 0 | 2 |

#### 1.1 Outliers (All Methods)

| Outliers overall |
|---|
| ILD_1 |
| ILD_7 |
| ILD_9 |

#### 1.2 Outliers (At Least One Method)

| Outliers at least 1 |
|---|
| ILD_4 |
| ILD_5 |
| ILD_13 |
| ILD_16 |
| ILD_17 |
| ILD_18 |
| ILD_23 |
| Control_5 |
| ILD_1 |
| ILD_2 |
| ILD_7 |
| ILD_9 |

#### 2 RNA Degradation

##### 2.1 Summarized Mean QC

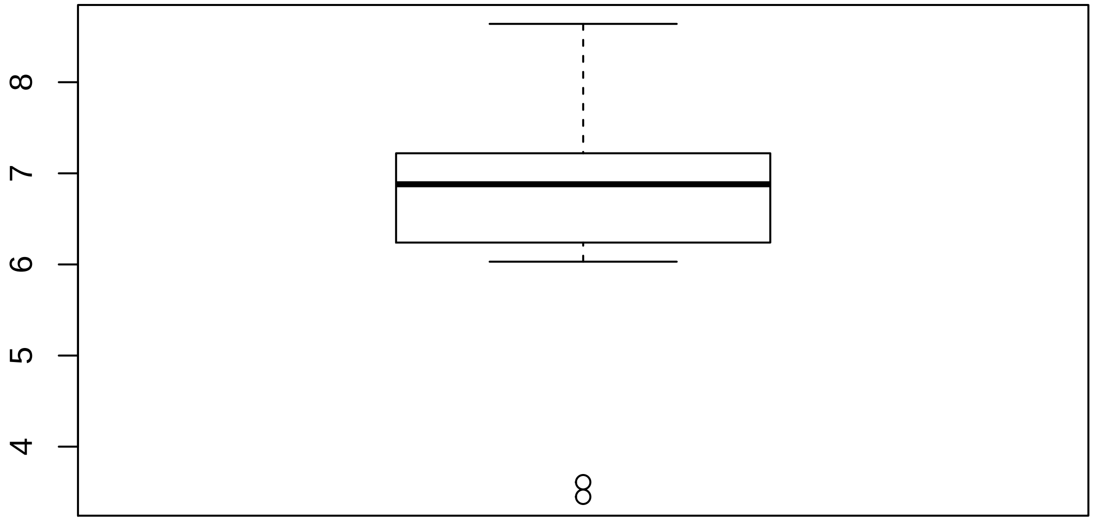

#### 2.2 Discrete QC Plots

**Sample Group [1]**

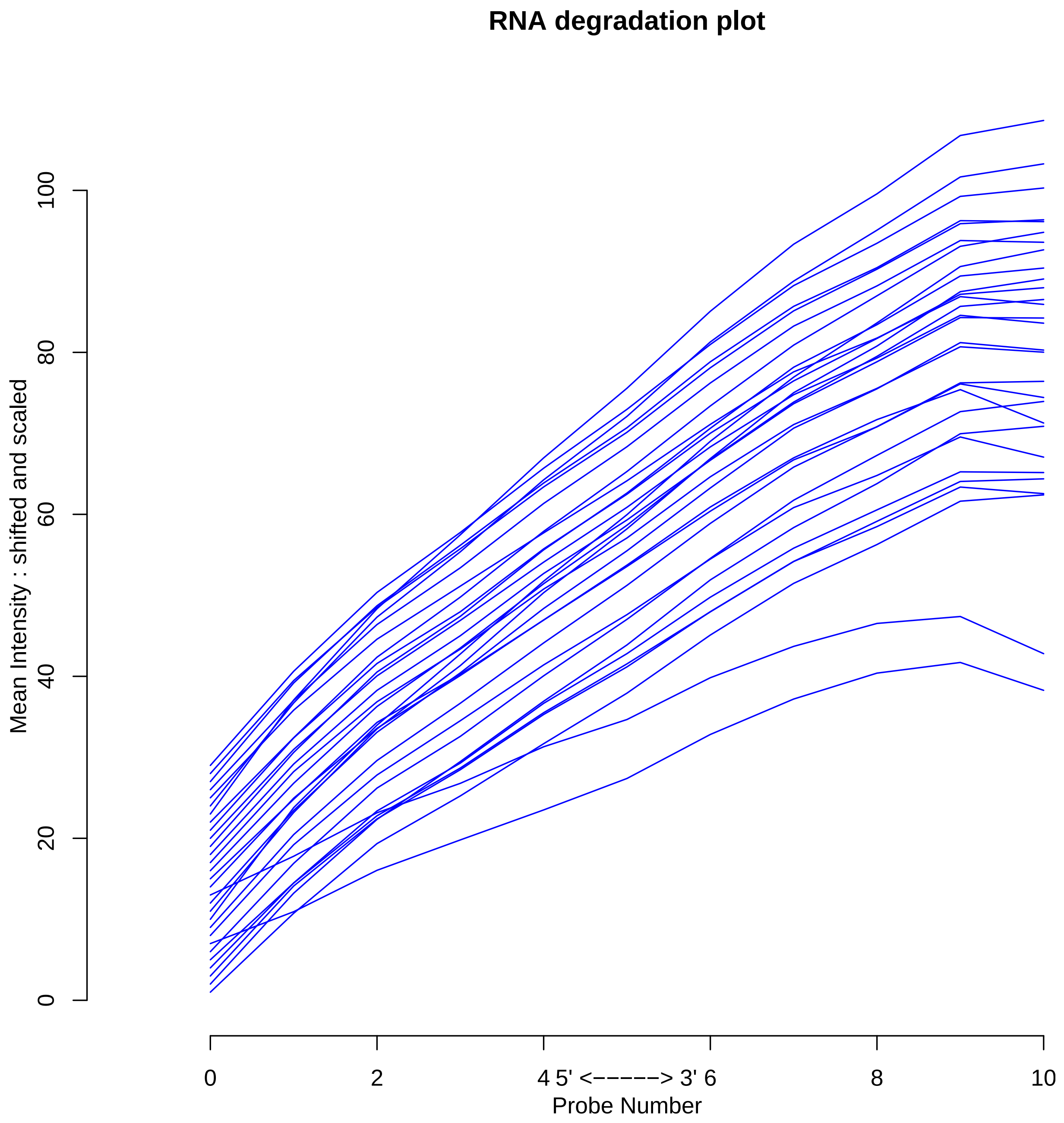

### 3 Relative Log Expression

#### 3.1 Summarized Median QC

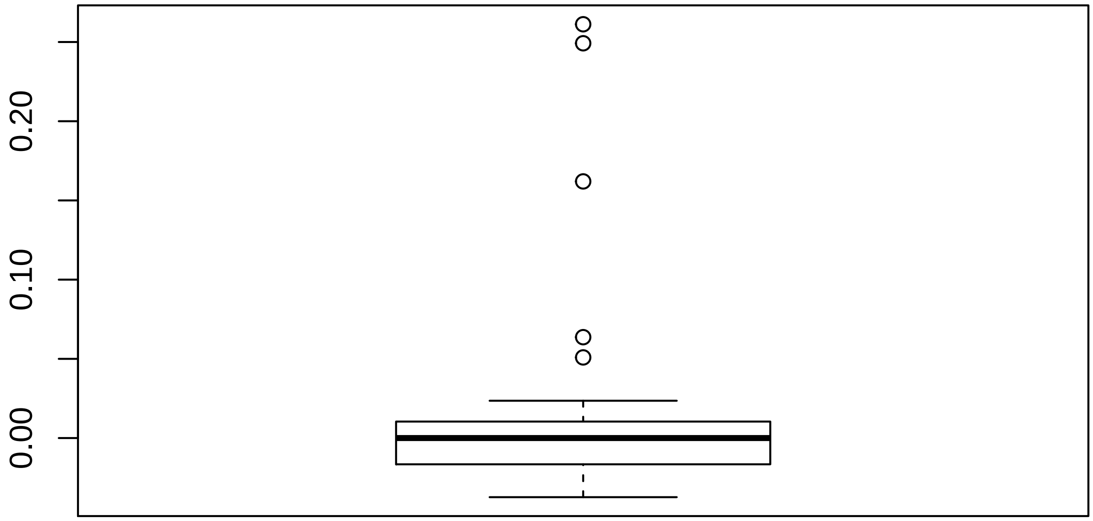

#### 3.2 Discrete QC Plots

Sample Group [1]

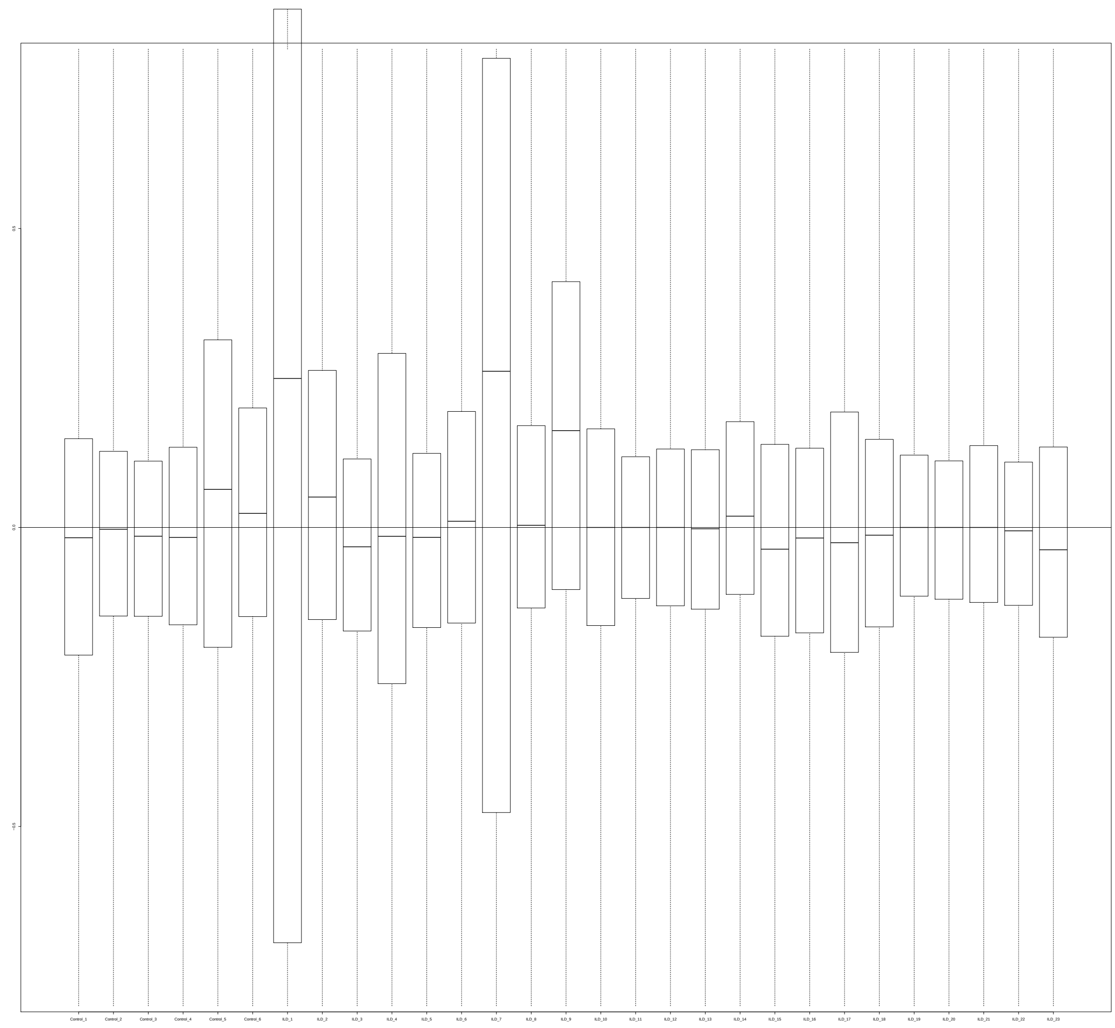

### 4 Normalized Unscaled Standard Errors

#### 4.1 Summarized Median QC

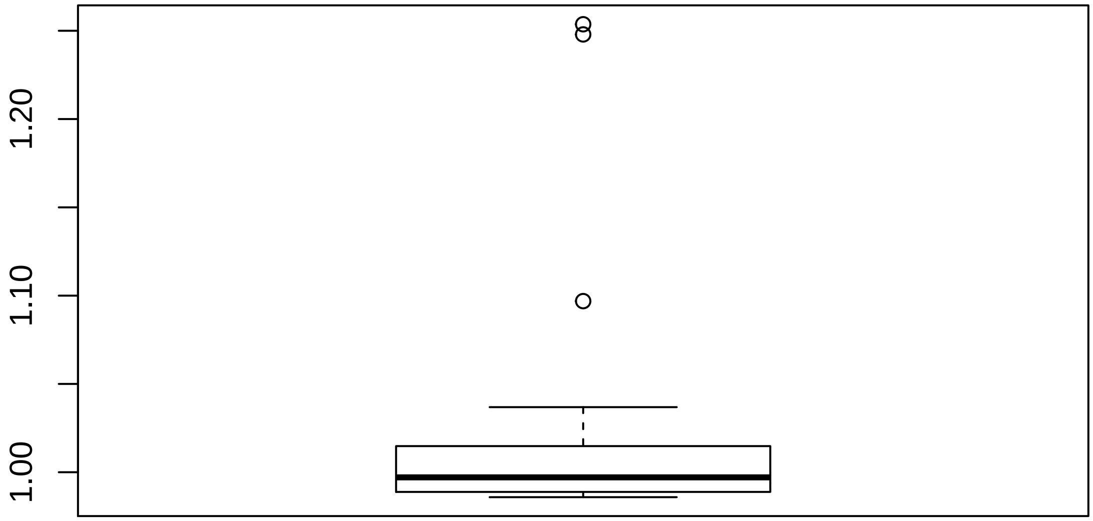

#### 4.2 Discrete QC Plots

Sample Group [1]

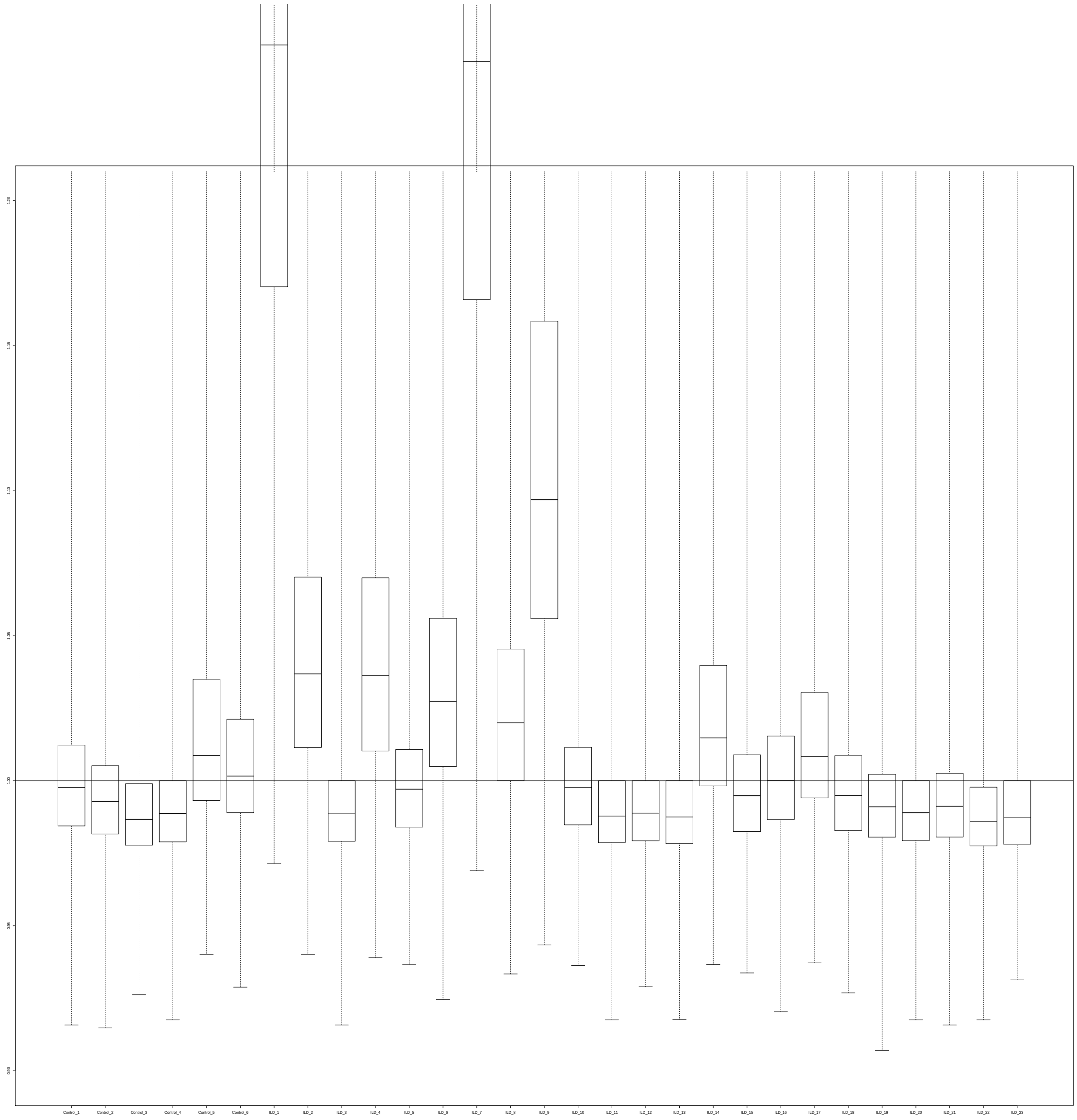

### 5 YAQC Plots

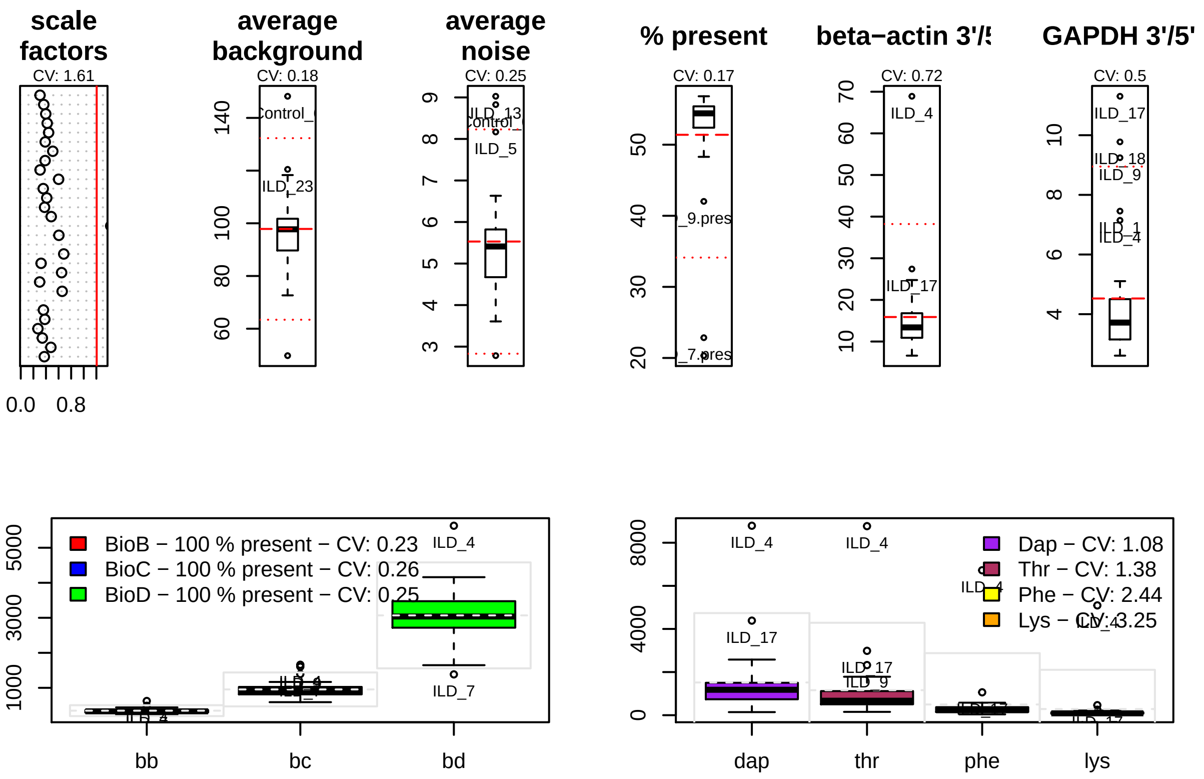
scale factors
CV: 1.61
0.0
0.8
average background
CV: 0.18
average noise
CV: 0.25
% present
CV: 0.17
beta−actin 3'/5
CV: 0.72
GAPDH 3'/5'
CV: 0.5
BioB − 100 % present − CV: 0.23
BioC − 100 % present − CV: 0.26
BioD − 100 % present − CV: 0.25
bb
bc
bd
Dap − CV: 1.08
Thr − CV: 1.38
Phe − CV: 2.44
Lys − CV: 3.25
dap
thr
phe
lys
