## Supplementary material for "Curated and harmonised transcriptomics datasets of interstitial lung diseases": GSE24206_eUTOPIA_Affymetrix_QC_Report_2024-02-12.pdf

*eUTOPIA*

**5 YAQC Plots** **8**

### 1 Outliers Table

| | RLE | NUSE | DEG | SUM |
|---|---|---|---|---|
| Early__IPF__surgical__biopsy__upper__lobe__rep140 | 1 | 0 | 1 | 2 |
| Early__IPF__surgical__biopsy__upper__lobe__rep142 | 0 | 0 | 1 | 1 |
| Early__IPF__surgical__biopsy__upper__lobe__rep144 | 0 | 0 | 1 | 1 |
| Early__IPF__surgical__biopsy__upper__lobe__rep145 | 0 | 0 | 1 | 1 |
| Advanced__IPF__explant__lower__lobe__rep146 | 0 | 0 | 1 | 1 |
| Early__IPF__surgical__biopsy__lower__lobe__rep149 | 0 | 0 | 1 | 1 |
| Healthy__donor__biological__replicate2 | 1 | 0 | 0 | 1 |
| Healthy__donor__biological__replicate4 | 1 | 1 | 0 | 2 |
| Healthy__donor__biological__replicate5 | 1 | 1 | 0 | 2 |
| Healthy__donor__biological__replicate6 | 1 | 1 | 0 | 2 |
| Advanced__IPF__explant__lower__lobe__rep152 | 1 | 1 | 0 | 2 |
| Advanced__IPF__explant__lower__lobe__rep158 | 1 | 0 | 0 | 1 |

#### 1.1 Outliers (All Methods)

| Outliers overall |
|---|
| Early__IPF__surgical__biopsy__upper__lobe__rep140 |
| Healthy__donor__biological__replicate4 |
| Healthy__donor__biological__replicate5 |
| Healthy__donor__biological__replicate6 |
| Advanced__IPF__explant__lower__lobe__rep152 |

##### 1.2 Outliers (At Least One Method)

| Outliers at least 1 |
|---|
| Early_IPF_surgical_biopsy_upper_lobe_rep140 |
| Early_IPF_surgical_biopsy_upper_lobe_rep142 |
| Early_IPF_surgical_biopsy_upper_lobe_rep144 |
| Early_IPF_surgical_biopsy_upper_lobe_rep145 |
| Advanced_IPF_explant_lower_lobe_rep146 |
| Early_IPF_surgical_biopsy_lower_lobe_rep149 |
| Healthy_donor_biological_replicate2 |
| Healthy_donor_biological_replicate4 |
| Healthy_donor_biological_replicate5 |
| Healthy_donor_biological_replicate6 |
| Advanced_IPF_explant_lower_lobe_rep152 |
| Advanced_IPF_explant_lower_lobe_rep158 |

#### 2 RNA Degradation

##### 2.1 Summarized Mean QC

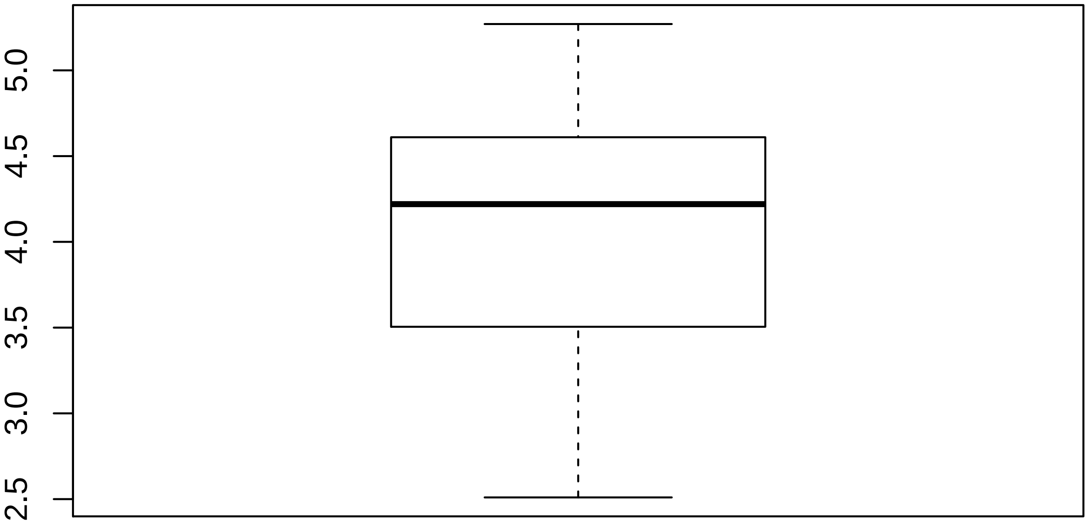

#### 2.2 Discrete QC Plots

**Sample Group [1]**

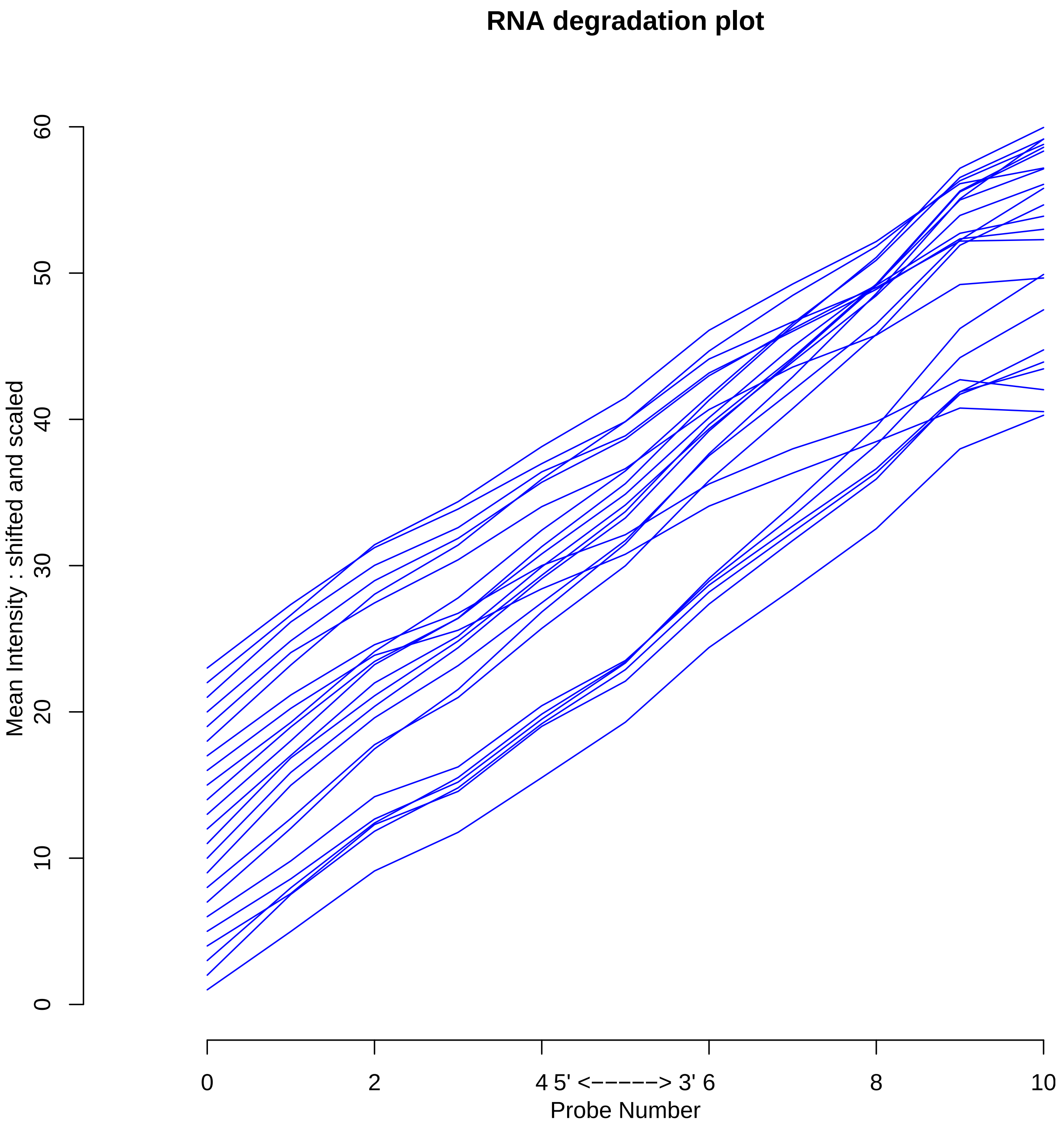

### 3 Relative Log Expression

#### 3.1 Summarized Median QC

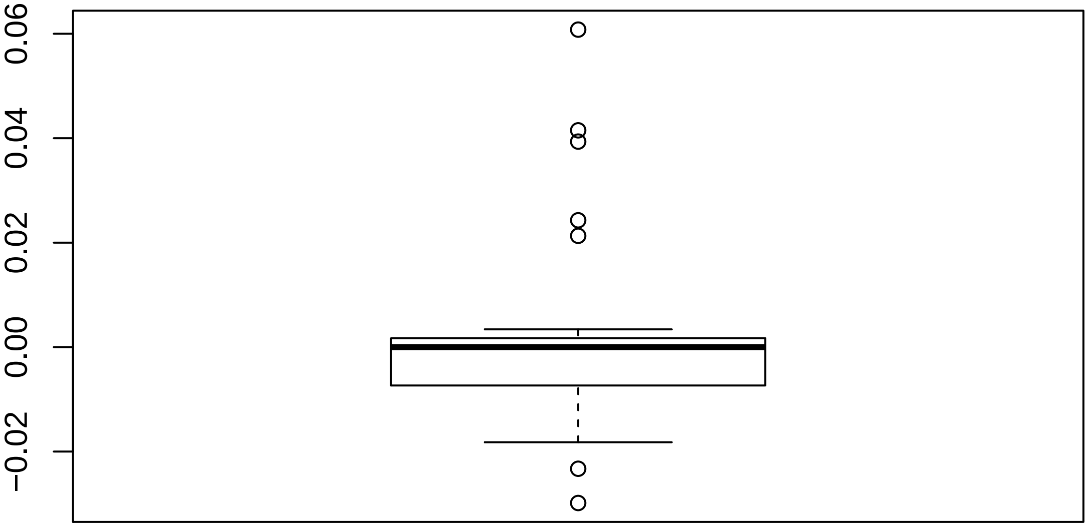

#### 3.2 Discrete QC Plots

**Sample Group [1]**

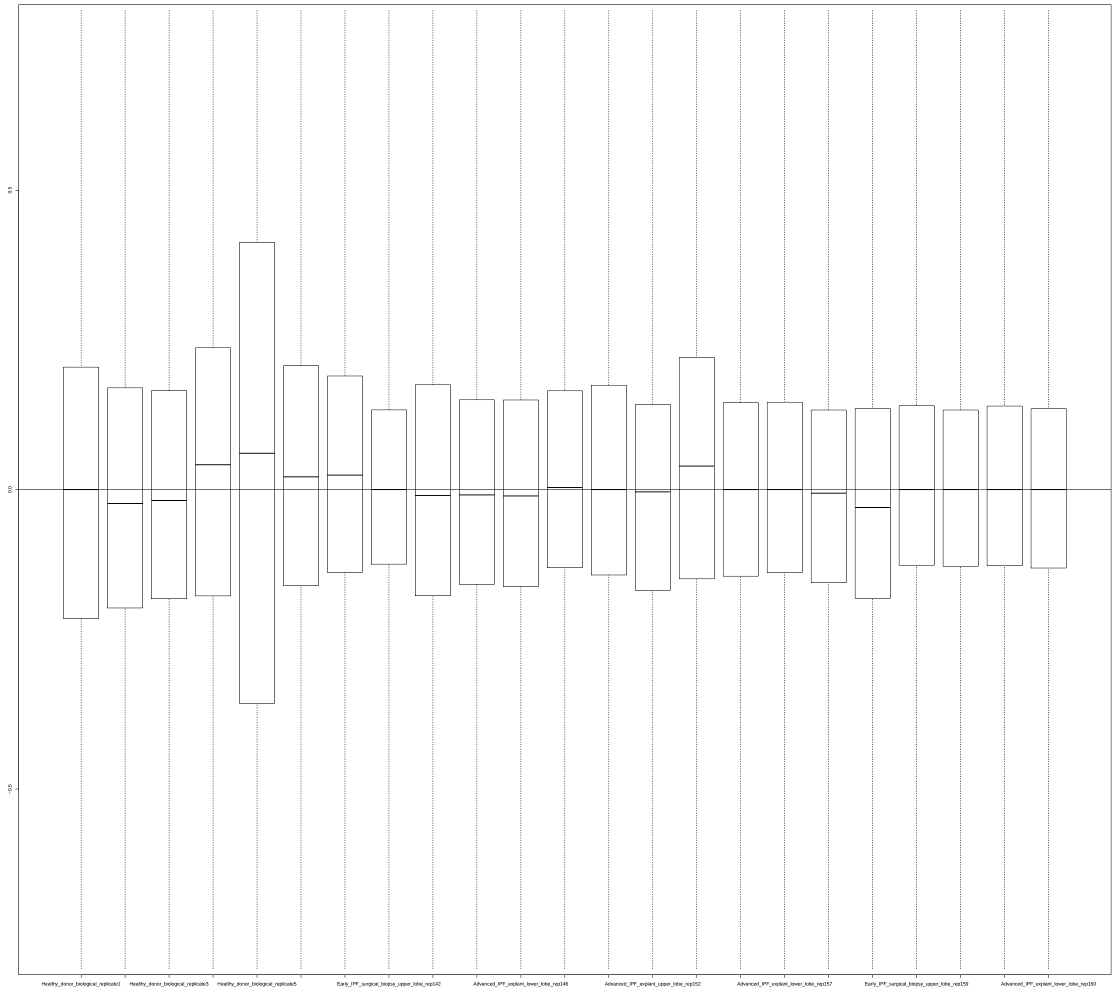

#### 4 Normalized Unscaled Standard Errors

##### 4.1 Summarized Median QC

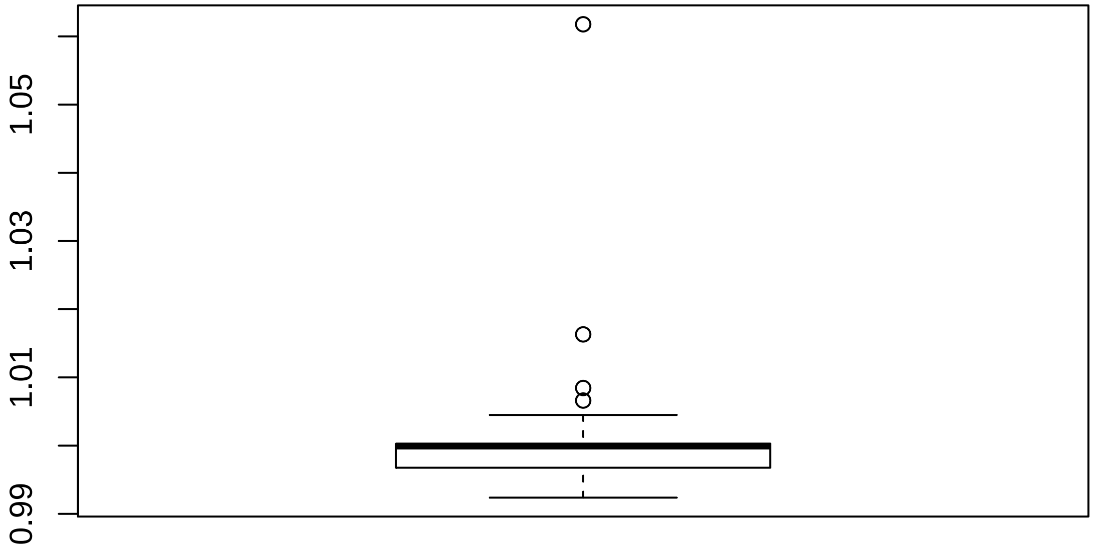

#### 4.2 Discrete QC Plots

Sample Group [1]

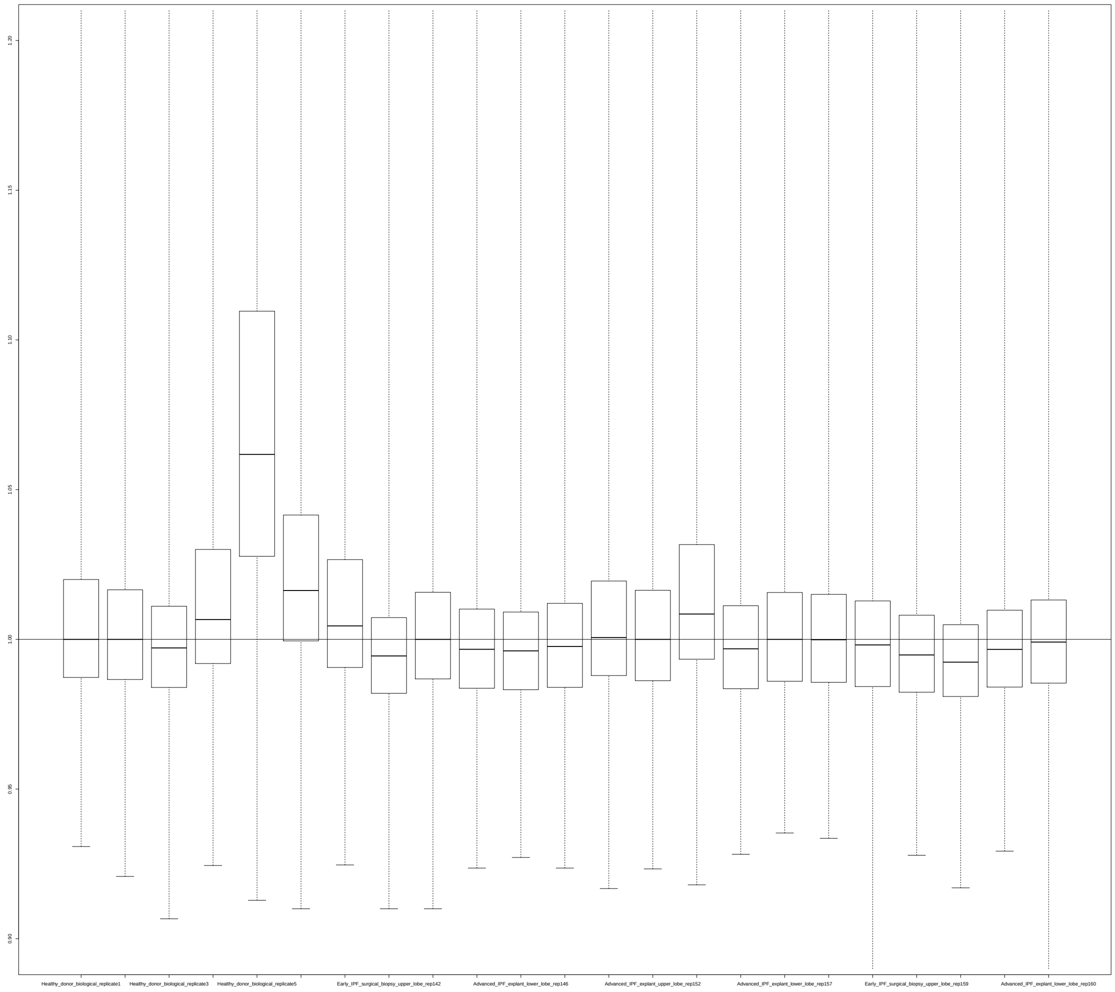

#### 5 YAQC Plots

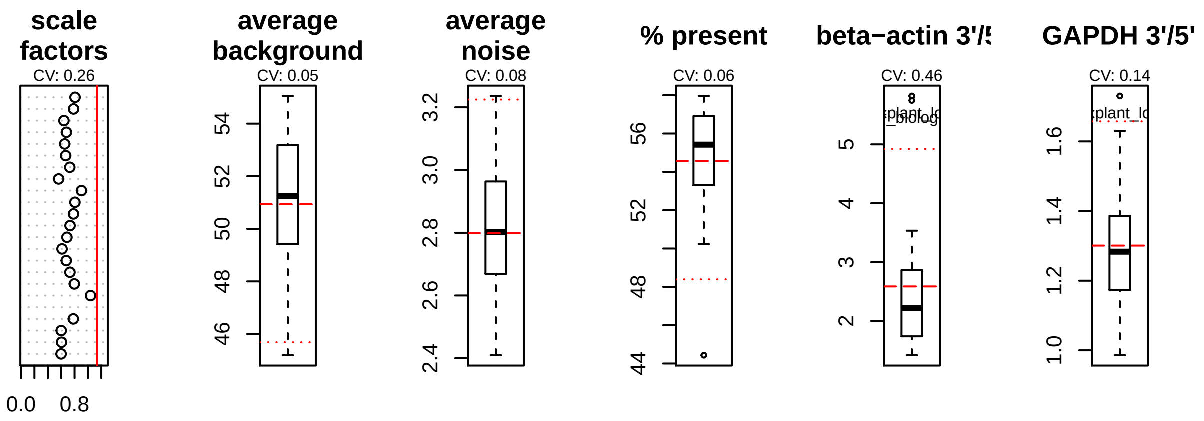
scale factors
CV: 0.26
0.0
0.8
average background
CV: 0.05
average noise
CV: 0.08
% present
CV: 0.06
beta−actin 3'/5'
CV: 0.46
GAPDH 3'/5'
CV: 0.14

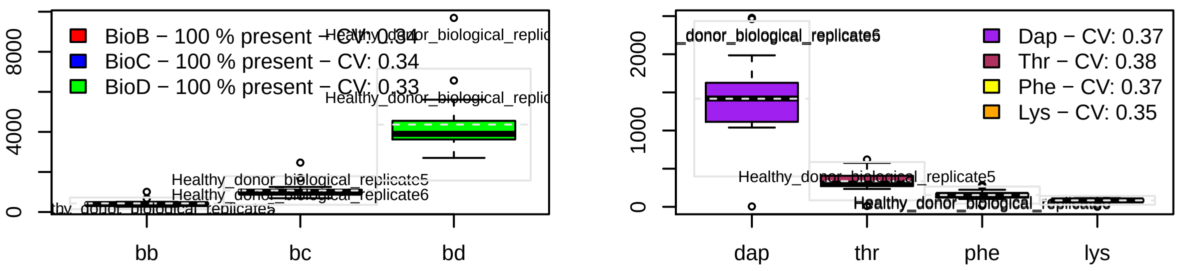
BioB − 100 % present − CV: 0.34
BioC − 100 % present − CV: 0.34
BioD − 100 % present − CV: 0.33
Healthy_donor_biological_replicate5
Healthy_donor_biological_replicate6
bb
bc
bd
Dap − CV: 0.37
Thr − CV: 0.38
Phe − CV: 0.37
Lys − CV: 0.35
dap
thr
phe
lys
