## Supplementary material for "Curated and harmonised transcriptomics datasets of interstitial lung diseases": GSE40839_eUTOPIA_Affymetrix_QC_Report_2024-02-12.pdf

*eUTOPIA*

**5 YAQC Plots** **8**

### 1 Outliers Table

| | RLE | NUSE | DEG | SU |
|---|---|---|---|---|
| Pulmonary_fibroblasts_control_individual___biological_rep4 | 1 | 0 | 1 | |
| Pulmonary_fibroblasts_control_individual___biological_rep6 | 1 | 0 | 1 | |
| Pulmonary_fibroblasts_scleroderma_associated_interstitial_lung_disease___biological_rep7 | 0 | 0 | 1 | |
| Pulmonary_fibroblasts_scleroderma_associated_interstitial_lung_disease___biological_rep8 | 0 | 0 | 1 | |
| Pulmonary_fibroblasts_Usual_Interstitial_Pneumonia___biological_rep1 | 1 | 0 | 1 | |
| Pulmonary_fibroblasts_control_individual___biological_rep5 | 1 | 0 | 0 | |
| Pulmonary_fibroblasts_control_individual___biological_rep7 | 1 | 1 | 0 | |
| Pulmonary_fibroblasts_control_individual___biological_rep8 | 1 | 0 | 0 | |
| Pulmonary_fibroblasts_scleroderma_associated_interstitial_lung_disease___biological_rep1 | 1 | 0 | 0 | |
| Pulmonary_fibroblasts_scleroderma_associated_interstitial_lung_disease___biological_rep2 | 1 | 0 | 0 | |
| Pulmonary_fibroblasts_scleroderma_associated_interstitial_lung_disease___biological_rep4 | 1 | 0 | 0 | |
| Pulmonary_fibroblasts_Usual_Interstitial_Pneumonia___biological_rep2 | 1 | 0 | 0 | |

#### 1.1 Outliers (All Methods)

| Outliers overall |
|---|
| Pulmonary_fibroblasts_control_individual___biological_rep4 |
| Pulmonary_fibroblasts_control_individual___biological_rep6 |
| Pulmonary_fibroblasts_Usual_Interstitial_Pneumonia___biological_rep1 |
| Pulmonary_fibroblasts_control_individual___biological_rep7 |

##### 1.2 Outliers (At Least One Method)

| Outliers at least 1 |
|---|
| Pulmonary__fibroblasts__control__individual___biological__rep4 |
| Pulmonary__fibroblasts__control__individual___biological__rep6 |
| Pulmonary__fibroblasts__scleroderma__associated__interstitial__lung__disease___biological__rep7 |
| Pulmonary__fibroblasts__scleroderma__associated__interstitial__lung__disease___biological__rep8 |
| Pulmonary__fibroblasts__Usual__Interstitial__Pneumonia___biological__rep1 |
| Pulmonary__fibroblasts__control__individual___biological__rep5 |
| Pulmonary__fibroblasts__control__individual___biological__rep7 |
| Pulmonary__fibroblasts__control__individual___biological__rep8 |
| Pulmonary__fibroblasts__scleroderma__associated__interstitial__lung__disease___biological__rep1 |
| Pulmonary__fibroblasts__scleroderma__associated__interstitial__lung__disease___biological__rep2 |
| Pulmonary__fibroblasts__scleroderma__associated__interstitial__lung__disease___biological__rep4 |
| Pulmonary__fibroblasts__Usual__Interstitial__Pneumonia___biological__rep2 |

#### 2 RNA Degradation

##### 2.1 Summarized Mean QC

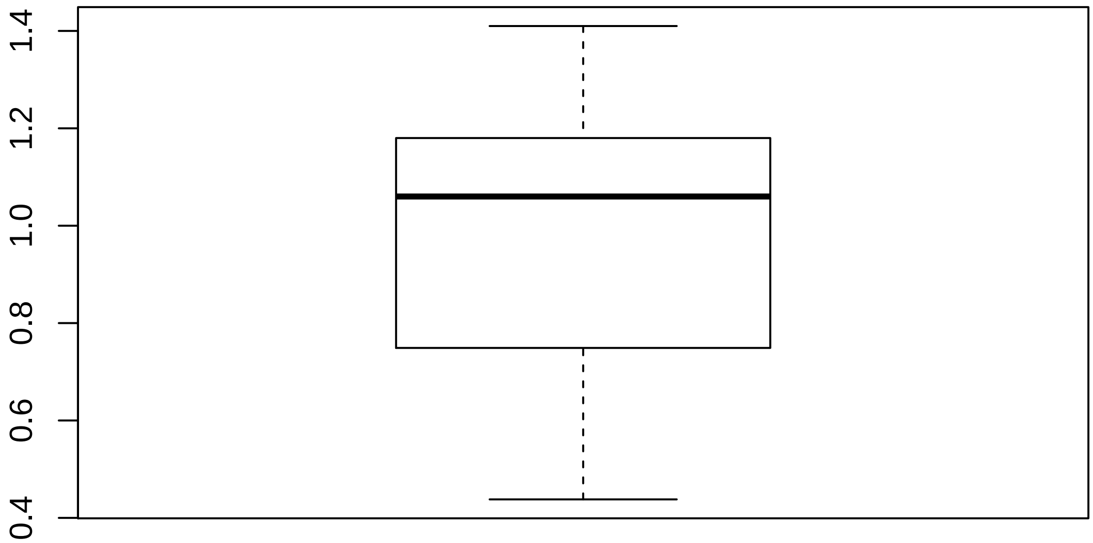

#### 2.2 Discrete QC Plots

**Sample Group [1]**

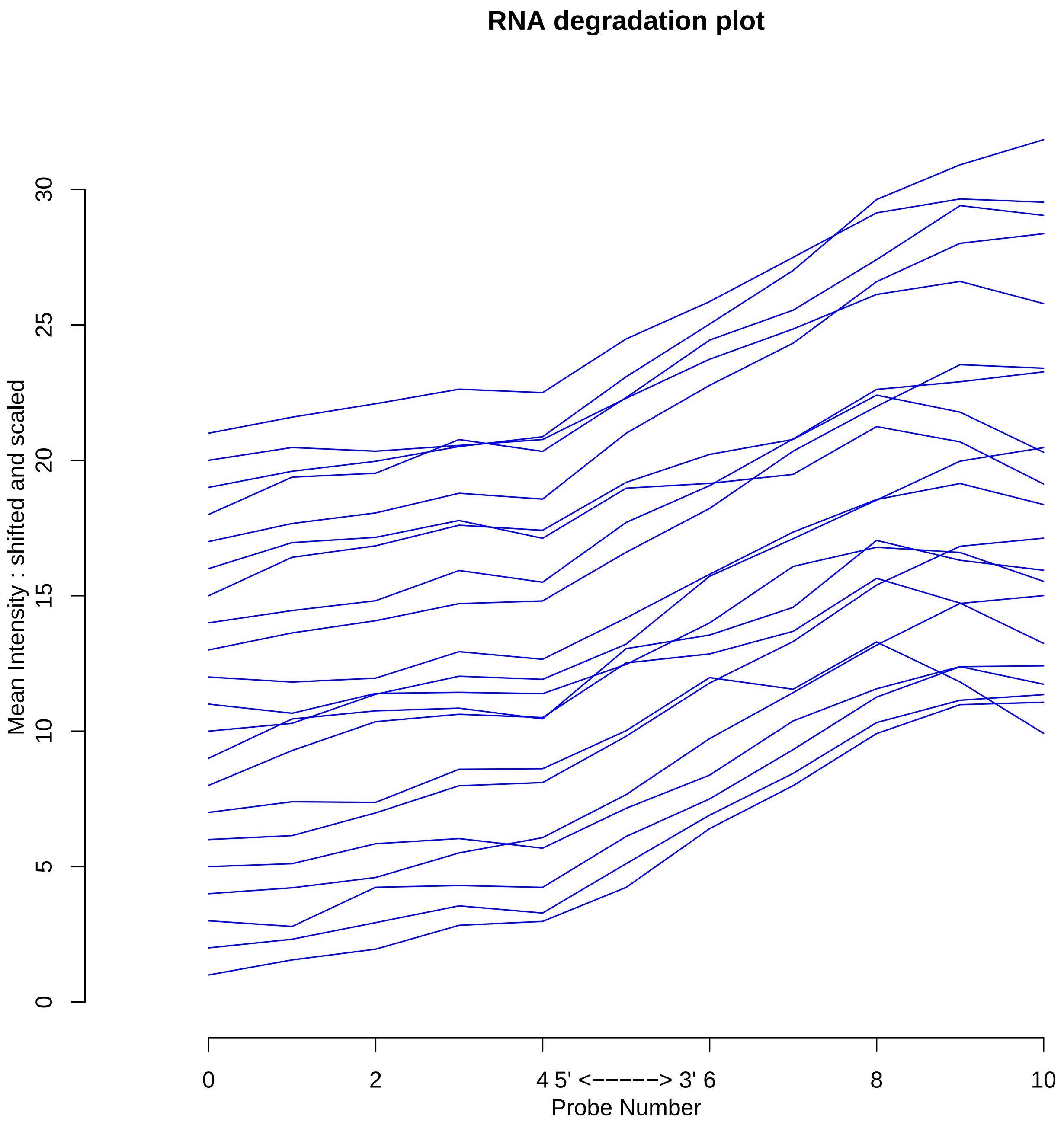

### 3 Relative Log Expression

#### 3.1 Summarized Median QC

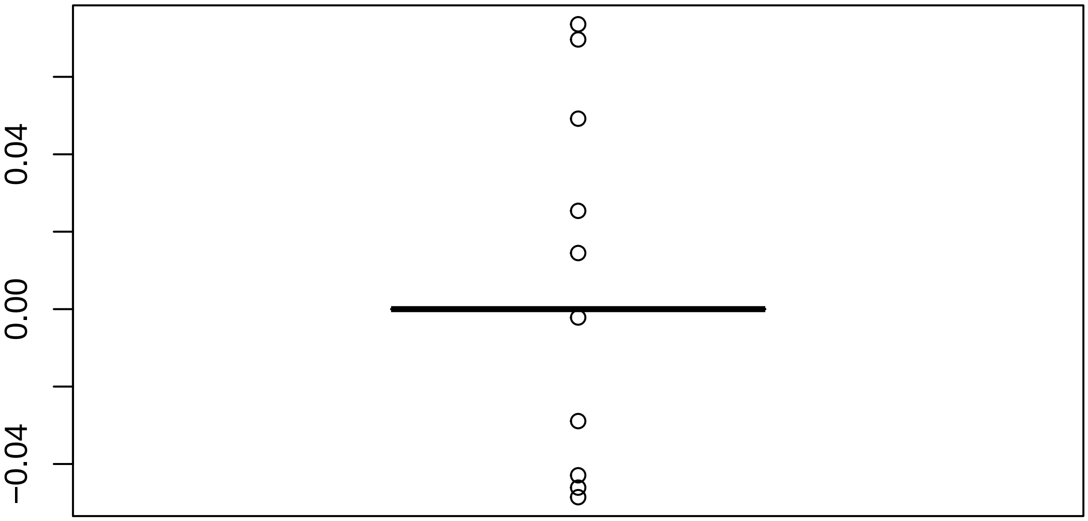

#### 3.2 Discrete QC Plots

Sample Group [1]

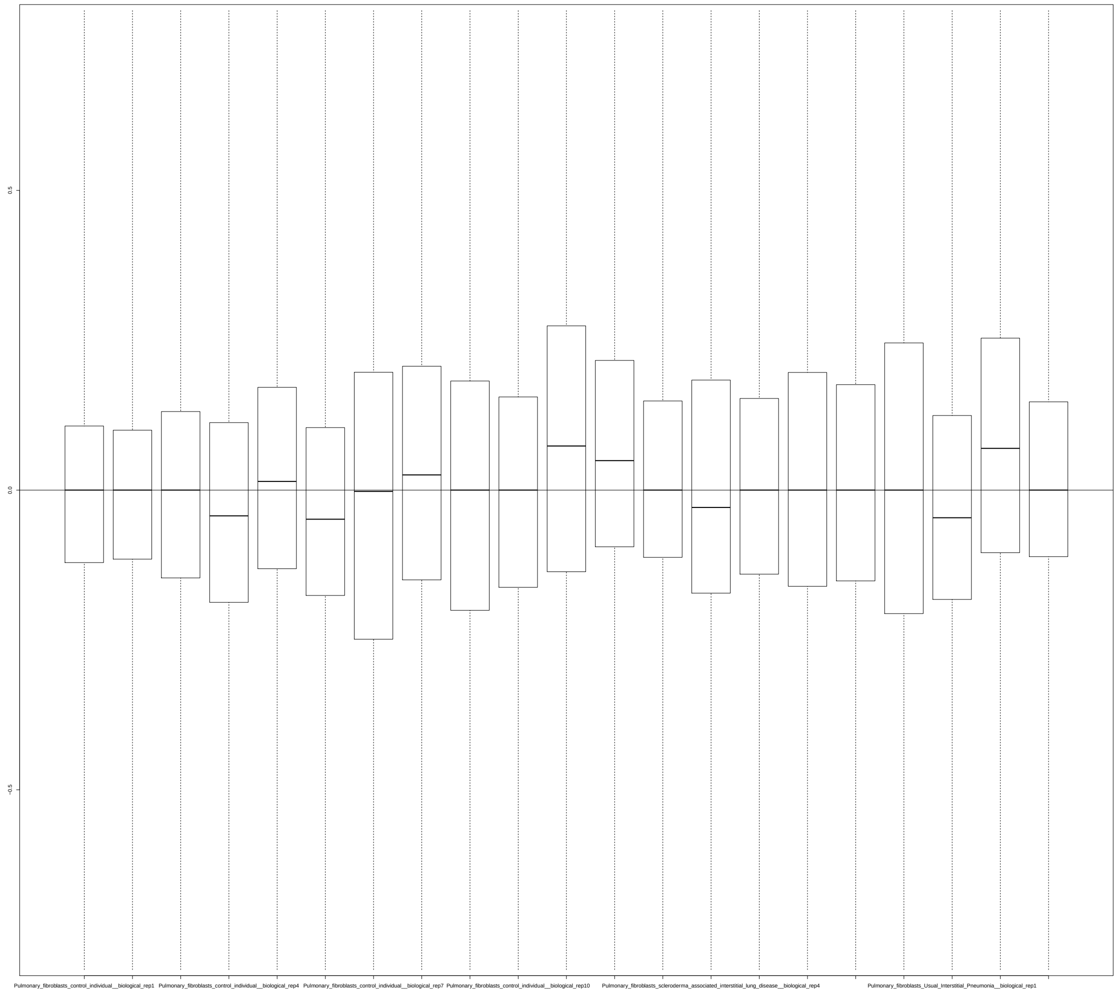

### 4 Normalized Unscaled Standard Errors

#### 4.1 Summarized Median QC

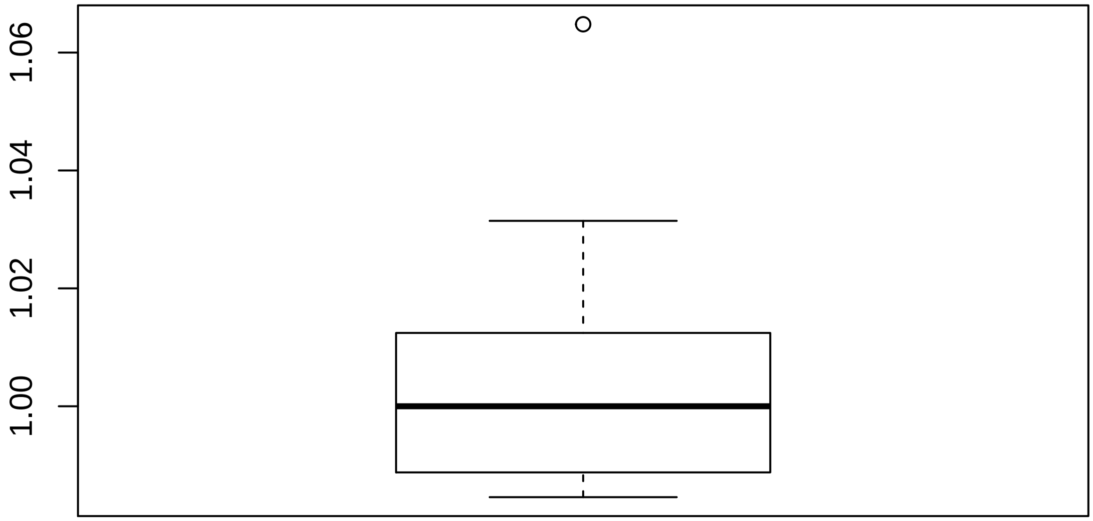

#### 4.2 Discrete QC Plots

**Sample Group [1]**

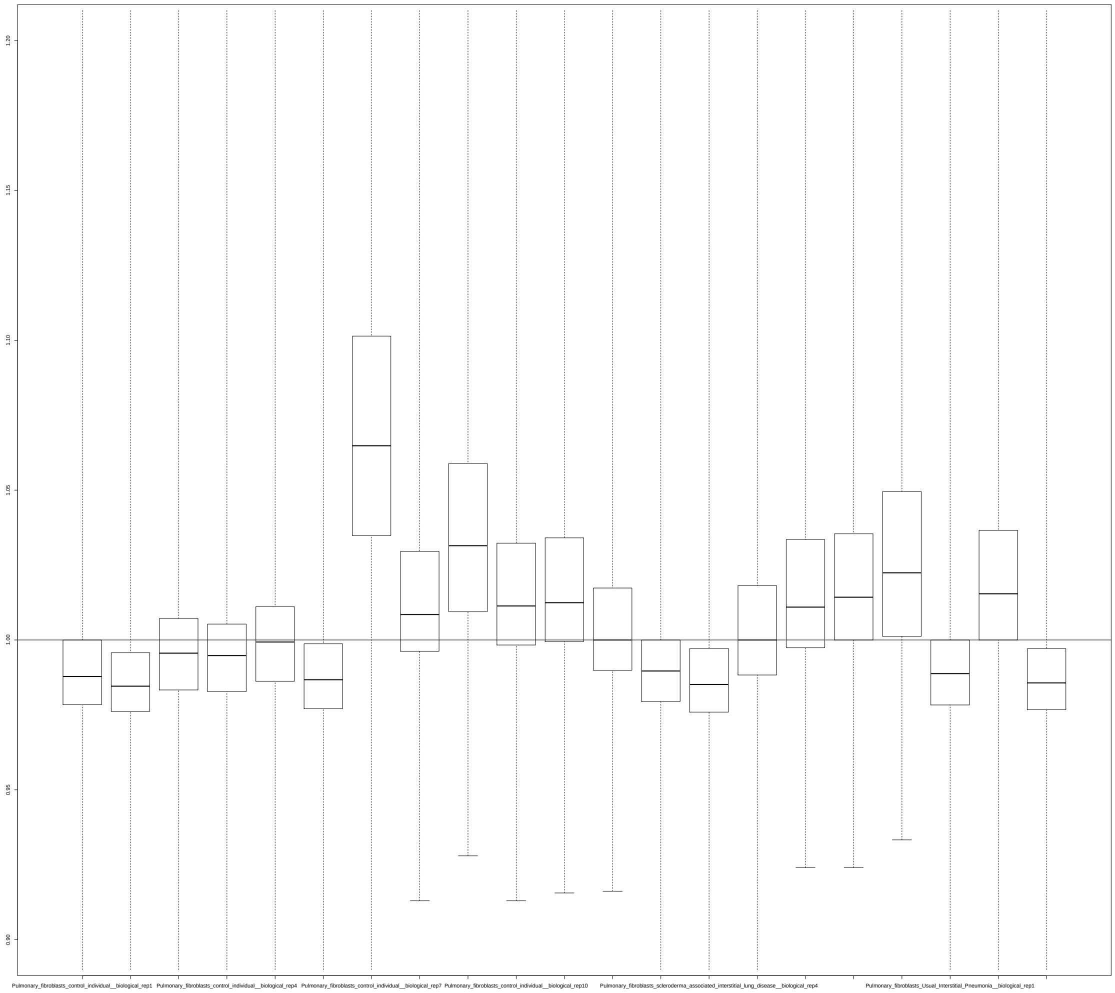

### 5 YAQC Plots

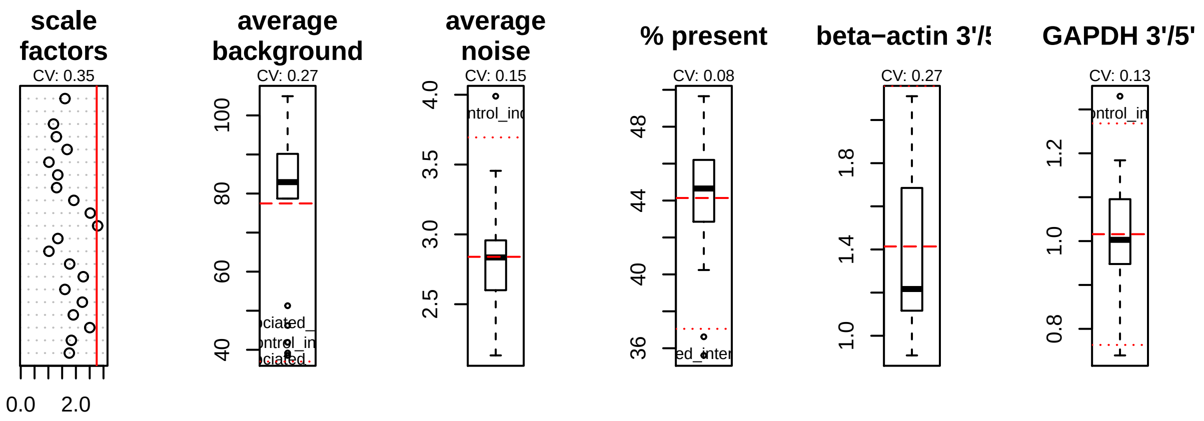
scale factors
CV: 0.35
0.0
2.0
average background
CV: 0.27
average noise
CV: 0.15
% present
CV: 0.08
beta−actin 3'/5
CV: 0.27
GAPDH 3'/5'
CV: 0.13

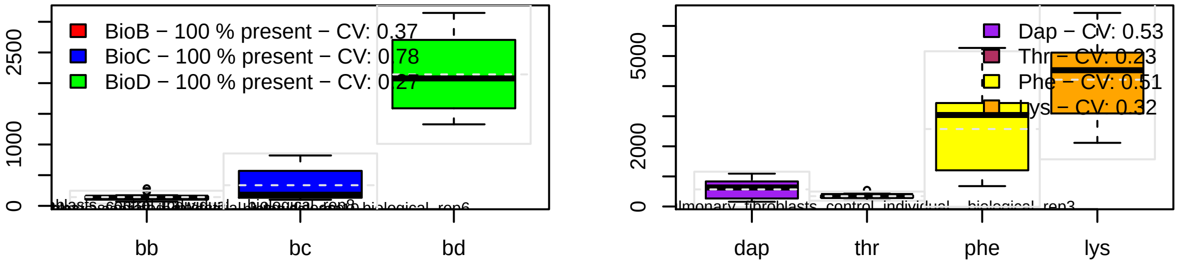
BioB − 100 % present − CV: 0.37
BioC − 100 % present − CV: 0.78
BioD − 100 % present − CV: 0.27
bb
bc
bd
Dap − CV: 0.53
Thr − CV: 0.23
Phe − CV: 0.51
lys − CV: 0.32
dap
thr
phe
lys
