## Supplementary material for "Curated and harmonised transcriptomics datasets of interstitial lung diseases": GSE44723_eUTOPIA_Affymetrix_QC_Report_2024-02-12-3.pdf

*eUTOPIA*

**5 YAQC Plots** **8**

### 1 Outliers Table

| | RLE | NUSE | DEG | SUM |
|---|---|---|---|---|
| fibroblast__Normal__D133__Media__3 | 0 | 0 | 1 | 1 |
| fibroblast__Rapid__D200__Media__14 | 0 | 0 | 1 | 1 |
| fibroblast__Rapid__D26__Media__15 | 0 | 0 | 1 | 1 |
| fibroblast__Rapid__D10__Media__13 | 0 | 1 | 0 | 1 |
| fibroblast__Normal__D207__Media__6 | 0 | 1 | 0 | 1 |

#### 1.1 Outliers (All Methods)

| Outliers overall |
|---|
| NA |

#### 1.2 Outliers (At Least One Method)

| Outliers at least 1 |
|---|
| fibroblast__Normal__D133__Media__3 |
| fibroblast__Rapid__D200__Media__14 |
| fibroblast__Rapid__D26__Media__15 |
| fibroblast__Rapid__D10__Media__13 |
| fibroblast__Normal__D207__Media__6 |

### 2 RNA Degradation

#### 2.1 Summarized Mean QC

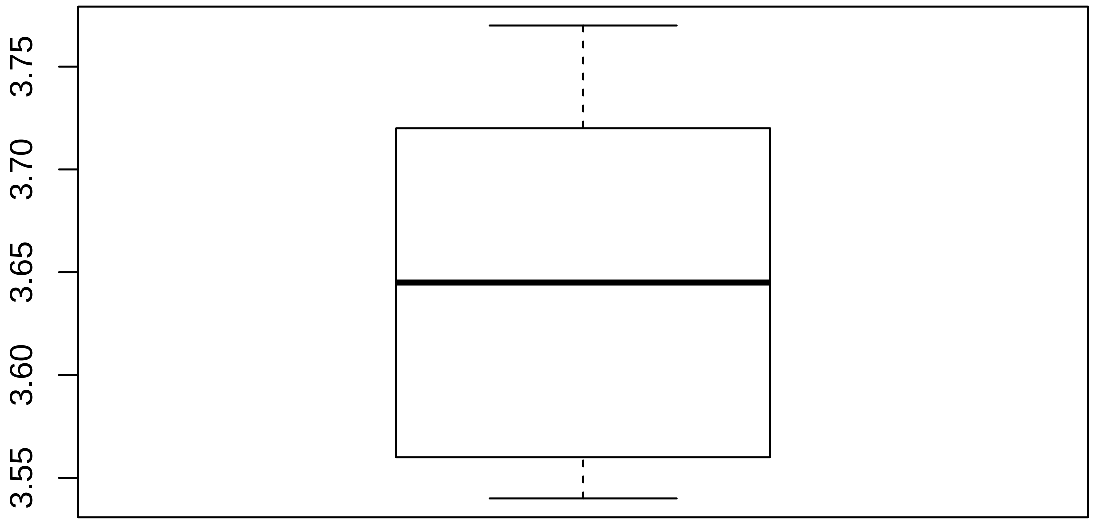

#### 2.2 Discrete QC Plots

**Sample Group [1]**

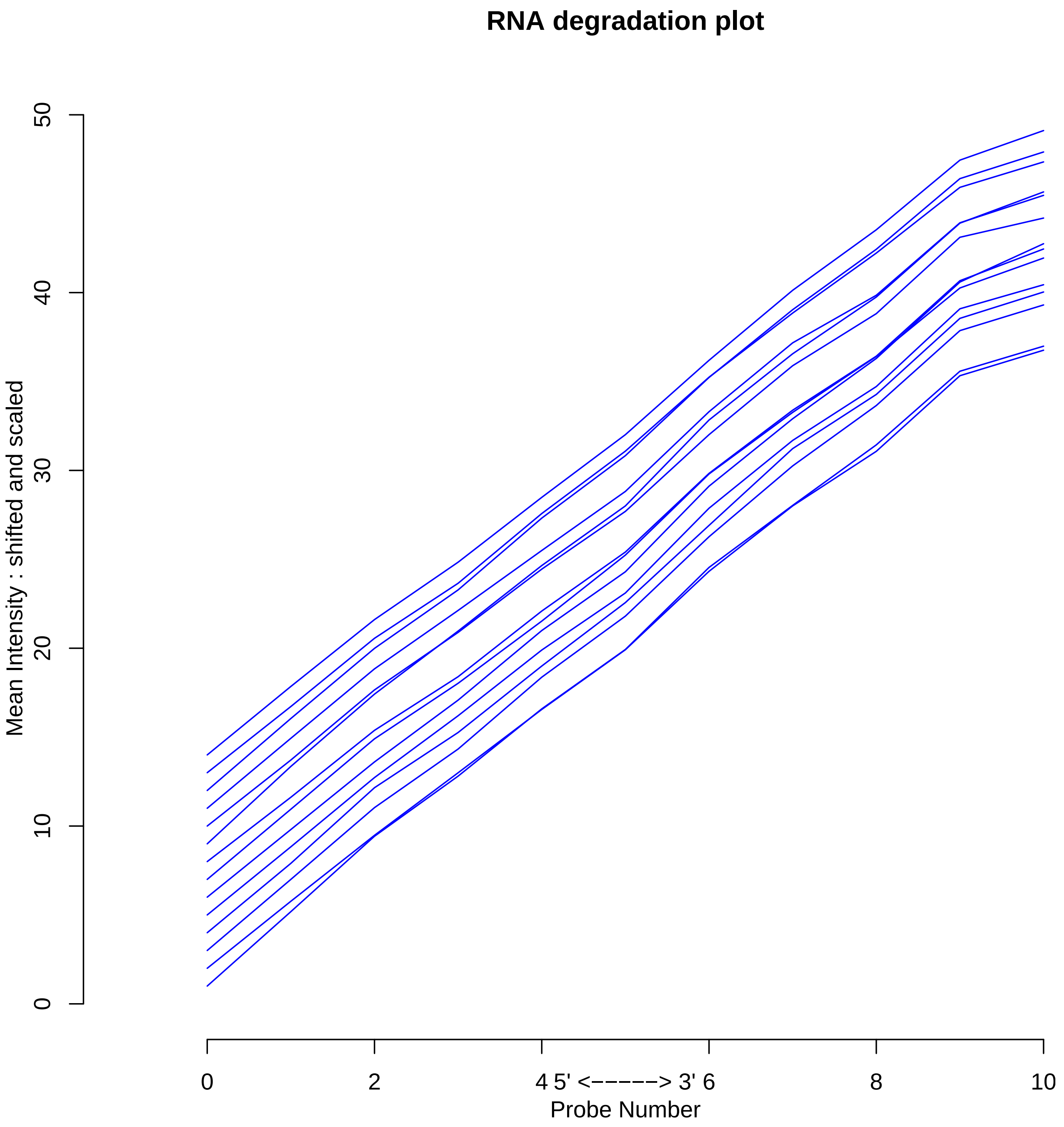

### 3 Relative Log Expression

#### 3.1 Summarized Median QC

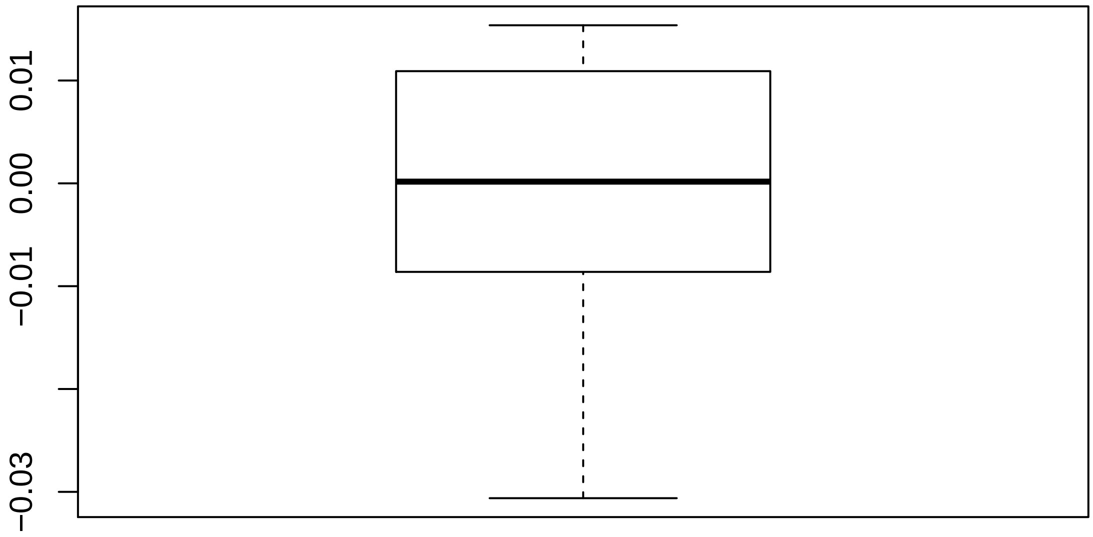

#### 3.2 Discrete QC Plots

**Sample Group [1]**

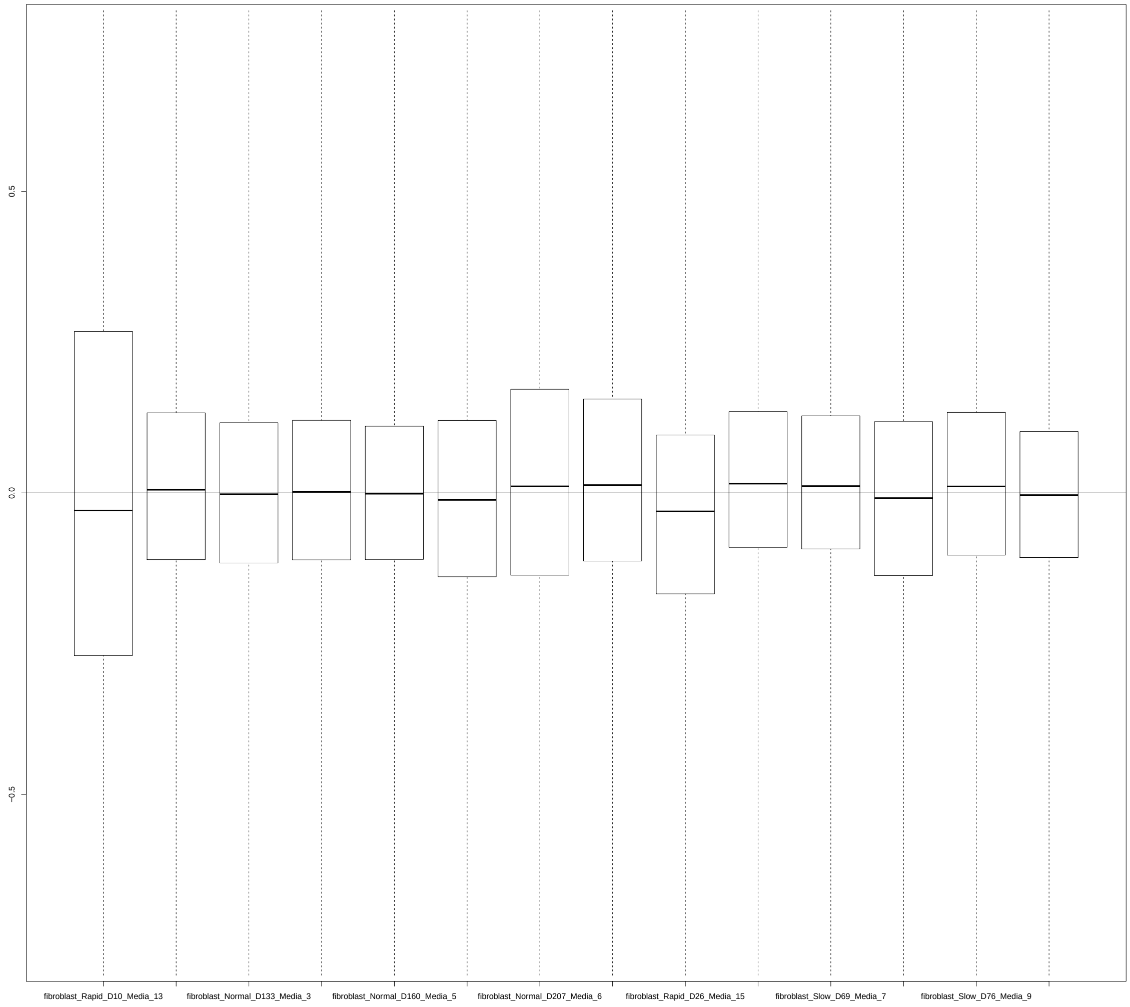

### 4 Normalized Unscaled Standard Errors

#### 4.1 Summarized Median QC

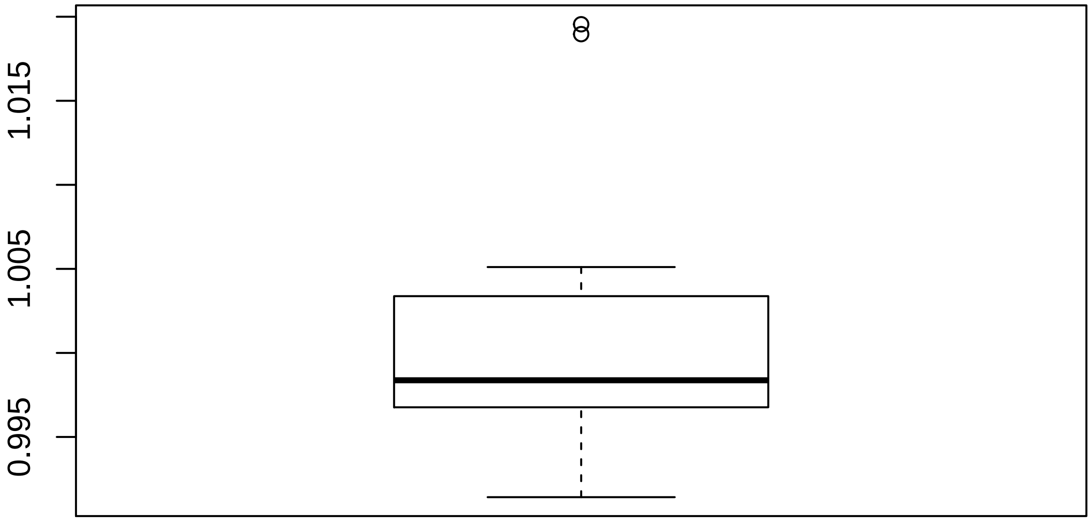

#### 4.2 Discrete QC Plots

**Sample Group [1]**

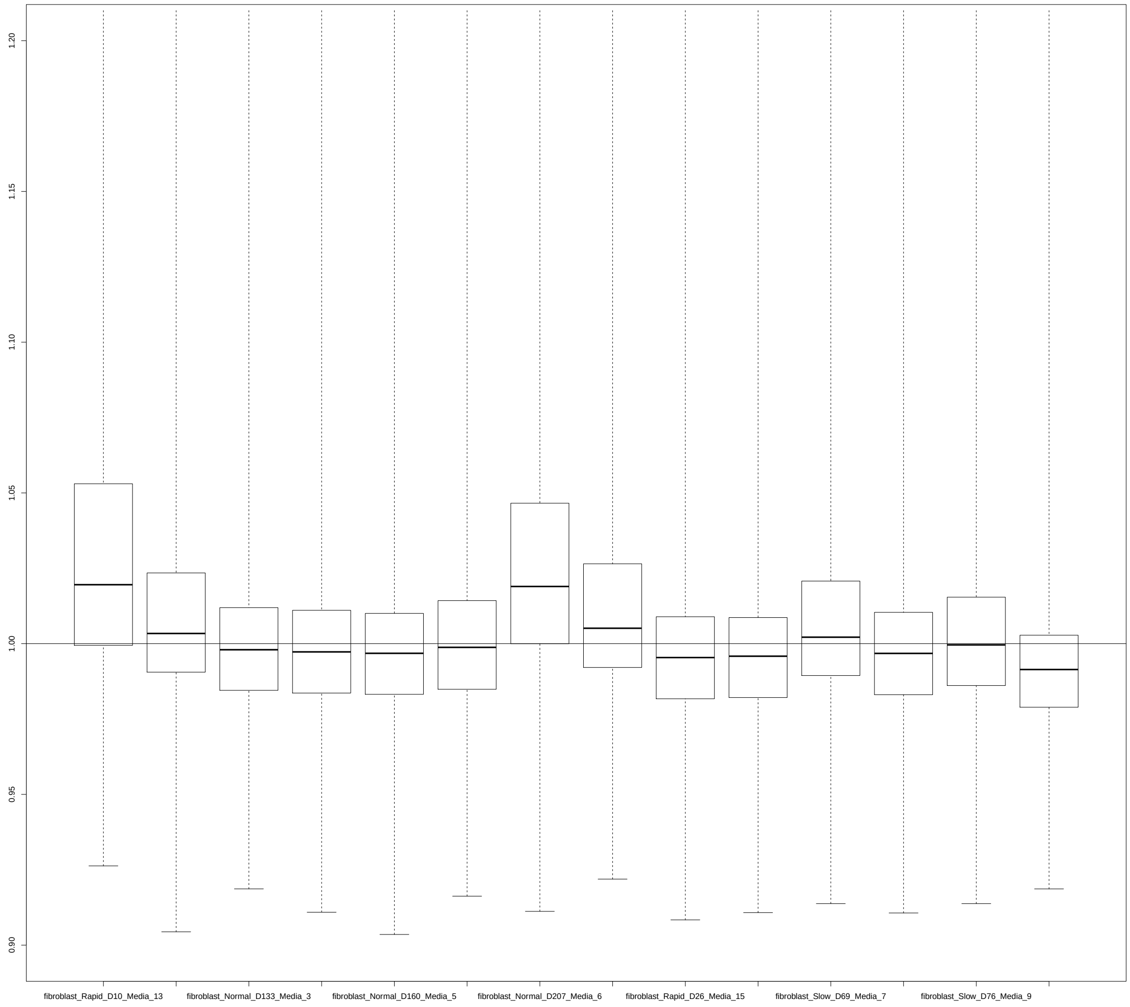

### 5 YAQC Plots

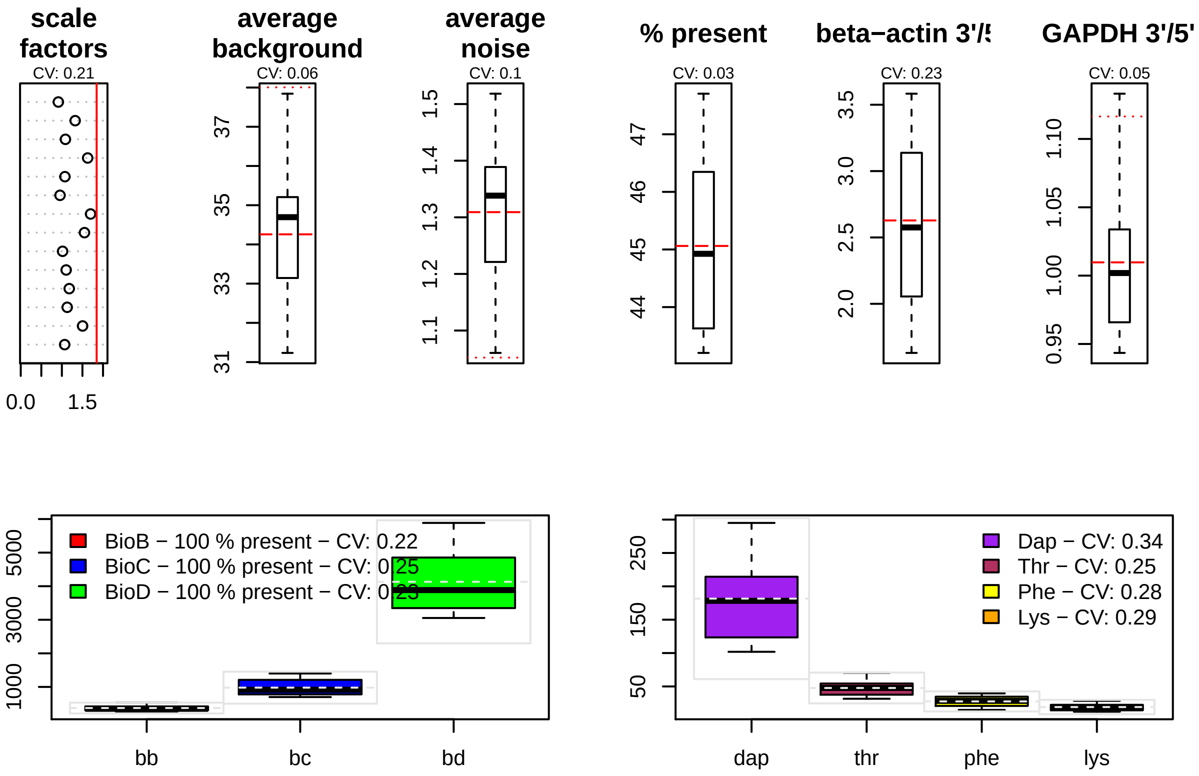

scale factors
CV: 0.21
0.0
1.5
average background
CV: 0.06
31
33
35
37
average noise
CV: 0.1
1.1
1.2
1.3
1.4
1.5
% present
CV: 0.03
44
45
46
47
beta−actin 3'/5
CV: 0.23
2.0
2.5
3.0
3.5
GAPDH 3'/5'
CV: 0.05
0.95
1.00
1.05
1.10
BioB − 100 % present − CV: 0.22
BioC − 100 % present − CV: 0.25
BioD − 100 % present − CV: 0.23
1000
3000
5000
bb
bc
bd
Dap − CV: 0.34
Thr − CV: 0.25
Phe − CV: 0.28
Lys − CV: 0.29
50
150
250
dap
thr
phe
lys
