## Supplementary material for "Curated and harmonised transcriptomics datasets of interstitial lung diseases": GSE49072_eUTOPIA_Affymetrix_QC_Report_2024-02-12-4.pdf

*eUTOPIA*

**5 YAQC Plots** **12**

### 1 Outliers Table

| | RLE | NUSE | DEG | SUM |
|---|---|---|---|---|
| Normal_Volunteer,_Replicate_21 | 0 | 0 | 1 | 1 |
| Familial_IPF,_Replicate_3 | 0 | 0 | 1 | 1 |
| Normal_Volunteer,_Replicate_25 | 0 | 0 | 1 | 1 |
| Spontaneous_IPF,_Replicate_11 | 1 | 0 | 1 | 2 |
| Normal_Volunteer,_Replicate_42 | 1 | 0 | 1 | 2 |
| Spontaneous_IPF,_Replicate_14 | 0 | 1 | 1 | 2 |
| Normal_Volunteer,_Replicate_5 | 0 | 0 | 1 | 1 |
| Spontaneous_IPF,_Replicate_5 | 0 | 0 | 1 | 1 |
| Normal_Volunteer,_Replicate_28 | 0 | 0 | 1 | 1 |
| Normal_Volunteer,_Replicate_39 | 0 | 0 | 1 | 1 |
| Spontaneous_IPF,_Replicate_12 | 0 | 1 | 1 | 2 |
| Familial_IPF,_Replicate_2 | 0 | 0 | 1 | 1 |
| Spontaneous_IPF,_Replicate_9 | 0 | 0 | 1 | 1 |
| Normal_Volunteer,_Replicate_41 | 0 | 0 | 1 | 1 |
| Spontaneous_IPF,_Replicate_13 | 0 | 0 | 1 | 1 |
| Normal_Volunteer,_Replicate_45 | 0 | 1 | 1 | 2 |
| Familial_IPF,_Replicate_1 | 0 | 0 | 1 | 1 |
| Normal_Volunteer,_Replicate_24 | 0 | 0 | 1 | 1 |
| Spontaneous_IPF,_Replicate_6 | 0 | 0 | 1 | 1 |
| Normal_Volunteer,_Replicate_40 | 0 | 0 | 1 | 1 |
| Normal_Volunteer,_Replicate_43 | 0 | 0 | 1 | 1 |
| Normal_Relative,_Replicate_9 | 1 | 1 | 0 | 2 |
| Normal_Volunteer,_Replicate_18 | 1 | 0 | 0 | 1 |
| Normal_Relative,_Replicate_1 | 1 | 0 | 0 | 1 |
| Normal_Volunteer,_Replicate_12 | 1 | 0 | 0 | 1 |

| | RLE | NUSE | DEG | SUM |
|---|---|---|---|---|
| Normal_Volunteer,_Replicate_15 | 1 | 0 | 0 | 1 |
| Normal_Volunteer,_Replicate_32 | 1 | 1 | 0 | 2 |
| Normal_Volunteer,_Replicate_35 | 0 | 1 | 0 | 1 |
| Normal_Volunteer,_Replicate_23 | 0 | 1 | 0 | 1 |
| Spontaneous_IPF,_Replicate_3 | 0 | 1 | 0 | 1 |
| Familial_IPF,_Replicate_6 | 0 | 1 | 0 | 1 |
| Spontaneous_IPF,_Replicate_15 | 0 | 1 | 0 | 1 |

#### 1.1 Outliers (All Methods)

| Outliers overall |
|---|
| Spontaneous_IPF,_Replicate_11 |
| Normal_Volunteer,_Replicate_42 |
| Spontaneous_IPF,_Replicate_14 |
| Spontaneous_IPF,_Replicate_12 |
| Normal_Volunteer,_Replicate_45 |
| Normal_Relative,_Replicate_9 |
| Normal_Volunteer,_Replicate_32 |

#### 1.2 Outliers (At Least One Method)

| Outliers at least 1 |
|---|
| Normal_Volunteer,_Replicate_21 |
| Familial_IPF,_Replicate_3 |
| Normal_Volunteer,_Replicate_25 |
| Spontaneous_IPF,_Replicate_11 |
| Normal_Volunteer,_Replicate_42 |
| Spontaneous_IPF,_Replicate_14 |
| Normal_Volunteer,_Replicate_5 |
| Spontaneous_IPF,_Replicate_5 |
| Normal_Volunteer,_Replicate_28 |
| Normal_Volunteer,_Replicate_39 |
| Spontaneous_IPF,_Replicate_12 |
| Familial_IPF,_Replicate_2 |
| Spontaneous_IPF,_Replicate_9 |
| Normal_Volunteer,_Replicate_41 |
| Spontaneous_IPF,_Replicate_13 |
| Normal_Volunteer,_Replicate_45 |
| Familial_IPF,_Replicate_1 |
| Normal_Volunteer,_Replicate_24 |
| Spontaneous_IPF,_Replicate_6 |
| Normal_Volunteer,_Replicate_40 |
| Normal_Volunteer,_Replicate_43 |
| Normal_Relative,_Replicate_9 |
| Normal_Volunteer,_Replicate_18 |
| Normal_Relative,_Replicate_1 |
| Normal_Volunteer,_Replicate_12 |
| Normal_Volunteer,_Replicate_15 |
| Normal_Volunteer,_Replicate_32 |

| Outliers at least 1 |
|---|
| Normal_Volunteer,_Replicate_35 |
| Normal_Volunteer,_Replicate_23 |
| Spontaneous_IPF,_Replicate_3 |
| Familial_IPF,_Replicate_6 |
| Spontaneous_IPF,_Replicate_15 |

### 2 RNA Degradation

#### 2.1 Summarized Mean QC

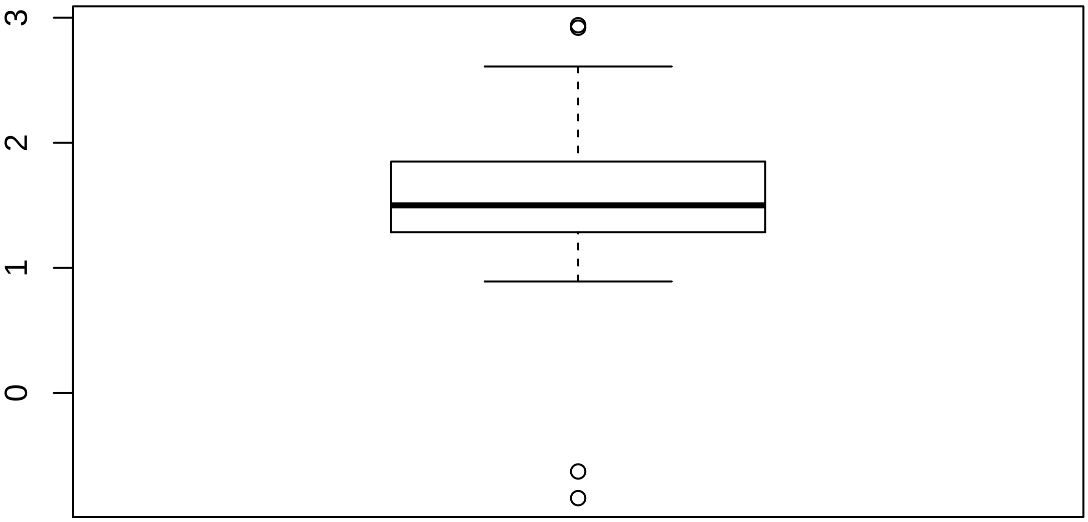

#### 2.2 Discrete QC Plots

**Sample Group [1]**

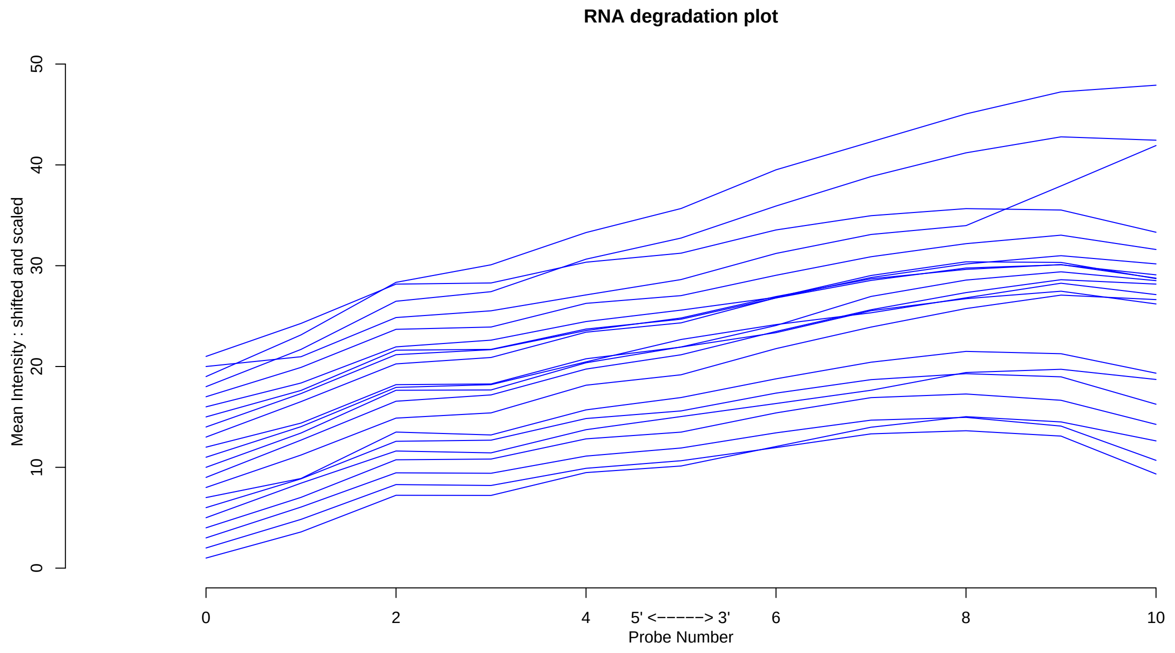

**Sample Group [2]**

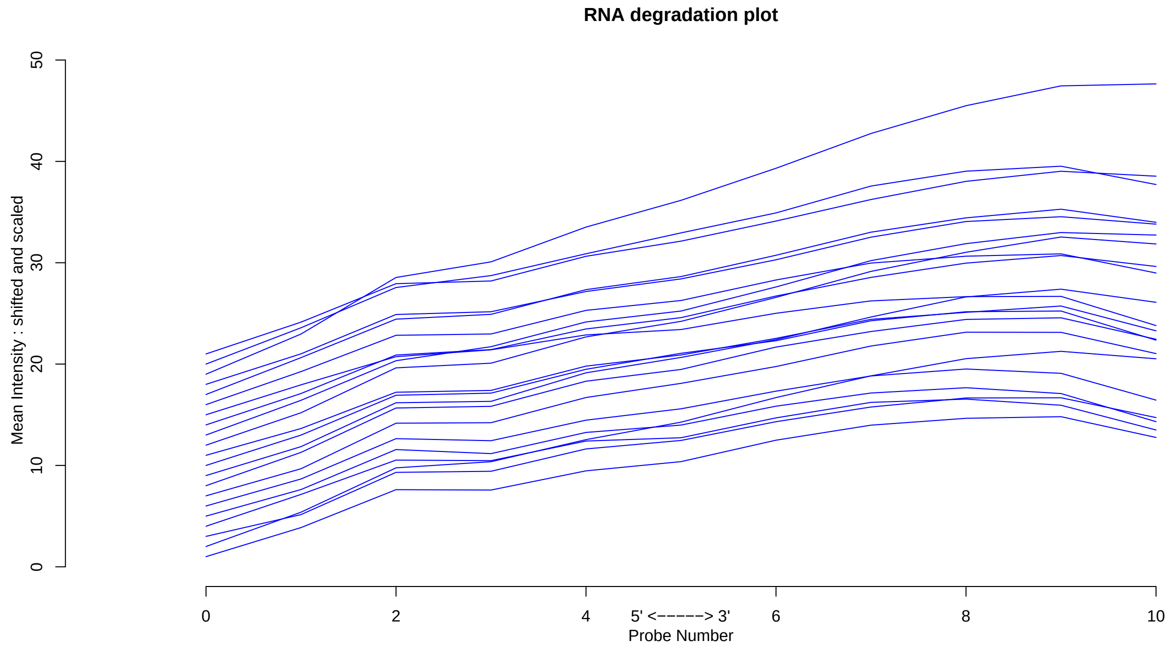

**Sample Group [3]**

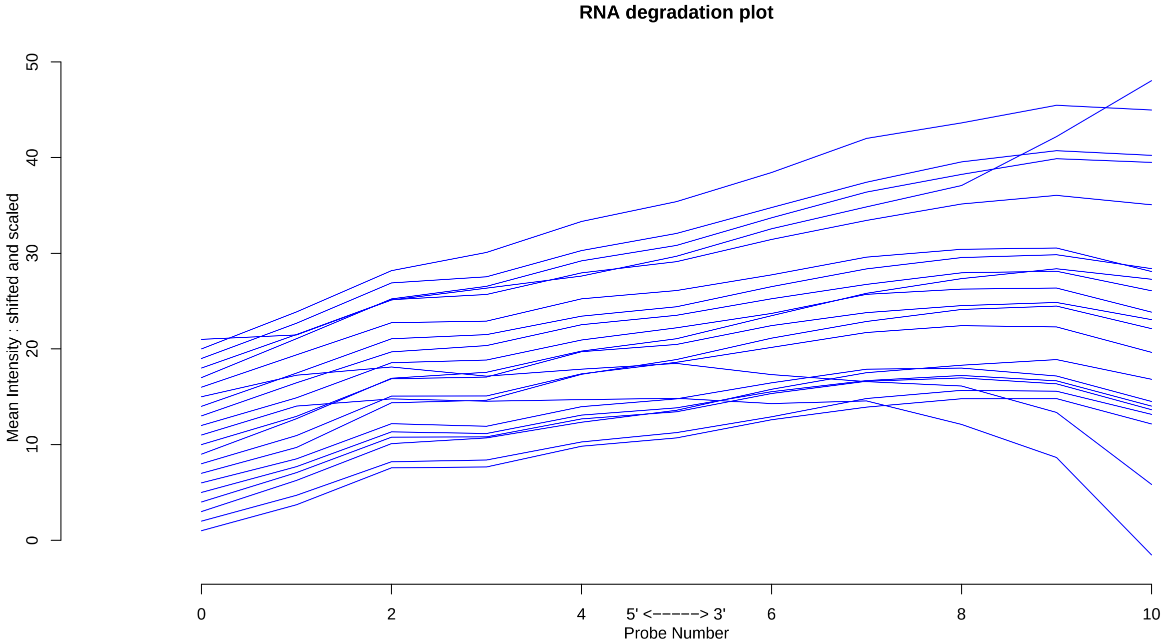

**Sample Group [4]**

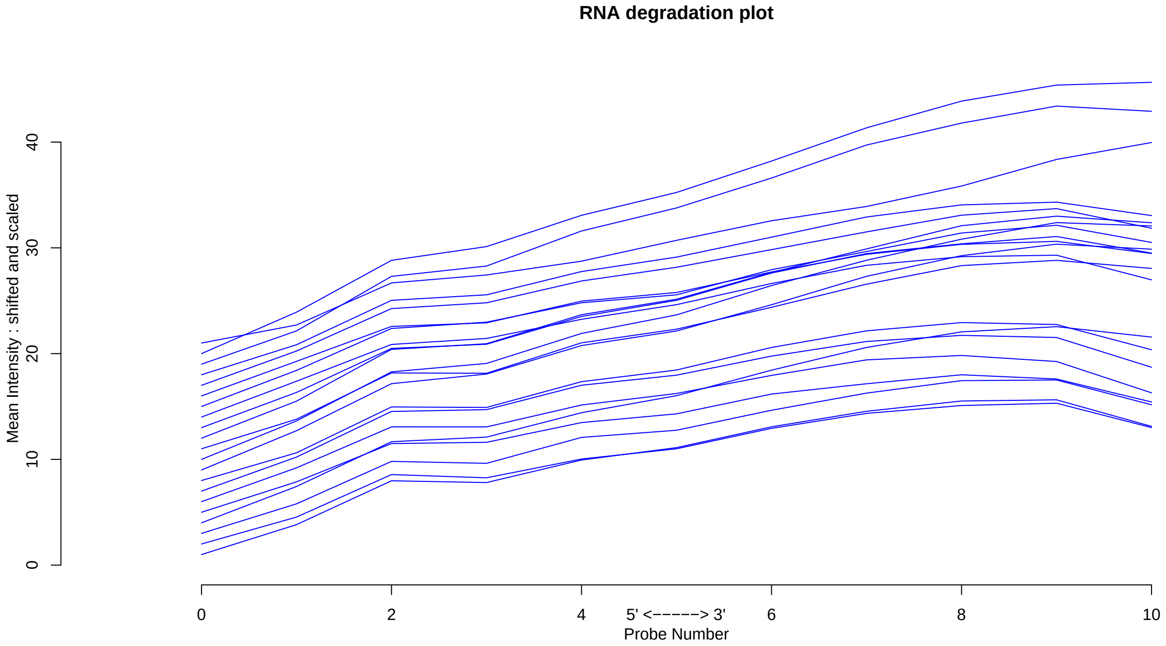

#### 3 Relative Log Expression

##### 3.1 Summarized Median QC

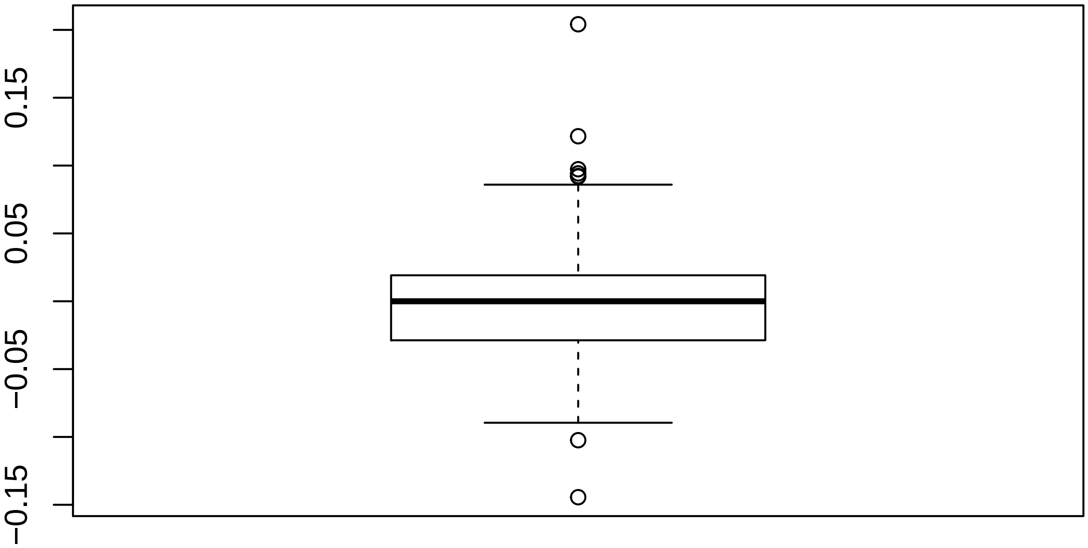

#### 3.2 Discrete QC Plots

Sample Group [1]

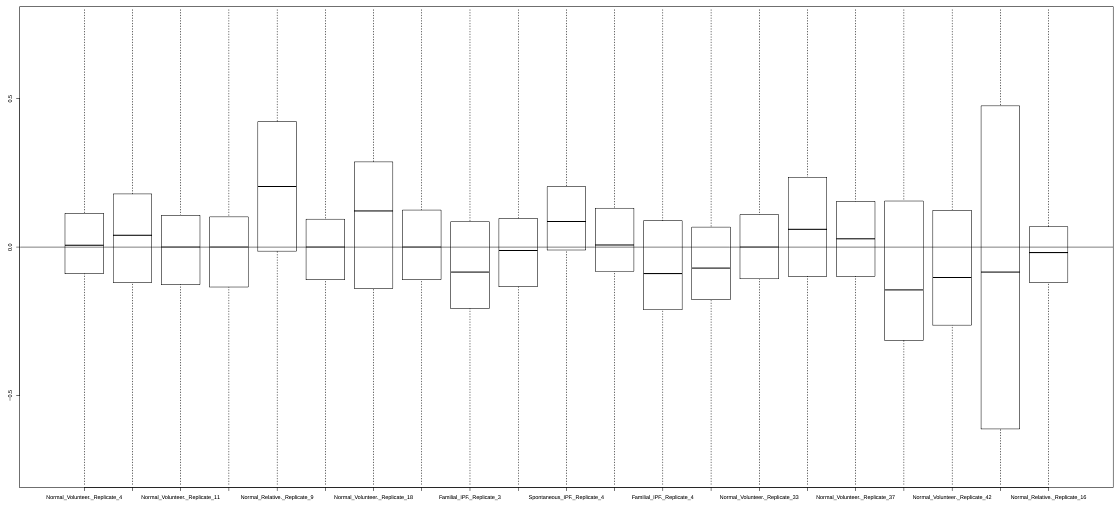

Sample Group [2]

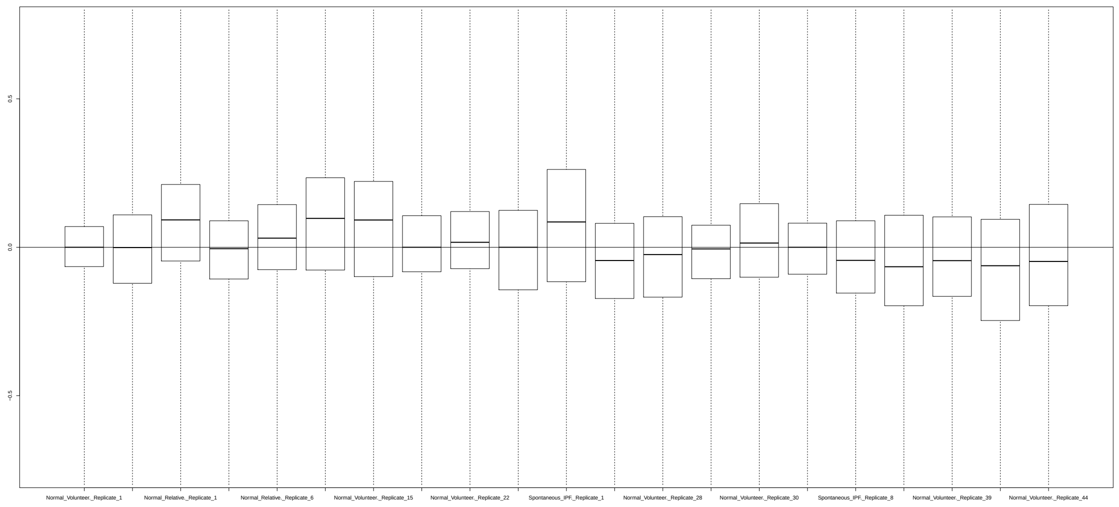

**Sample Group [3]**

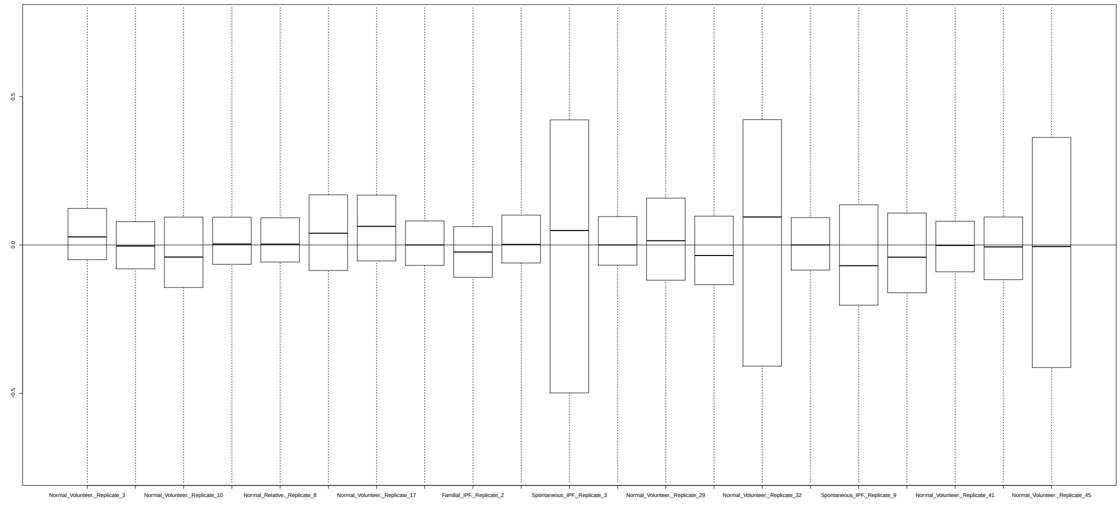

**Sample Group [4]**

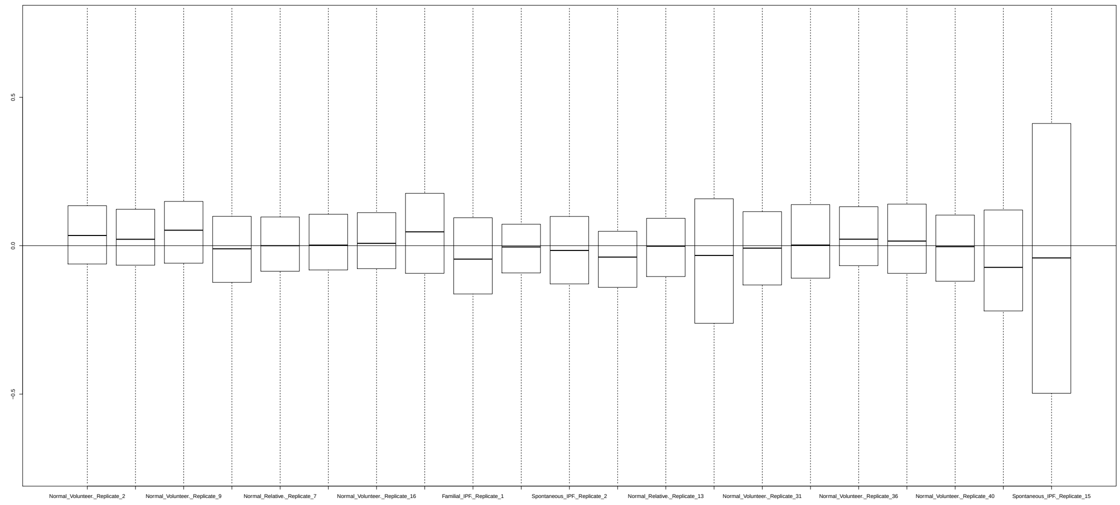

### 4 Normalized Unscaled Standard Errors

#### 4.1 Summarized Median QC

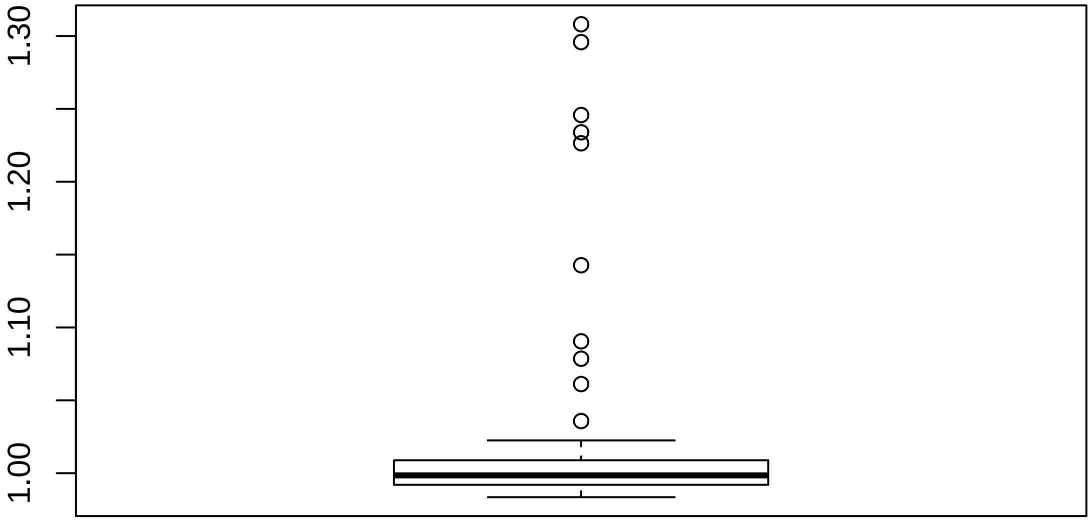

#### 4.2 Discrete QC Plots

Sample Group [1]

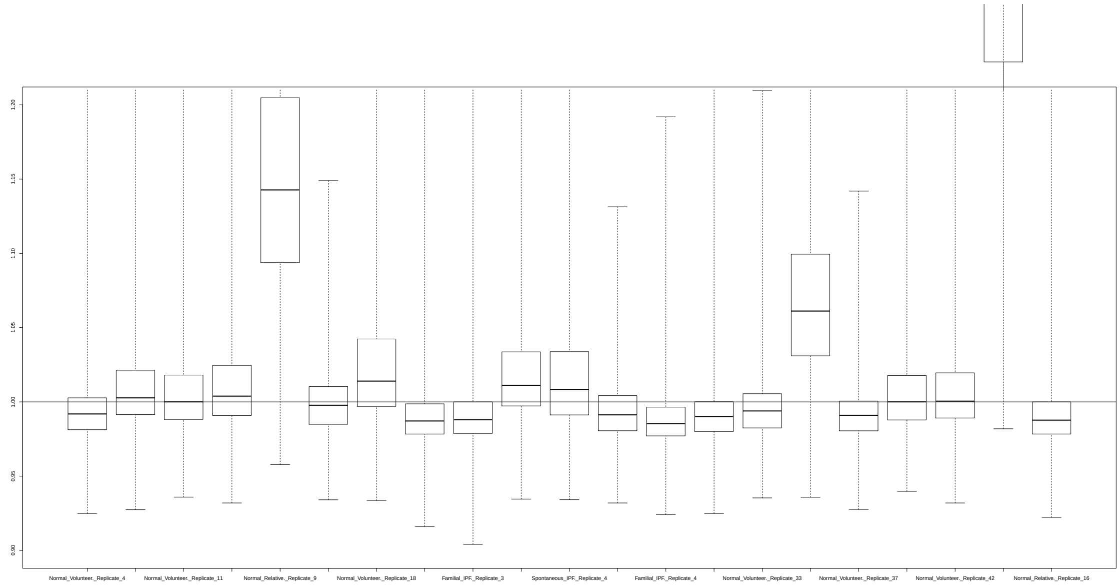

Sample Group [2]

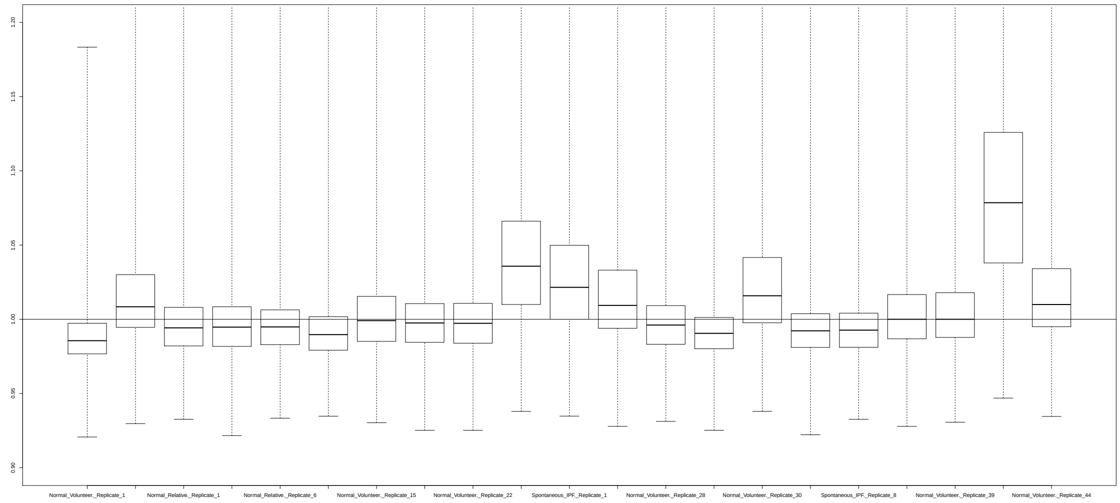

Sample Group [3]

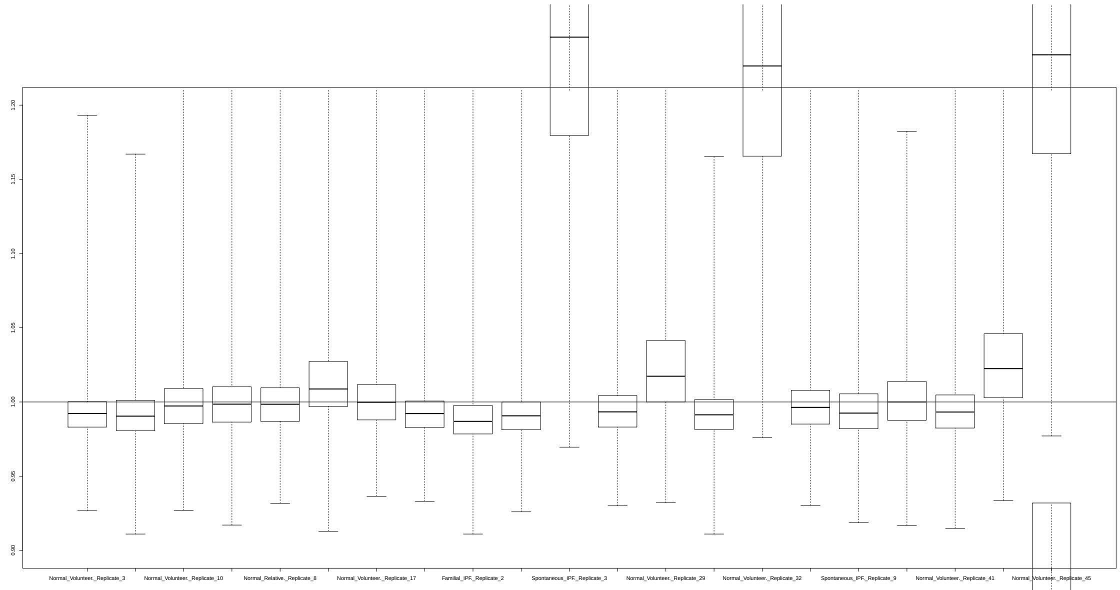

Sample Group [4]

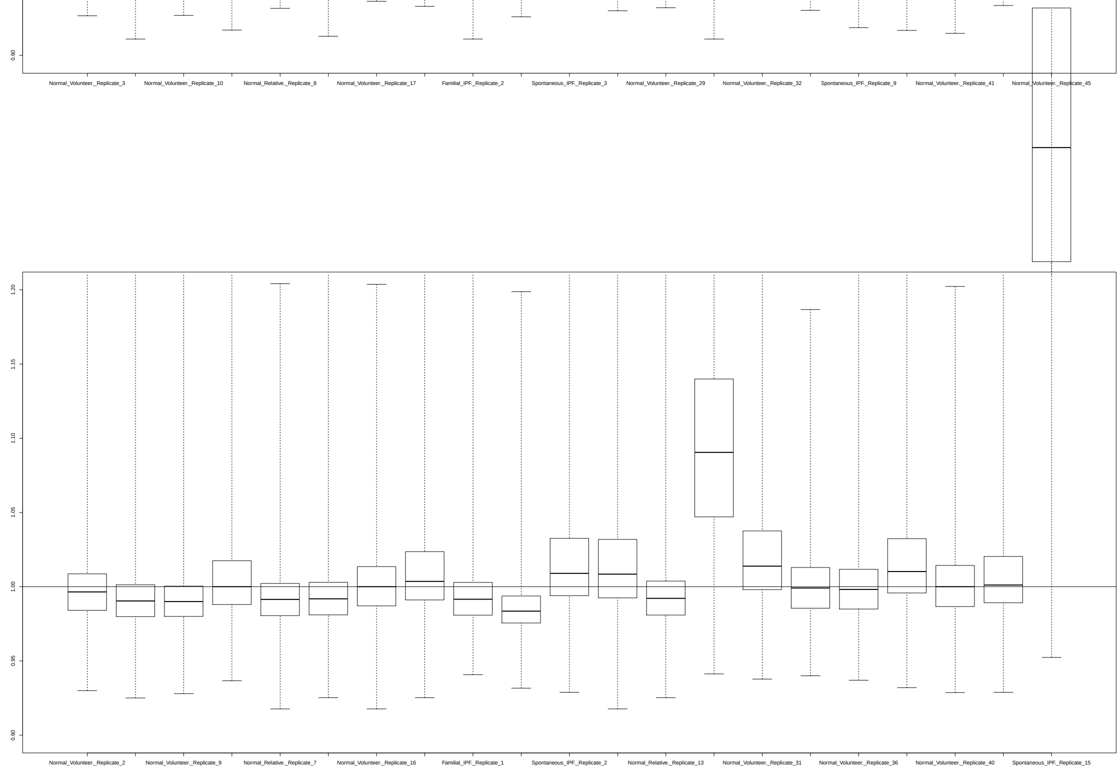

### 5 YAQC Plots

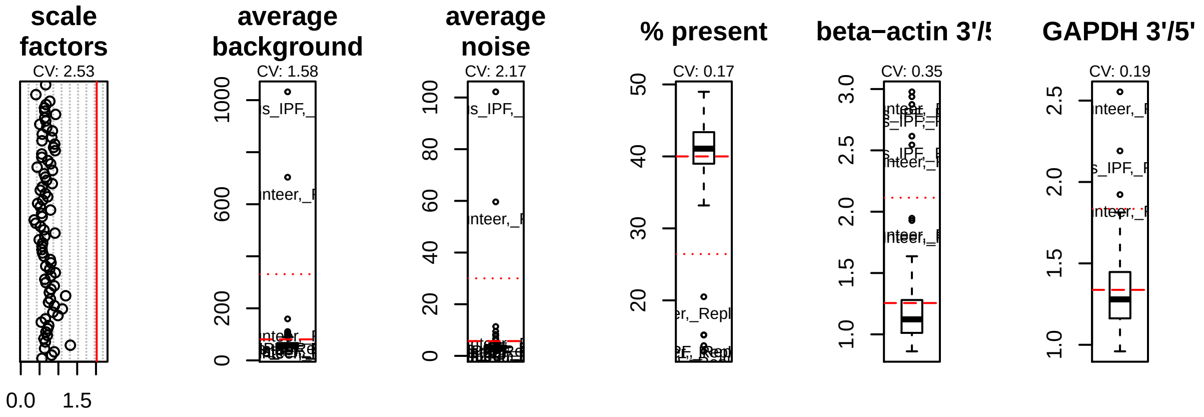
scale factors
CV: 2.53
0.0
1.5
average background
CV: 1.58
average noise
CV: 2.17
% present
CV: 0.17
beta−actin 3'/5
CV: 0.35
GAPDH 3'/5'
CV: 0.19

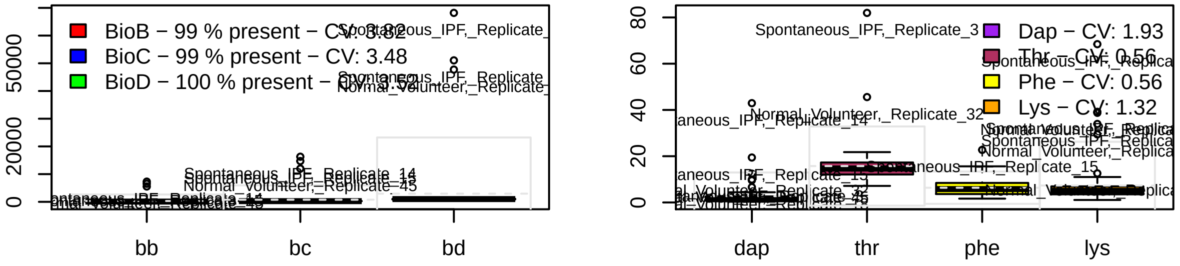
BioB − 99 % present − CV: 3.82
BioC − 99 % present − CV: 3.48
BioD − 100 % present − CV: 3.52
bb
bc
bd
Dap − CV: 1.93
Thr − CV: 0.56
Phe − CV: 0.56
Lys − CV: 1.32
dap
thr
phe
lys
Spontaneous_IPF,_Replicate_3
Normal_Volunteer,_Replicate_32
Spontaneous_IPF,_Replicate_14
