## Supplementary material for "Curated and harmonised transcriptomics datasets of interstitial lung diseases": GSE90010_eUTOPIA_Affymetrix_QC_Report_2024-02-12-4.pdf

*eUTOPIA*

### 1 Outliers Table

| | RLE | NUSE | DEG | SUM |
|---|---|---|---|---|
| Control_MDM_replicate_1 | 0 | 0 | 1 | 1 |
| Control_MDM_replicate_2 | 0 | 0 | 1 | 1 |
| Control_MDM_replicate_3 | 1 | 0 | 1 | 2 |
| Control_MDM_replicate_4 | 0 | 0 | 1 | 1 |
| MDM_co_cultured_with_AN_replicate_1 | 0 | 0 | 1 | 1 |
| AM_from_RB_ILD_patient_1 | 0 | 0 | 1 | 1 |
| MDM_co_cultured_with_AN_replicate_3 | 0 | 1 | 0 | 1 |
| AM_from_RB_ILD_patient_2 | 0 | 1 | 0 | 1 |

#### 1.1 Outliers (All Methods)

| Outliers overall |
|---|
| Control_MDM_replicate_3 |

#### 1.2 Outliers (At Least One Method)

| Outliers at least 1 |
|---|
| Control_MDM_replicate_1 |
| Control_MDM_replicate_2 |
| Control_MDM_replicate_3 |
| Control_MDM_replicate_4 |
| MDM_co_cultured_with_AN_replicate_1 |
| AM_from_RB_ILD_patient_1 |

| Outliers at least 1 |
| --- |
| MDM_co_cultured_with_AN_replicate_3 |
| AM_from_RB_ILD_patient_2 |

#### 2 RNA Degradation

##### 2.1 Summarized Mean QC

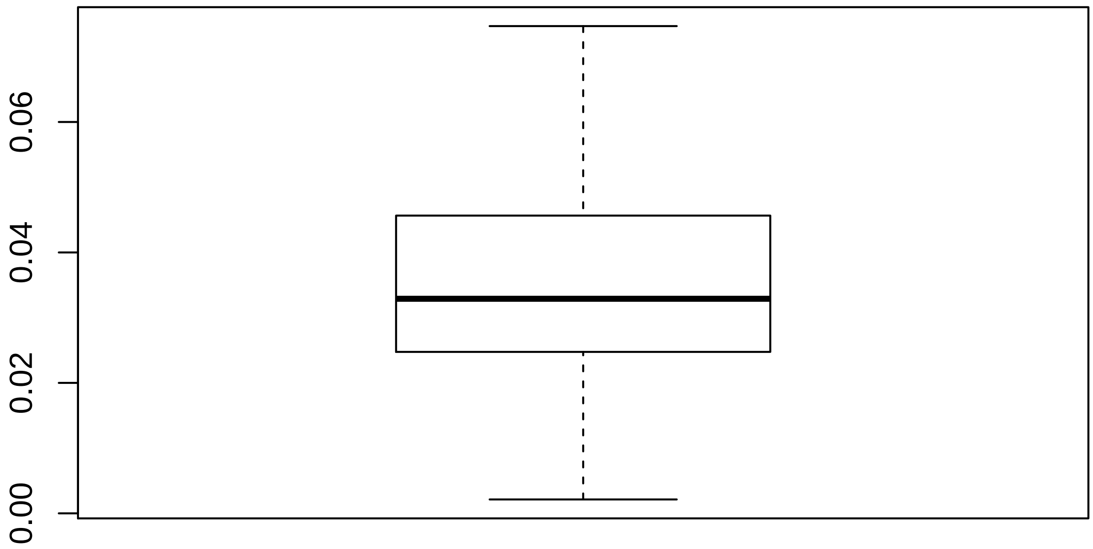

#### 2.2 Discrete QC Plots

**Sample Group [1]**

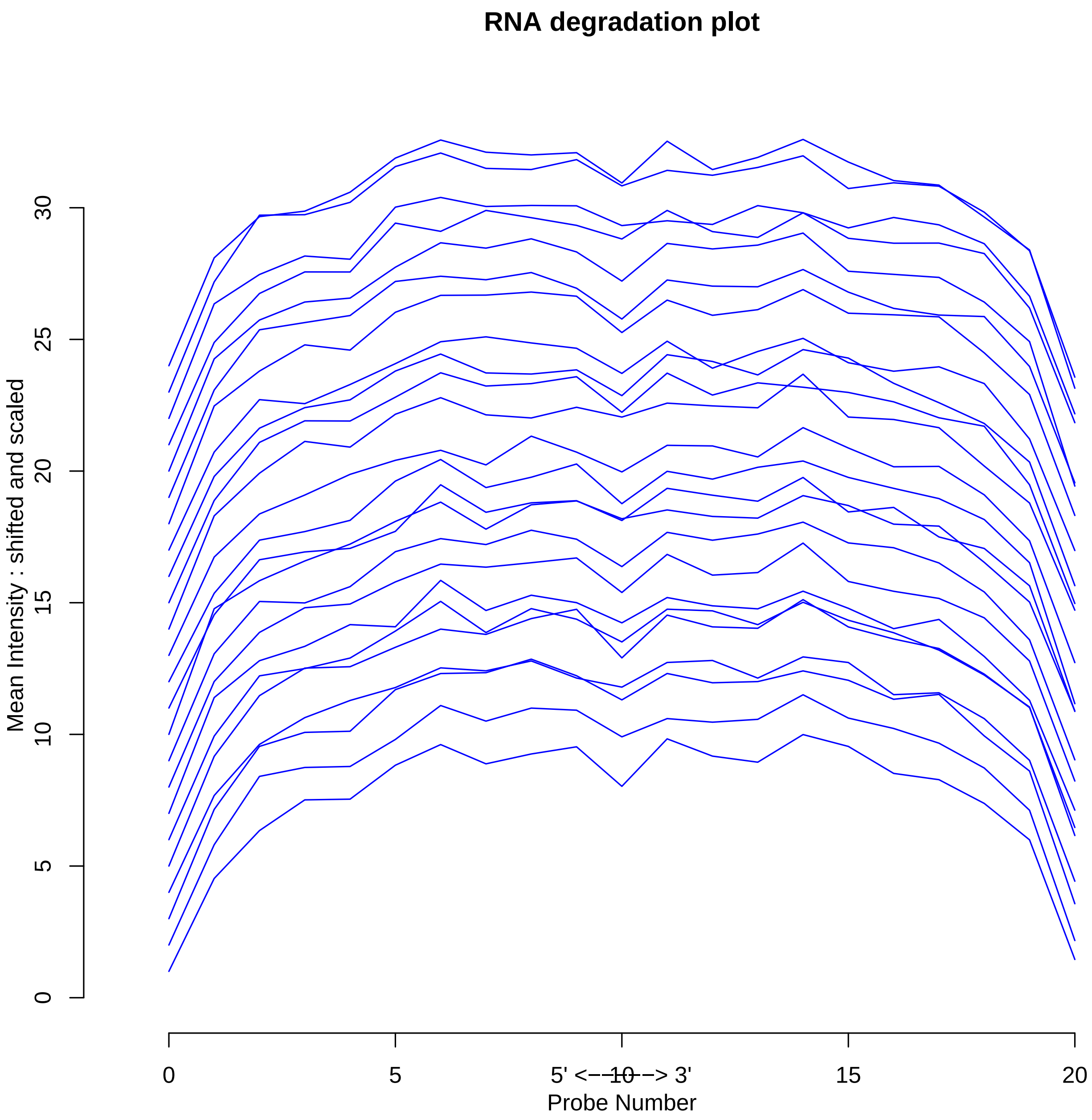

### 3 Relative Log Expression

#### 3.1 Summarized Median QC

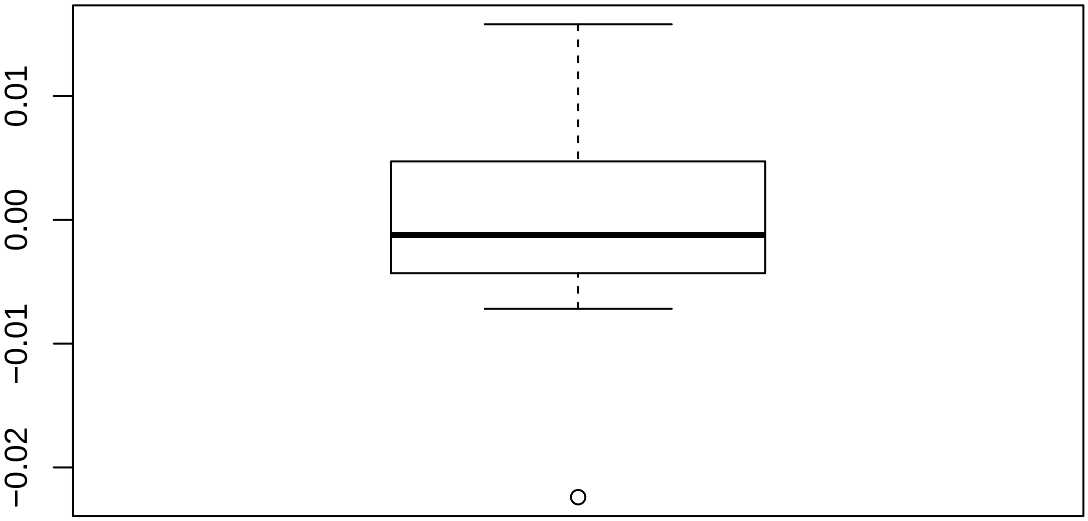

#### 3.2 Discrete QC Plots

Sample Group [1]

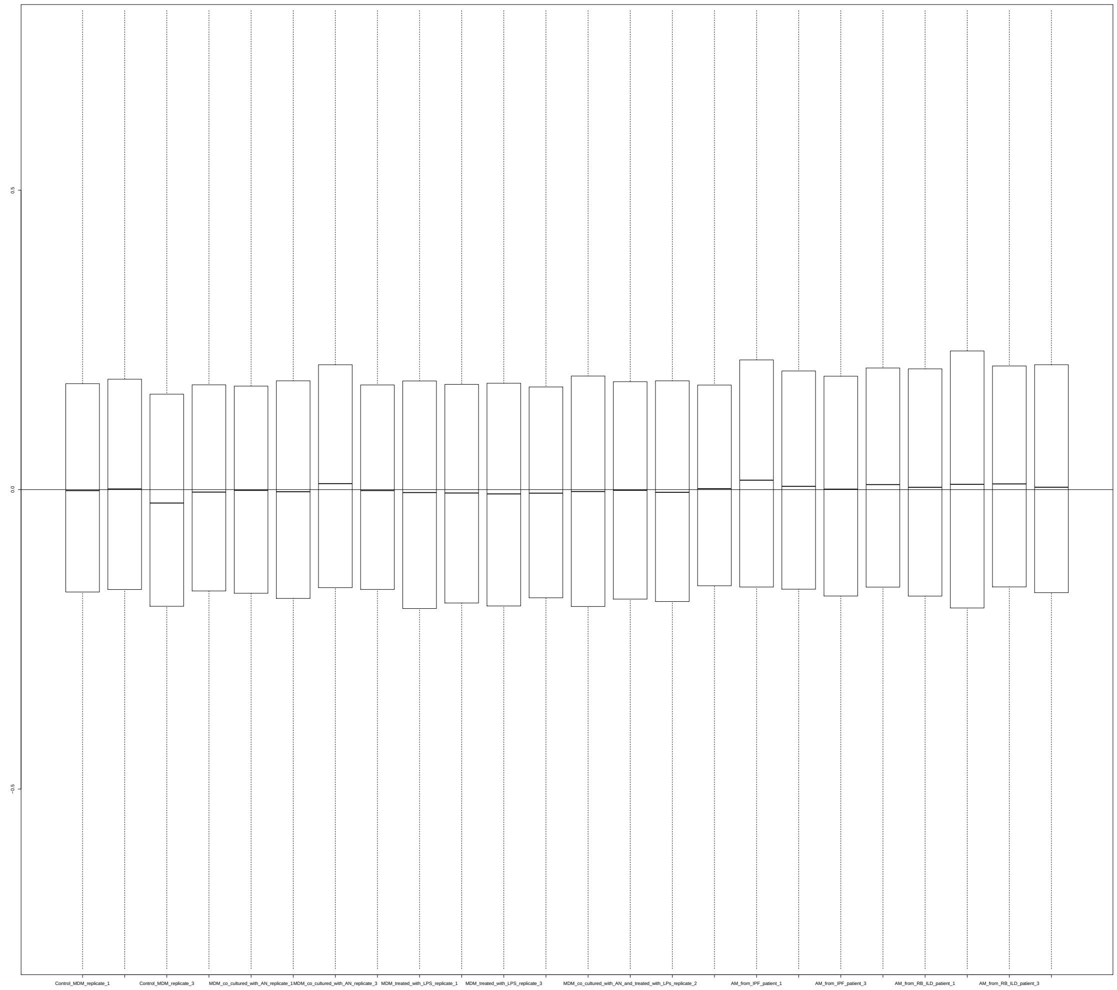

### 4 Normalized Unscaled Standard Errors

#### 4.1 Summarized Median QC

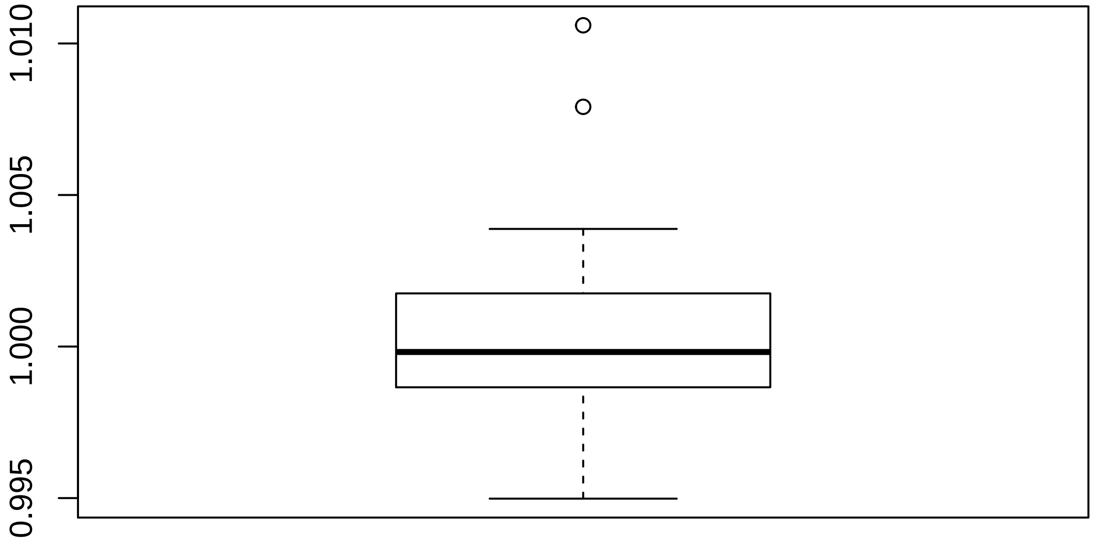

#### 4.2 Discrete QC Plots

Sample Group [1]

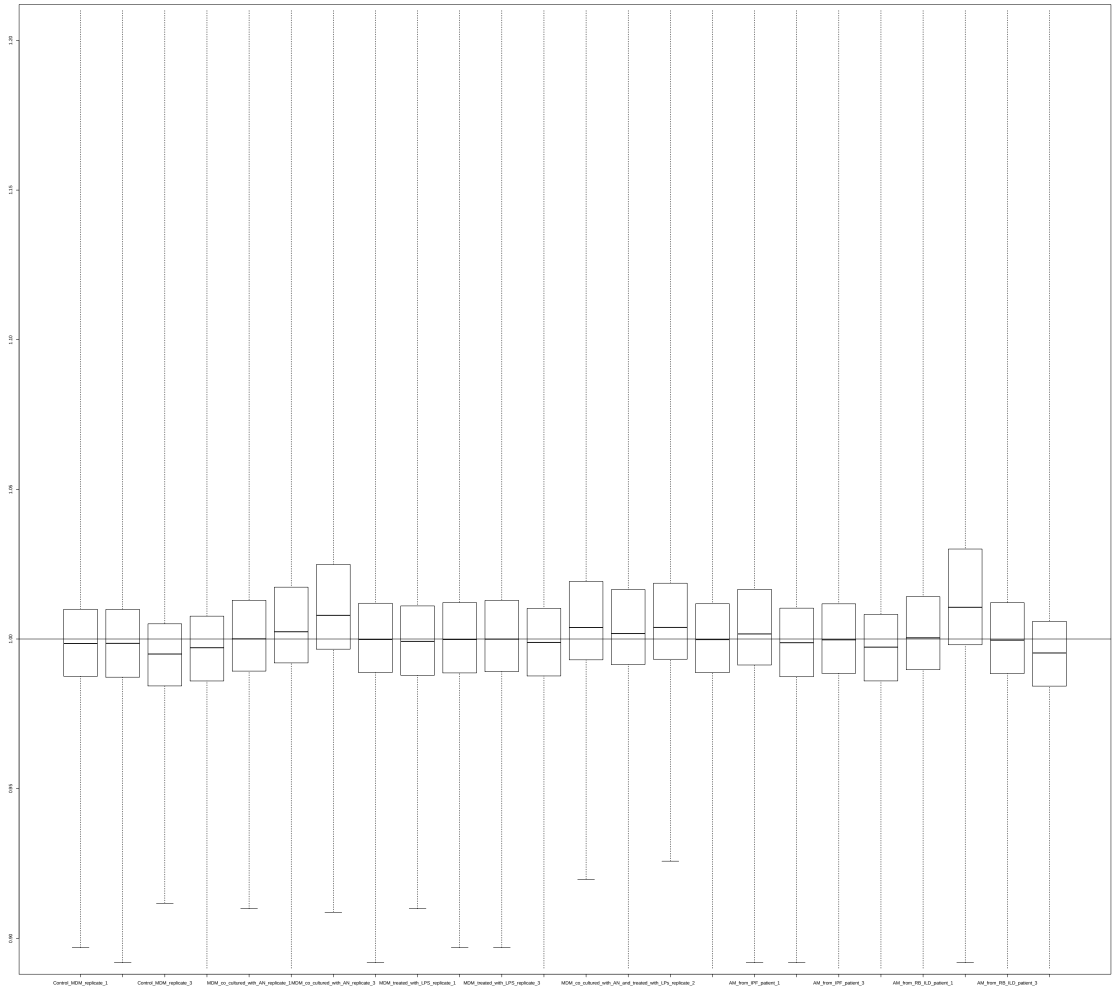
