## Supplementary material for "Curated and harmonised transcriptomics datasets of interstitial lung diseases": GSE110147_eUTOPIA_Affymetrix_QC_Report_2024-02-21.pdf

eUTOPIA

### 1 Outliers Table

| | RLE | NUSE | DEG | SUM |
|---|---|---|---|---|
| GSM2978758 | 0 | NA | 1 | 1 |
| GSM2978759 | 0 | NA | 1 | 1 |
| GSM2978761 | 0 | NA | 1 | 1 |
| GSM2978762 | 0 | NA | 1 | 1 |
| GSM2978763 | 0 | NA | 1 | 1 |
| GSM2978764 | 0 | NA | 1 | 1 |
| GSM2978770 | 0 | NA | 1 | 1 |
| GSM2978772 | 0 | NA | 1 | 1 |
| GSM2978779 | 0 | NA | 1 | 1 |
| GSM2978780 | 0 | NA | 1 | 1 |
| GSM2978786 | 0 | NA | 1 | 1 |
| GSM2978787 | 0 | NA | 1 | 1 |
| GSM2978790 | 1 | NA | 0 | 1 |
| GSM2978791 | 1 | NA | 0 | 1 |
| GSM2978795 | 1 | NA | 0 | 1 |
| GSM2978799 | 1 | NA | 0 | 1 |

#### 1.1 Outliers (All Methods)

| Outliers overall |
|---|
| NA |

#### 1.2 Outliers (At Least One Method)

| Outliers at least 1 |
|---|
| GSM2978758 |
| GSM2978759 |
| GSM2978761 |
| GSM2978762 |
| GSM2978763 |
| GSM2978764 |
| GSM2978770 |
| GSM2978772 |
| GSM2978779 |
| GSM2978780 |
| GSM2978786 |
| GSM2978787 |
| GSM2978790 |
| GSM2978791 |
| GSM2978795 |
| GSM2978799 |

### 2 RNA Degradation

#### 2.1 Summarized Mean QC

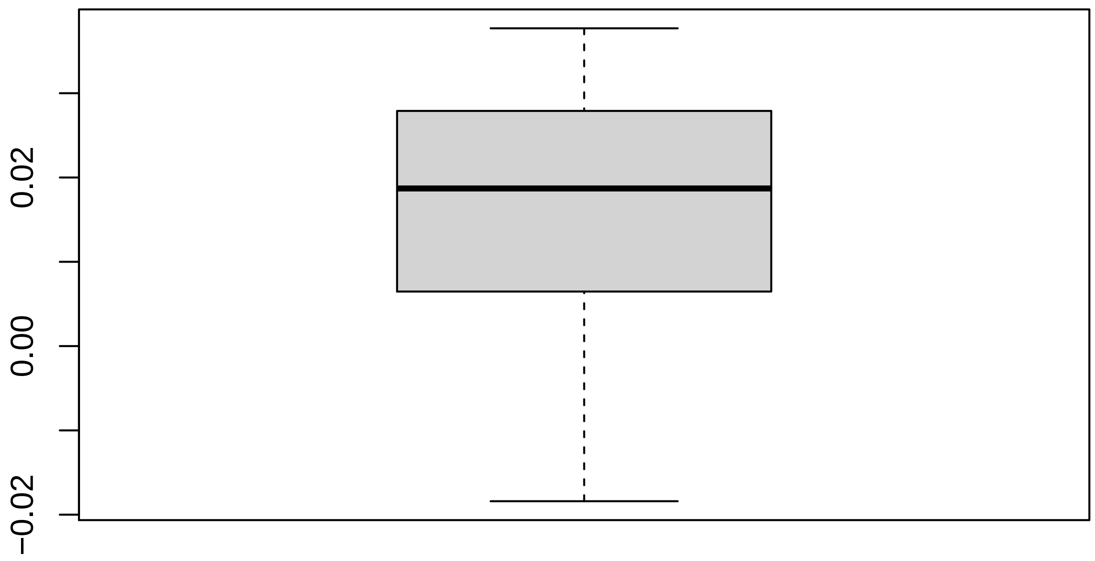

#### 2.2 Discrete QC Plots

**Sample Group [1]**

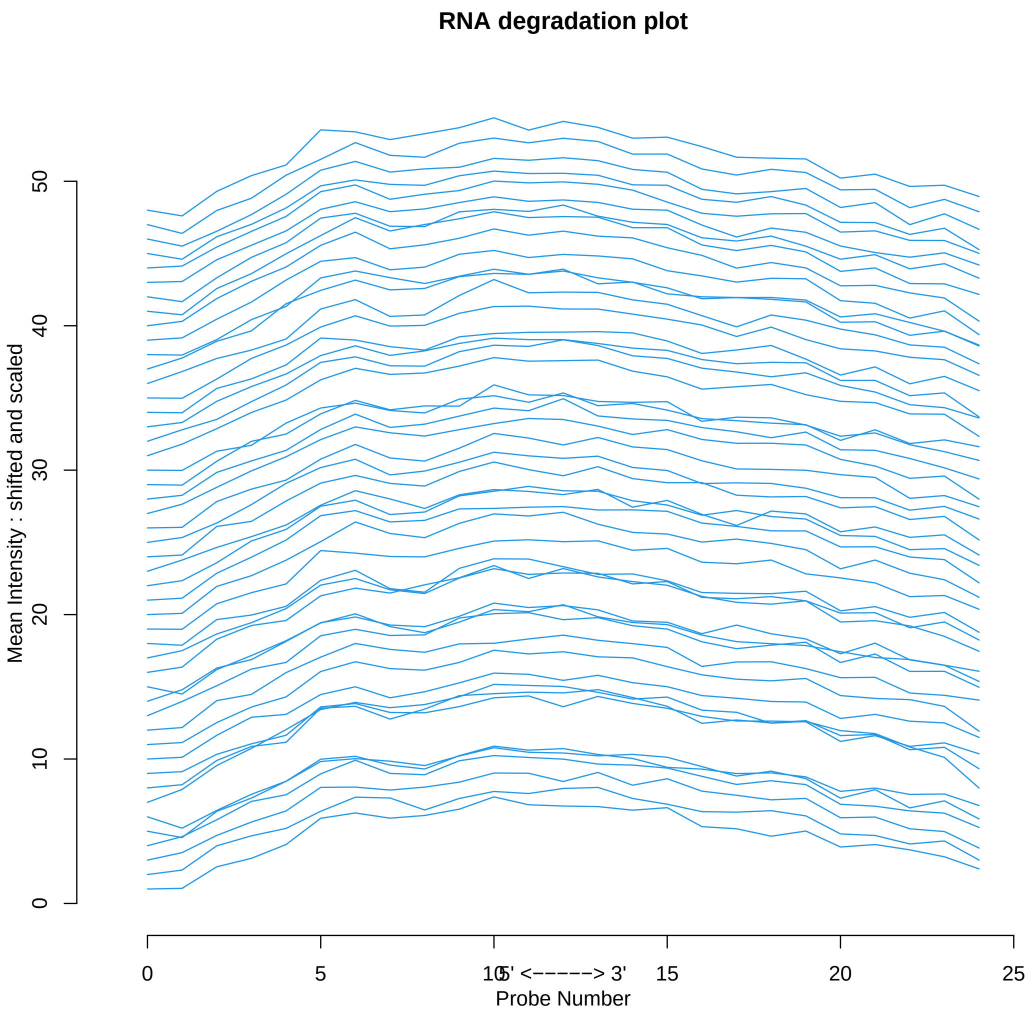

### 3 Relative Log Expression

#### 3.1 Summarized Median QC

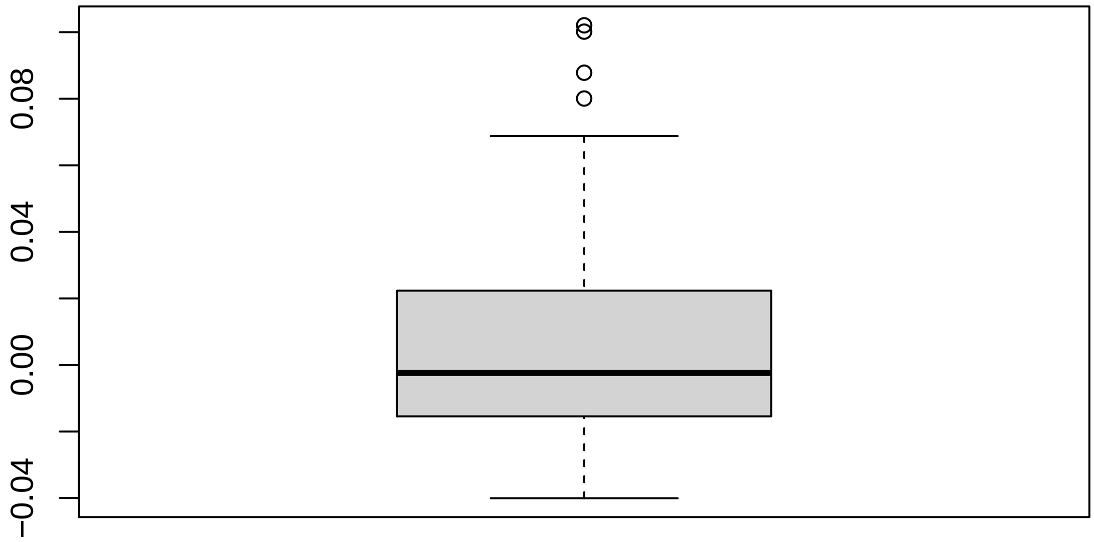

#### 3.2 Discrete QC Plots

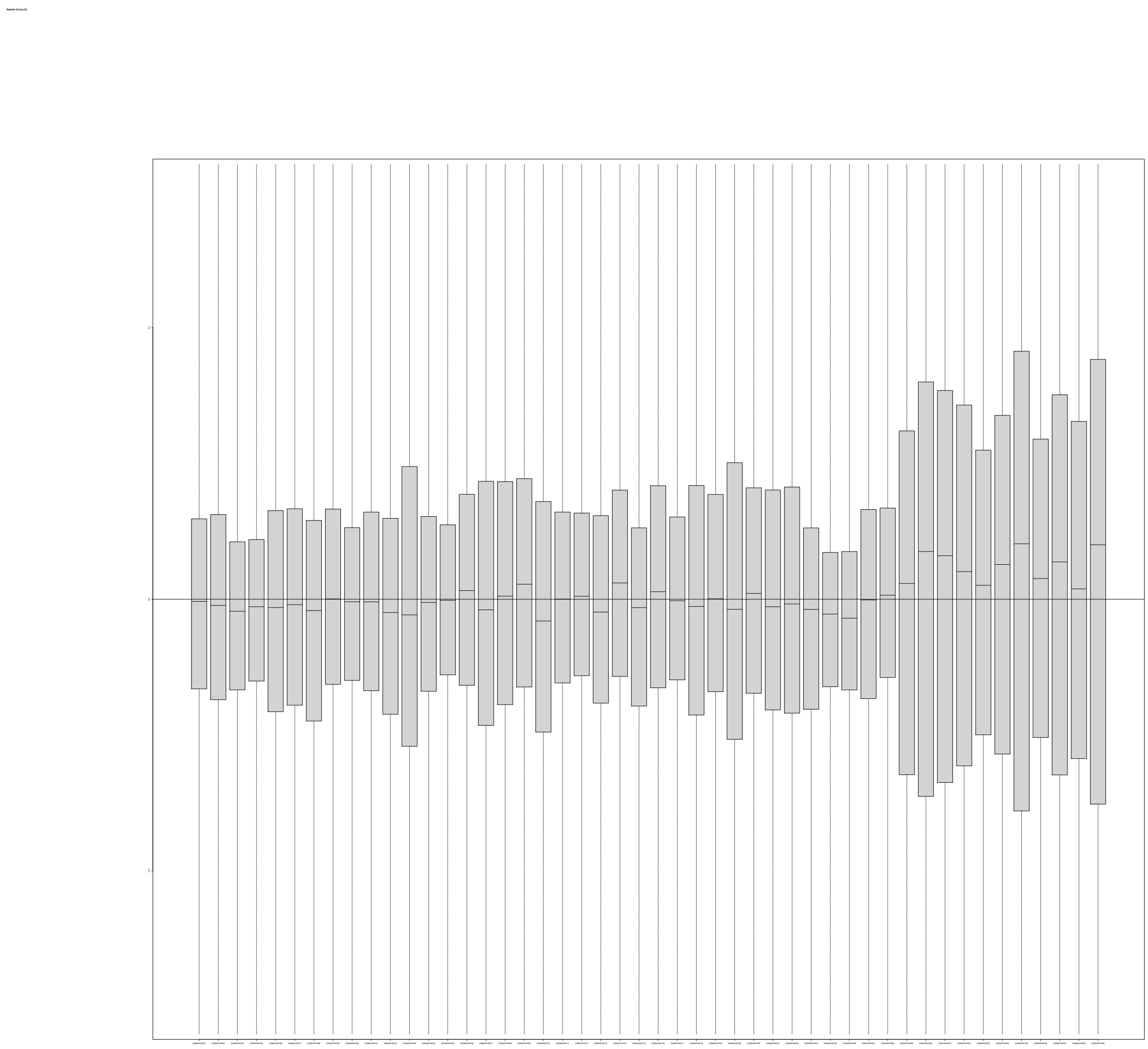
