## Supplementary material for "Curated and harmonised transcriptomics datasets of interstitial lung diseases": GSE129164_eUTOPIA_Affymetrix_QC_Report_2024-02-19.pdf

eUTOPIA

#### 1 Outliers Table

| | RLE | NUSE | DEG | SUM |
|---|---|---|---|---|
| donor_3_TGF_b1 | 0 | 0 | 1 | 1 |
| NL_59_NS | 0 | 0 | 1 | 1 |
| NL59_TGF_b1 | 0 | 1 | 1 | 2 |
| IPF8_NS | 0 | 0 | 1 | 1 |
| IPF8_TGF_b1 | 0 | 0 | 1 | 1 |

##### 1.1 Outliers (All Methods)

| Outliers overall |
|---|
| NL59_TGF_b1 |

##### 1.2 Outliers (At Least One Method)

| Outliers at least 1 |
|---|
| donor_3_TGF_b1 |
| NL_59_NS |
| NL59_TGF_b1 |
| IPF8_NS |

| Outliers at least 1 |
| --- |
| IPF8_TGF_b1 |

### 2 RNA Degradation

#### 2.1 Summarized Mean QC

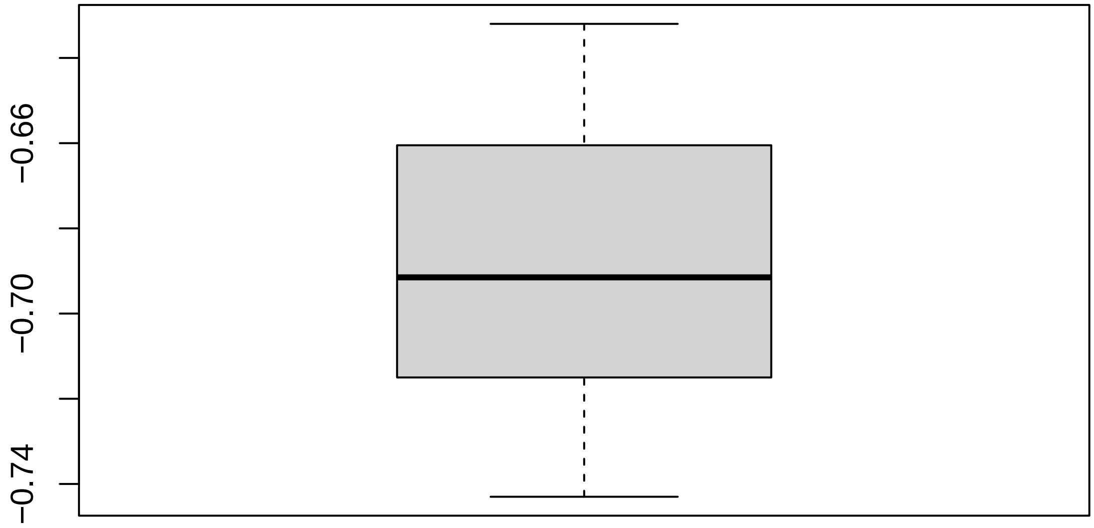

#### 2.2 Discrete QC Plots

**Sample Group [1]**

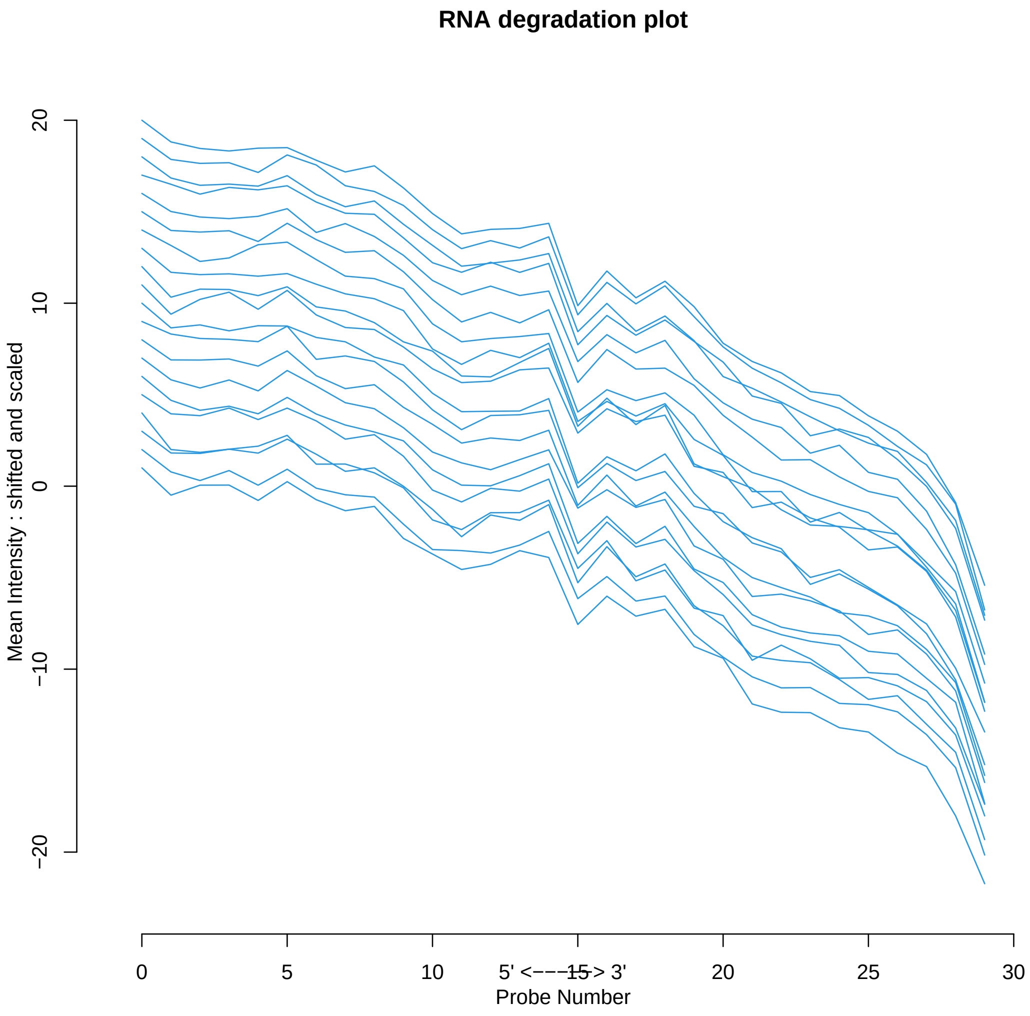

### 3 Relative Log Expression

#### 3.1 Summarized Median QC

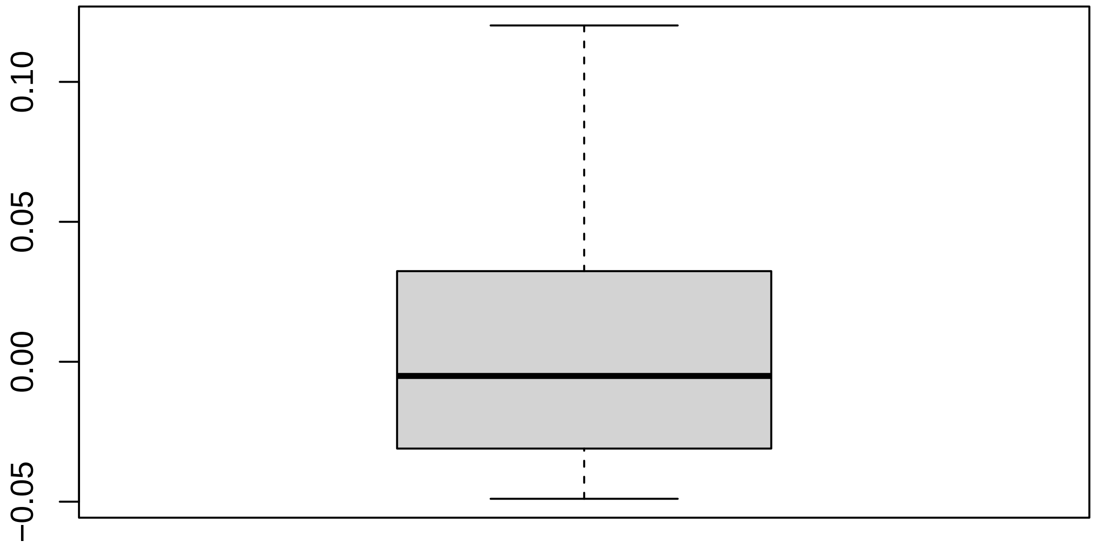

#### 3.2 Discrete QC Plots

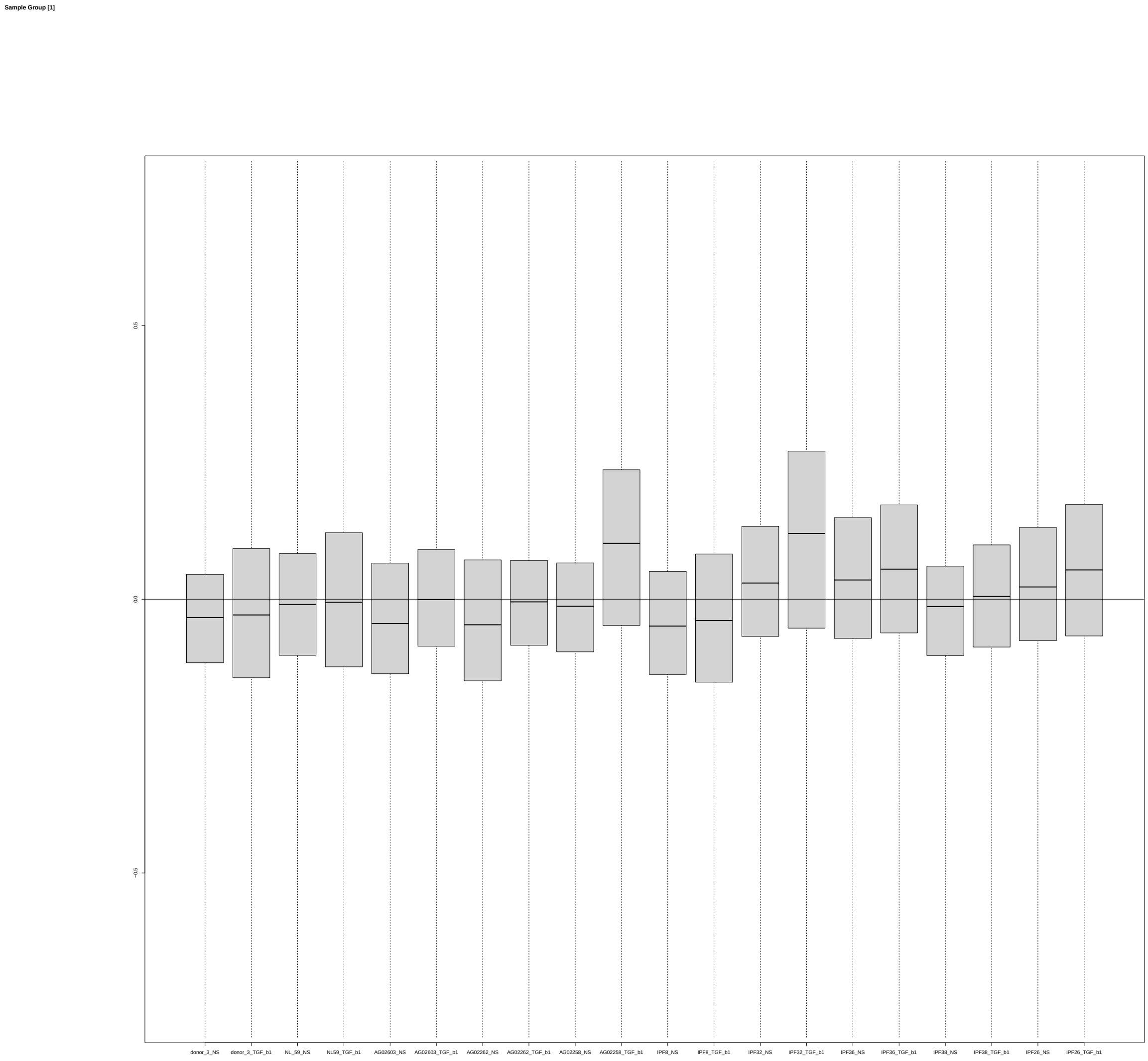

### 4 Normalized Unscaled Standard Errors

#### 4.1 Summarized Median QC

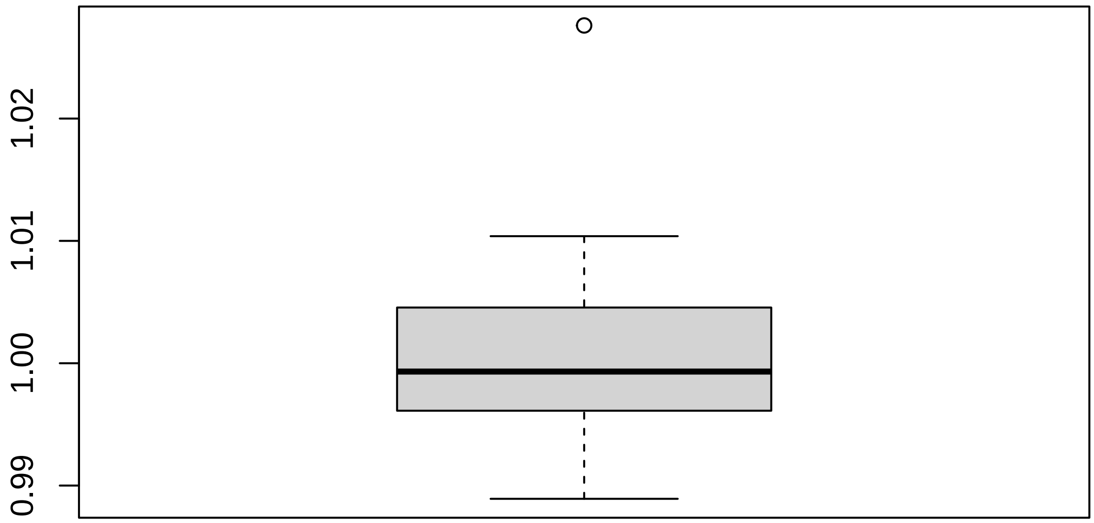

#### 4.2 Discrete QC Plots

Sample Group [1]

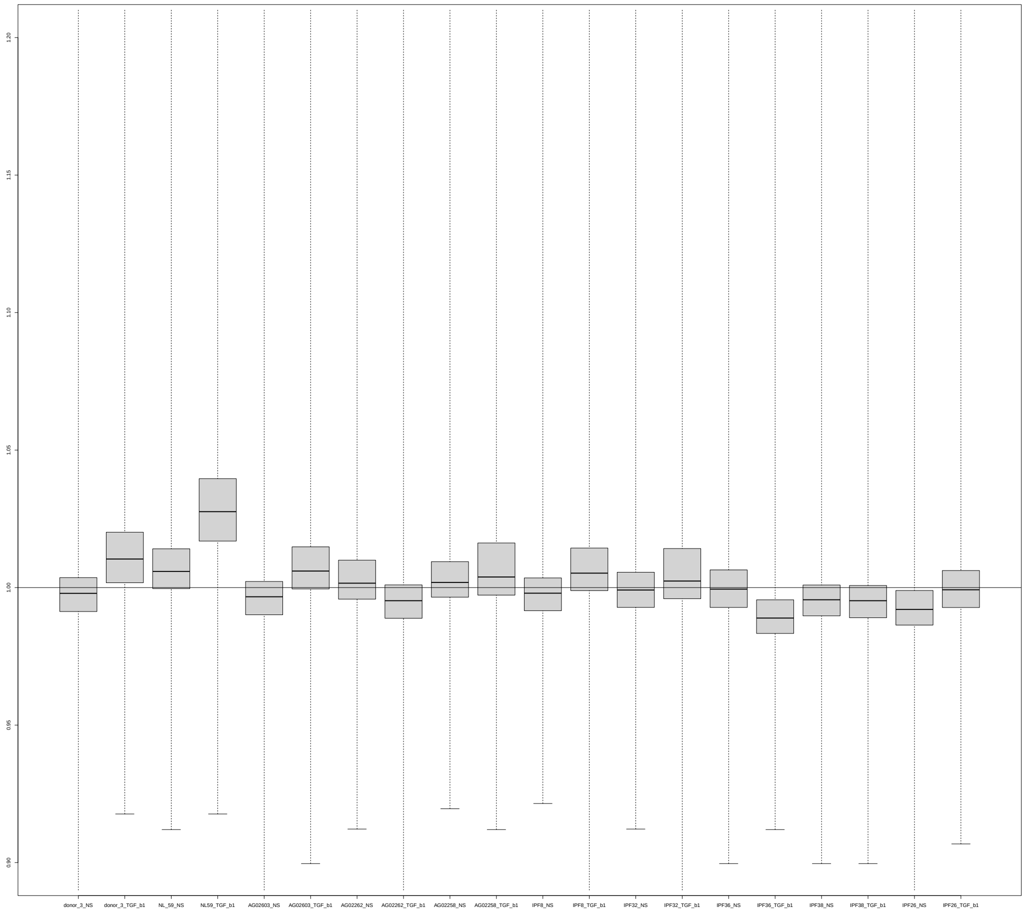
