## Supplementary material for "Curated and harmonised transcriptomics datasets of interstitial lung diseases": GSE144338_eUTOPIA_Affymetrix_QC_Report_2024-02-19.pdf

eUTOPIA

### 1 Outliers Table

| | RLE | NUSE | DEG | SUM |
|---|---|---|---|---|
| Primary__ADC__cell__line__3 | 0 | 1 | 1 | 2 |
| Primary__control__cell__line__3 | 0 | 0 | 1 | 1 |

#### 1.1 Outliers (All Methods)

| Outliers overall |
|---|
| Primary__ADC__cell__line__3 |

#### 1.2 Outliers (At Least One Method)

| Outliers at least 1 |
|---|
| Primary__ADC__cell__line__3 |
| Primary__control__cell__line__3 |

### 2 RNA Degradation

#### 2.1 Summarized Mean QC

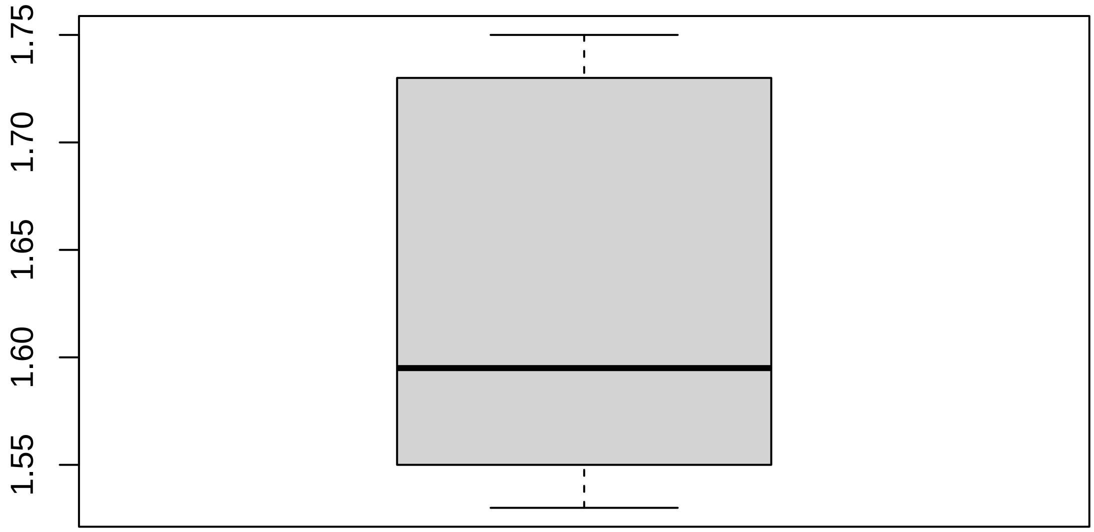

#### 2.2 Discrete QC Plots

**Sample Group [1]**

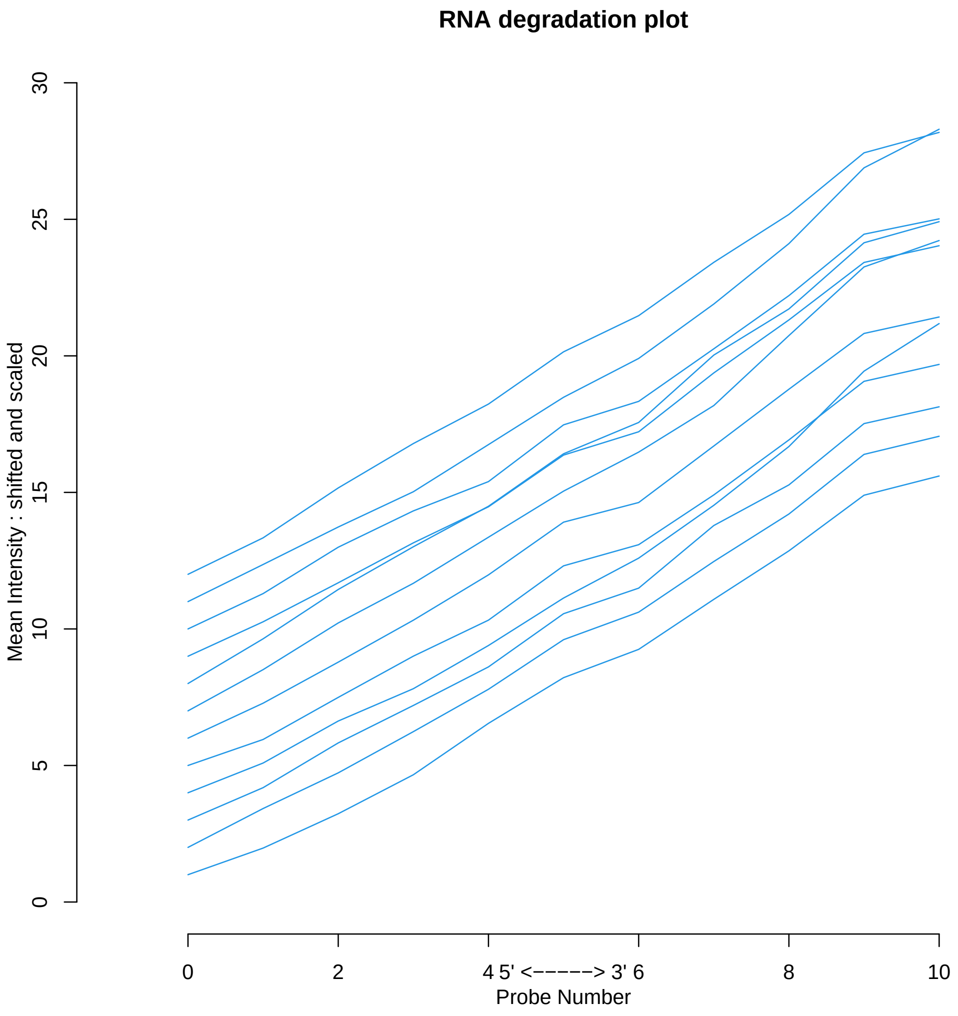

### 3 Relative Log Expression

#### 3.1 Summarized Median QC

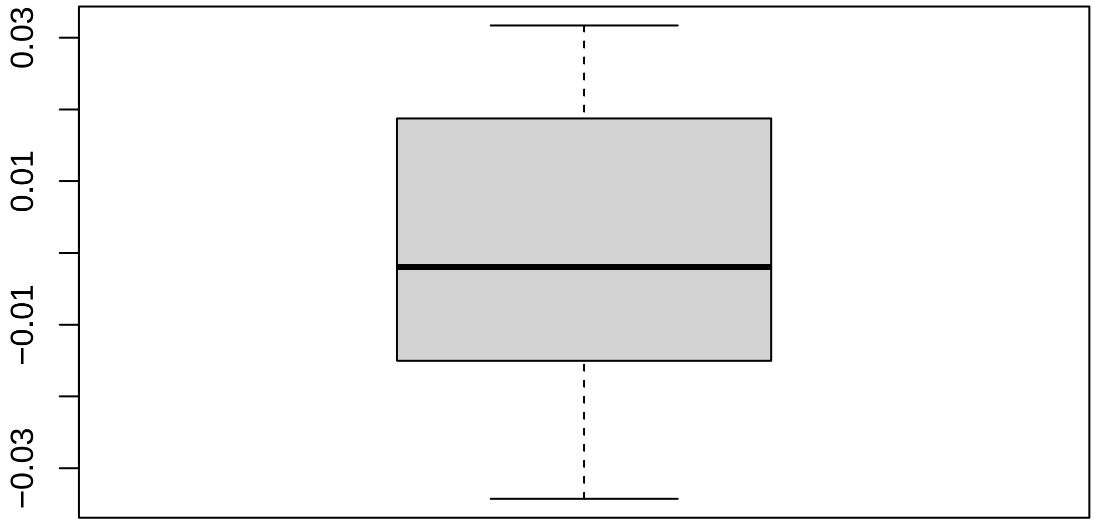

#### 3.2 Discrete QC Plots

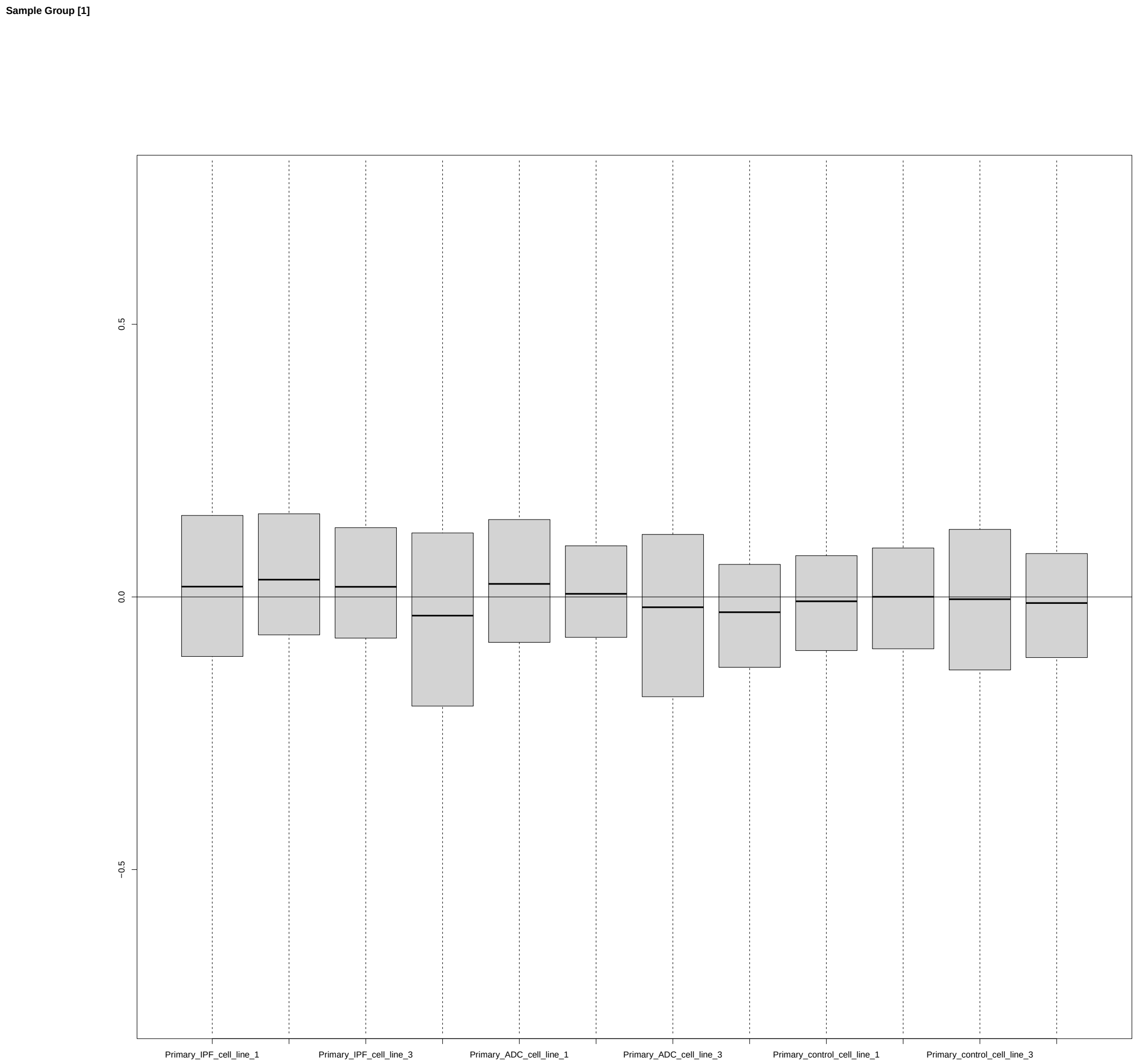

### 4 Normalized Unscaled Standard Errors

#### 4.1 Summarized Median QC

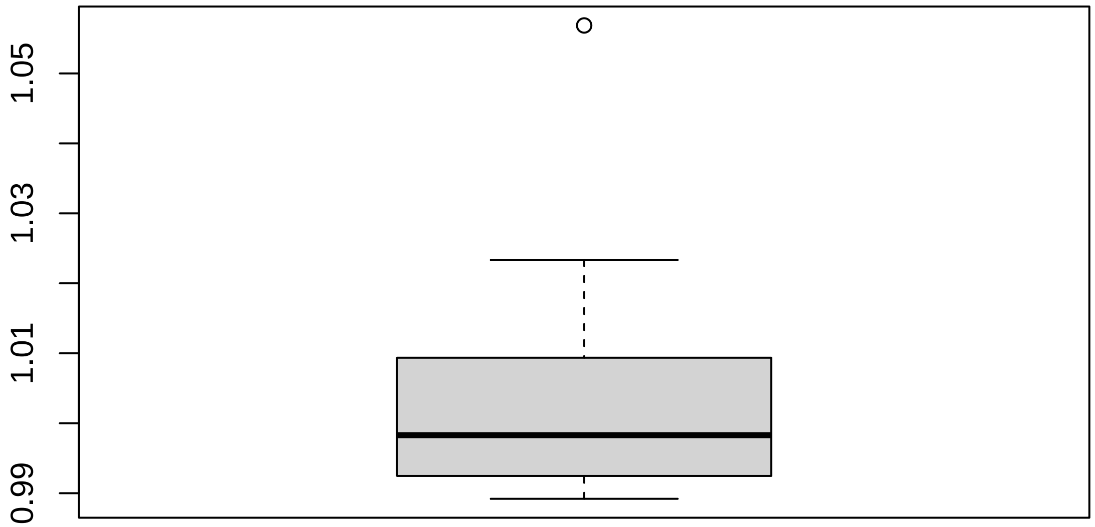

#### 4.2 Discrete QC Plots

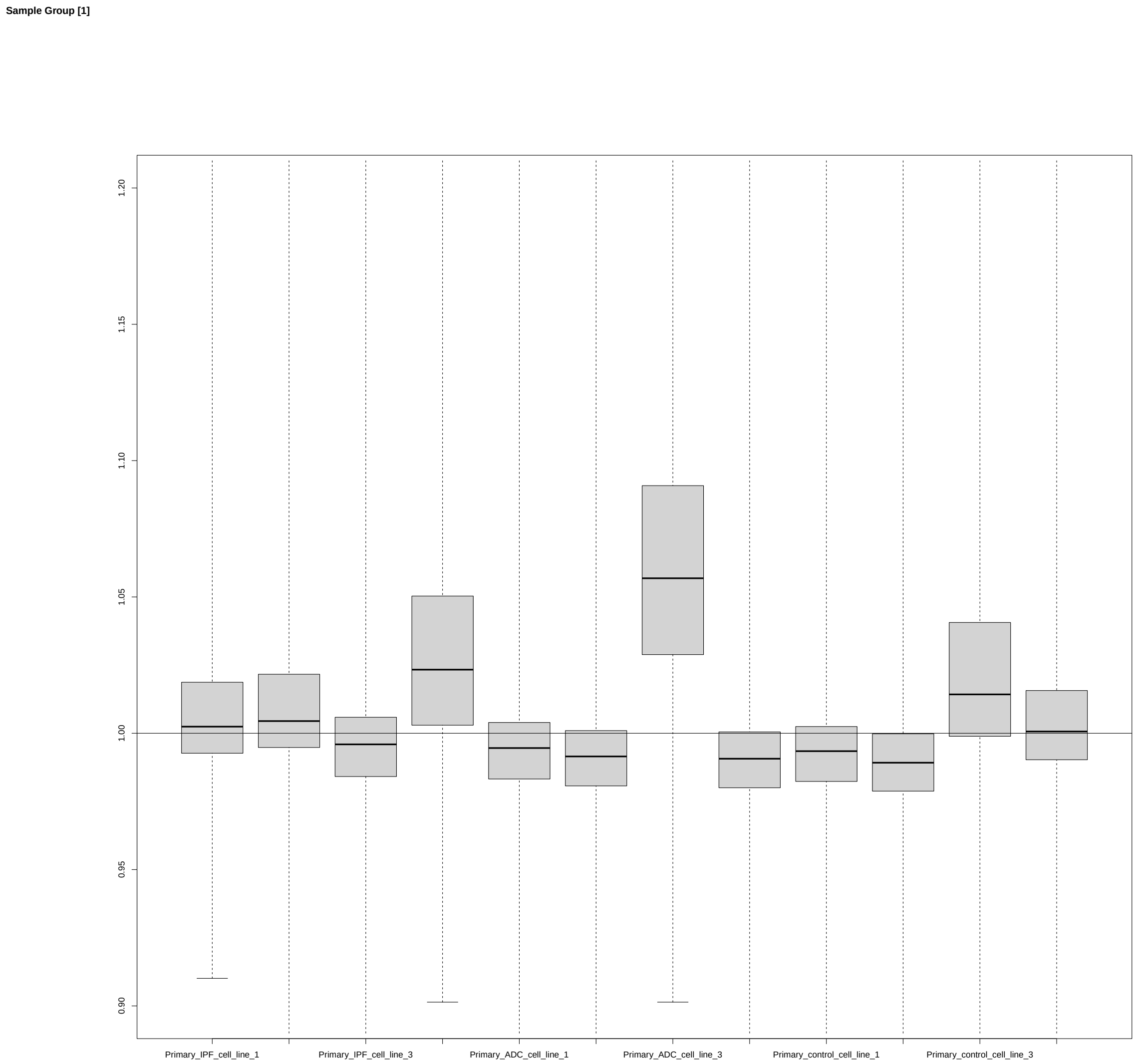
