## Supplementary material for "Curated and harmonised transcriptomics datasets of interstitial lung diseases": Excluded_samples.docx

Several datasets were excluded from the analysis for various reasons. The dataset GSE94060^54,55^ was removed since all the genes had adjusted p-value of 1 in the differential gene expression analysis. GSE45686^56^ was discarded as the sequencing process utilised the Illumina HumanHT-12 V4.0 microarray technique, incompatible with eUTOPIA, which processes only Affymetrix, Agilent, and Illumina methylation data. GSE173355^13^ and GSE176001^57^ were discarded because raw sequencing data was unavailable, and the dataset read alignment was performed with Kallisto pseudoalignment which is not in compliance with our pipeline. GSE76808^58^ was excluded as it only contains samples from patients with SSc-ILD and lacks IPF samples. Finally, GSE79544^59^ was discarded because the use of the NuGEN Ovation FFPE RNA-seq system indicated potentially degraded RNA, leading to an extremely low percentage of assigned fragments during the raw counts extraction phase. In the analysis of GSE70866^60^, only Freiburg samples were included (62 IPF and 20 healthy samples) due to the absence of healthy samples from Sienna and Leuven cohorts. From the GSE11196^61^ dataset, only the total RNA samples were used while the heavy polyribosomial RNA samples were discarded. GSE90010^62^ was included despite a slightly different experimental design involving two sets: one with human monocyte-derived macrophages and another with alveolar macrophages from IPF patients, selected for the lack of suitable datasets containing alveolar macrophages.

Part of the data was not included in the network inference. In dataset GSE49072^63^, only samples from *healthy volunteers* were included in network inference, while samples from *healthy relatives* were excluded. This led to the inclusion of 45 healthy samples instead of the original 61. In GSE129164^64^, only non-stimulated samples were employed for network inference, and TGF-b stimulated samples were omitted. Consequently, 5 IPF samples and 5 healthy samples were utilised, rather than the intended 10 of each condition. The inconsistency in the number of nodes in BAL RNA-Seq networks (Table 3) between healthy and disease nodes arises from specific transcripts in the adjusted expression matrix showing identical expression values across samples, resulting in a null Pearson correlation.
