## Supplementary material for "Curated and harmonised transcriptomics datasets of interstitial lung diseases": SRR8397171_fastqc.html

SRR8397171.fastq.gz FastQC Report 

FastQC Report

Tue 30 Jan 2024  
SRR8397171.fastq.gz

### Summary

- Basic Statistics
- Per base sequence quality
- Per sequence quality scores
- Per base sequence content
- Per sequence GC content
- Per base N content
- Sequence Length Distribution
- Sequence Duplication Levels
- Overrepresented sequences
- Adapter Content

### Basic Statistics

| Measure | Value |
| --- | --- |
| Filename | SRR8397171.fastq.gz |
| File type | Conventional base calls |
| Encoding | Sanger / Illumina 1.9 |
| Total Sequences | 26187137 |
| Sequences flagged as poor quality | 0 |
| Sequence length | 8-371 |
| %GC | 52 |

### Per base sequence quality

### Per sequence quality scores

### Per base sequence content

### Per sequence GC content

### Per base N content

### Sequence Length Distribution

### Sequence Duplication Levels

### Overrepresented sequences

| Sequence | Count | Percentage | Possible Source |
| --- | --- | --- | --- |
| GGGAATGTCTACGTGCGTATGCACGTGGCACTCTCTGCCCGAGGTCCGGG | 251020 | 0.9585622131965017 | No Hit |
| GGGCCACGAGCTGAGTGCGTCCTGTCACTCCACTCCCATGTCCCTTGGGA | 191633 | 0.7317829360269509 | No Hit |
| CCACCTTCTCCATCGGCTCCACTGGCCTCGTGGTGTATGACTACCAGCAG | 179442 | 0.6852295460935649 | No Hit |
| AAGCTCAGGGAGAGCCCTGTTAGGGCCGCCTCTGGCCCTAGTCTCAGACC | 159763 | 0.6100819650502459 | No Hit |
| ACACTGAGGACTCTGTTCCTCCCCTTTCCGCCTAGGGGAAAGTCCCCGGA | 147456 | 0.5630856095494516 | No Hit |
| ATGATGTAGCAGCAGGTGCCAGGGGCTGGCTTGTAGGCGATCAGCAGCTG | 126742 | 0.4839857064176202 | No Hit |
| AAAAGTGATGGACTGCCCATTGCTGGAGAAGACCTTCTCTCCTACTGTCA | 119357 | 0.4557848381821961 | No Hit |
| TGGTTCCTGGCAAGCCCATGTGTGTTGAGAGCTTCTCAGACTATCCACCT | 88999 | 0.33985769425653517 | No Hit |
| ATCAAATCCTGCAGACAAGGGGAGCCCTCAGTCTGCAGGGCTCCATAATG | 80774 | 0.3084491443260865 | No Hit |
| ACTTGATCCCAACTCATCTCTCATTTATTTCGGCTTCTTTTATTCCAGGA | 79162 | 0.30229345040658706 | No Hit |
| ACACCCACCGCAACTGTCTGTCTCATATCACGAACAGCAAAGCGACCCAA | 75365 | 0.28779396541133917 | No Hit |
| ACACTGAGGACTCTGTTCCTCCCCTTTCCGCCTAGGGAAAGTCCCCGGAC | 74238 | 0.28349032580384786 | No Hit |
| TCTGTCTTCTTCAGTTTCGACTTATCGAATTTCTCGATCTCAGCCATATC | 69356 | 0.26484758528586 | No Hit |
| TCTTTCTGGCCTGGAGGCTATCCAGCGTACTCCAAAGATTCAGGTTTACT | 63662 | 0.24310408579601506 | No Hit |
| TCTCTGGGCTTCTATTTCGACCGCGATGATGTGGCTCTGGAAGGCGTGAG | 57328 | 0.21891663834805614 | No Hit |
| AGCGGTTCCGCTGCCCTGAGGCACTCTTCCAGCCTTCCTTCCTGGGCATG | 55038 | 0.21017188706042972 | No Hit |
| TACTCGTGCGCCTCGCTTTGCTTTTCCTCCGCAACCATGTCTGACAAACC | 53494 | 0.20427586261147984 | No Hit |
| ACATAGCAATTCAGGAAATTTGACTTTCCATTCTCTGCTGGATGACGTGA | 52191 | 0.19930013731550722 | No Hit |
| TCACACTTCATGATGGAGTTGAAGGTAGTTTCGTGGATGCCACAGGACTC | 51082 | 0.19506523374433793 | No Hit |
| AATAAAAGCGAAAAGAAATGAAAATGTTACACTACATTAATCCTGGAATA | 46036 | 0.17579623156208332 | No Hit |
| GGCTGCCTGGAGCCCCTGGTGTCCCTGGAGAGCGTGGAGAGAAGGGGGAG | 43680 | 0.1667994481412764 | No Hit |
| CTAAGTGCTGTGTTGTCGTTCCCCCTGCTTAAAATAAAGTTGTTTCTTAA | 38890 | 0.14850802514226735 | No Hit |
| GCTGCTACAGGAGAATAGCAGACAGGTATAGTTAAGAAACAACTTTATTT | 38419 | 0.1467094321918429 | No Hit |
| GGGTACTTCAGGGTCAGGATGCCACGCTTGCTCTGGGCCTCGTCGCCCAC | 33363 | 0.12740224332274275 | No Hit |
| TGTCCATGTCTTACTACTTTGACCGCGATGATGTGGCTTTGAAGAACTTT | 32860 | 0.1254814529744126 | No Hit |
| TGTTCCCTCTCCTCATGAGATTGGTGAAGAAAGTATTTGGCAAAGTTCTT | 31331 | 0.119642708555731 | No Hit |
| GAGCTCCTCATCTAGATGAGCTGGAAGCCCTGGAGGGCCTCTCTCGCCAG | 30898 | 0.11798922501531954 | No Hit |
| TACATCAAAGATTACATGAAATCAATCAAAGGGAAACTTGAAGAACAGAG | 27768 | 0.10603679203266855 | No Hit |
| ACGCTCGTAGCCCTCGCGCTTCTCCTCGGCCAGTTCGCGGAAGAAGTGGC | 27319 | 0.10432220979330424 | No Hit |

### Adapter Content

Produced by FastQC (version 0.11.7)
