## Supplementary material for "Curated and harmonised transcriptomics datasets of interstitial lung diseases": SRR8397178_fastqc.html

SRR8397178.fastq.gz FastQC Report 

FastQC Report

Tue 30 Jan 2024  
SRR8397178.fastq.gz

### Summary

- Basic Statistics
- Per base sequence quality
- Per sequence quality scores
- Per base sequence content
- Per sequence GC content
- Per base N content
- Sequence Length Distribution
- Sequence Duplication Levels
- Overrepresented sequences
- Adapter Content

### Basic Statistics

| Measure | Value |
| --- | --- |
| Filename | SRR8397178.fastq.gz |
| File type | Conventional base calls |
| Encoding | Sanger / Illumina 1.9 |
| Total Sequences | 21323780 |
| Sequences flagged as poor quality | 0 |
| Sequence length | 8-368 |
| %GC | 52 |

### Per base sequence quality

### Per sequence quality scores

### Per base sequence content

### Per sequence GC content

### Per base N content

### Sequence Length Distribution

### Sequence Duplication Levels

### Overrepresented sequences

| Sequence | Count | Percentage | Possible Source |
| --- | --- | --- | --- |
| GGGCCACGAGCTGAGTGCGTCCTGTCACTCCACTCCCATGTCCCTTGGGA | 161364 | 0.7567326243283321 | No Hit |
| AAGCTCAGGGAGAGCCCTGTTAGGGCCGCCTCTGGCCCTAGTCTCAGACC | 161060 | 0.7553069859096276 | No Hit |
| GGGAATGTCTACGTGCGTATGCACGTGGCACTCTCTGCCCGAGGTCCGGG | 102103 | 0.47882223508214766 | No Hit |
| AGCGGTTCCGCTGCCCTGAGGCACTCTTCCAGCCTTCCTTCCTGGGCATG | 100585 | 0.47170342218874883 | No Hit |
| ACACTGAGGACTCTGTTCCTCCCCTTTCCGCCTAGGGGAAAGTCCCCGGA | 87314 | 0.40946773977221673 | No Hit |
| TCACACTTCATGATGGAGTTGAAGGTAGTTTCGTGGATGCCACAGGACTC | 86837 | 0.4072308005428681 | No Hit |
| CCACCTTCTCCATCGGCTCCACTGGCCTCGTGGTGTATGACTACCAGCAG | 71720 | 0.33633811641275607 | No Hit |
| AAAAGTGATGGACTGCCCATTGCTGGAGAAGACCTTCTCTCCTACTGTCA | 69579 | 0.3262976826810256 | No Hit |
| ACACCCACCGCAACTGTCTGTCTCATATCACGAACAGCAAAGCGACCCAA | 67488 | 0.3164917289523715 | No Hit |
| TGGTTCCTGGCAAGCCCATGTGTGTTGAGAGCTTCTCAGACTATCCACCT | 67404 | 0.31609780254720315 | No Hit |
| TCTTTCTGGCCTGGAGGCTATCCAGCGTACTCCAAAGATTCAGGTTTACT | 63843 | 0.29939813672810356 | No Hit |
| TCTCTGGGCTTCTATTTCGACCGCGATGATGTGGCTCTGGAAGGCGTGAG | 58574 | 0.27468863400391486 | No Hit |
| GGGTACTTCAGGGTCAGGATGCCACGCTTGCTCTGGGCCTCGTCGCCCAC | 56938 | 0.2670164483032558 | No Hit |
| ATCAAATCCTGCAGACAAGGGGAGCCCTCAGTCTGCAGGGCTCCATAATG | 53315 | 0.25002602728034146 | No Hit |
| ATGATGTAGCAGCAGGTGCCAGGGGCTGGCTTGTAGGCGATCAGCAGCTG | 52124 | 0.24444071360706218 | No Hit |
| TCGGGCGCCCCAGACACCAGGCGTCATGGTGGGCATGGGCCAGAAGGACT | 46695 | 0.21898087487302909 | No Hit |
| ACATAGCAATTCAGGAAATTTGACTTTCCATTCTCTGCTGGATGACGTGA | 45510 | 0.21342369880011894 | No Hit |
| AGCGGTTCCGGTGTCCGGAGGCGCTGTTCCAGCCTTCCTTCCTGGGTATG | 40614 | 0.19046341689888002 | No Hit |
| ACTTGATCCCAACTCATCTCTCATTTATTTCGGCTTCTTTTATTCCAGGA | 34580 | 0.16216637012762278 | No Hit |
| TCACACTTCATGATGGAGTTGAAGGTGGTCTCGTGGATGCCGCAAGATTC | 34525 | 0.16190844212423874 | No Hit |
| TGAGGCGGAGCACCAGAGAGCCTACCTGGAAGACACATGCGTGGAGTGGC | 34165 | 0.16022018610208885 | No Hit |
| ACACTGAGGACTCTGTTCCTCCCCTTTCCGCCTAGGGAAAGTCCCCGGAC | 34059 | 0.1597230884955669 | No Hit |
| GCTGCTACAGGAGAATAGCAGACAGGTATAGTTAAGAAACAACTTTATTT | 32342 | 0.15167104518992414 | No Hit |
| TCTGTCTTCTTCAGTTTCGACTTATCGAATTTCTCGATCTCAGCCATATC | 32285 | 0.15140373798641704 | No Hit |
| TGTCCATGTCTTACTACTTTGACCGCGATGATGTGGCTTTGAAGAACTTT | 30711 | 0.14402230748957268 | No Hit |
| TGCTATGATTGATATGACCACCACCATTGAACTTCAGTGCAGGCTGAAGA | 29677 | 0.13917326102595318 | No Hit |
| TGTTCCCTCTCCTCATGAGATTGGTGAAGAAAGTATTTGGCAAAGTTCTT | 28809 | 0.13510268817254728 | No Hit |
| TACTCGTGCGCCTCGCTTCGCTTTTCCTCCGCAACCATGTCTGACAAACC | 24882 | 0.11668662873092857 | No Hit |
| TACATCAAAGATTACATGAAATCAATCAAAGGGAAACTTGAAGAACAGAG | 22545 | 0.10572703338713868 | No Hit |
| ACCTACGGCGAGGGCGAGAGCGGCCCCATGGGCAACATCATGATCGATCC | 22122 | 0.10374333256111253 | No Hit |
| ACAGTGTATCTAGCTTTGGAAAACACAGGGGTCTGCCCTGTGAGCTGCTC | 21479 | 0.1007279197215503 | No Hit |

### Adapter Content

Produced by FastQC (version 0.11.7)
