## Supplementary material for "Curated and harmonised transcriptomics datasets of interstitial lung diseases": SRR8397185_fastqc.html

SRR8397185.fastq.gz FastQC Report 

FastQC Report

Tue 30 Jan 2024  
SRR8397185.fastq.gz

### Summary

- Basic Statistics
- Per base sequence quality
- Per sequence quality scores
- Per base sequence content
- Per sequence GC content
- Per base N content
- Sequence Length Distribution
- Sequence Duplication Levels
- Overrepresented sequences
- Adapter Content

### Basic Statistics

| Measure | Value |
| --- | --- |
| Filename | SRR8397185.fastq.gz |
| File type | Conventional base calls |
| Encoding | Sanger / Illumina 1.9 |
| Total Sequences | 16315589 |
| Sequences flagged as poor quality | 0 |
| Sequence length | 8-366 |
| %GC | 53 |

### Per base sequence quality

### Per sequence quality scores

### Per base sequence content

### Per sequence GC content

### Per base N content

### Sequence Length Distribution

### Sequence Duplication Levels

### Overrepresented sequences

| Sequence | Count | Percentage | Possible Source |
| --- | --- | --- | --- |
| AAGCTCAGGGAGAGCCCTGTTAGGGCCGCCTCTGGCCCTAGTCTCAGACC | 355562 | 2.179277744738483 | No Hit |
| GGGCCACGAGCTGAGTGCGTCCTGTCACTCCACTCCCATGTCCCTTGGGA | 341878 | 2.095407036791623 | No Hit |
| GGGAATGTCTACGTGCGTATGCACGTGGCACTCTCTGCCCGAGGTCCGGG | 209975 | 1.2869593613813144 | No Hit |
| ACACTGAGGACTCTGTTCCTCCCCTTTCCGCCTAGGGGAAAGTCCCCGGA | 190945 | 1.17032244438126 | No Hit |
| GCTGCTACAGGAGAATAGCAGACAGGTATAGTTAAGAAACAACTTTATTT | 144861 | 0.8878686512635247 | No Hit |
| CTAAGTGCTGTGTTGTCGTTCCCCCTGCTTAAAATAAAGTTGTTTCTTAA | 138057 | 0.8461662033776409 | No Hit |
| TGGTTCCTGGCAAGCCCATGTGTGTTGAGAGCTTCTCAGACTATCCACCT | 73002 | 0.44743711060630414 | No Hit |
| TCTCTGGGCTTCTATTTCGACCGCGATGATGTGGCTCTGGAAGGCGTGAG | 58617 | 0.35926989825497563 | No Hit |
| ACACCCACCGCAACTGTCTGTCTCATATCACGAACAGCAAAGCGACCCAA | 57269 | 0.3510078612546565 | No Hit |
| TTTACACCTGCTAGGGTGTTCAAAGGTCAGTGCTATAGAAATTCAGTATC | 53955 | 0.3306959987776108 | No Hit |
| AAAGCCAAGAAAACCAACGATGCCAGATACTGAATTTCTATAGCACTGAC | 53790 | 0.3296846960290554 | No Hit |
| AGCTTAATGATGCTTTCTCTGGGCTTTTGGGGGAGGGTGTCCACCAGCTT | 42913 | 0.263018393022771 | No Hit |
| ATCAAGACATGAGGGAGGCAGGGGCTCAGCTGAAGAAGCTGGTGGACACC | 40656 | 0.2491849972440468 | No Hit |
| ACACTGAGGACTCTGTTCCTCCCCTTTCCGCCTAGGGAAAGTCCCCGGAC | 40309 | 0.247058196918297 | No Hit |
| CCTCGGCCTCCCAAAGTGCTGGG | 38681 | 0.23708000979921717 | No Hit |
| CCACCTTCTCCATCGGCTCCACTGGCCTCGTGGTGTATGACTACCAGCAG | 36172 | 0.22170207891360835 | No Hit |
| AAAAGTGATGGACTGCCCATTGCTGGAGAAGACCTTCTCTCCTACTGTCA | 35600 | 0.218196229385283 | No Hit |
| ACTTGATCCCAACTCATCTCTCATTTATTTCGGCTTCTTTTATTCCAGGA | 34601 | 0.21207325092584767 | No Hit |
| AGCGGTTCCGCTGCCCTGAGGCACTCTTCCAGCCTTCCTTCCTGGGCATG | 32170 | 0.197173390430465 | No Hit |
| TTGGCTTTCTCTTTCCTCTTCTCCTCCAGGGTGGCTGTCACTGCCTGGTA | 32026 | 0.1962907989408167 | No Hit |
| GGGGAATGTCTACGTGCGTATGCACGTGGCACTCTCTGCCCGAGGTCCGG | 31447 | 0.19274204565952233 | No Hit |
| GGGCCACGAGCTGAGTGCGTCCTGTCACTCCACTCCCCATGTCCCTTGGG | 31396 | 0.1924294611736052 | No Hit |
| TCTTTCTGGCCTGGAGGCTATCCAGCGTACTCCAAAGATTCAGGTTTACT | 29359 | 0.17994446905962144 | No Hit |
| TGAATCTTGAACTGAGTTCCACTTGTAAACTTCTTGTTTCTTGTGGTTCC | 28057 | 0.17196437100738443 | No Hit |
| TCACACTTCATGATGGAGTTGAAGGTAGTTTCGTGGATGCCACAGGACTC | 27793 | 0.1703462866096958 | No Hit |
| ATCAAATCCTGCAGACAAGGGGAGCCCTCAGTCTGCAGGGCTCCATAATG | 26098 | 0.15995744928362685 | No Hit |
| ATGATGTAGCAGCAGGTGCCAGGGGCTGGCTTGTAGGCGATCAGCAGCTG | 25594 | 0.15686837906985768 | No Hit |
| ACACTGAGGACTCTGTTCCTCCCCCTTTCCGCCTAGGGGAAAGTCCCCGG | 23267 | 0.14260594576144323 | No Hit |
| CTTCCGCATAGCTGCTGTGGTCAAAAAGGAGCCCAGAGTGACAGTTTTCC | 22103 | 0.13547166455345253 | No Hit |
| TGACTGGTGATGCTGATATTGGAGTGCATCTGAGGCTGTTGTAGAAATAA | 21811 | 0.13368196514388786 | No Hit |
| TTCTCCTCAGCTGGAGCAGCAGCAGTGGAGGGGGCAGGACCTCCTGCTGG | 21609 | 0.13244388541535337 | No Hit |
| GGTTATAATCATTGGCAATGTTGTAATCAACAACCCATGTTATTTCTACA | 21344 | 0.13081967191009775 | No Hit |
| GGGTACTTCAGGGTCAGGATGCCACGCTTGCTCTGGGCCTCGTCGCCCAC | 20722 | 0.1270073670034223 | No Hit |
| AGAACGGCGACCGTCAAGGAAAACTGTCACTCTGGGCTCCTTTTTGACCACAGCAGCTATGCGGAAGCAGCTGC | 20358 | 0.12477637184903345 | No Hit |
| ACAGTGTATCTAGCTTTGGAAAACACAGGGGTCTGCCCTGTGAGCTGCTC | 19976 | 0.12243505275843856 | No Hit |
| AACATCTGTGTTGGGGAGAGTGGAGACAGACTGACGCGAGCAGCCAAGGT | 19694 | 0.12070664442454392 | No Hit |
| ATTCTTCCTACCCATGAGCATGGAATGTTCTTCCATTTGTTTGTATCCTC | 19673 | 0.12057793316563688 | No Hit |
| CCGCCTCCGCCGCAGACGCCGCCGCGATGCGCTACGTCGCCTCCTACCTG | 19153 | 0.11739079723079565 | No Hit |
| TTTGGGAGGCCGAGGCGGGTGG | 18999 | 0.1164469146654773 | No Hit |
| ACGCTCGTAGCCCTCGCGCTTCTCCTCGGCCAGTTCGCGGAAGAAGTGGC | 18510 | 0.11344978106521315 | No Hit |
| ACATAGCAATTCAGGAAATTTGACTTTCCATTCTCTGCTGGATGACGTGA | 17863 | 0.10948424846936265 | No Hit |
| ACGCTCGTAGCCCTCGCGCTTCTCCTCGGCCAATTCGCGGAAGAAGTGGC | 17614 | 0.10795810068517907 | No Hit |
| TACATCAAAGATTACATGAAATCAATCAAAGGGAAACTTGAAGAACAGAG | 17480 | 0.10713680027120076 | No Hit |
| GGCCTTCTTCTGGCGCAGAGGAAGCAGGCGCATCTGGGTGTGGGCAATGA | 17438 | 0.10687937775338664 | No Hit |
| TGTTCCCTCTCCTCATGAGATTGGTGAAGAAAGTATTTGGCAAAGTTCTT | 17349 | 0.10633388717992344 | No Hit |
| CAGCTGGAGAAGGACTTCAGCAGCATGAAGAAGTACTGCCAAGTCATCCG | 17181 | 0.10530419710866705 | No Hit |
| AGAGGCCACAGTAAGGCCCATCCGAGGTCCTGAGGAGATCTTTTCTCTCT | 17160 | 0.10517548584976001 | No Hit |
| TCGGGCGCCCCAGACACCAGGCGTCATGGTGGGCATGGGCCAGAAGGACT | 16991 | 0.1041396666709366 | No Hit |
| CGGTTGTACTTGCGGATGTAGTGCAGATAGTCTCGGCGGATGACAATGGT | 16498 | 0.10111801664040446 | No Hit |

### Adapter Content

Produced by FastQC (version 0.11.7)
