## Supplementary material for "Curated and harmonised transcriptomics datasets of interstitial lung diseases": SRR8397190_fastqc.html

SRR8397190.fastq.gz FastQC Report 

FastQC Report

Tue 30 Jan 2024  
SRR8397190.fastq.gz

### Summary

- Basic Statistics
- Per base sequence quality
- Per sequence quality scores
- Per base sequence content
- Per sequence GC content
- Per base N content
- Sequence Length Distribution
- Sequence Duplication Levels
- Overrepresented sequences
- Adapter Content

### Basic Statistics

| Measure | Value |
| --- | --- |
| Filename | SRR8397190.fastq.gz |
| File type | Conventional base calls |
| Encoding | Sanger / Illumina 1.9 |
| Total Sequences | 24579447 |
| Sequences flagged as poor quality | 0 |
| Sequence length | 8-368 |
| %GC | 53 |

### Per base sequence quality

### Per sequence quality scores

### Per base sequence content

### Per sequence GC content

### Per base N content

### Sequence Length Distribution

### Sequence Duplication Levels

### Overrepresented sequences

| Sequence | Count | Percentage | Possible Source |
| --- | --- | --- | --- |
| AAGCTCAGGGAGAGCCCTGTTAGGGCCGCCTCTGGCCCTAGTCTCAGACC | 664171 | 2.702139718603108 | No Hit |
| GGGCCACGAGCTGAGTGCGTCCTGTCACTCCACTCCCATGTCCCTTGGGA | 580190 | 2.3604680772516975 | No Hit |
| GGGAATGTCTACGTGCGTATGCACGTGGCACTCTCTGCCCGAGGTCCGGG | 172091 | 0.7001418705636462 | No Hit |
| ACACTGAGGACTCTGTTCCTCCCCTTTCCGCCTAGGGGAAAGTCCCCGGA | 164651 | 0.6698726785838591 | No Hit |
| TCTCTGGGCTTCTATTTCGACCGCGATGATGTGGCTCTGGAAGGCGTGAG | 158771 | 0.64595025266435 | No Hit |
| GCTGCTACAGGAGAATAGCAGACAGGTATAGTTAAGAAACAACTTTATTT | 144615 | 0.58835741910711 | No Hit |
| GGGCCACGAGCTGAGTGCGTCCTGTCACTCCACTCCCCATGTCCCTTGGG | 120380 | 0.48975878098477965 | No Hit |
| CTAAGTGCTGTGTTGTCGTTCCCCCTGCTTAAAATAAAGTTGTTTCTTAA | 113109 | 0.4601771553281894 | No Hit |
| TGGTTCCTGGCAAGCCCATGTGTGTTGAGAGCTTCTCAGACTATCCACCT | 88053 | 0.3582383281446487 | No Hit |
| ACGCTCGTAGCCCTCGCGCTTCTCCTCGGCCAATTCGCGGAAGAAGTGGC | 84022 | 0.3418384473824818 | No Hit |
| ACTTGATCCCAACTCATCTCTCATTTATTTCGGCTTCTTTTATTCCAGGA | 82665 | 0.33631757459799644 | No Hit |
| ACGCTCGTAGCCCTCGCGCTTCTCCTCGGCCAGTTCGCGGAAGAAGTGGC | 82587 | 0.33600023629498255 | No Hit |
| AAAAGTGATGGACTGCCCATTGCTGGAGAAGACCTTCTCTCCTACTGTCA | 74536 | 0.3032452276082534 | No Hit |
| AGCGGTTCCGCTGCCCTGAGGCACTCTTCCAGCCTTCCTTCCTGGGCATG | 71823 | 0.2922075504790649 | No Hit |
| ATCAAATCCTGCAGACAAGGGGAGCCCTCAGTCTGCAGGGCTCCATAATG | 66302 | 0.26974569444137614 | No Hit |
| TCACACTTCATGATGGAGTTGAAGGTAGTTTCGTGGATGCCACAGGACTC | 63588 | 0.2587039488724055 | No Hit |
| ACACCCACCGCAACTGTCTGTCTCATATCACGAACAGCAAAGCGACCCAA | 63571 | 0.25863478539610757 | No Hit |
| TCTCTGGCTTCTATTTCGACCGCGATGATGTGGCTCTGGAAGGCGTGAGC | 56729 | 0.23079852040609378 | No Hit |
| CCACCTTCTCCATCGGCTCCACTGGCCTCGTGGTGTATGACTACCAGCAG | 52902 | 0.21522860135950167 | No Hit |
| TTTACACCTGCTAGGGTGTTCAAAGGTCAGTGCTATAGAAATTCAGTATC | 51945 | 0.21133510448790813 | No Hit |
| AATAAAAGCGAAAAGAAATGAAAATGTTACACTACATTAATCCTGGAATA | 50337 | 0.20479305331808317 | No Hit |
| AAAGCCAAGAAAACCAACGATGCCAGATACTGAATTTCTATAGCACTGAC | 48605 | 0.1977465156152618 | No Hit |
| ACACTGAGGACTCTGTTCCTCCCCTTTCCGCCTAGGGAAAGTCCCCGGAC | 48440 | 0.19707522305119396 | No Hit |
| ACACTGAGGACTCTGTTCCTCCCCCTTTCCGCCTAGGGGAAAGTCCCCGG | 42982 | 0.1748696787197857 | No Hit |
| TGACTGGTGATGCTGATATTGGAGTGCATCTGAGGCTGTTGTAGAAATAA | 41710 | 0.16969462331678983 | No Hit |
| ATGATGTAGCAGCAGGTGCCAGGGGCTGGCTTGTAGGCGATCAGCAGCTG | 40467 | 0.16463755266747865 | No Hit |
| GGGTACTTCAGGGTCAGGATGCCACGCTTGCTCTGGGCCTCGTCGCCCAC | 38391 | 0.15619147167957032 | No Hit |
| AGCGGTTCCGGTGTCCGGAGGCGCTGTTCCAGCCTTCCTTCCTGGGTATG | 38175 | 0.15531268868660877 | No Hit |
| TCACACTTCATGATGGAGTTGAAGGTGGTCTCGTGGATGCCGCAAGATTC | 34942 | 0.142159422870661 | No Hit |
| GGGGCCACGAGCTGAGTGCGTCCTGTCACTCCACTCCCATGTCCCTTGGG | 33562 | 0.13654497597118437 | No Hit |
| TGAATCTTGAACTGAGTTCCACTTGTAAACTTCTTGTTTCTTGTGGTTCC | 31512 | 0.12820467441761405 | No Hit |
| CCGCCTCCGCCGCAGACGCCGCCGCGATGCGCTACGTCGCCTCCTACCTG | 30158 | 0.12269600695247539 | No Hit |
| TCGGGCGCCCCAGACACCAGGCGTCATGGTGGGCATGGGCCAGAAGGACT | 29965 | 0.1219107980745051 | No Hit |
| CAGCTGGAGAAGGACTTCAGCAGCATGAAGAAGTACTGCCAAGTCATCCG | 27632 | 0.11241912806256382 | No Hit |
| GGTTATAATCATTGGCAATGTTGTAATCAACAACCCATGTTATTTCTACA | 27584 | 0.1122238429530168 | No Hit |
| GGCCTTCTTCTGGCGCAGAGGAAGCAGGCGCATCTGGGTGTGGGCAATGA | 27384 | 0.11141015499657092 | No Hit |
| TGTCCATGTCTTACTACTTTGACCGCGATGATGTGGCTTTGAAGAACTTT | 27107 | 0.11028319717689336 | No Hit |
| TCTGCAATGCAATCACAATGCCCAAACTAGACCTGCCATTTCTCACACAC | 25945 | 0.10555567014994276 | No Hit |
| ACAAGCAGTTCACAGTAGAACACTGCAGACTGCAGGGCATTTTCAGGCTA | 25784 | 0.10490065134500383 | No Hit |
| GGTTATATCATTGGCAATGTTGTAATCAACAACCCATGTTATTTCTACAA | 25366 | 0.10320004351603192 | No Hit |
| AGAGGCCACAGTAAGGCCCATCCGAGGTCCTGAGGAGATCTTTTCTCTCT | 25339 | 0.10309019564191171 | No Hit |
| TGTTCCCTCTCCTCATGAGATTGGTGAAGAAAGTATTTGGCAAAGTTCTT | 25121 | 0.1022032757693857 | No Hit |

### Adapter Content

Produced by FastQC (version 0.11.7)
