## Supplementary material for "Curated and harmonised transcriptomics datasets of interstitial lung diseases": SRR8397204_fastqc.html

SRR8397204.fastq.gz FastQC Report 

FastQC Report

Tue 30 Jan 2024  
SRR8397204.fastq.gz

### Summary

- Basic Statistics
- Per base sequence quality
- Per sequence quality scores
- Per base sequence content
- Per sequence GC content
- Per base N content
- Sequence Length Distribution
- Sequence Duplication Levels
- Overrepresented sequences
- Adapter Content

### Basic Statistics

| Measure | Value |
| --- | --- |
| Filename | SRR8397204.fastq.gz |
| File type | Conventional base calls |
| Encoding | Sanger / Illumina 1.9 |
| Total Sequences | 25169033 |
| Sequences flagged as poor quality | 0 |
| Sequence length | 8-341 |
| %GC | 53 |

### Per base sequence quality

### Per sequence quality scores

### Per base sequence content

### Per sequence GC content

### Per base N content

### Sequence Length Distribution

### Sequence Duplication Levels

### Overrepresented sequences

| Sequence | Count | Percentage | Possible Source |
| --- | --- | --- | --- |
| GGGCCACGAGCTGAGTGCGTCCTGTCACTCCACTCCCATGTCCCTTGGGA | 237280 | 0.9427457940080575 | No Hit |
| AAGCTCAGGGAGAGCCCTGTTAGGGCCGCCTCTGGCCCTAGTCTCAGACC | 224056 | 0.8902050388666104 | No Hit |
| TCTCTGGGCTTCTATTTCGACCGCGATGATGTGGCTCTGGAAGGCGTGAG | 144344 | 0.5734983938397633 | No Hit |
| GGGAATGTCTACGTGCGTATGCACGTGGCACTCTCTGCCCGAGGTCCGGG | 127934 | 0.5082992262753996 | No Hit |
| CCACCTTCTCCATCGGCTCCACTGGCCTCGTGGTGTATGACTACCAGCAG | 115612 | 0.459342240124998 | No Hit |
| AGCGGTTCCGCTGCCCTGAGGCACTCTTCCAGCCTTCCTTCCTGGGCATG | 103609 | 0.41165268447142966 | No Hit |
| TGGTTCCTGGCAAGCCCATGTGTGTTGAGAGCTTCTCAGACTATCCACCT | 92251 | 0.36652580176600347 | No Hit |
| ACACTGAGGACTCTGTTCCTCCCCTTTCCGCCTAGGGGAAAGTCCCCGGA | 91439 | 0.3632996150467918 | No Hit |
| TCACACTTCATGATGGAGTTGAAGGTAGTTTCGTGGATGCCACAGGACTC | 89607 | 0.35602082924679707 | No Hit |
| ATGATGTAGCAGCAGGTGCCAGGGGCTGGCTTGTAGGCGATCAGCAGCTG | 84107 | 0.3341685793013979 | No Hit |
| ACACCCACCGCAACTGTCTGTCTCATATCACGAACAGCAAAGCGACCCAA | 76550 | 0.30414358787641943 | No Hit |
| TCTTTCTGGCCTGGAGGCTATCCAGCGTACTCCAAAGATTCAGGTTTACT | 70273 | 0.2792042109841884 | No Hit |
| AAAAGTGATGGACTGCCCATTGCTGGAGAAGACCTTCTCTCCTACTGTCA | 69476 | 0.27603762131028237 | No Hit |
| AAGACCAACGTCAAGGCCGCCTGGGGTAAGGTCGGCGCGCACGCTGGCGA | 68887 | 0.27369744399794776 | No Hit |
| ACCAATGTCATAGGGTGCAATATCTACAATAGGTAGTCTCACAGCCTTGC | 63041 | 0.25047048887416534 | No Hit |
| AGAGCATGACCGATGGATTCCAGTTCGAGTATGGCGGCCAGGGCTCCGAC | 63023 | 0.2503989724197986 | No Hit |
| TTGGTGGTGGGGAAGGACAGGAACATCCTCTCCAGGGCCTCCGCACCATA | 60711 | 0.2412130811700235 | No Hit |
| GGGTACTTCAGGGTCAGGATGCCACGCTTGCTCTGGGCCTCGTCGCCCAC | 58748 | 0.23341381450769283 | No Hit |
| TCTGTCTTCTTCAGTTTCGACTTATCGAATTTCTCGATCTCAGCCATATC | 51202 | 0.20343252758260516 | No Hit |
| ACGCTCGTAGCCCTCGCGCTTCTCCTCGGCCAGTTCGCGGAAGAAGTGGC | 49216 | 0.19554187878413923 | No Hit |
| ATCAAATCCTGCAGACAAGGGGAGCCCTCAGTCTGCAGGGCTCCATAATG | 48524 | 0.19279246842737263 | No Hit |
| ACATAGCAATTCAGGAAATTTGACTTTCCATTCTCTGCTGGATGACGTGA | 47157 | 0.1873611910318525 | No Hit |
| GCTGCTACAGGAGAATAGCAGACAGGTATAGTTAAGAAACAACTTTATTT | 46924 | 0.18643545026143835 | No Hit |
| ACGCTCGTAGCCCTCGCGCTTCTCCTCGGCCAATTCGCGGAAGAAGTGGC | 45676 | 0.18147697609200958 | No Hit |
| ACTTGATCCCAACTCATCTCTCATTTATTTCGGCTTCTTTTATTCCAGGA | 41836 | 0.16622013249376724 | No Hit |
| AGCGGTTCCGGTGTCCGGAGGCGCTGTTCCAGCCTTCCTTCCTGGGTATG | 39934 | 0.15866322714901282 | No Hit |
| CTAAGTGCTGTGTTGTCGTTCCCCCTGCTTAAAATAAAGTTGTTTCTTAA | 39647 | 0.15752293701549838 | No Hit |
| TACTCGTGCGCCTCGCTTCGCTTTTCCTCCGCAACCATGTCTGACAAACC | 38922 | 0.15464241315905938 | No Hit |
| TCGGGCGCCCCAGACACCAGGCGTCATGGTGGGCATGGGCCAGAAGGACT | 38727 | 0.15386765157008614 | No Hit |
| TGGTTGCACGAAACACACTGGGGAATGGAGCAAAACAGTCTTTGAATATC | 36593 | 0.14538897859127126 | No Hit |
| TGTCCATGTCTTACTACTTTGACCGCGATGATGTGGCTTTGAAGAACTTT | 36393 | 0.14459435132052947 | No Hit |
| ACCCGCAGCTTCTGCTTCTCAGTCAGAAGGTTGTTGTCCTCATCCCTCTC | 36352 | 0.14443145273002742 | No Hit |
| TCAGCACGGGCTCGTACTCTGCCACAAACTGATCACACTGCTTCTGGTAA | 35712 | 0.14188864546365368 | No Hit |
| TGTTCCCTCTCCTCATGAGATTGGTGAAGAAAGTATTTGGCAAAGTTCTT | 35437 | 0.1407960329663837 | No Hit |
| AAAGAAGAACATGTGATCATCCAGGCCGAGTTCTATCTGAATCCTGACCA | 35315 | 0.14031131033123123 | No Hit |
| TCACACTTCATGATGGAGTTGAAGGTGGTCTCGTGGATGCCGCAAGATTC | 34306 | 0.1363024157503389 | No Hit |
| AATACCAAGACCTGCTCAATGTTAAGATGGCCCTTGACATTGAGATTGCC | 33001 | 0.13111747280874875 | No Hit |
| AGCACCAAGCAGGAGATCCTGGCTGCTCTTGAGAAAGGCTGCAGCTTCCT | 31666 | 0.1258133357765473 | No Hit |
| TACATCAAAGATTACATGAAATCAATCAAAGGGAAACTTGAAGAACAGAG | 30897 | 0.12275799392054514 | No Hit |
| ATGTTCTGGGAGGCTCGGTGGACATCAGGCGCAGGAAGGTCAGCTGGATG | 30797 | 0.12236068028517424 | No Hit |
| TGAGTTGGCCAGAACTCTGAAAAGATTGGGAATGGATGGCTACAGGGGAA | 30598 | 0.12157002615078617 | No Hit |
| GAAAAGTTTGGAAGAGGCAGAGAAATCCTGCTCTCCTCGCCTTCCAGCAG | 30090 | 0.11955167288310202 | No Hit |
| TTGGCTTTCTCTTTCCTCTTCTCCTCCAGGGTGGCTGTCACTGCCTGGTA | 28671 | 0.11391379239718905 | No Hit |
| ACAGTGTATCTAGCTTTGGAAAACACAGGGGTCTGCCCTGTGAGCTGCTC | 26187 | 0.10404452169457602 | No Hit |
| ACCTACGGCGAGGGCGAGAGCGGCCCCATGGGCAACATCATGATCGATCC | 26103 | 0.10371077824086448 | No Hit |
| AAACCAGACATGGGGGAAATCGCCAGCTTCGATAAGGCCAAGCTGAAGAA | 26097 | 0.10368693942274224 | No Hit |
| TTCCCCCTGCGCATGCGGGACTGGCTCAAGAACGTCCTGGTCACCCTGTA | 26049 | 0.10349622887776419 | No Hit |
| TTCTTGTAGATGGCTGCTCTAAAAAGACAAATGAATGGGGAAAGACAATC | 25829 | 0.10262213887994823 | No Hit |

### Adapter Content

Produced by FastQC (version 0.11.7)
