## Supplementary material for "Curated and harmonised transcriptomics datasets of interstitial lung diseases": SRR8397218_fastqc.html

SRR8397218.fastq.gz FastQC Report 

FastQC Report

Tue 30 Jan 2024  
SRR8397218.fastq.gz

### Summary

- Basic Statistics
- Per base sequence quality
- Per sequence quality scores
- Per base sequence content
- Per sequence GC content
- Per base N content
- Sequence Length Distribution
- Sequence Duplication Levels
- Overrepresented sequences
- Adapter Content

### Basic Statistics

| Measure | Value |
| --- | --- |
| Filename | SRR8397218.fastq.gz |
| File type | Conventional base calls |
| Encoding | Sanger / Illumina 1.9 |
| Total Sequences | 21009611 |
| Sequences flagged as poor quality | 0 |
| Sequence length | 8-369 |
| %GC | 53 |

### Per base sequence quality

### Per sequence quality scores

### Per base sequence content

### Per sequence GC content

### Per base N content

### Sequence Length Distribution

### Sequence Duplication Levels

### Overrepresented sequences

| Sequence | Count | Percentage | Possible Source |
| --- | --- | --- | --- |
| AAGCTCAGGGAGAGCCCTGTTAGGGCCGCCTCTGGCCCTAGTCTCAGACC | 207492 | 0.9876051489006628 | No Hit |
| GGGCCACGAGCTGAGTGCGTCCTGTCACTCCACTCCCATGTCCCTTGGGA | 201943 | 0.9611934271415115 | No Hit |
| CCACCTTCTCCATCGGCTCCACTGGCCTCGTGGTGTATGACTACCAGCAG | 194740 | 0.9269091179270288 | No Hit |
| ATGATGTAGCAGCAGGTGCCAGGGGCTGGCTTGTAGGCGATCAGCAGCTG | 162475 | 0.773336545831334 | No Hit |
| GGGAATGTCTACGTGCGTATGCACGTGGCACTCTCTGCCCGAGGTCCGGG | 132062 | 0.6285789870169419 | No Hit |
| TCTCTGGGCTTCTATTTCGACCGCGATGATGTGGCTCTGGAAGGCGTGAG | 105159 | 0.5005280678447592 | No Hit |
| AAAAGTGATGGACTGCCCATTGCTGGAGAAGACCTTCTCTCCTACTGTCA | 95575 | 0.45491085008665794 | No Hit |
| ATCAAATCCTGCAGACAAGGGGAGCCCTCAGTCTGCAGGGCTCCATAATG | 85496 | 0.4069375677636297 | No Hit |
| TGGTTCCTGGCAAGCCCATGTGTGTTGAGAGCTTCTCAGACTATCCACCT | 72735 | 0.3461986992524516 | No Hit |
| AGCGGTTCCGCTGCCCTGAGGCACTCTTCCAGCCTTCCTTCCTGGGCATG | 69848 | 0.33245736915357454 | No Hit |
| ACACTGAGGACTCTGTTCCTCCCCTTTCCGCCTAGGGGAAAGTCCCCGGA | 69086 | 0.32883045764150515 | No Hit |
| TCACACTTCATGATGGAGTTGAAGGTAGTTTCGTGGATGCCACAGGACTC | 64432 | 0.3066786910047978 | No Hit |
| ACACCCACCGCAACTGTCTGTCTCATATCACGAACAGCAAAGCGACCCAA | 54858 | 0.26110907051063437 | No Hit |
| TCTTTCTGGCCTGGAGGCTATCCAGCGTACTCCAAAGATTCAGGTTTACT | 50133 | 0.23861936329996783 | No Hit |
| TCTGTCTTCTTCAGTTTCGACTTATCGAATTTCTCGATCTCAGCCATATC | 49897 | 0.23749606787103295 | No Hit |
| GGCTGCCTGGAGCCCCTGGTGTCCCTGGAGAGCGTGGAGAGAAGGGGGAG | 41579 | 0.19790466372747217 | No Hit |
| ACATAGCAATTCAGGAAATTTGACTTTCCATTCTCTGCTGGATGACGTGA | 40592 | 0.1932068137767996 | No Hit |
| TACTCGTGCGCCTCGCTTCGCTTTTCCTCCGCAACCATGTCTGACAAACC | 38830 | 0.18482017587093832 | No Hit |
| ACGCTCGTAGCCCTCGCGCTTCTCCTCGGCCAGTTCGCGGAAGAAGTGGC | 38726 | 0.18432516432598395 | No Hit |
| ACGCTCGTAGCCCTCGCGCTTCTCCTCGGCCAATTCGCGGAAGAAGTGGC | 37684 | 0.17936552942365283 | No Hit |
| ACTTGATCCCAACTCATCTCTCATTTATTTCGGCTTCTTTTATTCCAGGA | 36954 | 0.17589092915618476 | No Hit |
| GGGTACTTCAGGGTCAGGATGCCACGCTTGCTCTGGGCCTCGTCGCCCAC | 34736 | 0.1653338560147544 | No Hit |
| CTAAGTGCTGTGTTGTCGTTCCCCCTGCTTAAAATAAAGTTGTTTCTTAA | 34613 | 0.164748409668318 | No Hit |
| GCTGCTACAGGAGAATAGCAGACAGGTATAGTTAAGAAACAACTTTATTT | 32531 | 0.15483865931644333 | No Hit |
| ACACTGAGGACTCTGTTCCTCCCCTTTCCGCCTAGGGAAAGTCCCCGGAC | 31099 | 0.14802273112053335 | No Hit |
| TCGGGCGCCCCAGACACCAGGCGTCATGGTGGGCATGGGCCAGAAGGACT | 30020 | 0.14288698634163194 | No Hit |
| ACCAATGTCATAGGGTGCAATATCTACAATAGGTAGTCTCACAGCCTTGC | 29786 | 0.14177321036548463 | No Hit |
| AGCGGTTCCGGTGTCCGGAGGCGCTGTTCCAGCCTTCCTTCCTGGGTATG | 28164 | 0.13405293415475422 | No Hit |
| TGTCCATGTCTTACTACTTTGACCGCGATGATGTGGCTTTGAAGAACTTT | 27015 | 0.12858400852828736 | No Hit |
| TGTTCCCTCTCCTCATGAGATTGGTGAAGAAAGTATTTGGCAAAGTTCTT | 26826 | 0.1276844202398607 | No Hit |
| AGAGCATGACCGATGGATTCCAGTTCGAGTATGGCGGCCAGGGCTCCGAC | 25296 | 0.12040203885735913 | No Hit |
| TACATCAAAGATTACATGAAATCAATCAAAGGGAAACTTGAAGAACAGAG | 24870 | 0.11837439541360381 | No Hit |
| GGGCCACGAGCTGAGTGCGTCCTGTCACTCCACTCCCCATGTCCCTTGGG | 24839 | 0.11822684389539626 | No Hit |
| TGGCTCTCCCAGAGGCAGAAGGACAATGGCTGTTTCAGGAGCTCTGGGTC | 24542 | 0.11681320515644006 | No Hit |
| TCACACTTCATGATGGAGTTGAAGGTGGTCTCGTGGATGCCGCAAGATTC | 24012 | 0.11429055016773038 | No Hit |
| GAGCTCCTCATCTAGATGAGCTGGAAGCCCTGGAGGGCCTCTCTCGCCAG | 23175 | 0.11030665917612659 | No Hit |
| AATAAAAGCGAAAAGAAATGAAAATGTTACACTACATTAATCCTGGAATA | 22910 | 0.10904533168177173 | No Hit |
| ACCCGCAGCTTCTGCTTCTCAGTCAGAAGGTTGTTGTCCTCATCCCTCTC | 22701 | 0.10805054886546923 | No Hit |
| GAGAGGGTCACTTCATCTTCTACTCCTCCCTTTATGGCATTGTTGAGCAG | 21941 | 0.10443315680618741 | No Hit |
| AAAGAAGAACATGTGATCATCCAGGCCGAGTTCTATCTGAATCCTGACCA | 21895 | 0.10421420939207299 | No Hit |

### Adapter Content

Produced by FastQC (version 0.11.7)
