## Supplementary material for "Curated and harmonised transcriptomics datasets of interstitial lung diseases": SRR8397223_fastqc.html

SRR8397223.fastq.gz FastQC Report 

FastQC Report

Tue 30 Jan 2024  
SRR8397223.fastq.gz

### Summary

- Basic Statistics
- Per base sequence quality
- Per sequence quality scores
- Per base sequence content
- Per sequence GC content
- Per base N content
- Sequence Length Distribution
- Sequence Duplication Levels
- Overrepresented sequences
- Adapter Content

### Basic Statistics

| Measure | Value |
| --- | --- |
| Filename | SRR8397223.fastq.gz |
| File type | Conventional base calls |
| Encoding | Sanger / Illumina 1.9 |
| Total Sequences | 24069955 |
| Sequences flagged as poor quality | 0 |
| Sequence length | 8-368 |
| %GC | 53 |

### Per base sequence quality

### Per sequence quality scores

### Per base sequence content

### Per sequence GC content

### Per base N content

### Sequence Length Distribution

### Sequence Duplication Levels

### Overrepresented sequences

| Sequence | Count | Percentage | Possible Source |
| --- | --- | --- | --- |
| AAGCTCAGGGAGAGCCCTGTTAGGGCCGCCTCTGGCCCTAGTCTCAGACC | 217194 | 0.9023448527427659 | No Hit |
| GGGCCACGAGCTGAGTGCGTCCTGTCACTCCACTCCCATGTCCCTTGGGA | 203158 | 0.8440314907111377 | No Hit |
| GGGAATGTCTACGTGCGTATGCACGTGGCACTCTCTGCCCGAGGTCCGGG | 178131 | 0.7400553927084617 | No Hit |
| CCACCTTCTCCATCGGCTCCACTGGCCTCGTGGTGTATGACTACCAGCAG | 163521 | 0.6793573149596666 | No Hit |
| ATGATGTAGCAGCAGGTGCCAGGGGCTGGCTTGTAGGCGATCAGCAGCTG | 130152 | 0.5407239024750982 | No Hit |
| ACACTGAGGACTCTGTTCCTCCCCTTTCCGCCTAGGGGAAAGTCCCCGGA | 118382 | 0.49182476660218105 | No Hit |
| TCTCTGGGCTTCTATTTCGACCGCGATGATGTGGCTCTGGAAGGCGTGAG | 102500 | 0.4258420923512321 | No Hit |
| TGGTTCCTGGCAAGCCCATGTGTGTTGAGAGCTTCTCAGACTATCCACCT | 85179 | 0.35388101057937166 | No Hit |
| AGCGGTTCCGCTGCCCTGAGGCACTCTTCCAGCCTTCCTTCCTGGGCATG | 79317 | 0.32952699745387976 | No Hit |
| ACACCCACCGCAACTGTCTGTCTCATATCACGAACAGCAAAGCGACCCAA | 78374 | 0.3256092502042484 | No Hit |
| AAAAGTGATGGACTGCCCATTGCTGGAGAAGACCTTCTCTCCTACTGTCA | 77651 | 0.32260550549429773 | No Hit |
| ACACTGAGGACTCTGTTCCTCCCCTTTCCGCCTAGGGAAAGTCCCCGGAC | 77576 | 0.32229391371940663 | No Hit |
| AAGACCAACGTCAAGGCCGCCTGGGGTAAGGTCGGCGCGCACGCTGGCGA | 73908 | 0.3070549986487303 | No Hit |
| ATCAAATCCTGCAGACAAGGGGAGCCCTCAGTCTGCAGGGCTCCATAATG | 72352 | 0.3005905079589887 | No Hit |
| TCACACTTCATGATGGAGTTGAAGGTAGTTTCGTGGATGCCACAGGACTC | 67060 | 0.2786045923226695 | No Hit |
| GGCTGCCTGGAGCCCCTGGTGTCCCTGGAGAGCGTGGAGAGAAGGGGGAG | 58610 | 0.24349858568493377 | No Hit |
| TCTGTCTTCTTCAGTTTCGACTTATCGAATTTCTCGATCTCAGCCATATC | 51220 | 0.2127964094656596 | No Hit |
| GGGTACTTCAGGGTCAGGATGCCACGCTTGCTCTGGGCCTCGTCGCCCAC | 50293 | 0.20894513512800503 | No Hit |
| TCTTTCTGGCCTGGAGGCTATCCAGCGTACTCCAAAGATTCAGGTTTACT | 49496 | 0.20563395320016176 | No Hit |
| TTGGTGGTGGGGAAGGACAGGAACATCCTCTCCAGGGCCTCCGCACCATA | 48602 | 0.2019197792434593 | No Hit |
| GAGCTCCTCATCTAGATGAGCTGGAAGCCCTGGAGGGCCTCTCTCGCCAG | 44796 | 0.18610753530698335 | No Hit |
| ACGCTCGTAGCCCTCGCGCTTCTCCTCGGCCAATTCGCGGAAGAAGTGGC | 41902 | 0.17408424735318367 | No Hit |
| ACGCTCGTAGCCCTCGCGCTTCTCCTCGGCCAGTTCGCGGAAGAAGTGGC | 41794 | 0.17363555519734042 | No Hit |
| ACTTGATCCCAACTCATCTCTCATTTATTTCGGCTTCTTTTATTCCAGGA | 40304 | 0.167445265269503 | No Hit |
| TCGGGCGCCCCAGACACCAGGCGTCATGGTGGGCATGGGCCAGAAGGACT | 38273 | 0.1590073600054508 | No Hit |
| AGCGGTTCCGGTGTCCGGAGGCGCTGTTCCAGCCTTCCTTCCTGGGTATG | 37698 | 0.15661848973128534 | No Hit |
| CCCATAACAGCATCAGGAGTGGACAGATCCCCAAAGGACTCAAAGAACCT | 37401 | 0.1553845863027164 | No Hit |
| TACTCGTGCGCCTCGCTTCGCTTTTCCTCCGCAACCATGTCTGACAAACC | 36974 | 0.15361059046433614 | No Hit |
| GCTGCTACAGGAGAATAGCAGACAGGTATAGTTAAGAAACAACTTTATTT | 34890 | 0.14495249367936083 | No Hit |
| ACATAGCAATTCAGGAAATTTGACTTTCCATTCTCTGCTGGATGACGTGA | 34756 | 0.14439578304155534 | No Hit |
| TGGCTCTCCCAGAGGCAGAAGGACAATGGCTGTTTCAGGAGCTCTGGGTC | 33314 | 0.13840491184964823 | No Hit |
| GAGAGGGTCACTTCATCTTCTACTCCTCCCTTTATGGCATTGTTGAGCAG | 33299 | 0.13834259349467 | No Hit |
| TCACACTTCATGATGGAGTTGAAGGTGGTCTCGTGGATGCCGCAAGATTC | 31902 | 0.13253867736769762 | No Hit |
| ACCTACGGCGAGGGCGAGAGCGGCCCCATGGGCAACATCATGATCGATCC | 30749 | 0.12774847314837107 | No Hit |
| AAGGCCCACCCGTGGAGGCCAGACCCAAAGCCCACGGTACCGAGGACAGG | 29998 | 0.12462840084246107 | No Hit |
| AATAAAAGCGAAAAGAAATGAAAATGTTACACTACATTAATCCTGGAATA | 29827 | 0.12391797159570926 | No Hit |
| CAGCTGGAGAAGGACTTCAGCAGCATGAAGAAGTACTGCCAAGTCATCCG | 29689 | 0.12334464272990954 | No Hit |
| TACATCAAAGATTACATGAAATCAATCAAAGGGAAACTTGAAGAACAGAG | 28780 | 0.11956815041822887 | No Hit |
| TTGGCTTTCTCTTTCCTCTTCTCCTCCAGGGTGGCTGTCACTGCCTGGTA | 27888 | 0.11586228557552351 | No Hit |
| ACACTGAGGACTCTGTTCCTCCCCTTCCGCCTAGGGAAAGTCCCCGGACC | 27239 | 0.11316597808346547 | No Hit |
| GGCCTTCTTCTGGCGCAGAGGAAGCAGGCGCATCTGGGTGTGGGCAATGA | 26962 | 0.11201516579486749 | No Hit |
| AAAAGTGATGGACTGCCCATTGCTAGAGAAGACCTTCTCTCCTACTGTCA | 26481 | 0.11001682387856562 | No Hit |
| AACATCTGTGTTGGGGAGAGTGGAGACAGACTGACGCGAGCAGCCAAGGT | 26240 | 0.10901557564191541 | No Hit |
| AGAGCATGACCGATGGATTCCAGTTCGAGTATGGCGGCCAGGGCTCCGAC | 25414 | 0.10558391156111427 | No Hit |
| ACAGTGTATCTAGCTTTGGAAAACACAGGGGTCTGCCCTGTGAGCTGCTC | 24407 | 0.10140027266357582 | No Hit |

### Adapter Content

Produced by FastQC (version 0.11.7)
