## Supplementary material for "Curated and harmonised transcriptomics datasets of interstitial lung diseases": SRR8397236_fastqc.html

SRR8397236.fastq.gz FastQC Report 

FastQC Report

Tue 30 Jan 2024  
SRR8397236.fastq.gz

### Summary

- Basic Statistics
- Per base sequence quality
- Per sequence quality scores
- Per base sequence content
- Per sequence GC content
- Per base N content
- Sequence Length Distribution
- Sequence Duplication Levels
- Overrepresented sequences
- Adapter Content

### Basic Statistics

| Measure | Value |
| --- | --- |
| Filename | SRR8397236.fastq.gz |
| File type | Conventional base calls |
| Encoding | Sanger / Illumina 1.9 |
| Total Sequences | 25036247 |
| Sequences flagged as poor quality | 0 |
| Sequence length | 8-372 |
| %GC | 53 |

### Per base sequence quality

### Per sequence quality scores

### Per base sequence content

### Per sequence GC content

### Per base N content

### Sequence Length Distribution

### Sequence Duplication Levels

### Overrepresented sequences

| Sequence | Count | Percentage | Possible Source |
| --- | --- | --- | --- |
| GGGCCACGAGCTGAGTGCGTCCTGTCACTCCACTCCCATGTCCCTTGGGA | 326896 | 1.305690904870846 | No Hit |
| AAGCTCAGGGAGAGCCCTGTTAGGGCCGCCTCTGGCCCTAGTCTCAGACC | 315845 | 1.2615509025773712 | No Hit |
| TCTCTGGGCTTCTATTTCGACCGCGATGATGTGGCTCTGGAAGGCGTGAG | 178034 | 0.7111049831070927 | No Hit |
| ACACTGAGGACTCTGTTCCTCCCCTTTCCGCCTAGGGGAAAGTCCCCGGA | 154586 | 0.6174487733724627 | No Hit |
| GGGAATGTCTACGTGCGTATGCACGTGGCACTCTCTGCCCGAGGTCCGGG | 139750 | 0.5581906904816845 | No Hit |
| TGGTTCCTGGCAAGCCCATGTGTGTTGAGAGCTTCTCAGACTATCCACCT | 121415 | 0.4849568707322627 | No Hit |
| ACGCTCGTAGCCCTCGCGCTTCTCCTCGGCCAGTTCGCGGAAGAAGTGGC | 116391 | 0.46488996533705707 | No Hit |
| AGCGGTTCCGCTGCCCTGAGGCACTCTTCCAGCCTTCCTTCCTGGGCATG | 94789 | 0.3786070651883247 | No Hit |
| TCACACTTCATGATGGAGTTGAAGGTAGTTTCGTGGATGCCACAGGACTC | 88480 | 0.35340760138690114 | No Hit |
| ACACCCACCGCAACTGTCTGTCTCATATCACGAACAGCAAAGCGACCCAA | 84560 | 0.337750302591279 | No Hit |
| GGGTACTTCAGGGTCAGGATGCCACGCTTGCTCTGGGCCTCGTCGCCCAC | 84124 | 0.33600882752115363 | No Hit |
| CCACCTTCTCCATCGGCTCCACTGGCCTCGTGGTGTATGACTACCAGCAG | 79617 | 0.31800692811506454 | No Hit |
| AAAAGTGATGGACTGCCCATTGCTGGAGAAGACCTTCTCTCCTACTGTCA | 74386 | 0.2971132214824371 | No Hit |
| TCTTTCTGGCCTGGAGGCTATCCAGCGTACTCCAAAGATTCAGGTTTACT | 69390 | 0.2771581539357716 | No Hit |
| TCGGGCGCCCCAGACACCAGGCGTCATGGTGGGCATGGGCCAGAAGGACT | 59047 | 0.23584605152681232 | No Hit |
| ATCAAGACATGAGGGAGGCAGGGGCTCAGCTGAAGAAGCTGGTGGACACC | 57595 | 0.2300464602382298 | No Hit |
| AGCTTAATGATGCTTTCTCTGGGCTTTTGGGGGAGGGTGTCCACCAGCTT | 57319 | 0.22894405858833394 | No Hit |
| TGTTCCCTCTCCTCATGAGATTGGTGAAGAAAGTATTTGGCAAAGTTCTT | 56584 | 0.22600831506415478 | No Hit |
| AGCGGTTCCGGTGTCCGGAGGCGCTGTTCCAGCCTTCCTTCCTGGGTATG | 55280 | 0.22079986668928453 | No Hit |
| ATGATGTAGCAGCAGGTGCCAGGGGCTGGCTTGTAGGCGATCAGCAGCTG | 52313 | 0.20894904895290417 | No Hit |
| ATCAAATCCTGCAGACAAGGGGAGCCCTCAGTCTGCAGGGCTCCATAATG | 51782 | 0.20682812403951759 | No Hit |
| GCTGCTACAGGAGAATAGCAGACAGGTATAGTTAAGAAACAACTTTATTT | 51581 | 0.2060252880553543 | No Hit |
| TGTCCATGTCTTACTACTTTGACCGCGATGATGTGGCTTTGAAGAACTTT | 47647 | 0.19031207033546202 | No Hit |
| TCACACTTCATGATGGAGTTGAAGGTGGTCTCGTGGATGCCGCAAGATTC | 45803 | 0.1829467491673173 | No Hit |
| ACATAGCAATTCAGGAAATTTGACTTTCCATTCTCTGCTGGATGACGTGA | 45342 | 0.1811054188752811 | No Hit |
| ACACTGAGGACTCTGTTCCTCCCCTTTCCGCCTAGGGAAAGTCCCCGGAC | 42081 | 0.16808030372922905 | No Hit |
| AAGGCCCACCCGTGGAGGCCAGACCCAAAGCCCACGGTACCGAGGACAGG | 37654 | 0.1503979410332547 | No Hit |
| TTGGCTTTCTCTTTCCTCTTCTCCTCCAGGGTGGCTGTCACTGCCTGGTA | 37584 | 0.1501183464119043 | No Hit |
| ACCTACGGCGAGGGCGAGAGCGGCCCCATGGGCAACATCATGATCGATCC | 37524 | 0.14987869387931826 | No Hit |
| TCAGCACGGGCTCGTACTCTGCCACAAACTGATCACACTGCTTCTGGTAA | 34720 | 0.13867893218979666 | No Hit |
| CTAAGTGCTGTGTTGTCGTTCCCCCTGCTTAAAATAAAGTTGTTTCTTAA | 34587 | 0.1381477024092309 | No Hit |
| TCTGTCTTCTTCAGTTTCGACTTATCGAATTTCTCGATCTCAGCCATATC | 34529 | 0.13791603829439772 | No Hit |
| AACATCTGTGTTGGGGAGAGTGGAGACAGACTGACGCGAGCAGCCAAGGT | 34232 | 0.13672975825809675 | No Hit |
| ACAGTGTATCTAGCTTTGGAAAACACAGGGGTCTGCCCTGTGAGCTGCTC | 34030 | 0.135922928065057 | No Hit |
| TACATCAAAGATTACATGAAATCAATCAAAGGGAAACTTGAAGAACAGAG | 32854 | 0.1312257384263704 | No Hit |
| CGGTTGTACTTGCGGATGTAGTGCAGATAGTCTCGGCGGATGACAATGGT | 32261 | 0.12885717256264487 | No Hit |
| GGCTGCCTGGAGCCCCTGGTGTCCCTGGAGAGCGTGGAGAGAAGGGGGAG | 31186 | 0.12456339802047807 | No Hit |
| GGCCTTCTTCTGGCGCAGAGGAAGCAGGCGCATCTGGGTGTGGGCAATGA | 30275 | 0.12092467373404647 | No Hit |
| GCCAAGCACAAAGAGCTTGCTCCCTACGATGAGAACTGGTTCTACACGCG | 30170 | 0.1205052818020209 | No Hit |
| CAGCTGGAGAAGGACTTCAGCAGCATGAAGAAGTACTGCCAAGTCATCCG | 29987 | 0.11977434157763342 | No Hit |
| ATTCGAGGGCGGATCCTCTCTGGCGTGGTGACCAAGATGAAGATGCAGAG | 29210 | 0.116670841280644 | No Hit |
| CCCAGGGCAGAGGCTGACGACGCTGACTGGCAGGGCACCGACACAGTGTC | 28931 | 0.11555645700411887 | No Hit |
| AAAAGGTCAAGGCCTTCTTGGCTGATCCATCTGCCTTTGTGGCTGCTGCC | 28796 | 0.11501723880580025 | No Hit |
| TACTGCTTTACAAGAGGTTCTGAAGACTGCCCTCATCCACGATGGCCTAG | 28794 | 0.11500925038804737 | No Hit |
| TCTCTGGCTTCTATTTCGACCGCGATGATGTGGCTCTGGAAGGCGTGAGC | 27930 | 0.11155825391880819 | No Hit |
| GAGCTCCTCATCTAGATGAGCTGGAAGCCCTGGAGGGCCTCTCTCGCCAG | 27144 | 0.1084188057419309 | No Hit |
| TACTCGTGCGCCTCGCTTCGCTTTTCCTCCGCAACCATGTCTGACAAACC | 25901 | 0.10345400410852314 | No Hit |
| CCACAGCTCTGCCAGTACCCCAAGACTCAGCACTAGTCTGATGACCTGCT | 25357 | 0.10128115447974291 | No Hit |

### Adapter Content

Produced by FastQC (version 0.11.7)
