## Supplementary material for "Curated and harmonised transcriptomics datasets of interstitial lung diseases": TRIMMED_SRR8397173_fastqc.html

TRIMMED\_SRR8397173.fastq.gz FastQC Report 

FastQC Report

Sat 3 Dec 2022  
TRIMMED\_SRR8397173.fastq.gz

### Summary

- Basic Statistics
- Per base sequence quality
- Per sequence quality scores
- Per base sequence content
- Per sequence GC content
- Per base N content
- Sequence Length Distribution
- Sequence Duplication Levels
- Overrepresented sequences
- Adapter Content

### Basic Statistics

| Measure | Value |
| --- | --- |
| Filename | TRIMMED\_SRR8397173.fastq.gz |
| File type | Conventional base calls |
| Encoding | Sanger / Illumina 1.9 |
| Total Sequences | 25611252 |
| Sequences flagged as poor quality | 0 |
| Sequence length | 35-368 |
| %GC | 53 |

### Per base sequence quality

### Per sequence quality scores

### Per base sequence content

### Per sequence GC content

### Per base N content

### Sequence Length Distribution

### Sequence Duplication Levels

### Overrepresented sequences

| Sequence | Count | Percentage | Possible Source |
| --- | --- | --- | --- |
| GGGCCACGAGCTGAGTGCGTCCTGTCACTCCACTCCCATGTCCCTTGGGA | 352245 | 1.3753525208373256 | No Hit |
| AAGCTCAGGGAGAGCCCTGTTAGGGCCGCCTCTGGCCCTAGTCTCAGACC | 338177 | 1.3204235388414436 | No Hit |
| ACACTGAGGACTCTGTTCCTCCCCTTTCCGCCTAGGGGAAAGTCCCCGGA | 277876 | 1.0849762440352388 | No Hit |
| GGGAATGTCTACGTGCGTATGCACGTGGCACTCTCTGCCCGAGGTCCGGG | 247987 | 0.9682736322300838 | No Hit |
| AGCGGTTCCGCTGCCCTGAGGCACTCTTCCAGCCTTCCTTCCTGGGCATG | 140819 | 0.5498325501619367 | No Hit |
| CCACCTTCTCCATCGGCTCCACTGGCCTCGTGGTGTATGACTACCAGCAG | 139310 | 0.5439406086043743 | No Hit |
| TCACACTTCATGATGGAGTTGAAGGTAGTTTCGTGGATGCCACAGGACTC | 132724 | 0.5182253487646757 | No Hit |
| TCTCTGGGCTTCTATTTCGACCGCGATGATGTGGCTCTGGAAGGCGTGAG | 132191 | 0.5161442322304275 | No Hit |
| AAAAGTGATGGACTGCCCATTGCTGGAGAAGACCTTCTCTCCTACTGTCA | 113284 | 0.44232121100522537 | No Hit |
| ATGATGTAGCAGCAGGTGCCAGGGGCTGGCTTGTAGGCGATCAGCAGCTG | 108881 | 0.4251295485281235 | No Hit |
| TCTTTCTGGCCTGGAGGCTATCCAGCGTACTCCAAAGATTCAGGTTTACT | 105959 | 0.4137205006611937 | No Hit |
| GGGTACTTCAGGGTCAGGATGCCACGCTTGCTCTGGGCCTCGTCGCCCAC | 98559 | 0.38482695027950997 | No Hit |
| ATCAAATCCTGCAGACAAGGGGAGCCCTCAGTCTGCAGGGCTCCATAATG | 93428 | 0.3647927871702641 | No Hit |
| TGGTTCCTGGCAAGCCCATGTGTGTTGAGAGCTTCTCAGACTATCCACCT | 84498 | 0.3299253000204754 | No Hit |
| TGAGGCGGAGCACCAGAGAGCCTACCTGGAAGACACATGCGTGGAGTGGC | 70936 | 0.2769720121452868 | No Hit |
| ACACCCACCGCAACTGTCTGTCTCATATCACGAACAGCAAAGCGACCCAA | 67697 | 0.2643252270525471 | No Hit |
| ACATAGCAATTCAGGAAATTTGACTTTCCATTCTCTGCTGGATGACGTGA | 63740 | 0.24887498666601696 | No Hit |
| AGCGGTTCCGGTGTCCGGAGGCGCTGTTCCAGCCTTCCTTCCTGGGTATG | 61655 | 0.24073403361928578 | No Hit |
| TCACACTTCATGATGGAGTTGAAGGTGGTCTCGTGGATGCCGCAAGATTC | 53632 | 0.20940795865817102 | No Hit |
| TCTGTCTTCTTCAGTTTCGACTTATCGAATTTCTCGATCTCAGCCATATC | 49794 | 0.1944223577980491 | No Hit |
| TCGGGCGCCCCAGACACCAGGCGTCATGGTGGGCATGGGCCAGAAGGACT | 46465 | 0.18142416466012673 | No Hit |
| TGTTCCCTCTCCTCATGAGATTGGTGAAGAAAGTATTTGGCAAAGTTCTT | 45361 | 0.17711355930588635 | No Hit |
| GGCTGCCTGGAGCCCCTGGTGTCCCTGGAGAGCGTGGAGAGAAGGGGGAG | 41523 | 0.1621279584457644 | No Hit |
| TGTCCATGTCTTACTACTTTGACCGCGATGATGTGGCTTTGAAGAACTTT | 39403 | 0.15385034671479553 | No Hit |
| ACGCTCGTAGCCCTCGCGCTTCTCCTCGGCCAATTCGCGGAAGAAGTGGC | 38810 | 0.15153495815042545 | No Hit |
| GGGGCTCCAGGTGAAGCAGCGTCTCCTTCCCCTTCTCCAGGTATTTGTGG | 38797 | 0.15148419921056572 | No Hit |
| GCTGCTACAGGAGAATAGCAGACAGGTATAGTTAAGAAACAACTTTATTT | 36915 | 0.14413586653241317 | No Hit |
| ACGCTCGTAGCCCTCGCGCTTCTCCTCGGCCAGTTCGCGGAAGAAGTGGC | 36518 | 0.1425857665997742 | No Hit |
| CGTGCTGGCTACTACCGACCGGAGGCGCCGTGACCTTGGTGGCTCAGCCC | 35387 | 0.138169738831979 | No Hit |
| TACTCGTGCGCCTCGCTTCGCTTTTCCTCCGCAACCATGTCTGACAAACC | 35382 | 0.13815021616280218 | No Hit |
| GGGAATGTCTACGTGCGTATGCACGTGGCACTCTCTGCCCGGAGGTCCGG | 35125 | 0.13714675096711398 | No Hit |
| GAGCTCCTCATCTAGATGAGCTGGAAGCCCTGGAGGGCCTCTCTCGCCAG | 34856 | 0.13609643136540142 | No Hit |
| CTAAGTGCTGTGTTGTCGTTCCCCCTGCTTAAAATAAAGTTGTTTCTTAA | 34202 | 0.13354286623707423 | No Hit |
| ACACTGAGGACTCTGTTCCTCCCCTTTCCGCCTAGGGAAAGTCCCCGGAC | 32720 | 0.12775634709306674 | No Hit |
| CCAGTGTAGTCAATAGCCAGGAGGTGTCCAAGGGCTACAGAGAGGCCGAT | 32602 | 0.12729561210049395 | No Hit |
| TCGGCGCCCCAGACACCAGGCGTCATGGTGGGCATGGGCCAGAAGGACTC | 32514 | 0.12695201312298204 | No Hit |
| TCAGCACGGGCTCGTACTCTGCCACAAACTGATCACACTGCTTCTGGTAA | 31045 | 0.12121625291883426 | No Hit |
| CCGGACTCCACCTGGCTGTTGTCAATCTTCACCTCATAGGTGTTGTCTGG | 30070 | 0.11740933242935567 | No Hit |
| CGGTTGTACTTGCGGATGTAGTGCAGATAGTCTCGGCGGATGACAATGGT | 29578 | 0.11548830178235722 | No Hit |
| GACATCCGTTGCAAGGATGATGAGTTTACACACCTGTACACACTGATTGT | 28533 | 0.11140806392440322 | No Hit |
| GGGCTCCAGGTGAAGCAGCGTCTCCTTCCCCTTCTCCAGGTATTTGTGGA | 27328 | 0.1067031006527912 | No Hit |
| ACCTACGGCGAGGGCGAGAGCGGCCCCATGGGCAACATCATGATCGATCC | 26188 | 0.10225193208047774 | No Hit |
| TTCCTTCTTCTCCTTTAAGTCCTTGGTGGTGATTTCGGAGCTGGTGTCTA | 26113 | 0.10195909204282555 | No Hit |
| AGCACCAAGCAGGAGATCCTGGCTGCTCTTGAGAAAGGCTGCAGCTTCCT | 25865 | 0.10099076765165561 | No Hit |

### Adapter Content

Produced by FastQC (version 0.11.7)
