## Supplementary material for "Curated and harmonised transcriptomics datasets of interstitial lung diseases": TRIMMED_SRR8397192_fastqc.html

TRIMMED\_SRR8397192.fastq.gz FastQC Report 

FastQC Report

Sat 3 Dec 2022  
TRIMMED\_SRR8397192.fastq.gz

### Summary

- Basic Statistics
- Per base sequence quality
- Per sequence quality scores
- Per base sequence content
- Per sequence GC content
- Per base N content
- Sequence Length Distribution
- Sequence Duplication Levels
- Overrepresented sequences
- Adapter Content

### Basic Statistics

| Measure | Value |
| --- | --- |
| Filename | TRIMMED\_SRR8397192.fastq.gz |
| File type | Conventional base calls |
| Encoding | Sanger / Illumina 1.9 |
| Total Sequences | 21067458 |
| Sequences flagged as poor quality | 0 |
| Sequence length | 35-368 |
| %GC | 53 |

### Per base sequence quality

### Per sequence quality scores

### Per base sequence content

### Per sequence GC content

### Per base N content

### Sequence Length Distribution

### Sequence Duplication Levels

### Overrepresented sequences

| Sequence | Count | Percentage | Possible Source |
| --- | --- | --- | --- |
| GGGCCACGAGCTGAGTGCGTCCTGTCACTCCACTCCCATGTCCCTTGGGA | 447620 | 2.124698670337921 | No Hit |
| AAGCTCAGGGAGAGCCCTGTTAGGGCCGCCTCTGGCCCTAGTCTCAGACC | 407684 | 1.9351361706761205 | No Hit |
| ACACTGAGGACTCTGTTCCTCCCCTTTCCGCCTAGGGGAAAGTCCCCGGA | 189240 | 0.8982573977363573 | No Hit |
| TCTCTGGGCTTCTATTTCGACCGCGATGATGTGGCTCTGGAAGGCGTGAG | 166936 | 0.7923879568194702 | No Hit |
| GGGAATGTCTACGTGCGTATGCACGTGGCACTCTCTGCCCGAGGTCCGGG | 166844 | 0.7919512643623166 | No Hit |
| CCACCTTCTCCATCGGCTCCACTGGCCTCGTGGTGTATGACTACCAGCAG | 104434 | 0.49571239206932327 | No Hit |
| AGCGGTTCCGCTGCCCTGAGGCACTCTTCCAGCCTTCCTTCCTGGGCATG | 102970 | 0.4887632860120096 | No Hit |
| TCACACTTCATGATGGAGTTGAAGGTAGTTTCGTGGATGCCACAGGACTC | 91832 | 0.43589501875356773 | No Hit |
| GCTGCTACAGGAGAATAGCAGACAGGTATAGTTAAGAAACAACTTTATTT | 81689 | 0.3877496753523847 | No Hit |
| TGGTTCCTGGCAAGCCCATGTGTGTTGAGAGCTTCTCAGACTATCCACCT | 77868 | 0.3696126984090819 | No Hit |
| ATGATGTAGCAGCAGGTGCCAGGGGCTGGCTTGTAGGCGATCAGCAGCTG | 70239 | 0.33340045106533495 | No Hit |
| ACACCCACCGCAACTGTCTGTCTCATATCACGAACAGCAAAGCGACCCAA | 67764 | 0.3216524746364749 | No Hit |
| ACGCTCGTAGCCCTCGCGCTTCTCCTCGGCCAATTCGCGGAAGAAGTGGC | 64595 | 0.306610318150391 | No Hit |
| ACGCTCGTAGCCCTCGCGCTTCTCCTCGGCCAGTTCGCGGAAGAAGTGGC | 63679 | 0.302262380207427 | No Hit |
| AAAAGTGATGGACTGCCCATTGCTGGAGAAGACCTTCTCTCCTACTGTCA | 60725 | 0.28824075500708246 | No Hit |
| CTAAGTGCTGTGTTGTCGTTCCCCCTGCTTAAAATAAAGTTGTTTCTTAA | 57564 | 0.27323657177814237 | No Hit |
| ACACTGAGGACTCTGTTCCTCCCCTTTCCGCCTAGGGAAAGTCCCCGGAC | 49417 | 0.23456555603433507 | No Hit |
| GGGTACTTCAGGGTCAGGATGCCACGCTTGCTCTGGGCCTCGTCGCCCAC | 46828 | 0.22227646069117596 | No Hit |
| AGCGGTTCCGGTGTCCGGAGGCGCTGTTCCAGCCTTCCTTCCTGGGTATG | 44593 | 0.21166768197662952 | No Hit |
| ATCAAATCCTGCAGACAAGGGGAGCCCTCAGTCTGCAGGGCTCCATAATG | 41342 | 0.19623629960482178 | No Hit |
| TCACACTTCATGATGGAGTTGAAGGTGGTCTCGTGGATGCCGCAAGATTC | 41263 | 0.1958613136905269 | No Hit |
| AAAGCCAAGAAAACCAACGATGCCAGATACTGAATTTCTATAGCACTGAC | 36395 | 0.17275458671853053 | No Hit |
| TCGGGCGCCCCAGACACCAGGCGTCATGGTGGGCATGGGCCAGAAGGACT | 33911 | 0.16096389037538367 | No Hit |
| TGTTCCCTCTCCTCATGAGATTGGTGAAGAAAGTATTTGGCAAAGTTCTT | 33676 | 0.15984842594678486 | No Hit |
| TGTCCATGTCTTACTACTTTGACCGCGATGATGTGGCTTTGAAGAACTTT | 31764 | 0.15077281748941898 | No Hit |
| TCTTTCTGGCCTGGAGGCTATCCAGCGTACTCCAAAGATTCAGGTTTACT | 31193 | 0.14806247626078095 | No Hit |
| TTTACACCTGCTAGGGTGTTCAAAGGTCAGTGCTATAGAAATTCAGTATC | 30643 | 0.14545181483214537 | No Hit |
| TCTGTCTTCTTCAGTTTCGACTTATCGAATTTCTCGATCTCAGCCATATC | 28873 | 0.13705023168908181 | No Hit |
| TGAATCTTGAACTGAGTTCCACTTGTAAACTTCTTGTTTCTTGTGGTTCC | 27374 | 0.1299349926317641 | No Hit |
| CCGCCTCCGCCGCAGACGCCGCCGCGATGCGCTACGTCGCCTCCTACCTG | 26895 | 0.1276613438602797 | No Hit |
| GGCCTTCTTCTGGCGCAGAGGAAGCAGGCGCATCTGGGTGTGGGCAATGA | 25639 | 0.12169954248870461 | No Hit |
| TTGGCTTTCTCTTTCCTCTTCTCCTCCAGGGTGGCTGTCACTGCCTGGTA | 25600 | 0.12151442286012863 | No Hit |
| ACTTGATCCCAACTCATCTCTCATTTATTTCGGCTTCTTTTATTCCAGGA | 25306 | 0.12011890566009434 | No Hit |
| CGGTTGTACTTGCGGATGTAGTGCAGATAGTCTCGGCGGATGACAATGGT | 24347 | 0.11556686145998249 | No Hit |
| ACACTGAGGACTCTGTTCCTCCCCCTTTCCGCCTAGGGGAAAGTCCCCGG | 24034 | 0.11408115777423171 | No Hit |
| ACAGTGTATCTAGCTTTGGAAAACACAGGGGTCTGCCCTGTGAGCTGCTC | 23620 | 0.11211604171704057 | No Hit |
| ATCAAGACATGAGGGAGGCAGGGGCTCAGCTGAAGAAGCTGGTGGACACC | 23269 | 0.11044996505985677 | No Hit |
| TACTCGTGCGCCTCGCTTCGCTTTTCCTCCGCAACCATGTCTGACAAACC | 22854 | 0.10848010234552265 | No Hit |
| ACATAGCAATTCAGGAAATTTGACTTTCCATTCTCTGCTGGATGACGTGA | 22718 | 0.10783455697407822 | No Hit |
| AACATCTGTGTTGGGGAGAGTGGAGACAGACTGACGCGAGCAGCCAAGGT | 22359 | 0.10613050705975063 | No Hit |
| TCTCTGGCTTCTATTTCGACCGCGATGATGTGGCTCTGGAAGGCGTGAGC | 22088 | 0.10484416297400474 | No Hit |
| AAGGCCCACCCGTGGAGGCCAGACCCAAAGCCCACGGTACCGAGGACAGG | 21674 | 0.10287904691681359 | No Hit |
| TGAGGCGGAGCACCAGAGAGCCTACCTGGAAGACACATGCGTGGAGTGGC | 21270 | 0.10096139743105219 | No Hit |
| TGACTGGTGATGCTGATATTGGAGTGCATCTGAGGCTGTTGTAGAAATAA | 21259 | 0.10090918420247949 | No Hit |

### Adapter Content

Produced by FastQC (version 0.11.7)
