## Supplementary material for "Curated and harmonised transcriptomics datasets of interstitial lung diseases": TRIMMED_SRR8397206_fastqc.html

TRIMMED\_SRR8397206.fastq.gz FastQC Report 

FastQC Report

Sat 3 Dec 2022  
TRIMMED\_SRR8397206.fastq.gz

### Summary

- Basic Statistics
- Per base sequence quality
- Per sequence quality scores
- Per base sequence content
- Per sequence GC content
- Per base N content
- Sequence Length Distribution
- Sequence Duplication Levels
- Overrepresented sequences
- Adapter Content

### Basic Statistics

| Measure | Value |
| --- | --- |
| Filename | TRIMMED\_SRR8397206.fastq.gz |
| File type | Conventional base calls |
| Encoding | Sanger / Illumina 1.9 |
| Total Sequences | 27324854 |
| Sequences flagged as poor quality | 0 |
| Sequence length | 35-371 |
| %GC | 53 |

### Per base sequence quality

### Per sequence quality scores

### Per base sequence content

### Per sequence GC content

### Per base N content

### Sequence Length Distribution

### Sequence Duplication Levels

### Overrepresented sequences

| Sequence | Count | Percentage | Possible Source |
| --- | --- | --- | --- |
| AAGCTCAGGGAGAGCCCTGTTAGGGCCGCCTCTGGCCCTAGTCTCAGACC | 721311 | 2.639761588479119 | No Hit |
| GGGCCACGAGCTGAGTGCGTCCTGTCACTCCACTCCCATGTCCCTTGGGA | 684252 | 2.504137808019029 | No Hit |
| ACACTGAGGACTCTGTTCCTCCCCTTTCCGCCTAGGGGAAAGTCCCCGGA | 381798 | 1.3972554071103178 | No Hit |
| GGGAATGTCTACGTGCGTATGCACGTGGCACTCTCTGCCCGAGGTCCGGG | 381178 | 1.394986410540382 | No Hit |
| TCTCTGGGCTTCTATTTCGACCGCGATGATGTGGCTCTGGAAGGCGTGAG | 299210 | 1.0950104253073045 | No Hit |
| GCTGCTACAGGAGAATAGCAGACAGGTATAGTTAAGAAACAACTTTATTT | 144987 | 0.5306048478795166 | No Hit |
| ACACTGAGGACTCTGTTCCTCCCCTTTCCGCCTAGGGAAAGTCCCCGGAC | 138483 | 0.5068023419265113 | No Hit |
| ACGCTCGTAGCCCTCGCGCTTCTCCTCGGCCAGTTCGCGGAAGAAGTGGC | 119413 | 0.4370123990415466 | No Hit |
| ACGCTCGTAGCCCTCGCGCTTCTCCTCGGCCAATTCGCGGAAGAAGTGGC | 118426 | 0.43340030288908404 | No Hit |
| CCACCTTCTCCATCGGCTCCACTGGCCTCGTGGTGTATGACTACCAGCAG | 111733 | 0.40890611894943707 | No Hit |
| TGGTTCCTGGCAAGCCCATGTGTGTTGAGAGCTTCTCAGACTATCCACCT | 101425 | 0.37118222113830873 | No Hit |
| ACACCCACCGCAACTGTCTGTCTCATATCACGAACAGCAAAGCGACCCAA | 95704 | 0.3502452382728193 | No Hit |
| TCTTTCTGGCCTGGAGGCTATCCAGCGTACTCCAAAGATTCAGGTTTACT | 87601 | 0.3205909169725115 | No Hit |
| TCTGTCTTCTTCAGTTTCGACTTATCGAATTTCTCGATCTCAGCCATATC | 75333 | 0.2756940622628761 | No Hit |
| ATGATGTAGCAGCAGGTGCCAGGGGCTGGCTTGTAGGCGATCAGCAGCTG | 71453 | 0.2614945353413416 | No Hit |
| TACTCGTGCGCCTCGCTTCGCTTTTCCTCCGCAACCATGTCTGACAAACC | 65220 | 0.23868380046971155 | No Hit |
| ACATAGCAATTCAGGAAATTTGACTTTCCATTCTCTGCTGGATGACGTGA | 64049 | 0.23439832469004224 | No Hit |
| AAAAGTGATGGACTGCCCATTGCTGGAGAAGACCTTCTCTCCTACTGTCA | 62863 | 0.23005795383206806 | No Hit |
| TGTTCCCTCTCCTCATGAGATTGGTGAAGAAAGTATTTGGCAAAGTTCTT | 60614 | 0.22182735175821983 | No Hit |
| CTAAGTGCTGTGTTGTCGTTCCCCCTGCTTAAAATAAAGTTGTTTCTTAA | 60051 | 0.21976695648584255 | No Hit |
| TGTCCATGTCTTACTACTTTGACCGCGATGATGTGGCTTTGAAGAACTTT | 59651 | 0.21830308773104515 | No Hit |
| TCACACTTCATGATGGAGTTGAAGGTAGTTTCGTGGATGCCACAGGACTC | 56803 | 0.20788034219688784 | No Hit |
| AAGACCAACGTCAAGGCCGCCTGGGGTAAGGTCGGCGCGCACGCTGGCGA | 53809 | 0.19692328456722952 | No Hit |
| ATCAAATCCTGCAGACAAGGGGAGCCCTCAGTCTGCAGGGCTCCATAATG | 52286 | 0.19134960428333853 | No Hit |
| CCCAGGGCAGAGGCTGACGACGCTGACTGGCAGGGCACCGACACAGTGTC | 50034 | 0.18310802319382932 | No Hit |
| ACACTGAGGACTCTGTTCCTCCCCTTTCCGCCTAGGGGAAAGTCCCCGGC | 49413 | 0.18083536695200642 | No Hit |
| AGCGGTTCCGCTGCCCTGAGGCACTCTTCCAGCCTTCCTTCCTGGGCATG | 48430 | 0.17723790948709187 | No Hit |
| ACTTGATCCCAACTCATCTCTCATTTATTTCGGCTTCTTTTATTCCAGGA | 45753 | 0.16744096784561044 | No Hit |
| TCAGCACGGGCTCGTACTCTGCCACAAACTGATCACACTGCTTCTGGTAA | 45037 | 0.16482064277452316 | No Hit |
| AAAGCCAAGAAAACCAACGATGCCAGATACTGAATTTCTATAGCACTGAC | 44528 | 0.1629578697840435 | No Hit |
| TGAATCTTGAACTGAGTTCCACTTGTAAACTTCTTGTTTCTTGTGGTTCC | 43410 | 0.15886635661438483 | No Hit |
| TTTACACCTGCTAGGGTGTTCAAAGGTCAGTGCTATAGAAATTCAGTATC | 42668 | 0.1561508800742357 | No Hit |
| GGGTACTTCAGGGTCAGGATGCCACGCTTGCTCTGGGCCTCGTCGCCCAC | 40890 | 0.14964398345916138 | No Hit |
| TCTCTGGCTTCTATTTCGACCGCGATGATGTGGCTCTGGAAGGCGTGAGC | 37886 | 0.13865032911063313 | No Hit |
| TACATCAAAGATTACATGAAATCAATCAAAGGGAAACTTGAAGAACAGAG | 37179 | 0.13606294108652878 | No Hit |
| AGCACCAAGCAGGAGATCCTGGCTGCTCTTGAGAAAGGCTGCAGCTTCCT | 35482 | 0.1298524778943009 | No Hit |
| ACAGTGTATCTAGCTTTGGAAAACACAGGGGTCTGCCCTGTGAGCTGCTC | 34966 | 0.1279640872006123 | No Hit |
| GGCCTTCTTCTGGCGCAGAGGAAGCAGGCGCATCTGGGTGTGGGCAATGA | 34852 | 0.12754688460549507 | No Hit |
| ACACTGAGGACTCTGTTCCTCCCCCTTTCCGCCTAGGGGAAAGTCCCCGG | 34326 | 0.1256218971929365 | No Hit |
| AACATCTGTGTTGGGGAGAGTGGAGACAGACTGACGCGAGCAGCCAAGGT | 33785 | 0.12364201470207306 | No Hit |
| TCGGGCGCCCCAGACACCAGGCGTCATGGTGGGCATGGGCCAGAAGGACT | 33484 | 0.12254045346408804 | No Hit |
| TTCATGGTCTCGTCCATCAGCGCCCTCAGTTCCTGGGTGACCTGGGAGCT | 33332 | 0.12198418333726503 | No Hit |
| CGGTTGTACTTGCGGATGTAGTGCAGATAGTCTCGGCGGATGACAATGGT | 32927 | 0.1205020162230327 | No Hit |
| CTGGATGTTGAGCAGGAACGCAGTCTTGGCCTCTTCTATCACCCTCTGCC | 32463 | 0.11880392846746773 | No Hit |
| AATAAAAGCGAAAAGAAATGAAAATGTTACACTACATTAATCCTGGAATA | 32374 | 0.11847821766952533 | No Hit |
| ACACTGAGGACTCTGTTCCTCCCCTTTCCGCCTAGGGAAAGTCCCCGGCA | 31559 | 0.11549558508162569 | No Hit |
| AATACCAAGACCTGCTCAATGTTAAGATGGCCCTTGACATTGAGATTGCC | 30108 | 0.11018540117359822 | No Hit |
| TACAACCAAAAAAGTTGCTGGATGTTTCTTTGACTACTGGAACCACAAGA | 29814 | 0.10910945763882215 | No Hit |
| GGGAATGTCTACGTGCGTATGCACGTGGCACTCTCTGCCCGAGGTCCCGG | 29509 | 0.10799325771328916 | No Hit |
| TTGGCTTTCTCTTTCCTCTTCTCCTCCAGGGTGGCTGTCACTGCCTGGTA | 29442 | 0.10774805969686059 | No Hit |
| ATTCGAGGGCGGATCCTCTCTGGCGTGGTGACCAAGATGAAGATGCAGAG | 28875 | 0.10567302573693531 | No Hit |
| ACCAATGTCATAGGGTGCAATATCTACAATAGGTAGTCTCACAGCCTTGC | 28853 | 0.10559251295542146 | No Hit |
| AAGGCCCACCCGTGGAGGCCAGACCCAAAGCCCACGGTACCGAGGACAGG | 28791 | 0.10536561329842788 | No Hit |
| GAAAAGTTTGGAAGAGGCAGAGAAATCCTGCTCTCCTCGCCTTCCAGCAG | 28441 | 0.10408472813798017 | No Hit |
| AGCGGTTCCGGTGTCCGGAGGCGCTGTTCCAGCCTTCCTTCCTGGGTATG | 27962 | 0.10233174530411032 | No Hit |
| GCCAAGCACAAAGAGCTTGCTCCCTACGATGAGAACTGGTTCTACACGCG | 27500 | 0.10064097689231935 | No Hit |

### Adapter Content

Produced by FastQC (version 0.11.7)
