## Supplementary material for "Curated and harmonised transcriptomics datasets of interstitial lung diseases": TRIMMED_SRR8397213_fastqc.html

TRIMMED\_SRR8397213.fastq.gz FastQC Report 

FastQC Report

Sat 3 Dec 2022  
TRIMMED\_SRR8397213.fastq.gz

### Summary

- Basic Statistics
- Per base sequence quality
- Per sequence quality scores
- Per base sequence content
- Per sequence GC content
- Per base N content
- Sequence Length Distribution
- Sequence Duplication Levels
- Overrepresented sequences
- Adapter Content

### Basic Statistics

| Measure | Value |
| --- | --- |
| Filename | TRIMMED\_SRR8397213.fastq.gz |
| File type | Conventional base calls |
| Encoding | Sanger / Illumina 1.9 |
| Total Sequences | 8078760 |
| Sequences flagged as poor quality | 0 |
| Sequence length | 35-354 |
| %GC | 53 |

### Per base sequence quality

### Per sequence quality scores

### Per base sequence content

### Per sequence GC content

### Per base N content

### Sequence Length Distribution

### Sequence Duplication Levels

### Overrepresented sequences

| Sequence | Count | Percentage | Possible Source |
| --- | --- | --- | --- |
| AAGCTCAGGGAGAGCCCTGTTAGGGCCGCCTCTGGCCCTAGTCTCAGACC | 162939 | 2.016881303566389 | No Hit |
| GGGCCACGAGCTGAGTGCGTCCTGTCACTCCACTCCCATGTCCCTTGGGA | 147041 | 1.820093677742624 | No Hit |
| ACACTGAGGACTCTGTTCCTCCCCTTTCCGCCTAGGGGAAAGTCCCCGGA | 75333 | 0.9324822126167877 | No Hit |
| GGGAATGTCTACGTGCGTATGCACGTGGCACTCTCTGCCCGAGGTCCGGG | 70739 | 0.8756170501413583 | No Hit |
| GCTGCTACAGGAGAATAGCAGACAGGTATAGTTAAGAAACAACTTTATTT | 32786 | 0.4058296075140244 | No Hit |
| ACACTGAGGACTCTGTTCCTCCCCTTTCCGCCTAGGGAAAGTCCCCGGAC | 28875 | 0.3574187127727523 | No Hit |
| TCTCTGGGCTTCTATTTCGACCGCGATGATGTGGCTCTGGAAGGCGTGAG | 23795 | 0.2945377755992256 | No Hit |
| ATCAAGACATGAGGGAGGCAGGGGCTCAGCTGAAGAAGCTGGTGGACACC | 21404 | 0.2649416494610559 | No Hit |
| TGGTTCCTGGCAAGCCCATGTGTGTTGAGAGCTTCTCAGACTATCCACCT | 21177 | 0.2621318123078294 | No Hit |
| ACACCCACCGCAACTGTCTGTCTCATATCACGAACAGCAAAGCGACCCAA | 20357 | 0.2519817397719452 | No Hit |
| CTAAGTGCTGTGTTGTCGTTCCCCCTGCTTAAAATAAAGTTGTTTCTTAA | 18298 | 0.22649515519708469 | No Hit |
| TCACACTTCATGATGGAGTTGAAGGTAGTTTCGTGGATGCCACAGGACTC | 15686 | 0.1941634607291218 | No Hit |
| AGCGGTTCCGCTGCCCTGAGGCACTCTTCCAGCCTTCCTTCCTGGGCATG | 14428 | 0.17859176408260674 | No Hit |
| GGGCCACGAGCTGAGTGCGTCCTGTCACTCCACTCCCCATGTCCCTTGGG | 13893 | 0.17196946065980423 | No Hit |
| TCTTTCTGGCCTGGAGGCTATCCAGCGTACTCCAAAGATTCAGGTTTACT | 13360 | 0.16537191351147948 | No Hit |
| GACAGACTCCAGCTTTGAAGGACTTTCCAGAGCCTTCCACAGCCCAAGGT | 13294 | 0.1645549564537132 | No Hit |
| ACACCATTCAGCTCTACCTGGGGGCCAAGTTGTTGGACTCACAGGGAAAG | 13244 | 0.16393604959176902 | No Hit |
| AGCTTAATGATGCTTTCTCTGGGCTTTTGGGGGAGGGTGTCCACCAGCTT | 13189 | 0.16325525204363045 | No Hit |
| TGAGTTGGCCAGAACTCTGAAAAGATTGGGAATGGATGGCTACAGGGGAA | 12375 | 0.15317944833117952 | No Hit |
| TGAAGTCCAGCGGCCTCTTCCCCTTCCTGGTGCTGCTTGCCCTGGGAACT | 12165 | 0.15058003951101406 | No Hit |
| GGCCTTCTTCTGGCGCAGAGGAAGCAGGCGCATCTGGGTGTGGGCAATGA | 12084 | 0.14957741039466452 | No Hit |
| GGCATTGTCAGGGAAGCTGCAGAGTTATTGAACCACTTGGTCACCTTTCC | 11527 | 0.14268278795260658 | No Hit |
| ACATAGCAATTCAGGAAATTTGACTTTCCATTCTCTGCTGGATGACGTGA | 10858 | 0.13440181413979374 | No Hit |
| AAAGCCAAGAAAACCAACGATGCCAGATACTGAATTTCTATAGCACTGAC | 10723 | 0.1327307656125445 | No Hit |
| ACACTGAGGACTCTGTTCCTCCCCCTTTCCGCCTAGGGGAAAGTCCCCGG | 10505 | 0.13003233169446796 | No Hit |
| TTTACACCTGCTAGGGTGTTCAAAGGTCAGTGCTATAGAAATTCAGTATC | 10244 | 0.12680163787511944 | No Hit |
| GGGTACTTCAGGGTCAGGATGCCACGCTTGCTCTGGGCCTCGTCGCCCAC | 10190 | 0.12613321846421977 | No Hit |
| ACGCTCGTAGCCCTCGCGCTTCTCCTCGGCCAGTTCGCGGAAGAAGTGGC | 9929 | 0.12290252464487125 | No Hit |
| ACGCTCGTAGCCCTCGCGCTTCTCCTCGGCCAATTCGCGGAAGAAGTGGC | 9548 | 0.11818645435685675 | No Hit |
| ACTTGATCCCAACTCATCTCTCATTTATTTCGGCTTCTTTTATTCCAGGA | 9407 | 0.1164411370061742 | No Hit |
| TCGGGCGCCCCAGACACCAGGCGTCATGGTGGGCATGGGCCAGAAGGACT | 9138 | 0.11311141808891464 | No Hit |
| TCACACTTCATGATGGAGTTGAAGGTGGTCTCGTGGATGCCGCAAGATTC | 8911 | 0.11030158093568816 | No Hit |
| CAGCTGGAGAAGGACTTCAGCAGCATGAAGAAGTACTGCCAAGTCATCCG | 8804 | 0.10897712025112764 | No Hit |
| TTGGCTTTCTCTTTCCTCTTCTCCTCCAGGGTGGCTGTCACTGCCTGGTA | 8313 | 0.102899454866836 | No Hit |
| TAGGGGCCGGTGGACCTGCTCCAGCAGCTGGTGCTGCACCAGCAGGAGGT | 8279 | 0.10247859820071398 | No Hit |
| CGCTGCACTGACCTTCTTCCAAGCCTCAGTTCCTGTTCTAGGAACTTGAG | 8119 | 0.10049809624249266 | No Hit |

### Adapter Content

Produced by FastQC (version 0.11.7)
