## Supplementary material for "Curated and harmonised transcriptomics datasets of interstitial lung diseases": TRIMMED_SRR8397228_fastqc.html

TRIMMED\_SRR8397228.fastq.gz FastQC Report 

FastQC Report

Sat 3 Dec 2022  
TRIMMED\_SRR8397228.fastq.gz

### Summary

- Basic Statistics
- Per base sequence quality
- Per sequence quality scores
- Per base sequence content
- Per sequence GC content
- Per base N content
- Sequence Length Distribution
- Sequence Duplication Levels
- Overrepresented sequences
- Adapter Content

### Basic Statistics

| Measure | Value |
| --- | --- |
| Filename | TRIMMED\_SRR8397228.fastq.gz |
| File type | Conventional base calls |
| Encoding | Sanger / Illumina 1.9 |
| Total Sequences | 4671012 |
| Sequences flagged as poor quality | 0 |
| Sequence length | 35-370 |
| %GC | 51 |

### Per base sequence quality

### Per sequence quality scores

### Per base sequence content

### Per sequence GC content

### Per base N content

### Sequence Length Distribution

### Sequence Duplication Levels

### Overrepresented sequences

| Sequence | Count | Percentage | Possible Source |
| --- | --- | --- | --- |
| GGGCCACGAGCTGAGTGCGTCCTGTCACTCCACTCCCATGTCCCTTGGGA | 41819 | 0.8952877877427847 | No Hit |
| GGGAATGTCTACGTGCGTATGCACGTGGCACTCTCTGCC | 23674 | 0.5068280706622034 | No Hit |
| AGCGGTTCCGCTGCCCTGAGGCACTCTTCCAGCCTTCCTTCCTGGGCATG | 21282 | 0.4556186111275244 | No Hit |
| AAGCTCAGGGAGAGCCCTGTTAGGGCCGCCTCTGGCCCTAGTCTCAGACC | 21141 | 0.45259999332050527 | No Hit |
| TCTCTGGGCTTCTATTTCGACCGCGATGATGTGGCTCTGGAAGGCGTGAG | 20914 | 0.4477402327375738 | No Hit |
| ACACTGAGGACTCTGTTCCTCCCCTTTCCGCCTAGGGGAAAGTCCCCGGA | 20112 | 0.4305705059203444 | No Hit |
| TCACACTTCATGATGGAGTTGAAGGTAGTTTCGTGGATGCCACAGGACTC | 18445 | 0.39488230815934533 | No Hit |
| ATCAAGACATGAGGGAGGCAGGGGCTCAGCTGAAGAAGCTGGTGGACACC | 17950 | 0.3842850328793846 | No Hit |
| AAACTCAGGGAGAGCCCTGTTAGGGCCGCCTCTGGCCCTAGTCTCAGACC | 16621 | 0.3558329544004597 | No Hit |
| TCTTTCTGGCCTGGAGGCTATCCAGCGTACTCCAAAGATTCAGGTTTACT | 15011 | 0.32136504894442575 | No Hit |
| TGGTTCCTGGCAAGCCCATGTGTGTTGAGAGCTTCTCAGACTATCCACCT | 14154 | 0.3030178470960897 | No Hit |
| CCACCTTCTCCATCGGCTCCACTGGCCTCGTGGTGTATGACTACCAGCAG | 12727 | 0.27246772219810184 | No Hit |
| ATGATGTAGCAGCAGGTGCCAGGGGCTGGCTTGTAGGCGATCAGCAGCTG | 12719 | 0.2722964531026681 | No Hit |
| ACACCCACCGCAACTGTCTGTCTCATATCACGAACAGCAAAGCGACCCAA | 12666 | 0.2711617953454198 | No Hit |
| CTAAGTGCTGTGTTGTCGTTCCCCCTGCTTAAAATAAAGTTGTTTCTTAA | 11941 | 0.2556405335717399 | No Hit |
| AGCTTAATGATGCTTTCTCTGGGCTTTTGGGGGAGGGTGTCCACCAGCTT | 11269 | 0.2412539295553084 | No Hit |
| TCTGTCTTCTTCAGTTTCGACTTATCGAATTTCTCGATCTCAGCCATATC | 11155 | 0.23881334494537798 | No Hit |
| ACTTGATCCCAACTCATCTCTCATTTATTTCGGCTTCTTTTATTCCAGGA | 10709 | 0.22926509287494873 | No Hit |
| GCTGCTACAGGAGAATAGCAGACAGGTATAGTTAAGAAACAACTTTATTT | 10337 | 0.22130107993728124 | No Hit |
| GGGAATGTCTACGTGCGTATGCACGTGGCACTCTCT | 9390 | 0.20102710076531594 | No Hit |
| AAAAGTGATGGACTGCCCATTGCTGGAGAAGACCTTCTCTCCTACTGTCA | 9255 | 0.1981369347798721 | No Hit |
| ACATAGCAATTCAGGAAATTTGACTTTCCATTCTCTGCTGGATGACGTGA | 9005 | 0.1927847755475687 | No Hit |
| TGTTCCCTCTCCTCATGAGATTGGTGAAGAAAGTATTTGGCAAAGTTCTT | 8189 | 0.1753153278133304 | No Hit |
| TGTCCATGTCTTACTACTTTGACCGCGATGATGTGGCTTTGAAGAACTTT | 7652 | 0.16381888978234269 | No Hit |
| AGCGGTTCCGGTGTCCGGAGGCGCTGTTCCAGCCTTCCTTCCTGGGTATG | 7589 | 0.16247014565580223 | No Hit |
| TACATCAAAGATTACATGAAATCAATCAAAGGGAAACTTGAAGAACAGAG | 7579 | 0.16225605928651007 | No Hit |
| TACTCGTGCGCCTCGCTTCGCTTTTCCTCCGCAACCATGTCTGACAAACC | 7358 | 0.15752475052515386 | No Hit |
| AAAGAAGAACATGTGATCATCCAGGCCGAGTTCTATCTGAATCCTGACCA | 7047 | 0.1508666644401684 | No Hit |
| ACGCTCGTAGCCCTCGCGCTTCTCCTCGGCCAGTTCGCGGAAGAAGTGGC | 6909 | 0.14791227254393693 | No Hit |
| AAAGCCAAGAAAACCAACGATGCCAGATACTGAATTTCTATAGCACTGAC | 6494 | 0.1390276882183133 | No Hit |
| ACGCTCGTAGCCCTCGCGCTTCTCCTCGGCCAATTCGCGGAAGAAGTGGC | 6482 | 0.13877078457516273 | No Hit |
| AAGACCAACGTCAAGGCCGCCTGGGGTAAGGTCGGCGCGCACGCTGGCGA | 6054 | 0.1296078879694593 | No Hit |
| TCGGGCGCCCCAGACACCAGGCGTCATGGTGGGCATGGGCCAGAAGGACT | 6009 | 0.12864449930764468 | No Hit |
| TTTACACCTGCTAGGGTGTTCAAAGGTCAGTGCTATAGAAATTCAGTATC | 5995 | 0.1283447783906357 | No Hit |
| TCGGCTCAGCCAAACACTGTCAGGGCCCCCAGCAGGGCCTTCAGGGCCTT | 5855 | 0.1253475692205458 | No Hit |
| TGAATCTTGAACTGAGTTCCACTTGTAAACTTCTTGTTTCTTGTGGTTCC | 5782 | 0.1237847387247132 | No Hit |
| AATAAAAGCGAAAAGAAATGAAAATGTTACACTACATTAATCCTGGAATA | 5684 | 0.12168669230565025 | No Hit |
| GGCTCCCAGAAGTGTGTGGCTGAGCTGGGTCCCCAGGCCGTGGGGGCCGT | 5518 | 0.11813285857540079 | No Hit |
| ACACTGAGGACTCTGTTCCTCCCCTTTCCGCCTAGGGAAAGGTACCCGGC | 5367 | 0.11490015439908953 | No Hit |
| GGGTACTTCAGGGTCAGGATGCCACGCTTGCTCTGGGCCTCGTCGCCCAC | 5319 | 0.11387253982648729 | No Hit |
| ATCAAATCCTGCAGACAAGGGGAGCCCTCAGTCTGCAGGGCTCCATAATG | 5258 | 0.11256661297380526 | No Hit |
| TCACACTTCATGATGGAGTTGAAGGTGGTCTCGTGGATGCCGCAAGATTC | 5089 | 0.10894855333276815 | No Hit |
| ACACTGAGGACTCTGTTCCTCCCCTTTCCGCCTAGGGAAAGTCCCCGGAC | 4969 | 0.1063795169012625 | No Hit |
| GGCCTTCTTCTGGCGCAGAGGAAGCAGGCGCATCTGGGTGTGGGCAATGA | 4694 | 0.10049214174572876 | No Hit |

### Adapter Content

Produced by FastQC (version 0.11.7)
