## Supplementary material for "Curated and harmonised transcriptomics datasets of interstitial lung diseases": TRIMMED_SRR8397234_fastqc.html

TRIMMED\_SRR8397234.fastq.gz FastQC Report 

FastQC Report

Sat 3 Dec 2022  
TRIMMED\_SRR8397234.fastq.gz

### Summary

- Basic Statistics
- Per base sequence quality
- Per sequence quality scores
- Per base sequence content
- Per sequence GC content
- Per base N content
- Sequence Length Distribution
- Sequence Duplication Levels
- Overrepresented sequences
- Adapter Content

### Basic Statistics

| Measure | Value |
| --- | --- |
| Filename | TRIMMED\_SRR8397234.fastq.gz |
| File type | Conventional base calls |
| Encoding | Sanger / Illumina 1.9 |
| Total Sequences | 6785433 |
| Sequences flagged as poor quality | 0 |
| Sequence length | 35-367 |
| %GC | 54 |

### Per base sequence quality

### Per sequence quality scores

### Per base sequence content

### Per sequence GC content

### Per base N content

### Sequence Length Distribution

### Sequence Duplication Levels

### Overrepresented sequences

| Sequence | Count | Percentage | Possible Source |
| --- | --- | --- | --- |
| AAGCTCAGGGAGAGCCCTGTTAGGGCCGCCTCTGGCCCTAGTCTCAGACC | 97281 | 1.4336741664091295 | No Hit |
| GGGCCACGAGCTGAGTGCGTCCTGTCACTCCACTCCCATGTCCCTTGGGA | 87176 | 1.2847522037281924 | No Hit |
| ACACTGAGGACTCTGTTCCTCCCCTTTCCGCCTAGGGGAAAGTCCCCGGA | 54919 | 0.8093661819370996 | No Hit |
| GGGAATGTCTACGTGCGTATGCACGTGGCACTCTCTGCCCGAGGTCCGGG | 42324 | 0.623747961257594 | No Hit |
| TCACACTTCATGATGGAGTTGAAGGTAGTTTCGTGGATGCCACAGGACTC | 33066 | 0.4873086212773746 | No Hit |
| AGCGGTTCCGCTGCCCTGAGGCACTCTTCCAGCCTTCCTTCCTGGGCATG | 27285 | 0.4021114054180478 | No Hit |
| TGGTTCCTGGCAAGCCCATGTGTGTTGAGAGCTTCTCAGACTATCCACCT | 27017 | 0.39816176801097297 | No Hit |
| ATCAAGACATGAGGGAGGCAGGGGCTCAGCTGAAGAAGCTGGTGGACACC | 26202 | 0.38615074380662223 | No Hit |
| AGAGCATGACCGATGGATTCCAGTTCGAGTATGGCGGCCAGGGCTCCGAC | 26125 | 0.3850159599247388 | No Hit |
| GGGTACTTCAGGGTCAGGATGCCACGCTTGCTCTGGGCCTCGTCGCCCAC | 24793 | 0.3653856725134564 | No Hit |
| AGCTTAATGATGCTTTCTCTGGGCTTTTGGGGGAGGGTGTCCACCAGCTT | 24237 | 0.35719164863907726 | No Hit |
| CCACCTTCTCCATCGGCTCCACTGGCCTCGTGGTGTATGACTACCAGCAG | 23842 | 0.3513703546995453 | No Hit |
| ACACTGAGGACTCTGTTCCTCCCCTTTCCGCCTAGGGAAAGTCCCCGGAC | 23341 | 0.34398689074079725 | No Hit |
| ACACCCACCGCAACTGTCTGTCTCATATCACGAACAGCAAAGCGACCCAA | 20760 | 0.30594952451818475 | No Hit |
| ATGTTCTGGGAGGCTCGGTGGACATCAGGCGCAGGAAGGTCAGCTGGATG | 18199 | 0.26820690735580177 | No Hit |
| ATGATGTAGCAGCAGGTGCCAGGGGCTGGCTTGTAGGCGATCAGCAGCTG | 17872 | 0.26338776022105 | No Hit |
| ATGTTCTGGGAGCTCGGTGGACATCAGGCGCAGGAAGGTCAGCTGGATGG | 16471 | 0.24274058855197597 | No Hit |
| TACTCGTGCGCCTCGCTTCGCTTTTCCTCCGCAACCATGTCTGACAAACC | 14091 | 0.20766545038466963 | No Hit |
| TCACACTTCATGATGGAGTTGAAGGTGGTCTCGTGGATGCCGCAAGATTC | 13801 | 0.20339158901134238 | No Hit |
| ACCAATGTCATAGGGTGCAATATCTACAATAGGTAGTCTCACAGCCTTGC | 12675 | 0.1867972169204235 | No Hit |
| ACGCTCGTAGCCCTCGCGCTTCTCCTCGGCCAGTTCGCGGAAGAAGTGGC | 12027 | 0.17724734736898884 | No Hit |
| GCTGCTACAGGAGAATAGCAGACAGGTATAGTTAAGAAACAACTTTATTT | 11926 | 0.17575886461483003 | No Hit |
| TCGGGCGCCCCAGACACCAGGCGTCATGGTGGGCATGGGCCAGAAGGACT | 11583 | 0.1707039182318947 | No Hit |
| TCTTTCTGGCCTGGAGGCTATCCAGCGTACTCCAAAGATTCAGGTTTACT | 11317 | 0.16678375573084284 | No Hit |
| AGCGGTTCCGGTGTCCGGAGGCGCTGTTCCAGCCTTCCTTCCTGGGTATG | 11213 | 0.1652510606176496 | No Hit |
| GGGGCCACGAGCTGAGTGCGTCCTGTCACTCCACTCCCATGTCCCTTGGG | 11075 | 0.1632172921020663 | No Hit |
| ACGCTCGTAGCCCTCGCGCTTCTCCTCGGCCAATTCGCGGAAGAAGTGGC | 10980 | 0.16181723406597634 | No Hit |
| ACATAGCAATTCAGGAAATTTGACTTTCCATTCTCTGCTGGATGACGTGA | 10416 | 0.1535053105675054 | No Hit |
| AAGACCAACGTCAAGGCCGCCTGGGGTAAGGTCGGCGCGCACGCTGGCGA | 10178 | 0.14999779675077476 | No Hit |
| TTGGTGGTGGGGAAGGACAGGAACATCCTCTCCAGGGCCTCCGCACCATA | 10174 | 0.1499388469387289 | No Hit |
| GTCCGACGCACCTGTTTGCAGGCTGCCTGGCGCTCCTTGGTCTCGTTGTC | 10098 | 0.14881880050985694 | No Hit |
| ACCCGCAGCTTCTGCTTCTCAGTCAGAAGGTTGTTGTCCTCATCCCTCTC | 9731 | 0.14341015525464626 | No Hit |
| TTGGCTTTCTCTTTCCTCTTCTCCTCCAGGGTGGCTGTCACTGCCTGGTA | 9646 | 0.14215747174867102 | No Hit |
| GGCCTTCTTCTGGCGCAGAGGAAGCAGGCGCATCTGGGTGTGGGCAATGA | 9228 | 0.1359972163898752 | No Hit |
| GCCAAGCACAAAGAGCTTGCTCCCTACGATGAGAACTGGTTCTACACGCG | 9077 | 0.13377186098514274 | No Hit |
| ACCTACGGCGAGGGCGAGAGCGGCCCCATGGGCAACATCATGATCGATCC | 8984 | 0.13240127785507572 | No Hit |
| GGCTCCCAGAAGTGTGTGGCTGAGCTGGGTCCCCAGGCCGTGGGGGCCGT | 8859 | 0.13055909622864156 | No Hit |
| TCTGTCTTCTTCAGTTTCGACTTATCGAATTTCTCGATCTCAGCCATATC | 8684 | 0.12798004195163373 | No Hit |
| ATCACCGCCCTCGAGGCCAAGATTGCACAGCTGGAGGAGCAGCTGGACAA | 8682 | 0.1279505670456108 | No Hit |
| ACAGTGTATCTAGCTTTGGAAAACACAGGGGTCTGCCCTGTGAGCTGCTC | 8677 | 0.12787687978055343 | No Hit |
| ACACTGAGGACTCTGTTCCTCCCCCTTTCCGCCTAGGGGAAAGTCCCCGG | 8371 | 0.12336721915904261 | No Hit |
| TCGGCTCAGCCAAACACTGTCAGGGCCCCCAGCAGGGCCTTCAGGGCCTT | 7980 | 0.11760487503155656 | No Hit |
| AAGGCCCACCCGTGGAGGCCAGACCCAAAGCCCACGGTACCGAGGACAGG | 7924 | 0.11677957766291407 | No Hit |
| TGGTTGCACGAAACACACTGGGGAATGGAGCAAAACAGTCTTTGAATATC | 7787 | 0.11476054660034224 | No Hit |
| ATTCGAGGGCGGATCCTCTCTGGCGTGGTGACCAAGATGAAGATGCAGAG | 7703 | 0.11352260054737849 | No Hit |
| AAAAGTGATGGACTGCCCATTGCTGGAGAAGACCTTCTCTCCTACTGTCA | 7470 | 0.1100887739957052 | No Hit |
| AACATCTGTGTTGGGGAGAGTGGAGACAGACTGACGCGAGCAGCCAAGGT | 7133 | 0.10512225233083872 | No Hit |
| GGCTGCCTGGAGCCCCTGGTGTCCCTGGAGAGCGTGGAGAGAAGGGGGAG | 7080 | 0.10434116732123064 | No Hit |
| TCAGCACGGGCTCGTACTCTGCCACAAACTGATCACACTGCTTCTGGTAA | 7022 | 0.10348639504656519 | No Hit |
| ACACTGAGGACTCTGTTCCTCCCCTTCCGCCTAGGGAAAGTCCCCGGACC | 6953 | 0.10246951078877353 | No Hit |

### Adapter Content

Produced by FastQC (version 0.11.7)
