## Supplementary material for "Curated and harmonised transcriptomics datasets of interstitial lung diseases": TRIMMED_SRR8397244_fastqc.html

TRIMMED\_SRR8397244.fastq.gz FastQC Report 

FastQC Report

Sat 3 Dec 2022  
TRIMMED\_SRR8397244.fastq.gz

### Summary

- Basic Statistics
- Per base sequence quality
- Per sequence quality scores
- Per base sequence content
- Per sequence GC content
- Per base N content
- Sequence Length Distribution
- Sequence Duplication Levels
- Overrepresented sequences
- Adapter Content

### Basic Statistics

| Measure | Value |
| --- | --- |
| Filename | TRIMMED\_SRR8397244.fastq.gz |
| File type | Conventional base calls |
| Encoding | Sanger / Illumina 1.9 |
| Total Sequences | 23147628 |
| Sequences flagged as poor quality | 0 |
| Sequence length | 35-367 |
| %GC | 52 |

### Per base sequence quality

### Per sequence quality scores

### Per base sequence content

### Per sequence GC content

### Per base N content

### Sequence Length Distribution

### Sequence Duplication Levels

### Overrepresented sequences

| Sequence | Count | Percentage | Possible Source |
| --- | --- | --- | --- |
| AAGCTCAGGGAGAGCCCTGTTAGGGCCGCCTCTGGCCCTAGTCTCAGACC | 498851 | 2.155084745616268 | No Hit |
| GGGCCACGAGCTGAGTGCGTCCTGTCACTCCACTCCCATGTCCCTTGGGA | 494043 | 2.134313718882989 | No Hit |
| GGGAATGTCTACGTGCGTATGCACGTGGCACTCTCTGCCCGAGGTCCGGG | 120049 | 0.5186233336737569 | No Hit |
| GCTGCTACAGGAGAATAGCAGACAGGTATAGTTAAGAAACAACTTTATTT | 118866 | 0.5135126588348491 | No Hit |
| ACACTGAGGACTCTGTTCCTCCCCTTTCCGCCTAGGGGAAAGTCCCCGGA | 117127 | 0.5060000100226252 | No Hit |
| CTAAGTGCTGTGTTGTCGTTCCCCCTGCTTAAAATAAAGTTGTTTCTTAA | 93727 | 0.40490973848378764 | No Hit |
| TGGTTCCTGGCAAGCCCATGTGTGTTGAGAGCTTCTCAGACTATCCACCT | 80801 | 0.34906816370126564 | No Hit |
| ACTTGATCCCAACTCATCTCTCATTTATTTCGGCTTCTTTTATTCCAGGA | 71473 | 0.3087702981921085 | No Hit |
| ACACCCACCGCAACTGTCTGTCTCATATCACGAACAGCAAAGCGACCCAA | 66583 | 0.28764502349873605 | No Hit |
| AGCGGTTCCGCTGCCCTGAGGCACTCTTCCAGCCTTCCTTCCTGGGCATG | 61413 | 0.2653101216245569 | No Hit |
| TCACACTTCATGATGGAGTTGAAGGTAGTTTCGTGGATGCCACAGGACTC | 59298 | 0.25617311631239276 | No Hit |
| TCTCTGGGCTTCTATTTCGACCGCGATGATGTGGCTCTGGAAGGCGTGAG | 53997 | 0.2332722817214792 | No Hit |
| AAAGCCAAGAAAACCAACGATGCCAGATACTGAATTTCTATAGCACTGAC | 53568 | 0.2314189600766005 | No Hit |
| TTTACACCTGCTAGGGTGTTCAAAGGTCAGTGCTATAGAAATTCAGTATC | 51623 | 0.2230163712670689 | No Hit |
| TCTTTCTGGCCTGGAGGCTATCCAGCGTACTCCAAAGATTCAGGTTTACT | 45500 | 0.1965644168810731 | No Hit |
| TGAGTTGGCCAGAACTCTGAAAAGATTGGGAATGGATGGCTACAGGGGAA | 44057 | 0.19033051680284477 | No Hit |
| GGGCCACGAGCTGAGTGCGTCCTGTCACTCCACTCCCCATGTCCCTTGGG | 43128 | 0.18631714662081142 | No Hit |
| TCACTGATCAGACACACTAGTGTGGCCTTGTTGGCTTGGAGCTCCTCAGA | 41535 | 0.1794352319814367 | No Hit |
| CCACCTTCTCCATCGGCTCCACTGGCCTCGTGGTGTATGACTACCAGCAG | 40582 | 0.17531817946961997 | No Hit |
| GGGTACTTCAGGGTCAGGATGCCACGCTTGCTCTGGGCCTCGTCGCCCAC | 39485 | 0.17057903297910265 | No Hit |
| AATAAAAGCGAAAAGAAATGAAAATGTTACACTACATTAATCCTGGAATA | 35984 | 0.1554543731219458 | No Hit |
| ACGCTCGTAGCCCTCGCGCTTCTCCTCGGCCAGTTCGCGGAAGAAGTGGC | 33092 | 0.1429606523830433 | No Hit |
| ATGATGTAGCAGCAGGTGCCAGGGGCTGGCTTGTAGGCGATCAGCAGCTG | 31114 | 0.13441550037005953 | No Hit |
| AGAGCATGACCGATGGATTCCAGTTCGAGTATGGCGGCCAGGGCTCCGAC | 30506 | 0.13178888134887948 | No Hit |
| TCGGGCGCCCCAGACACCAGGCGTCATGGTGGGCATGGGCCAGAAGGACT | 30495 | 0.13174136028106206 | No Hit |
| ACACCATTCAGCTCTACCTGGGGGCCAAGTTGTTGGACTCACAGGGAAAG | 30286 | 0.1308384599925314 | No Hit |
| ATCAAGACATGAGGGAGGCAGGGGCTCAGCTGAAGAAGCTGGTGGACACC | 29827 | 0.12885553543542347 | No Hit |
| AGCGGTTCCGGTGTCCGGAGGCGCTGTTCCAGCCTTCCTTCCTGGGTATG | 29358 | 0.12682940990757238 | No Hit |
| ACAAGCAGTTCACAGTAGAACACTGCAGACTGCAGGGCATTTTCAGGCTA | 28945 | 0.12504520981588266 | No Hit |
| TGACTGGTGATGCTGATATTGGAGTGCATCTGAGGCTGTTGTAGAAATAA | 28142 | 0.12157617186521227 | No Hit |
| ACCAATGTCATAGGGTGCAATATCTACAATAGGTAGTCTCACAGCCTTGC | 27867 | 0.12038814516977722 | No Hit |
| TCACACTTCATGATGGAGTTGAAGGTGGTCTCGTGGATGCCGCAAGATTC | 27492 | 0.11876810876691123 | No Hit |
| TCCCATTTGGCCAAACACATCCAGTTTGCTAGGCTGATTCCCCTGTAGCC | 27117 | 0.11714807236404526 | No Hit |
| GGCCTTCTTCTGGCGCAGAGGAAGCAGGCGCATCTGGGTGTGGGCAATGA | 26913 | 0.11626677256088615 | No Hit |
| ACATAGCAATTCAGGAAATTTGACTTTCCATTCTCTGCTGGATGACGTGA | 26523 | 0.11458193470190552 | No Hit |
| ATCAAATCCTGCAGACAAGGGGAGCCCTCAGTCTGCAGGGCTCCATAATG | 26441 | 0.11422768674181218 | No Hit |
| AGCTTAATGATGCTTTCTCTGGGCTTTTGGGGGAGGGTGTCCACCAGCTT | 25624 | 0.11069816743210148 | No Hit |
| GGTTATAATCATTGGCAATGTTGTAATCAACAACCCATGTTATTTCTACA | 24509 | 0.10588125919424661 | No Hit |
| GGCATTGTCAGGGAAGCTGCAGAGTTATTGAACCACTTGGTCACCTTTCC | 24370 | 0.1052807657009176 | No Hit |
| TCTGTCTTCTTCAGTTTCGACTTATCGAATTTCTCGATCTCAGCCATATC | 24308 | 0.10501291968231044 | No Hit |
| AAAAGTGATGGACTGCCCATTGCTGGAGAAGACCTTCTCTCCTACTGTCA | 24125 | 0.10422234191771182 | No Hit |
| TACATCAAAGATTACATGAAATCAATCAAAGGGAAACTTGAAGAACAGAG | 23321 | 0.10074898386996714 | No Hit |
| ATTCTTCCTACCCATGAGCATGGAATGTTCTTCCATTTGTTTGTATCCTC | 23244 | 0.10041633639524535 | No Hit |

### Adapter Content

Produced by FastQC (version 0.11.7)
