## Supplementary material for "Curated and harmonised transcriptomics datasets of interstitial lung diseases": SRR13615069_fastqc.html

SRR13615069.fastq.gz FastQC Report 

FastQC Report

Mon 5 Feb 2024  
SRR13615069.fastq.gz

### Summary

- Basic Statistics
- Per base sequence quality
- Per sequence quality scores
- Per base sequence content
- Per sequence GC content
- Per base N content
- Sequence Length Distribution
- Sequence Duplication Levels
- Overrepresented sequences
- Adapter Content

### Basic Statistics

| Measure | Value |
| --- | --- |
| Filename | SRR13615069.fastq.gz |
| File type | Conventional base calls |
| Encoding | Sanger / Illumina 1.9 |
| Total Sequences | 34327852 |
| Sequences flagged as poor quality | 0 |
| Sequence length | 50 |
| %GC | 51 |

### Per base sequence quality

### Per sequence quality scores

### Per base sequence content

### Per sequence GC content

### Per base N content

### Sequence Length Distribution

### Sequence Duplication Levels

### Overrepresented sequences

| Sequence | Count | Percentage | Possible Source |
| --- | --- | --- | --- |
| ATCGGAAGAGCACACGTCTGAACTCCAGTCACAGCGATAGATCTCGTATG | 94805 | 0.2761751594594383 | TruSeq Adapter, Index 1 (97% over 35bp) |

### Adapter Content

Produced by FastQC (version 0.11.7)
