## Supplementary figures and images for "Curated and harmonised transcriptomics datasets of interstitial lung diseases"

### biopsy_disease_microarray.png

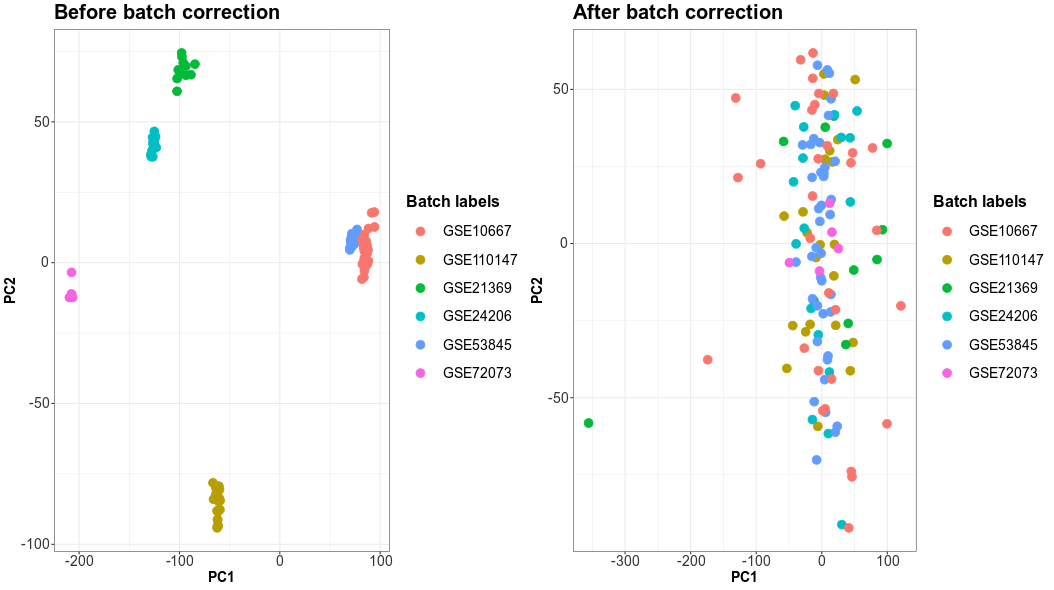

### biopsy_disease_RNAseq.png

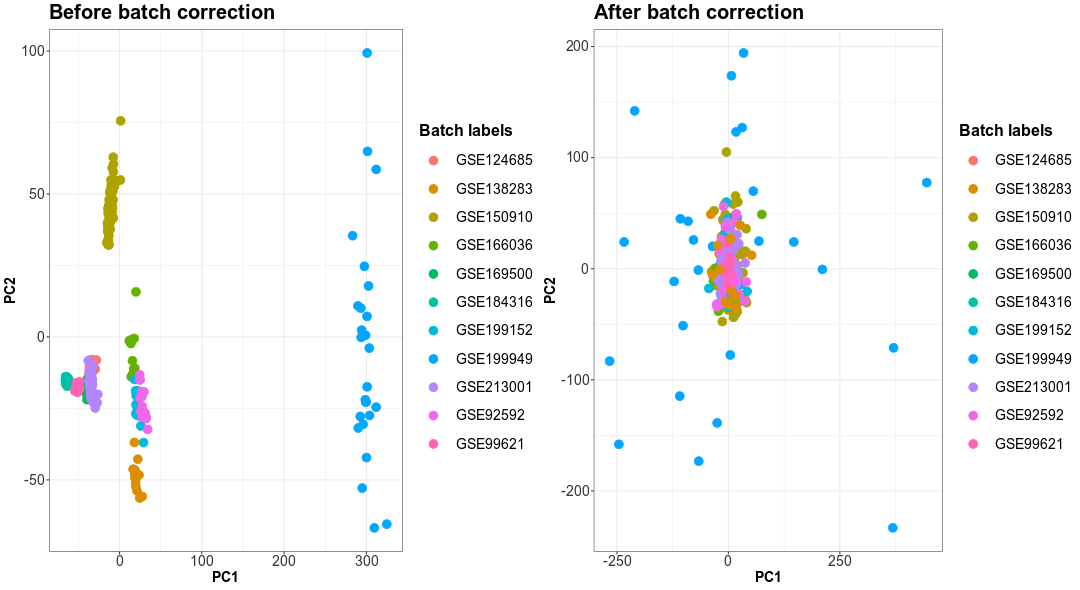

### epithelial_disease_RNAseq.png

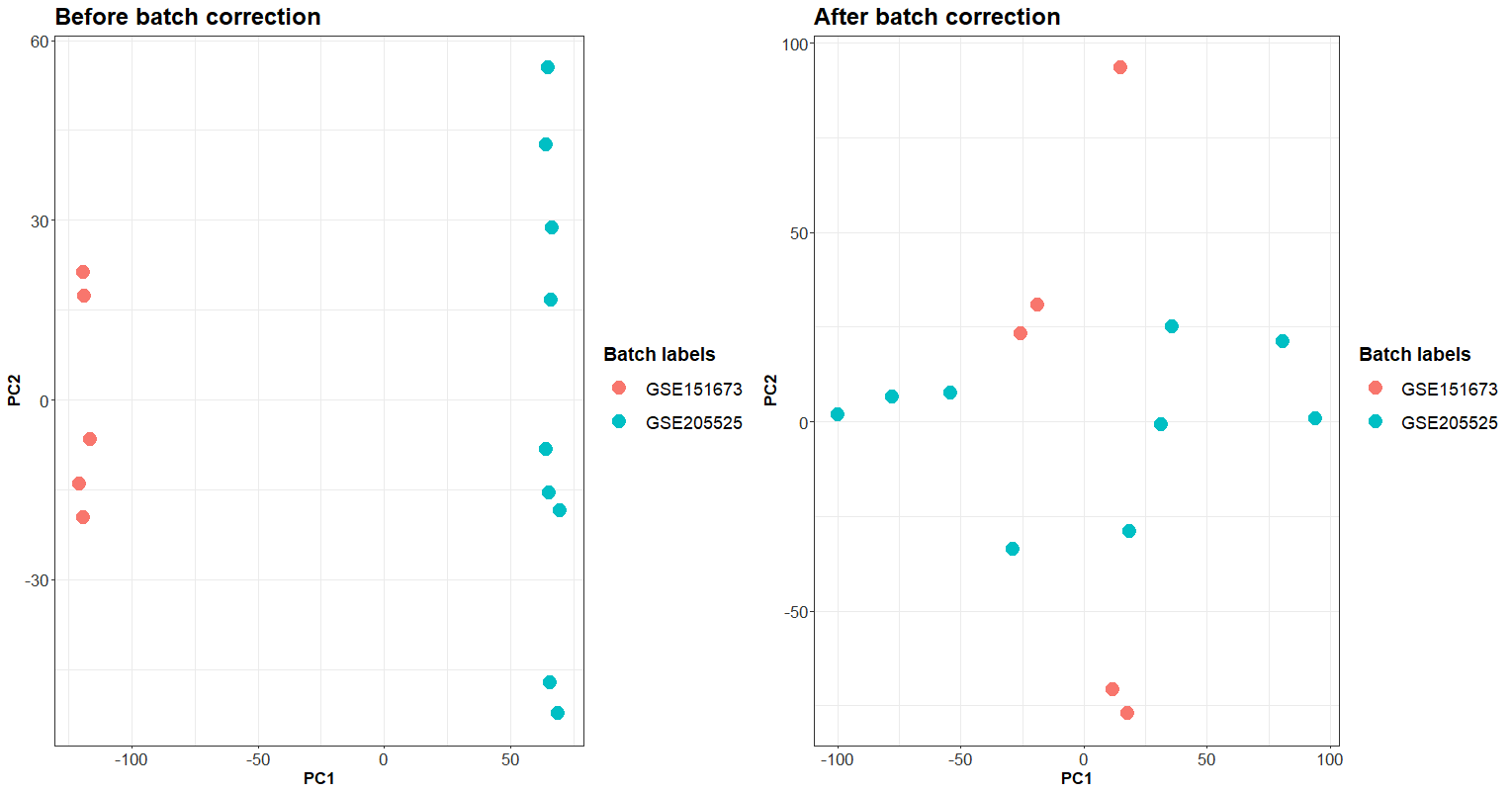

### epithelial_healthy_RNAseq.png

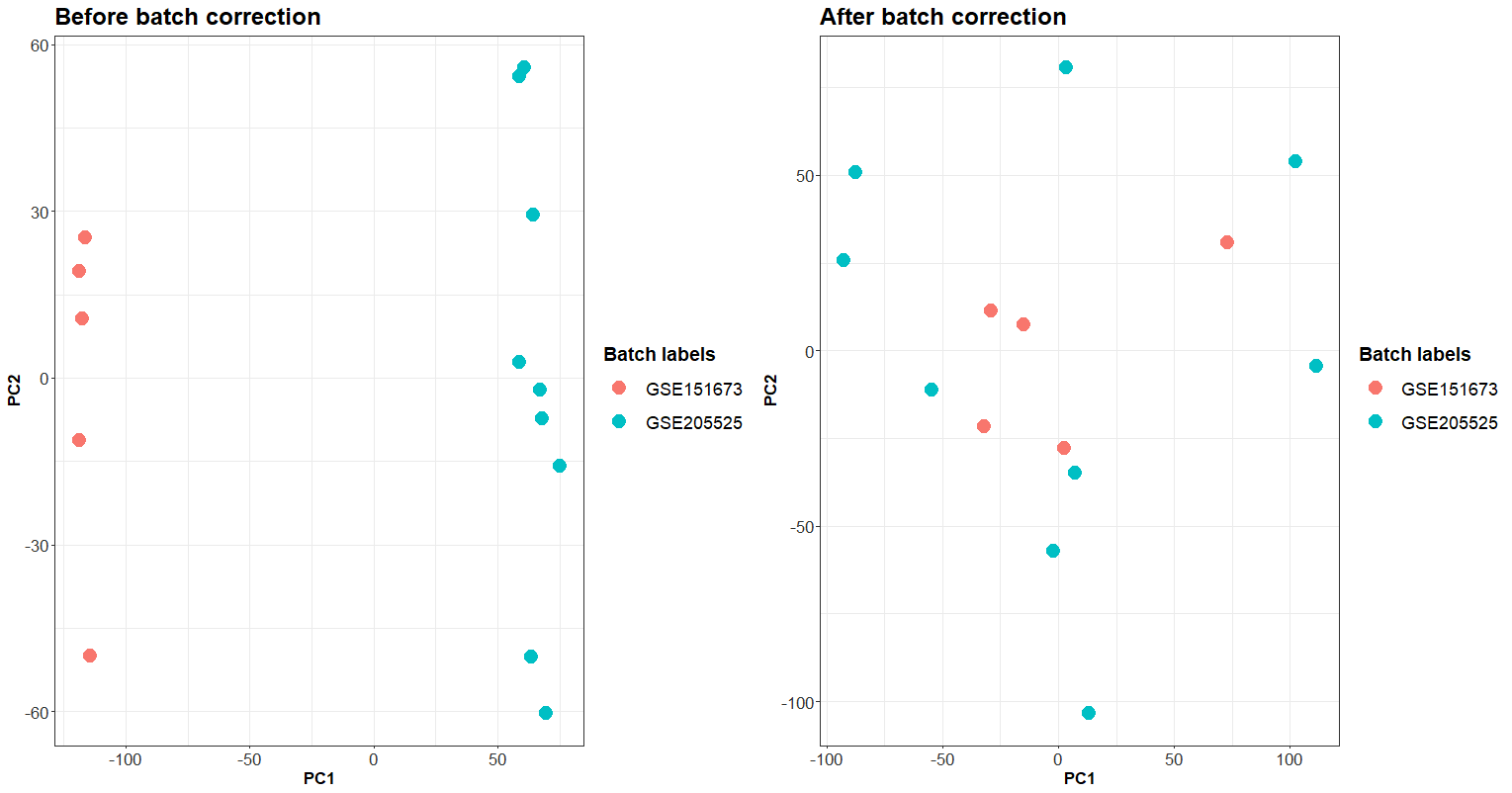

### fibroblast_disease_microarray.png

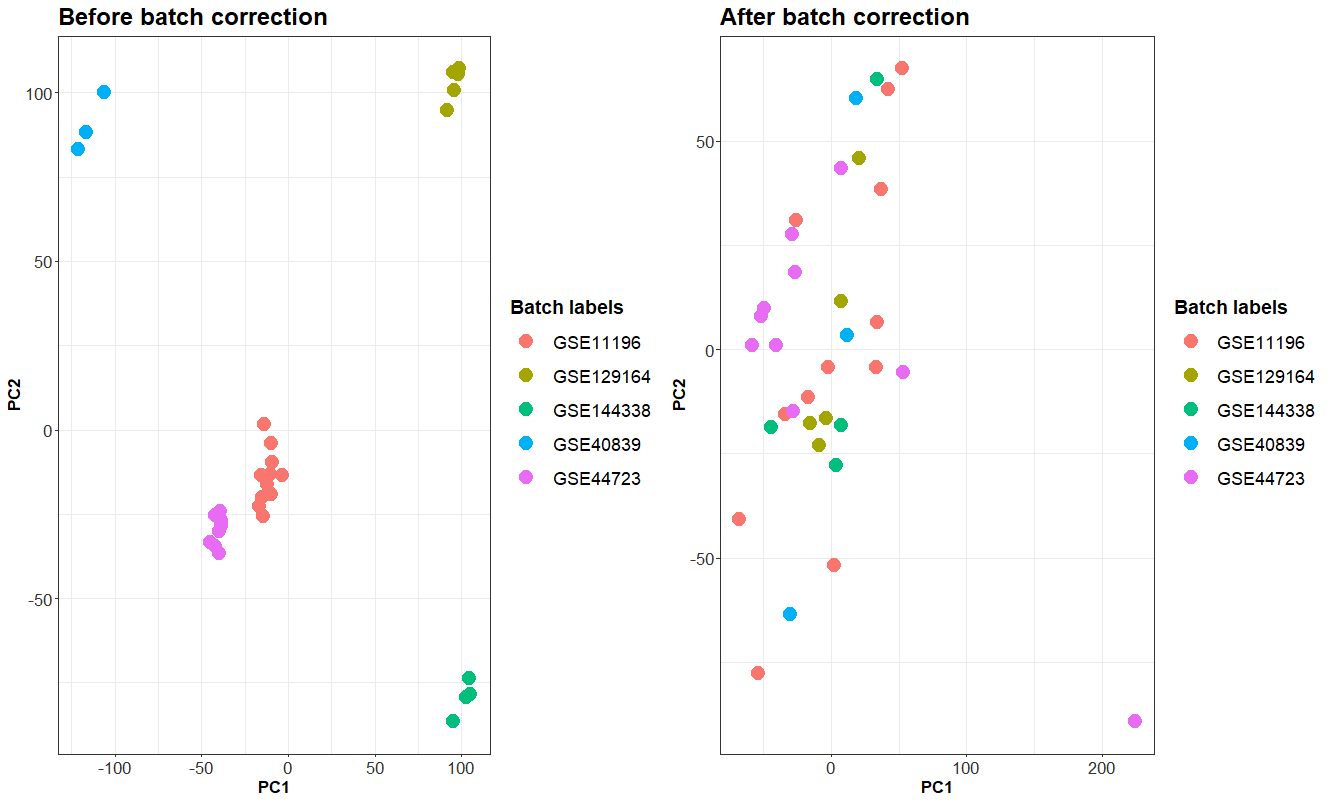

### fibroblast_disease_RNAseq.png

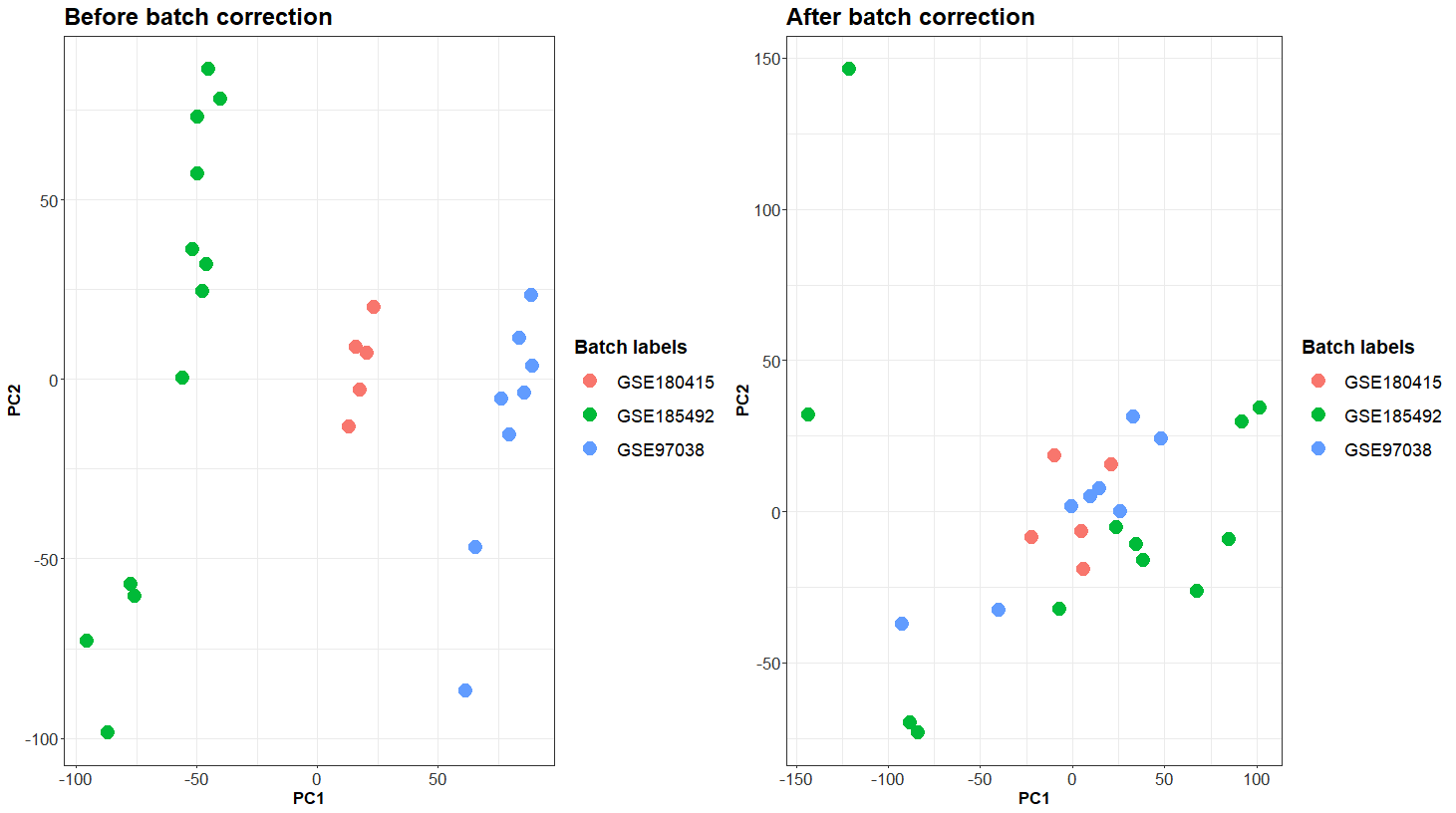

### fibroblast_healthy_microarray.png

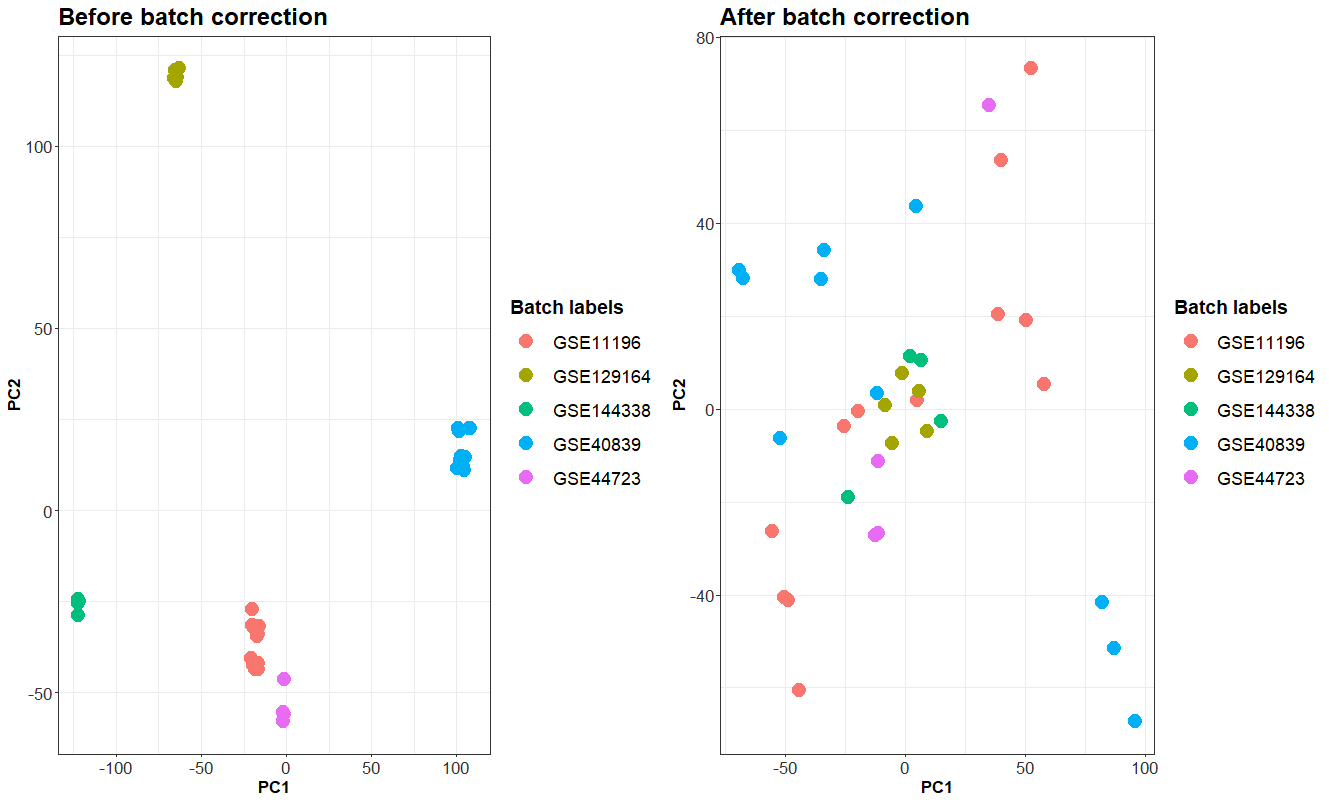

### fibroblast_healthy_RNAseq.png

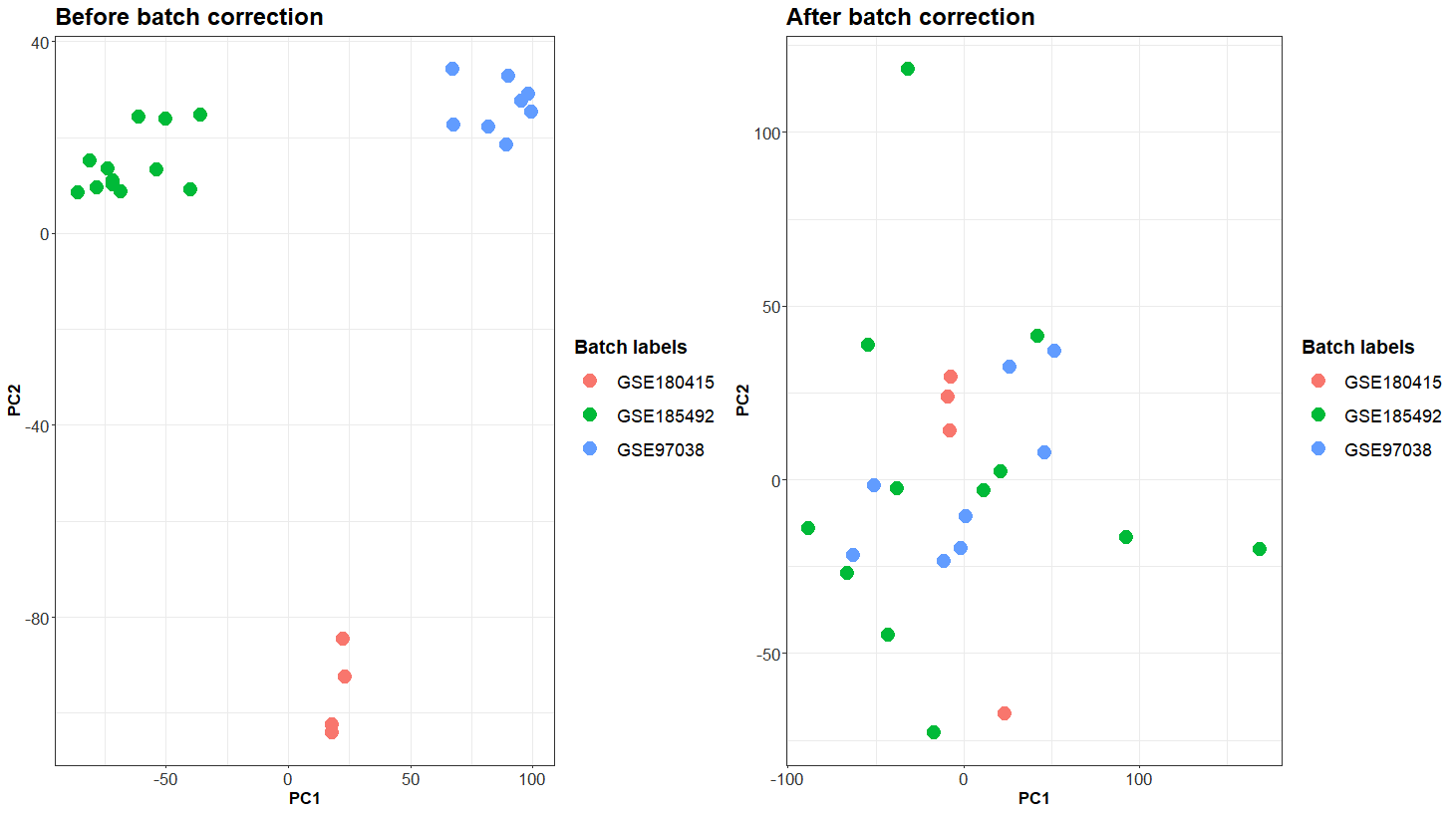

### GSE10667_boxplot_after_normalization.png

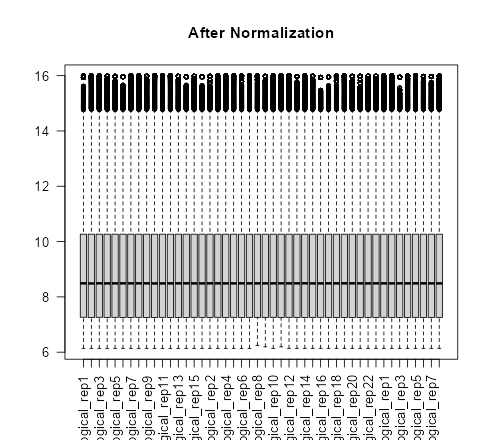

### GSE10667_boxplot_before_normalization.png

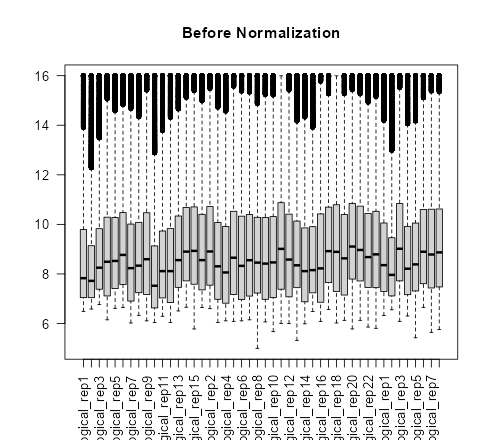

### GSE10667_MDS_after_aggregation_disease.png

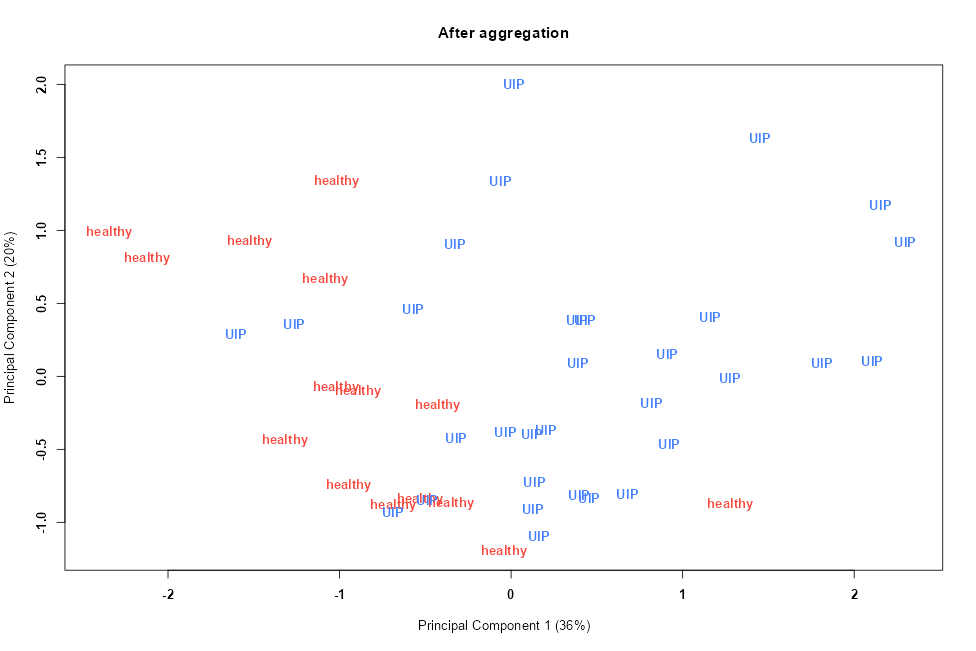

### GSE10667_MDS_after_aggregation_disease_state.png

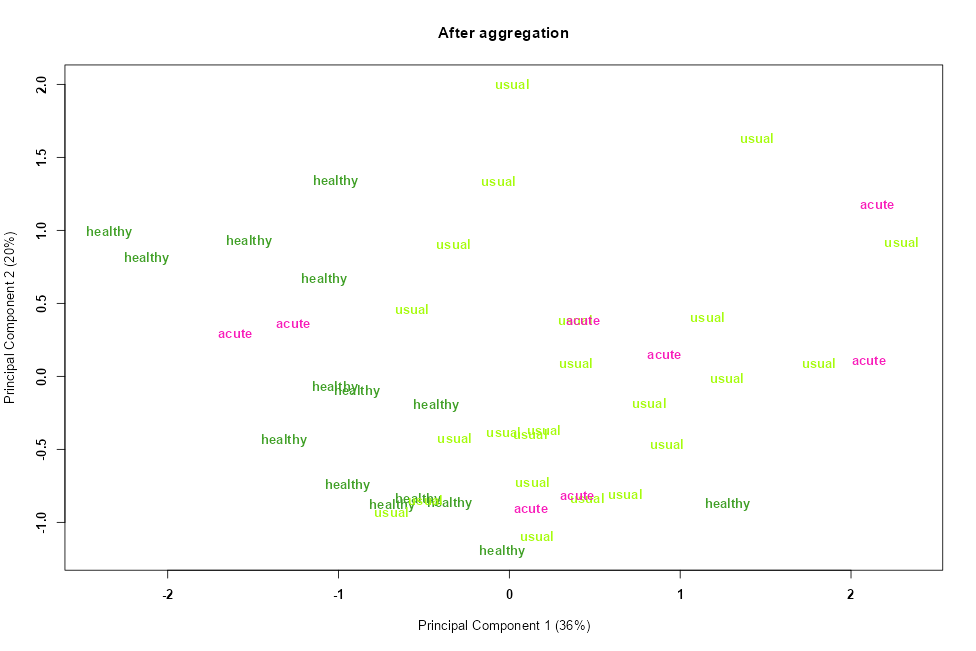

### GSE10667_MDS_before_correction_disease.png

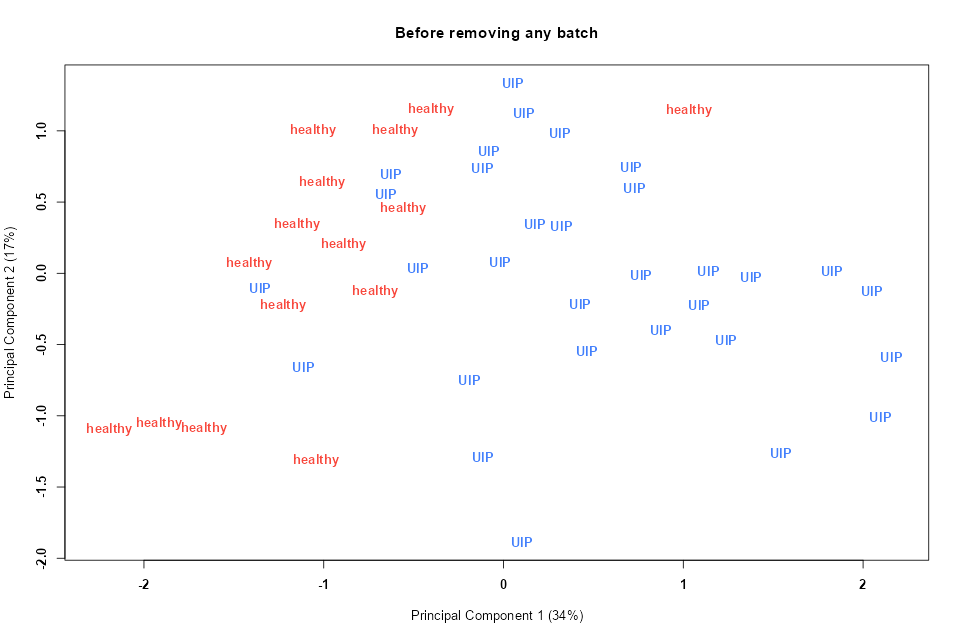

### GSE10667_MDS_before_correction_disease_state.png

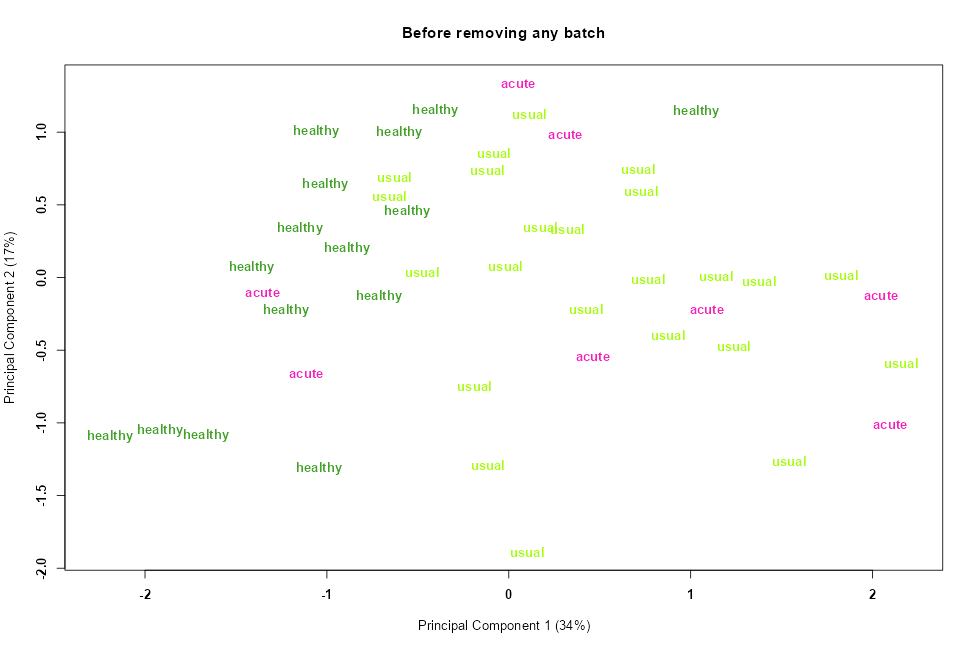

### GSE11196_boxplot_after_normalization_all_samples_RNA.png

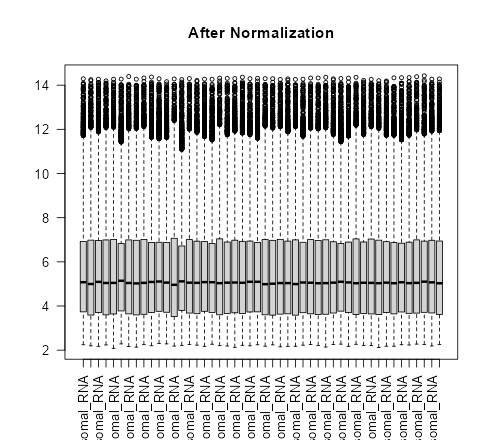

### GSE11196_boxplot_after_normalization_total_RNA.png

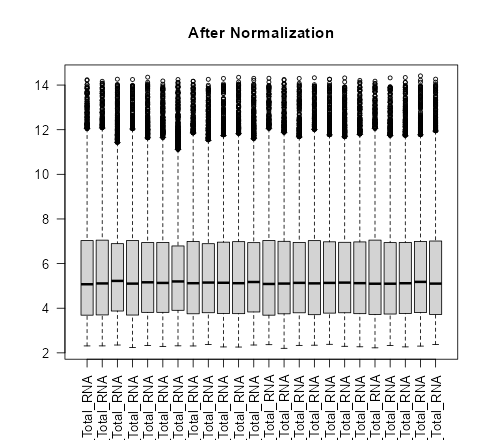

### GSE11196_MDS_before_correction_all_samples.png

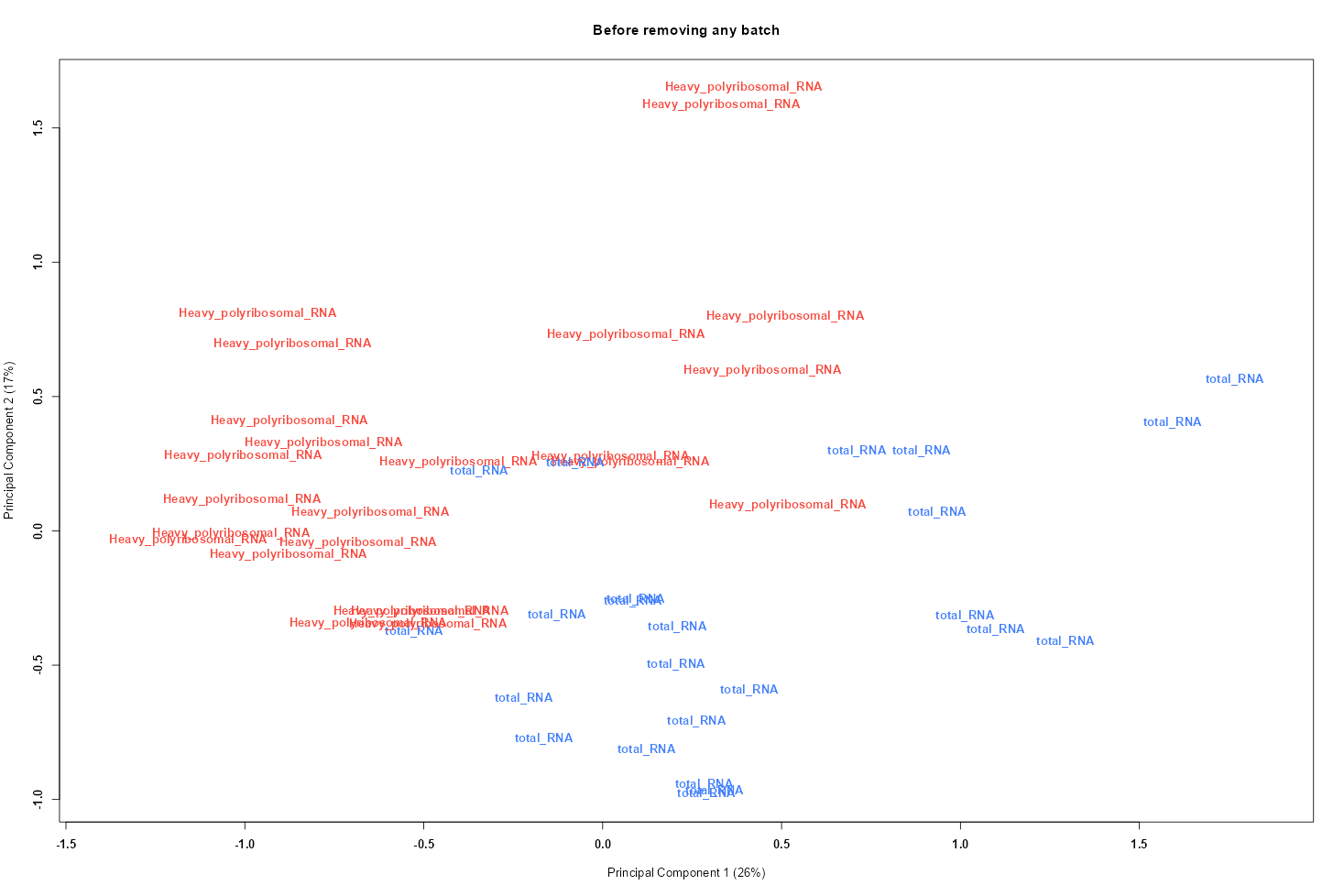

### GSE11196_MDS_before_correction_all_samples_disease.png

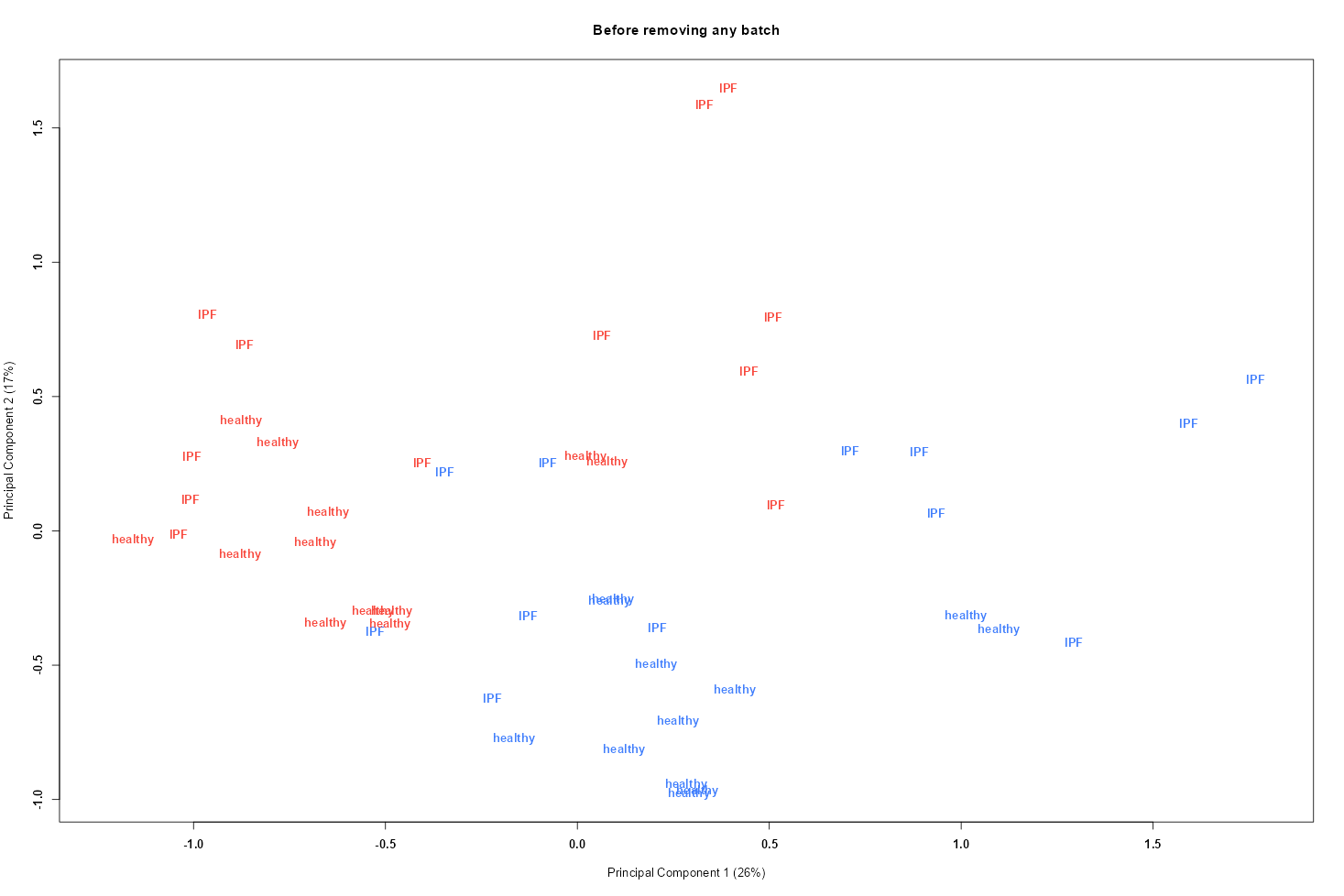

### GSE11196_MDS_before_correction_total_RNA.png

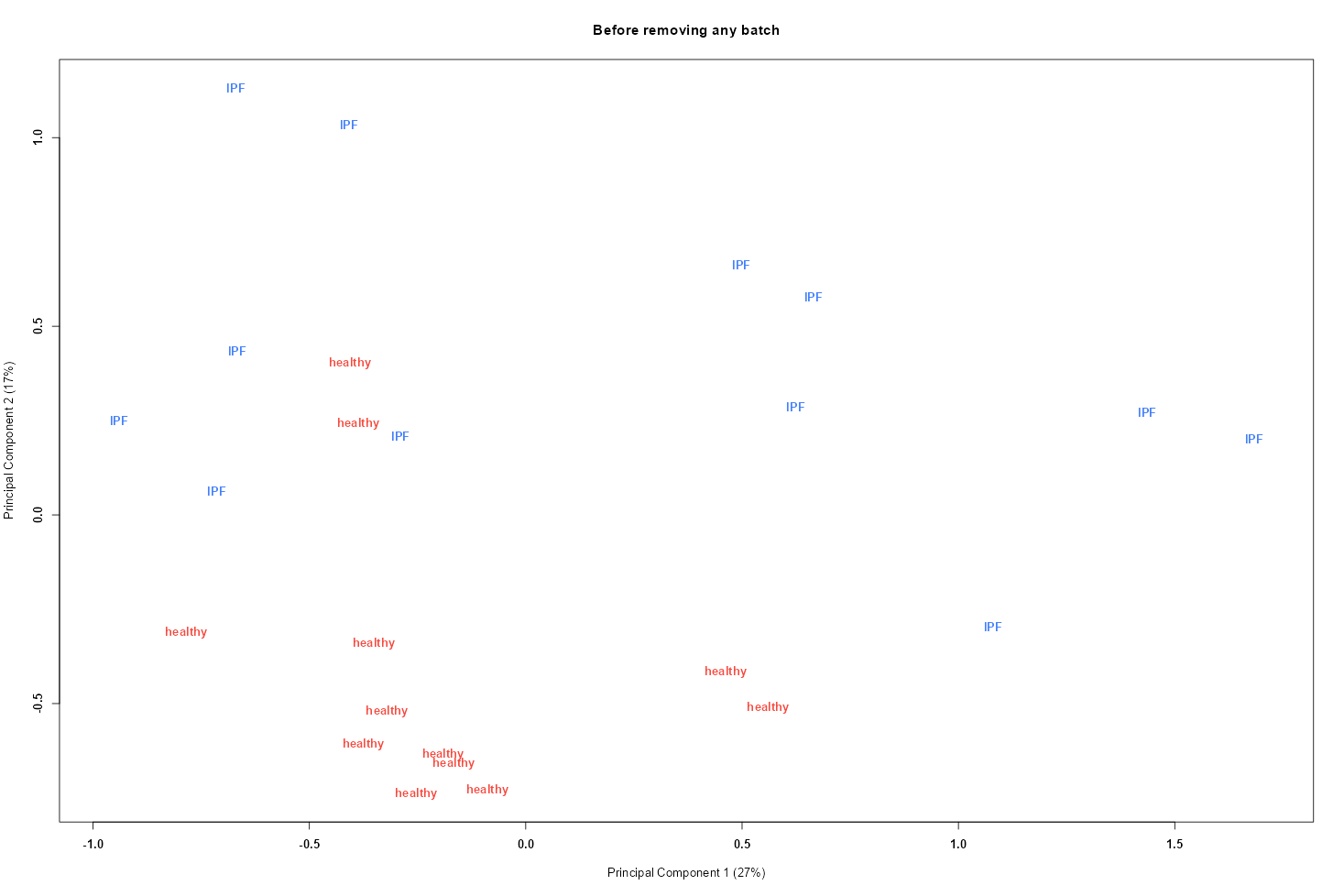

### GSE21369_boxplot_after_normalization.png

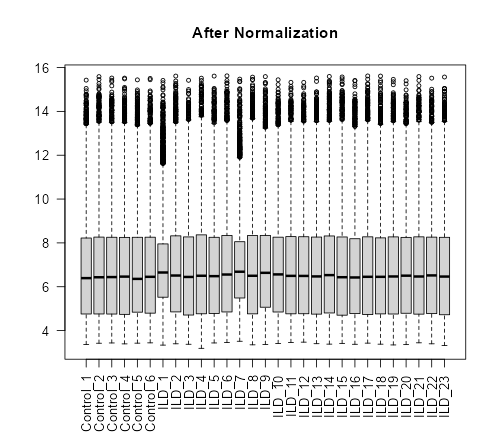

### GSE21369_MDS_before_correction.png

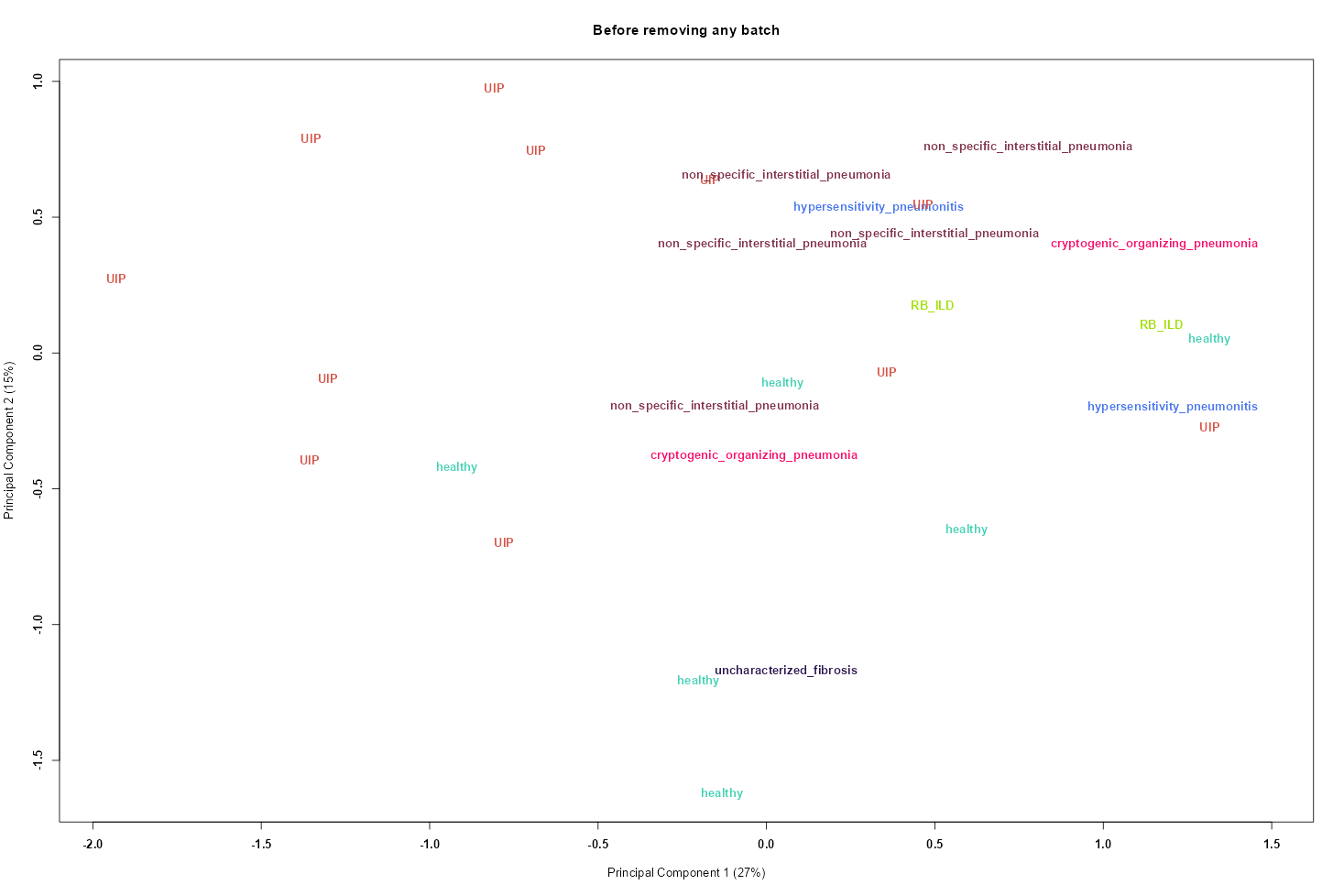

### GSE24206_boxplot_after_normalization.png

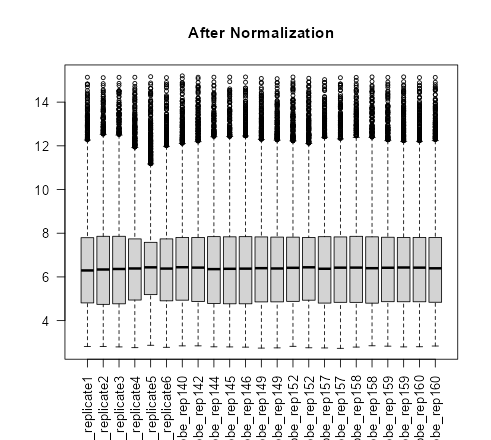

### GSE24206_MDS_before_correction_disease.png

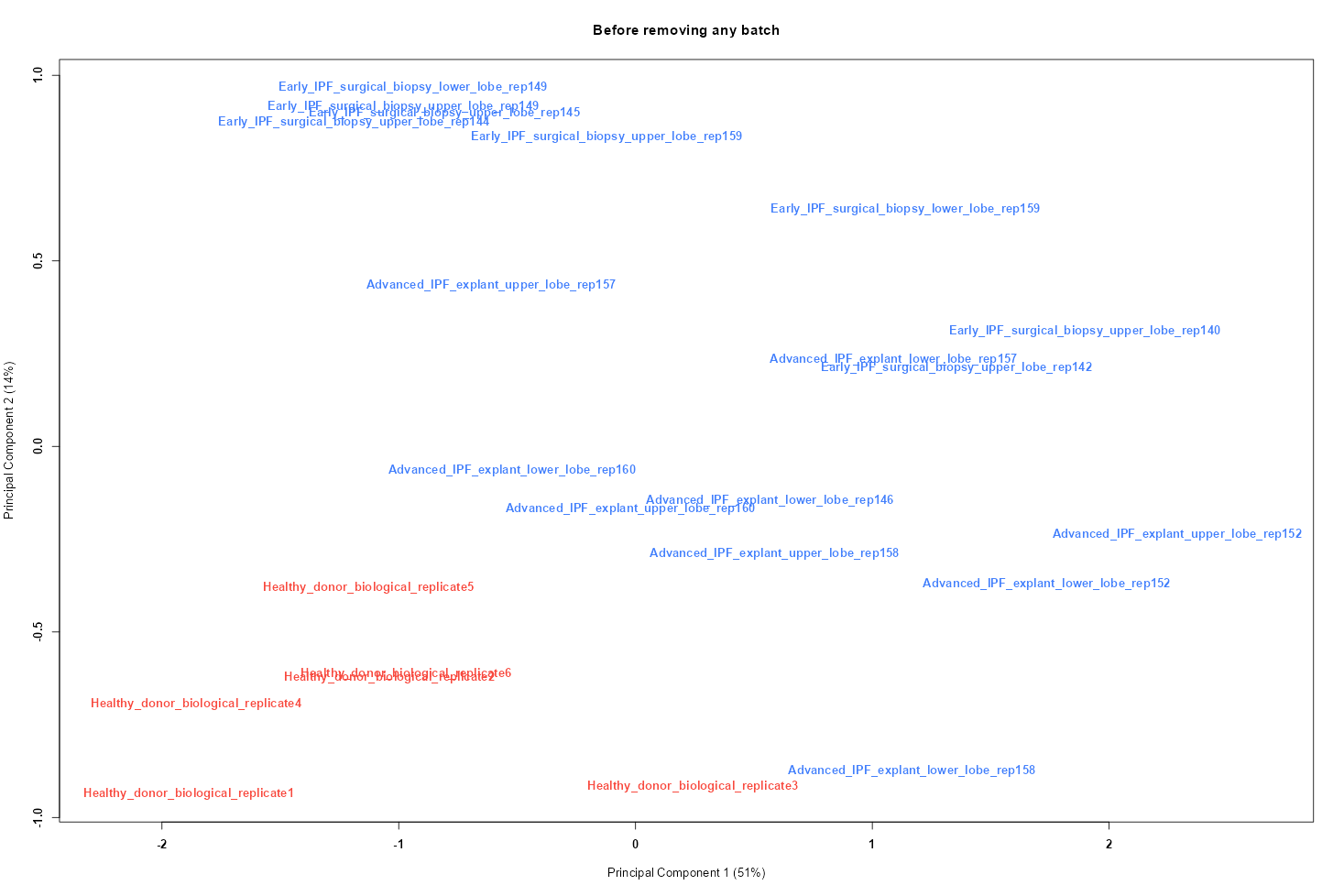

### GSE24206_MDS_before_correction_disease_state.png

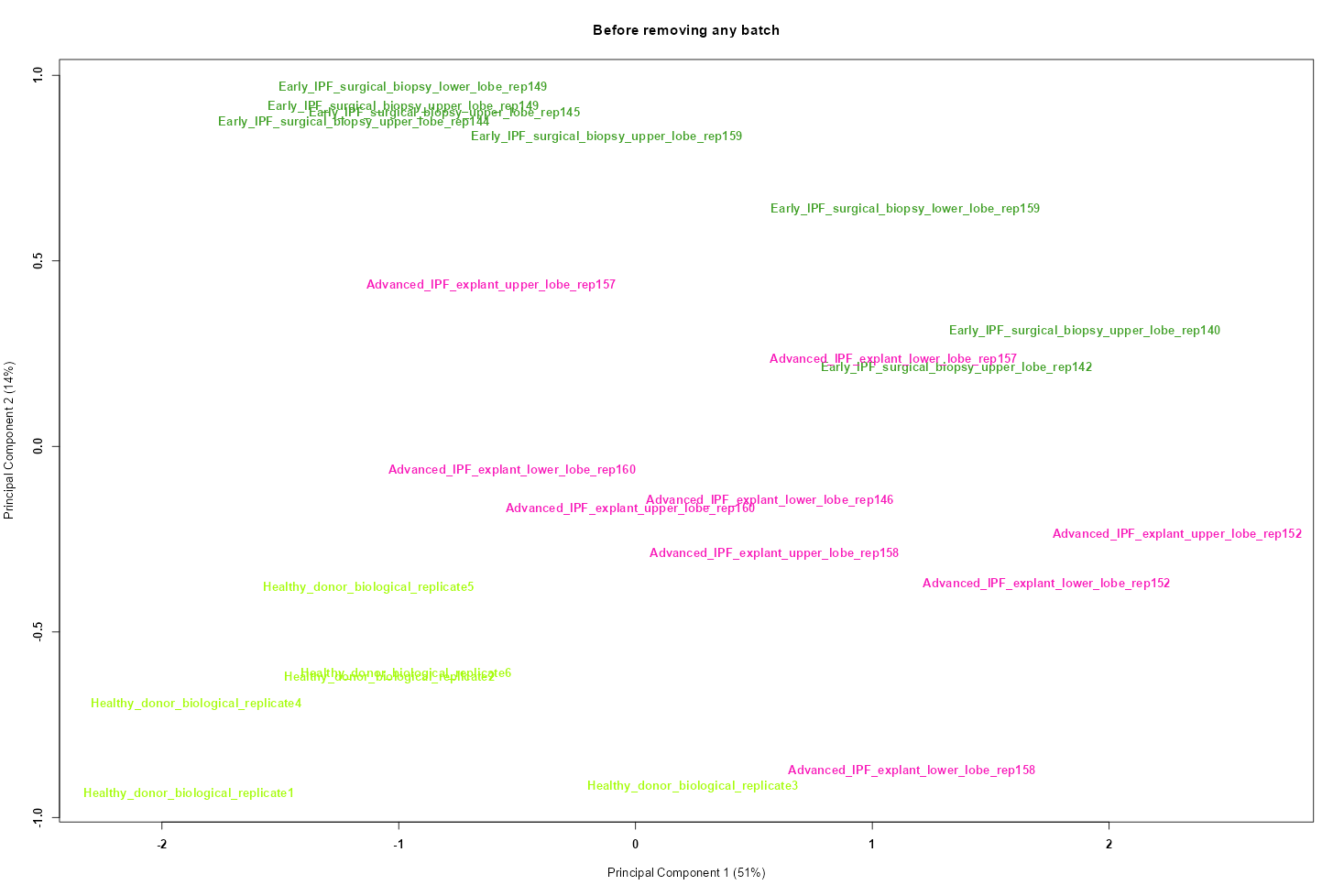

### GSE40839_boxplot_after_normalization.png

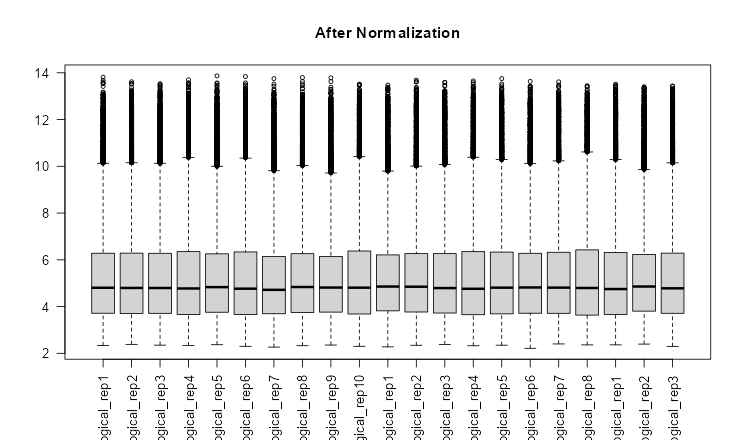

### GSE40839_MDS_before_correction.png

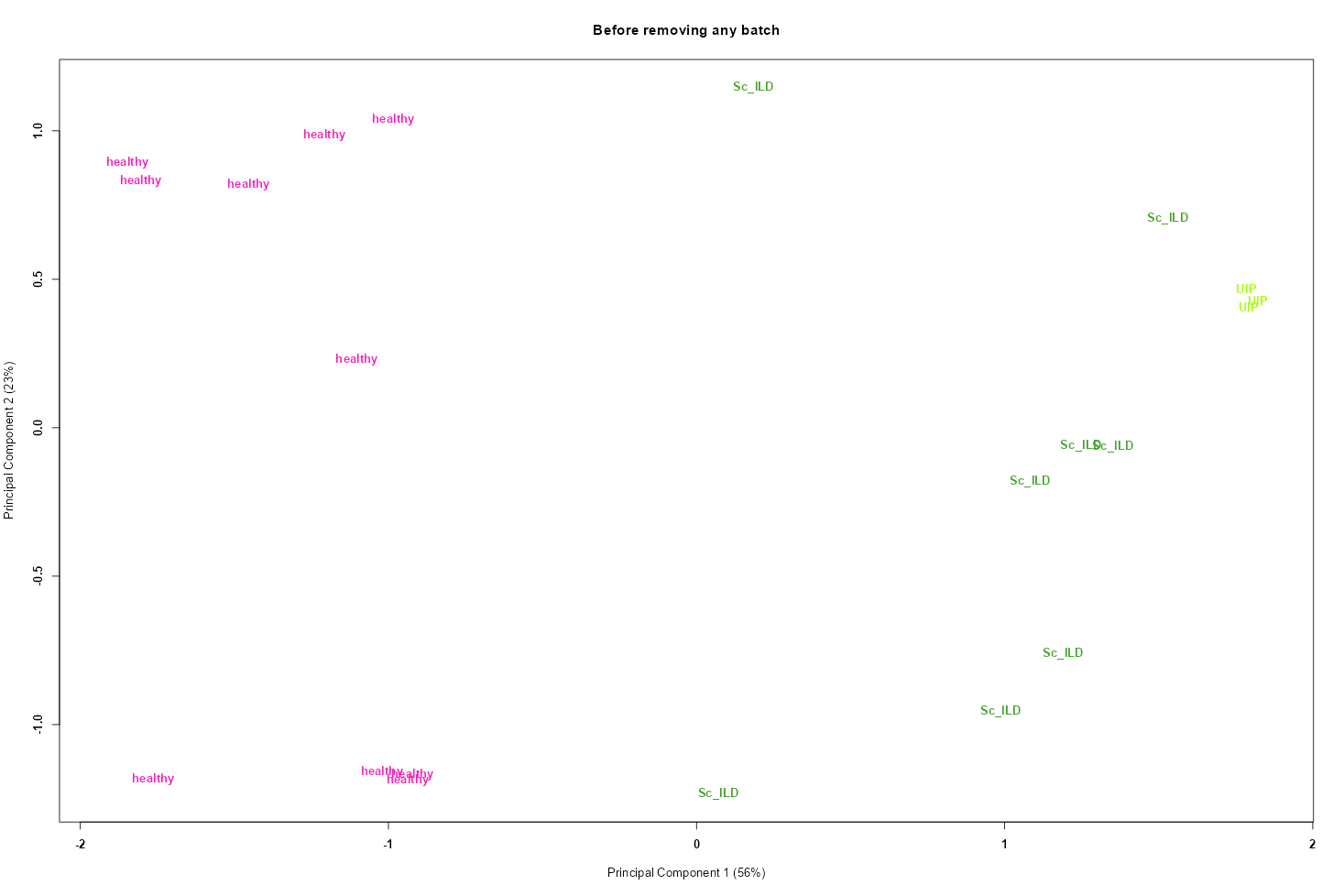

### GSE44723_boxplot_after_normalization.png

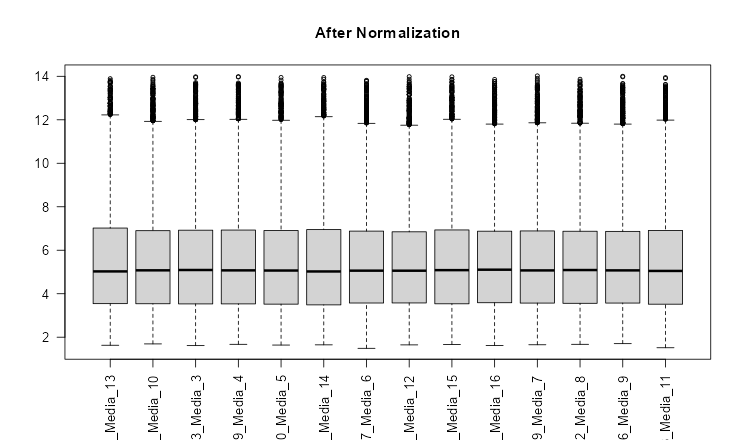

### GSE44723_MDS_before_correction.png

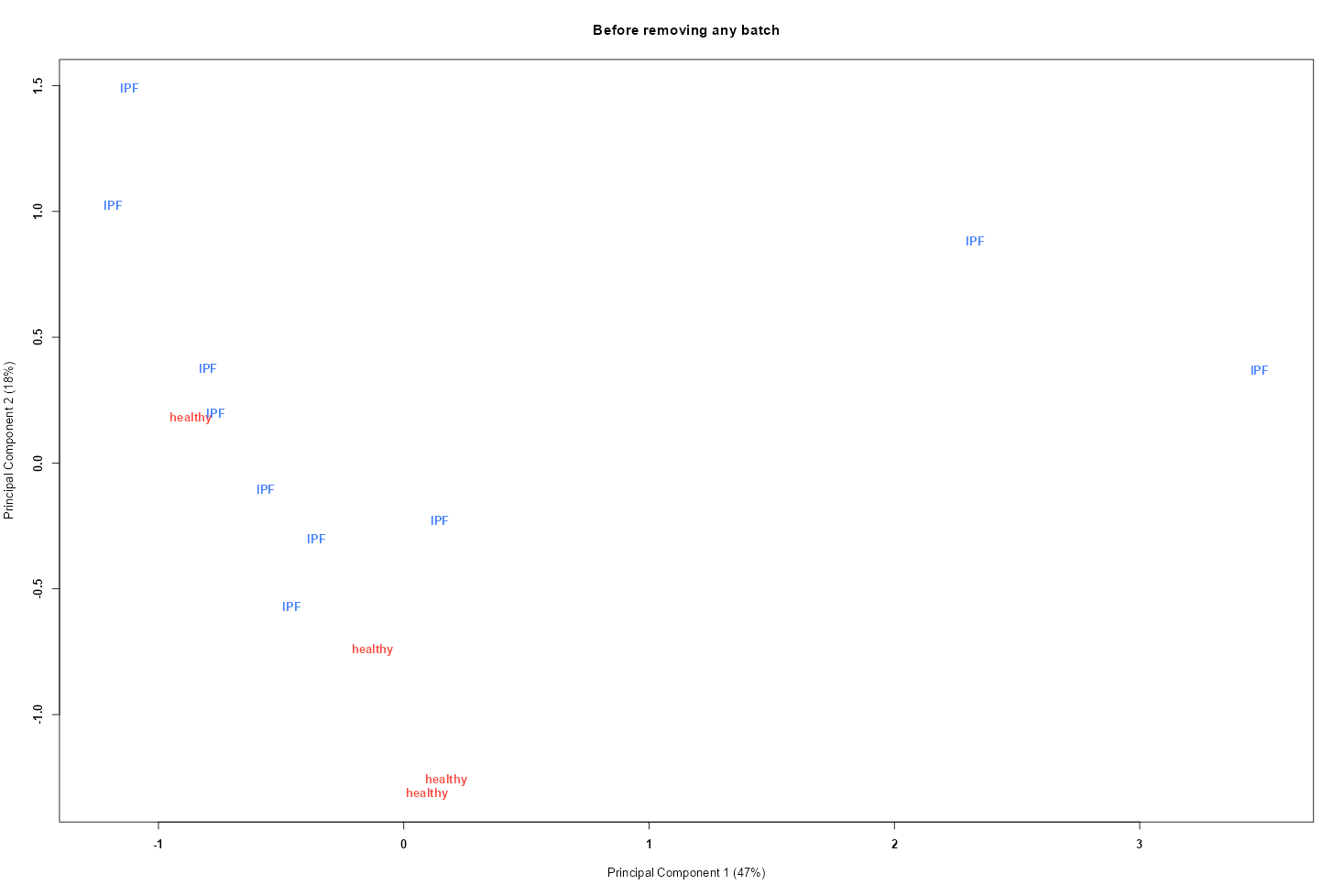

### GSE44723_MDS_before_correction_disease_state.png

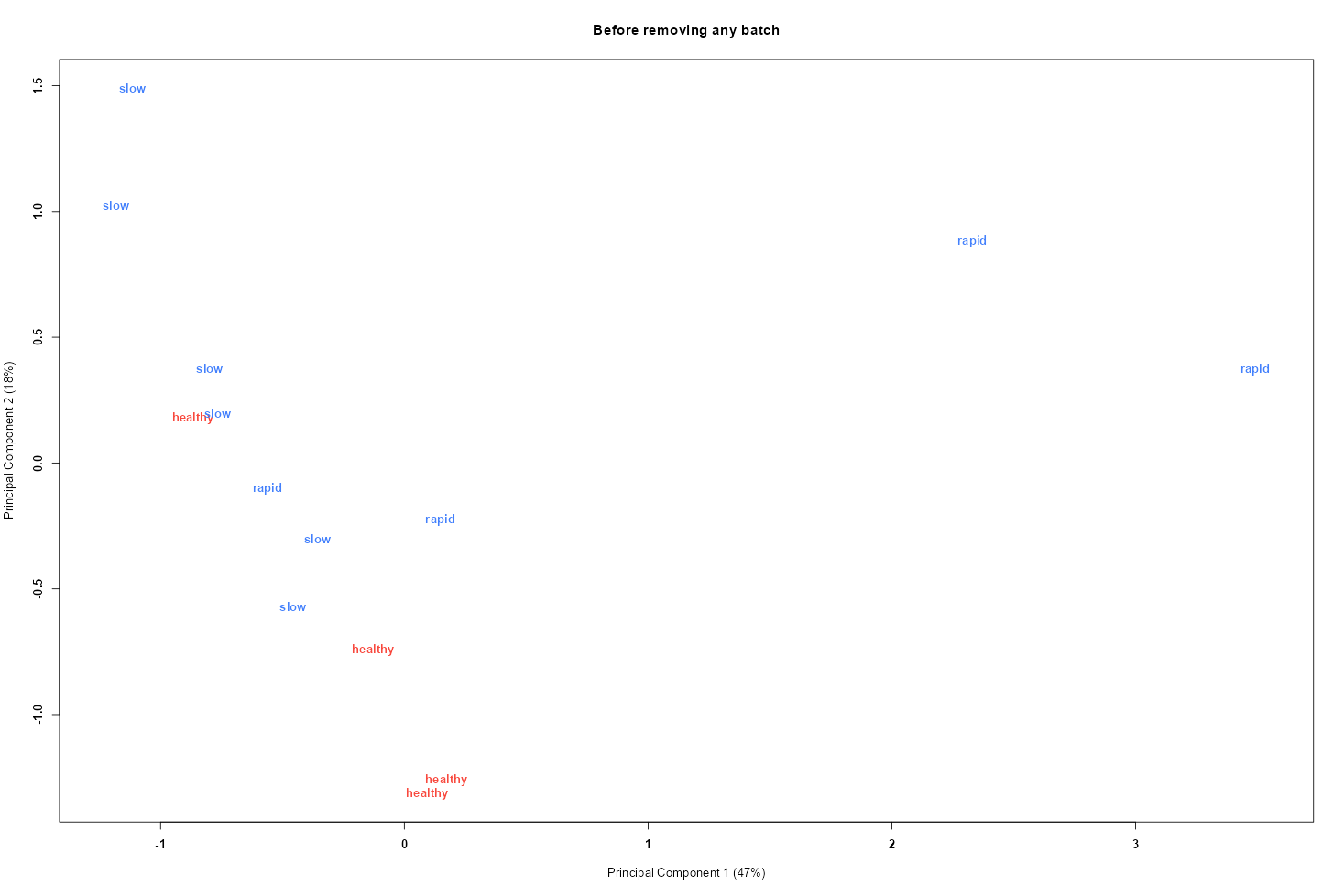

### GSE49072_boxplot_after_normalization.png

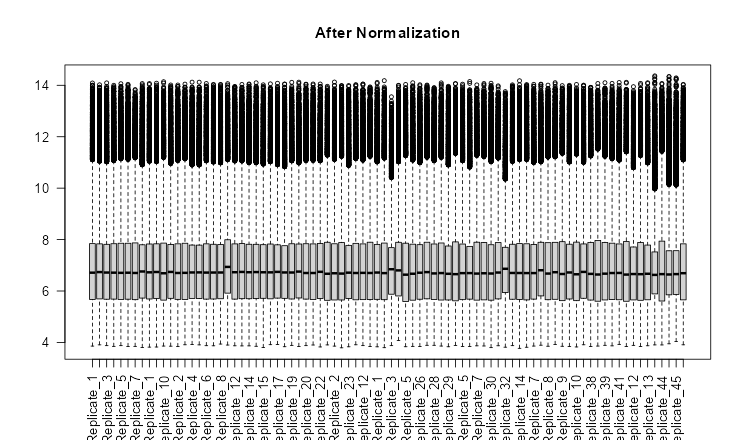

### GSE49072_MDS_before_correction.png

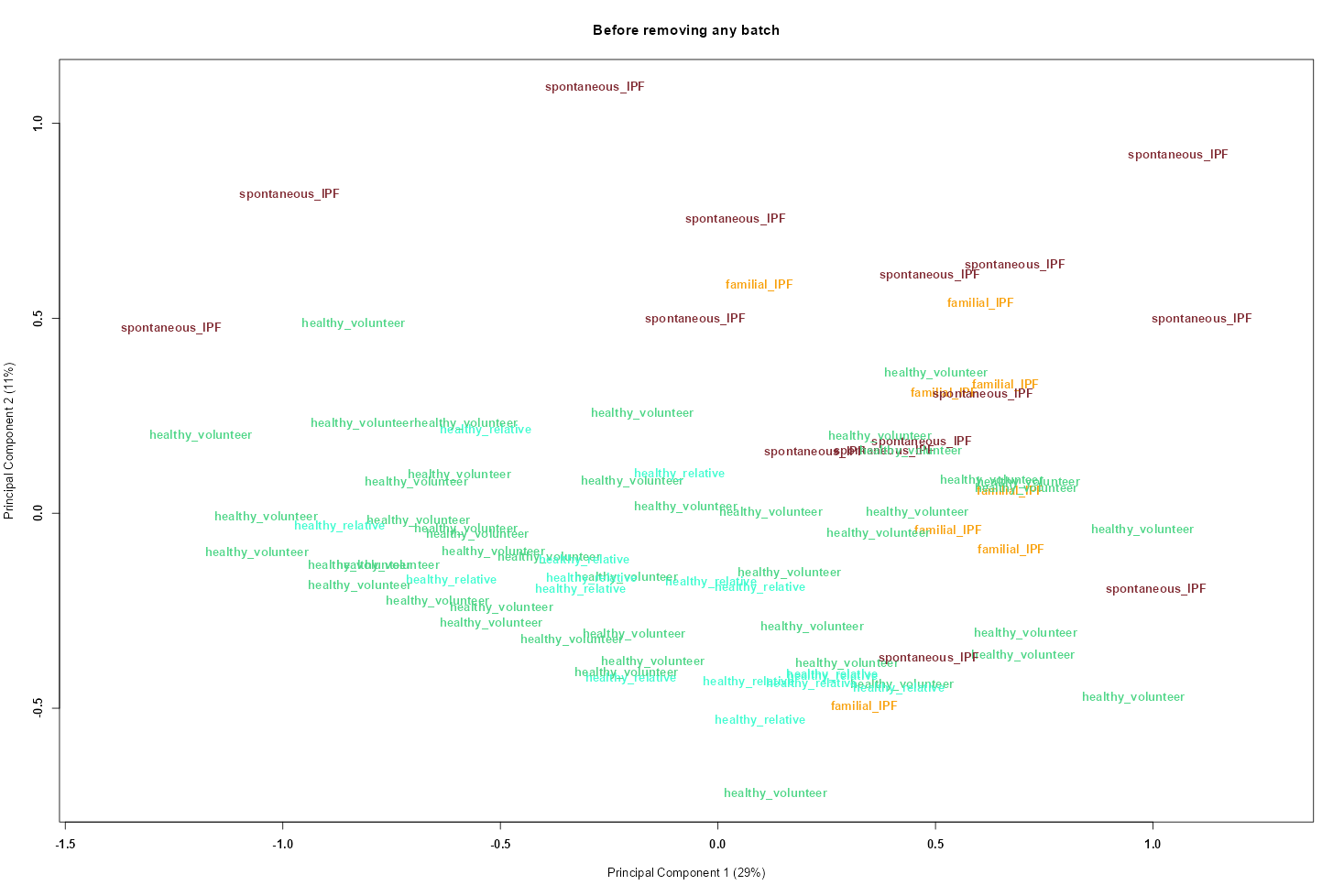

### GSE53845_boxplot_after_normalization.png

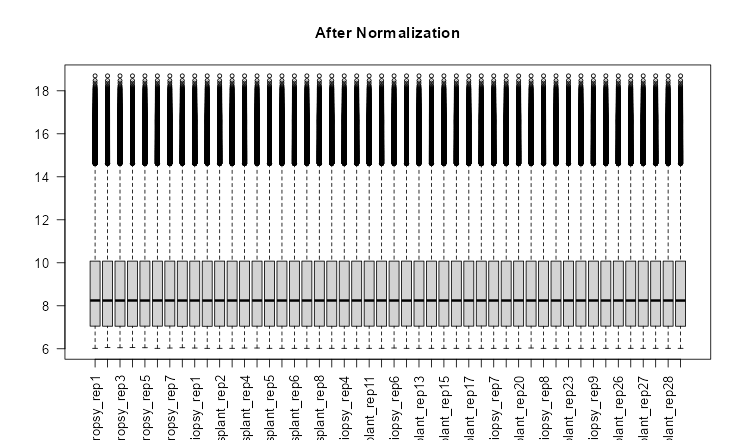

### GSE53845_boxplot_before_normalization.png

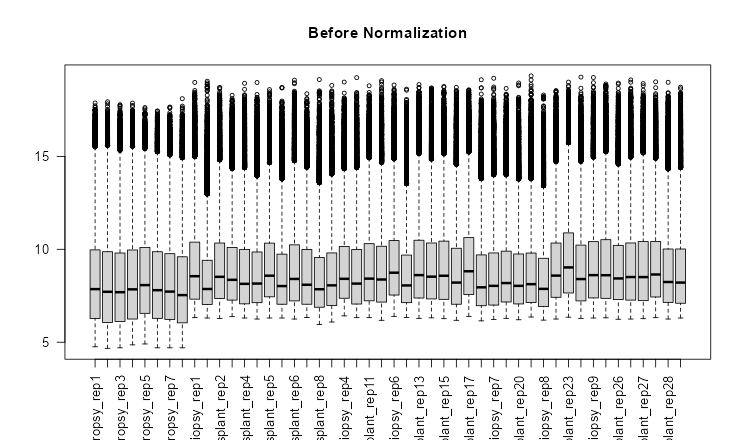

### GSE53845_MDS_after_aggregation.png

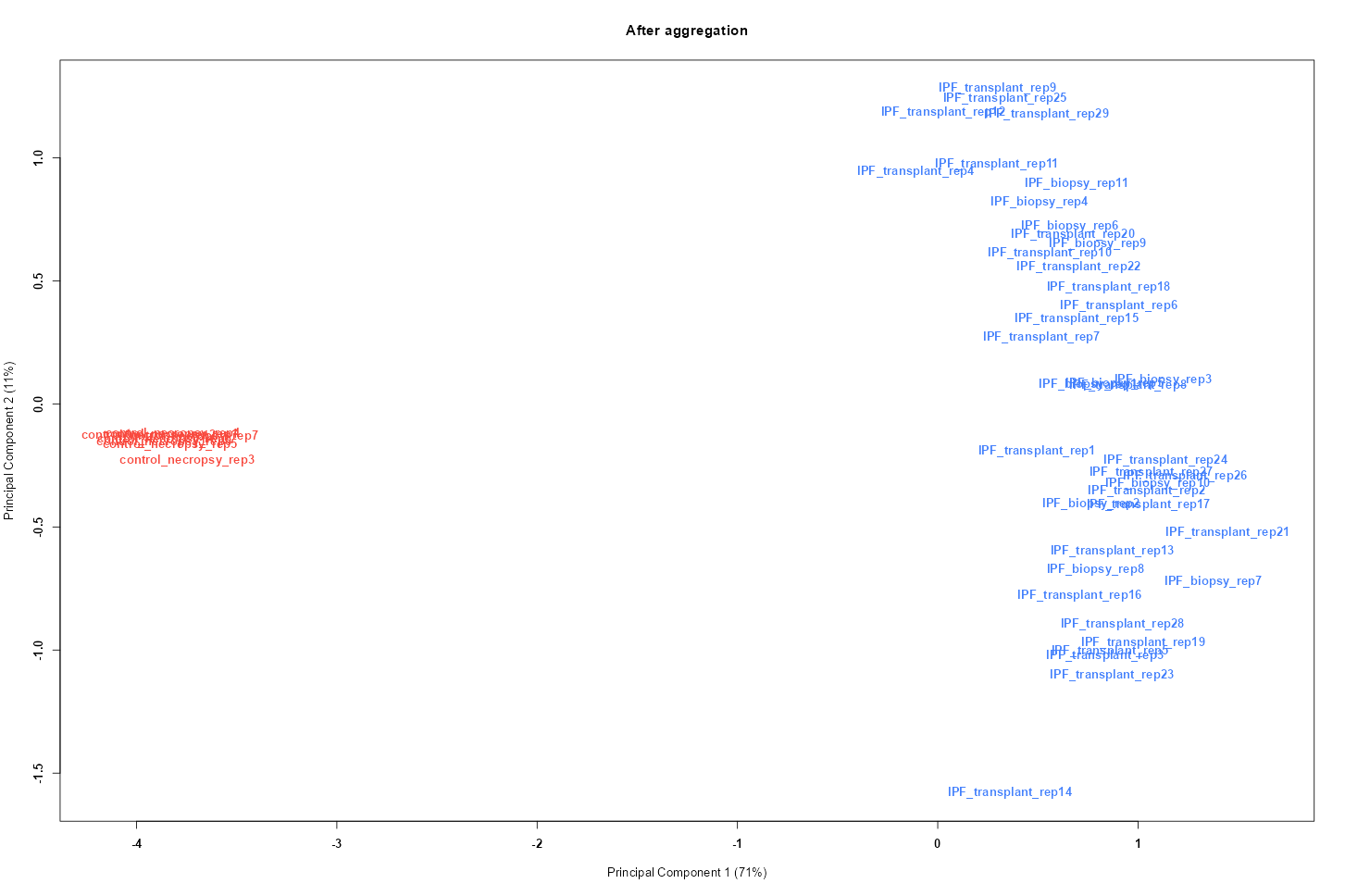

### GSE53845_MDS_before_correction.png

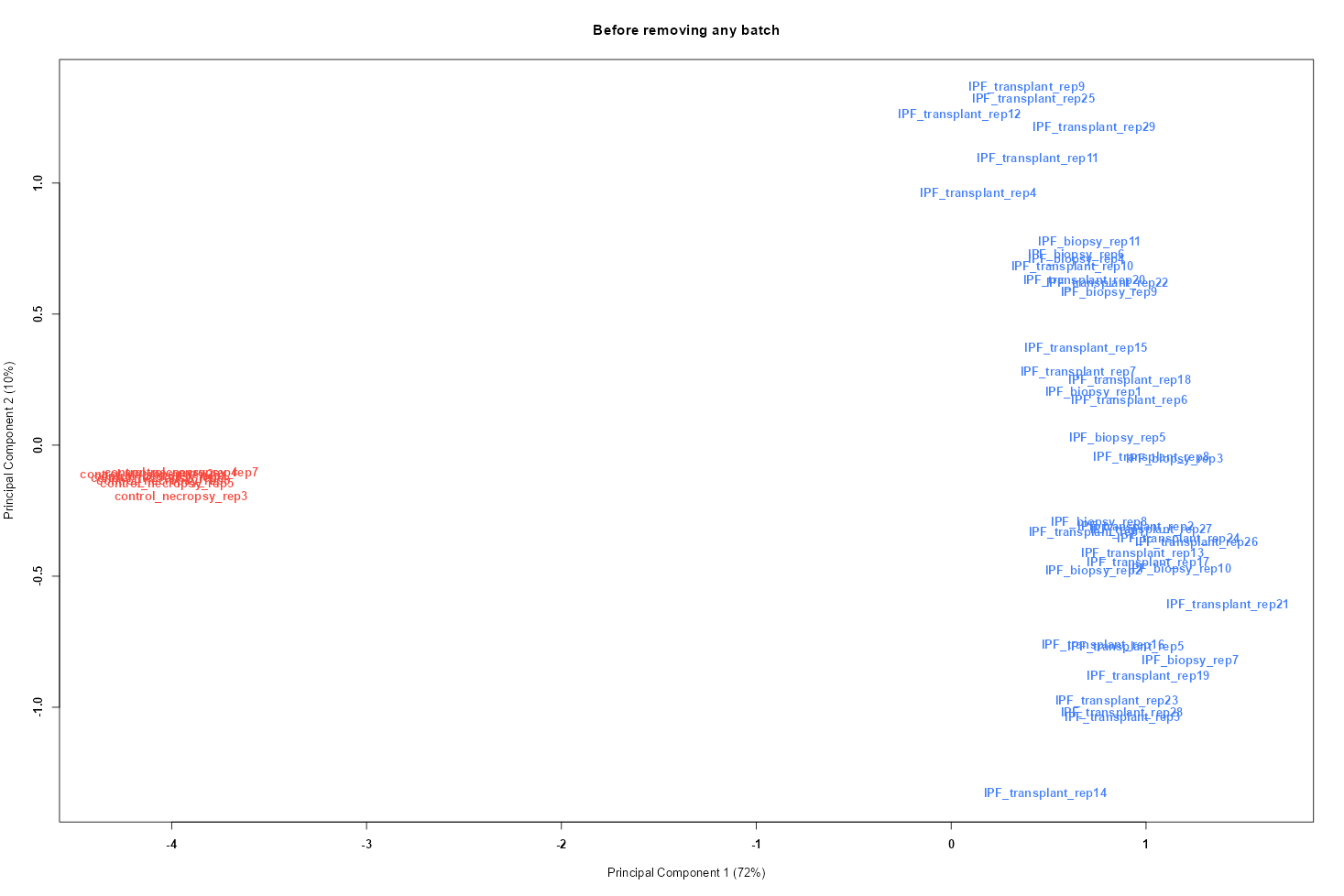

### GSE70866_boxplot_after_normalization_all_samples.png

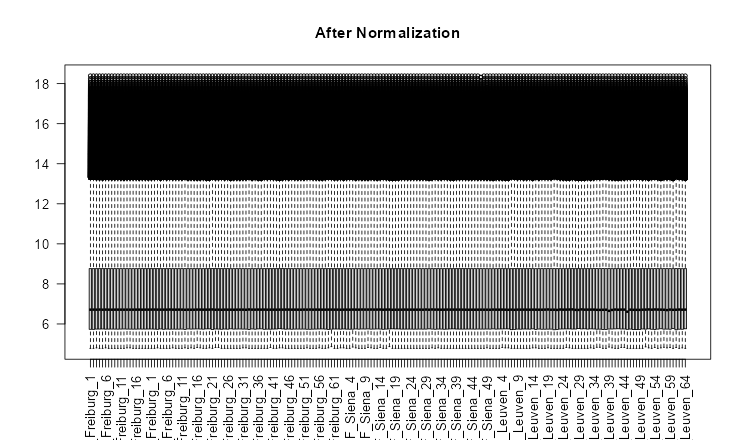

### GSE70866_boxplot_after_normalization_freiburg.png

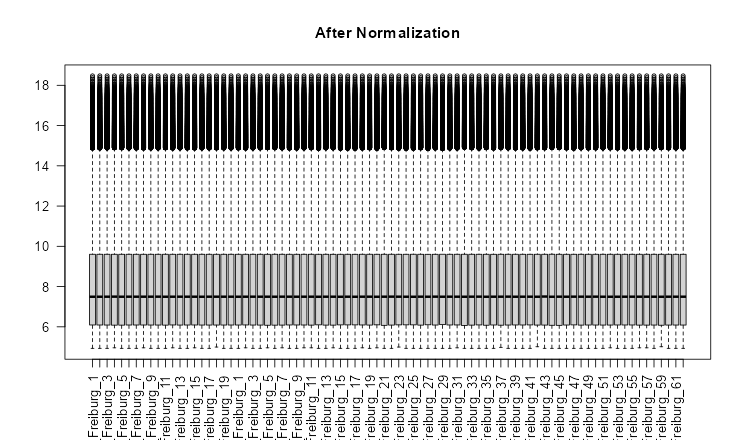

### GSE70866_boxplot_before_normalization_all_samples.png

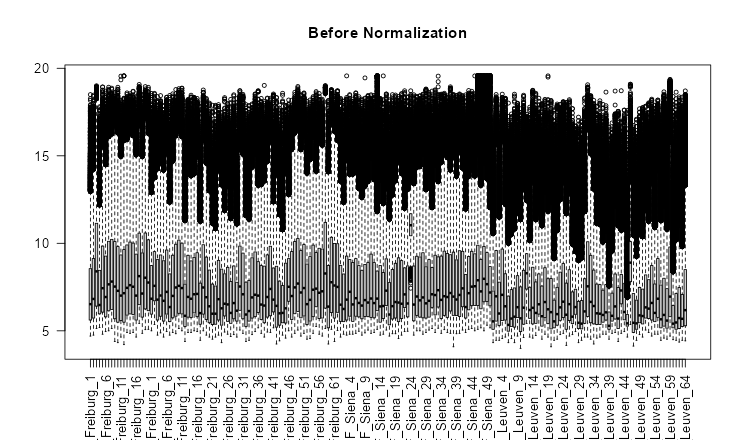

### GSE70866_boxplot_before_normalization_freiburgh.png

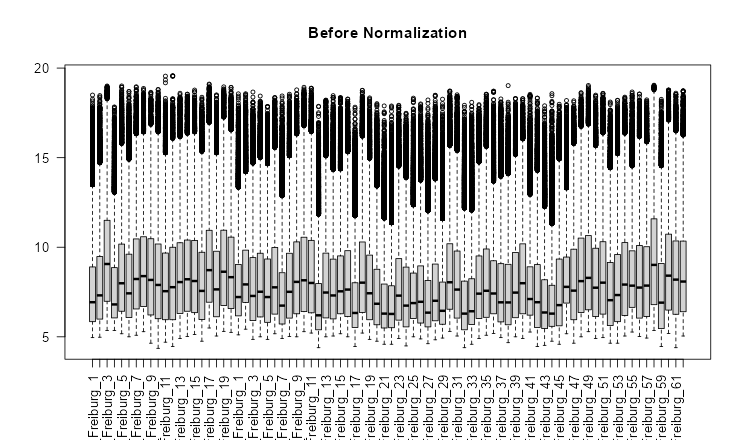

### GSE70866_MDS_after_aggregation_freiburgh.png

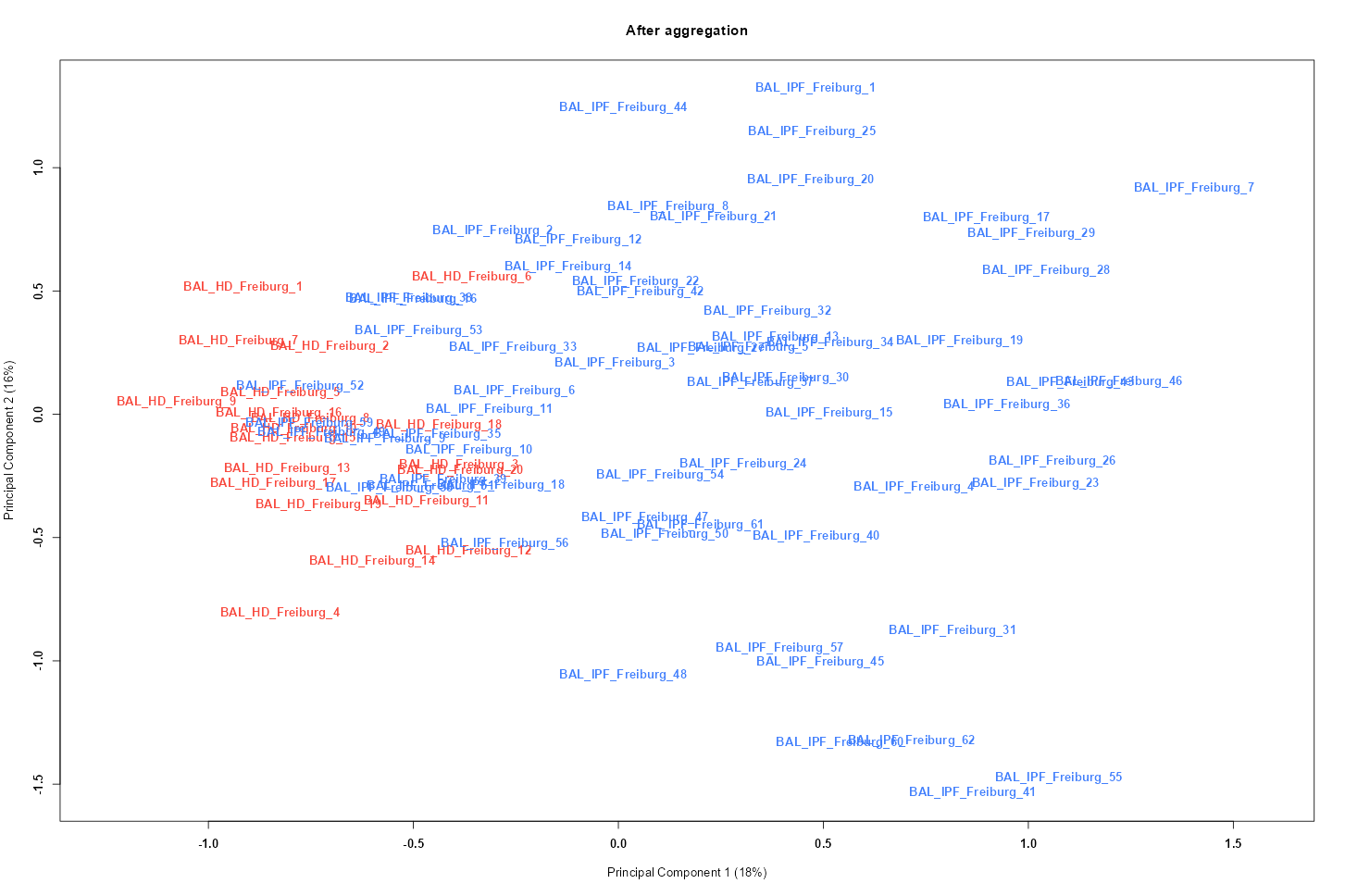

### GSE70866_MDS_before_correction_all_samples.png

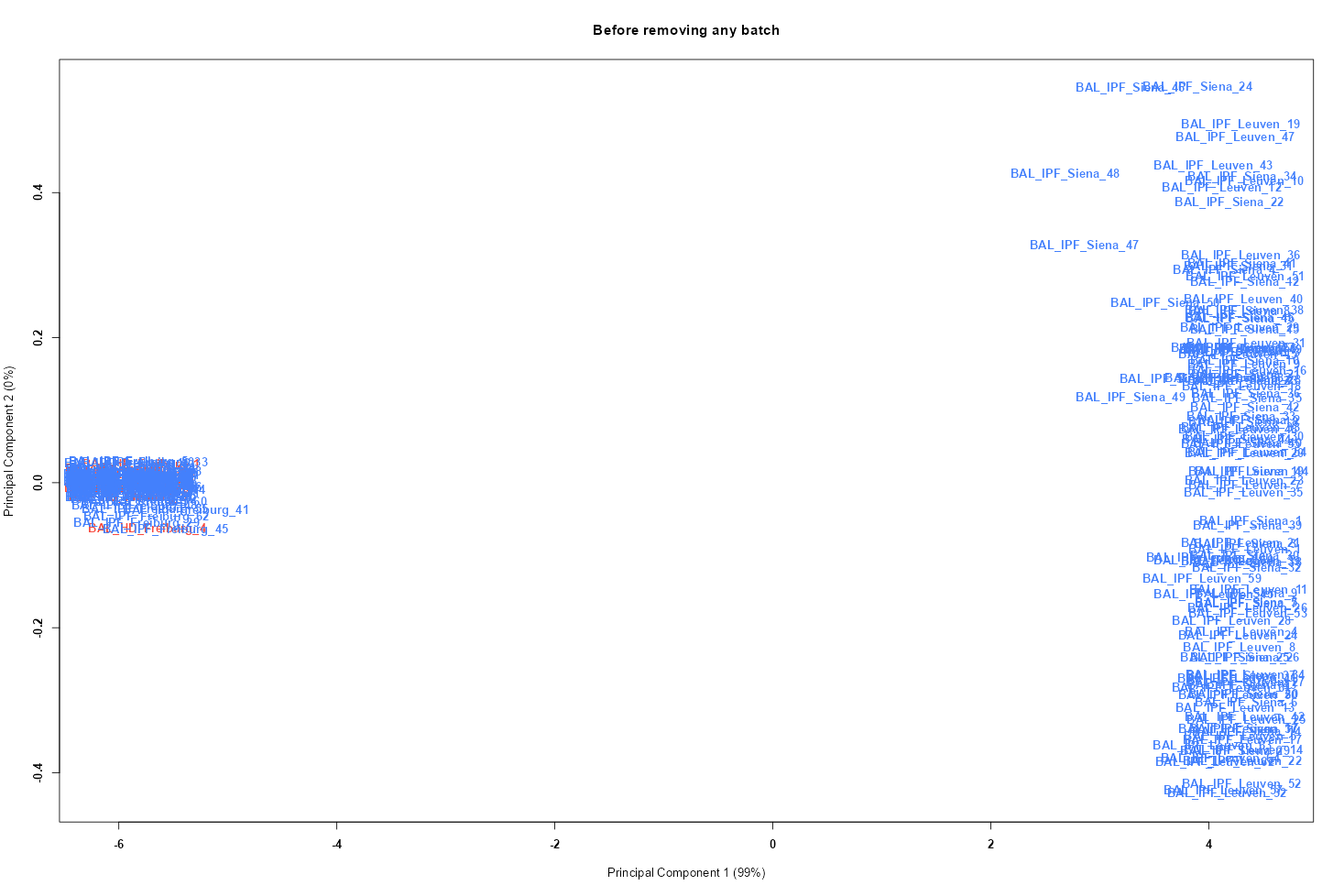

### GSE70866_MDS_before_correction_freiburgh.png

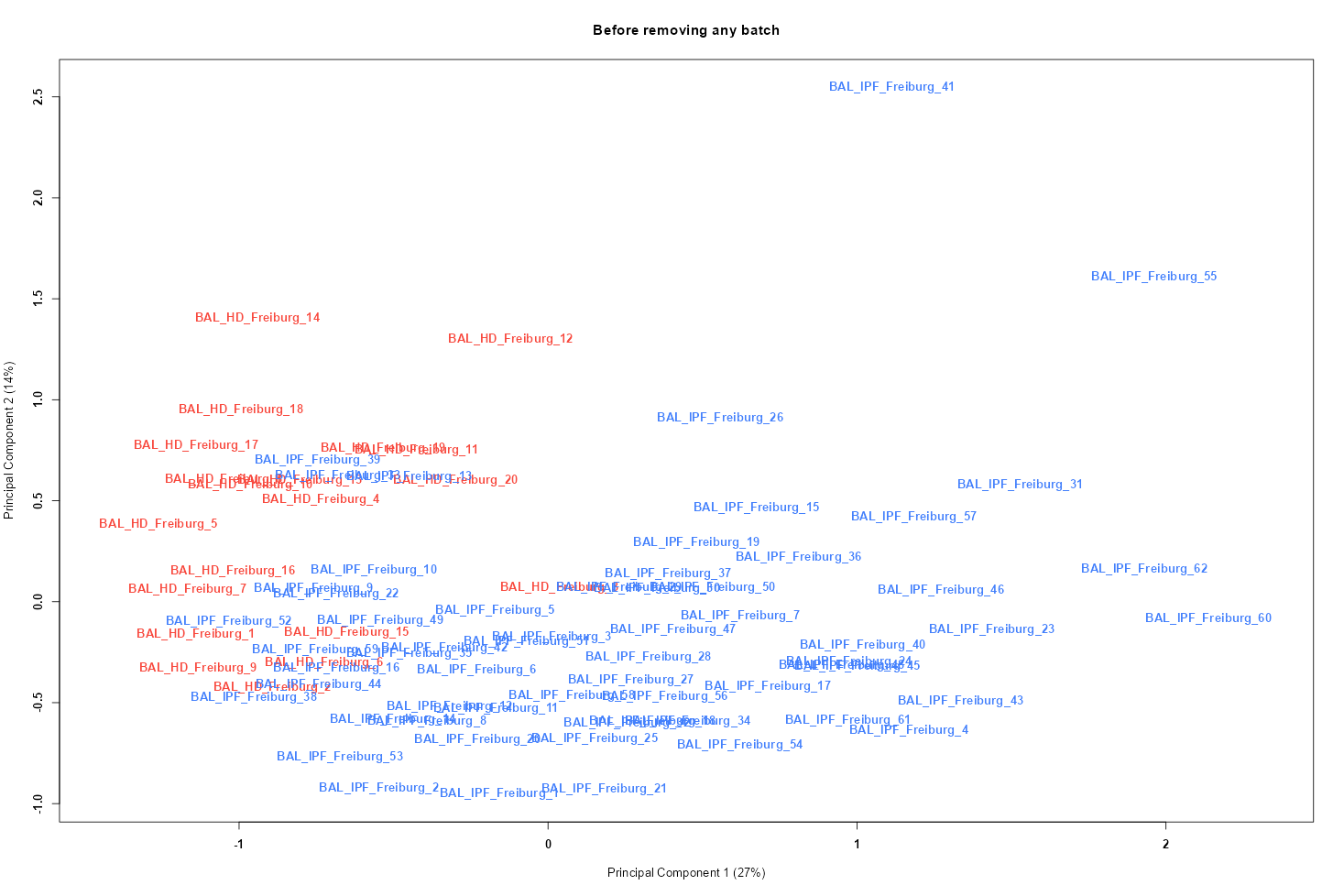

### GSE72073_boxplot_after_normalization.png

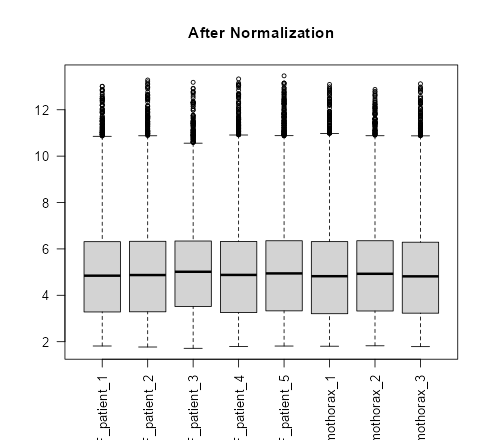

### GSE72073_MDS_before_correction.png

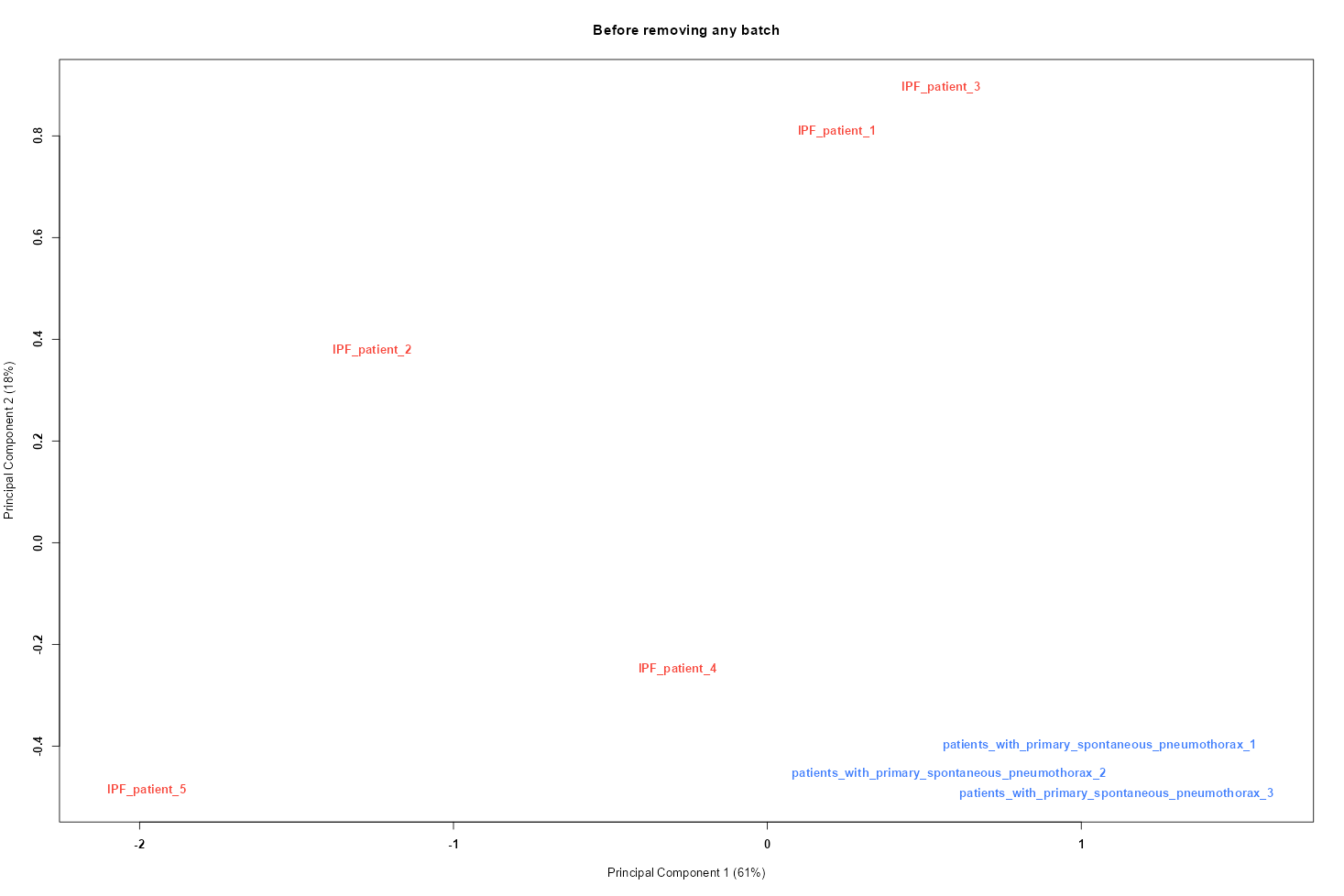

### GSE90010_boxplot_after_normalization.png

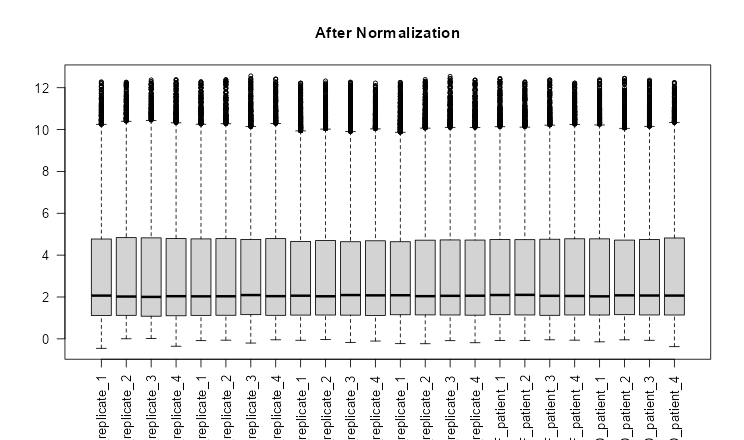

### GSE90010_MDS_before_correction.png

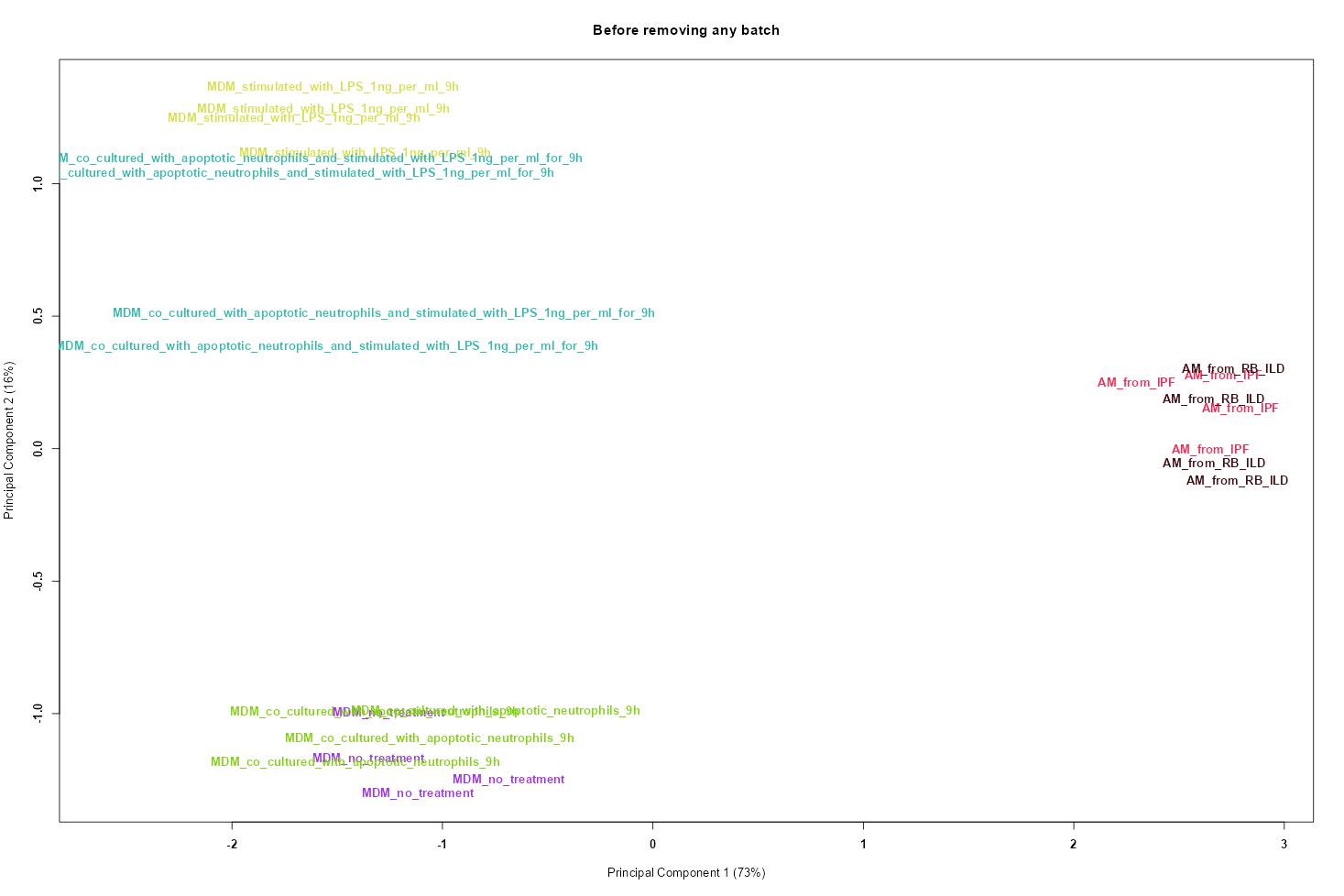

### GSE110147_boxplot_after_normalization.png

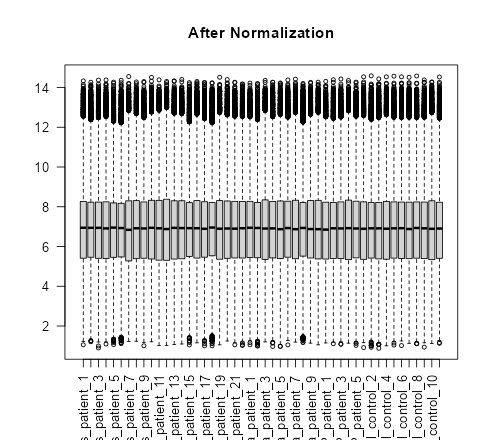

### GSE110147_MDS_before_correction.png

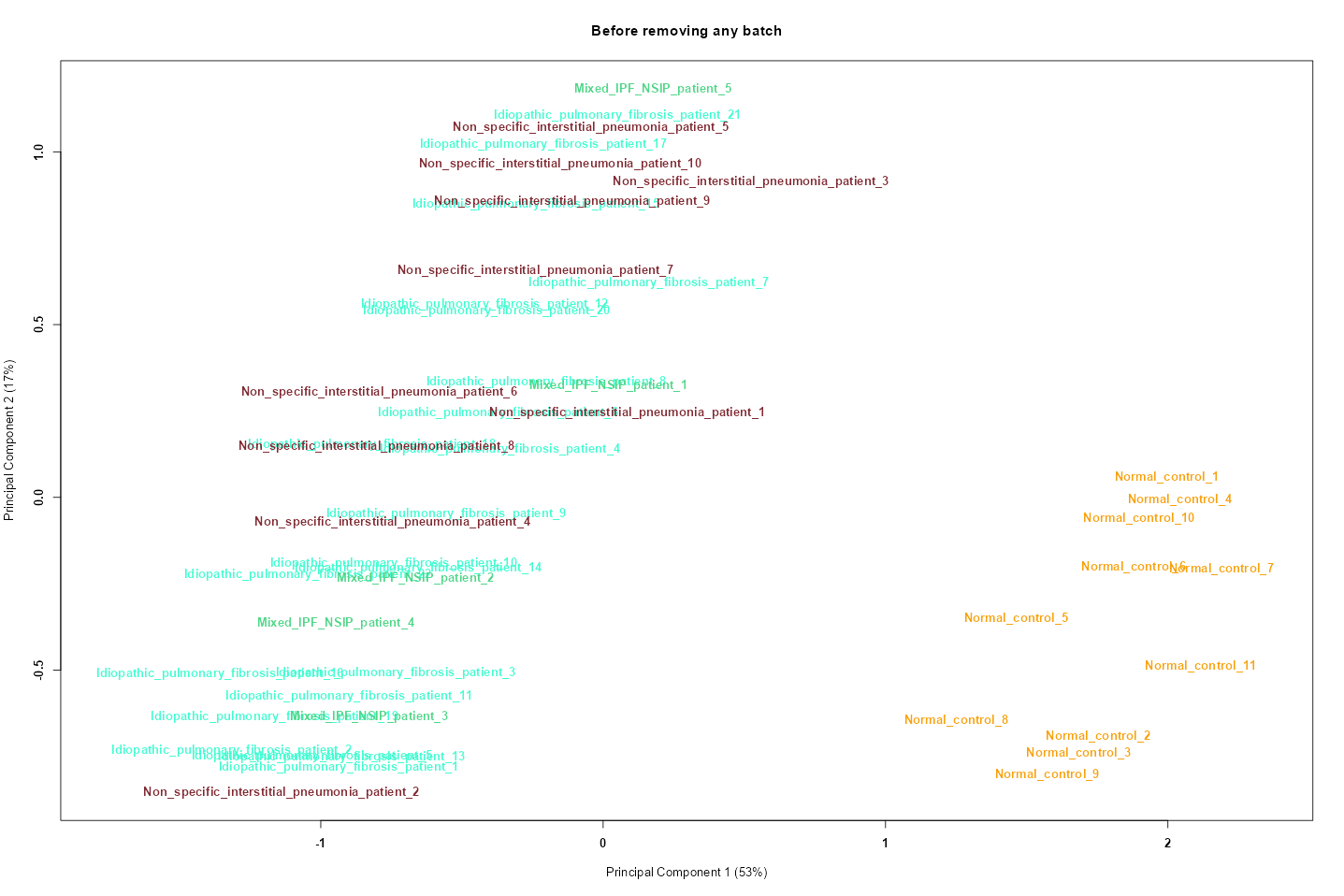

### GSE129164_boxplot_after_normalization.png

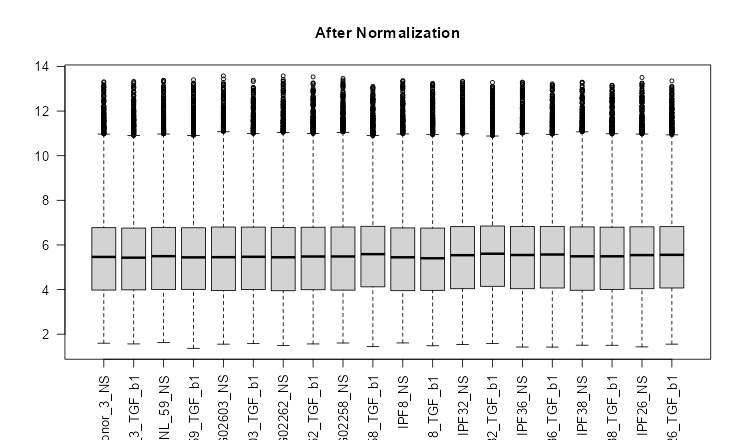

### GSE129164_MDS_before_correction.png

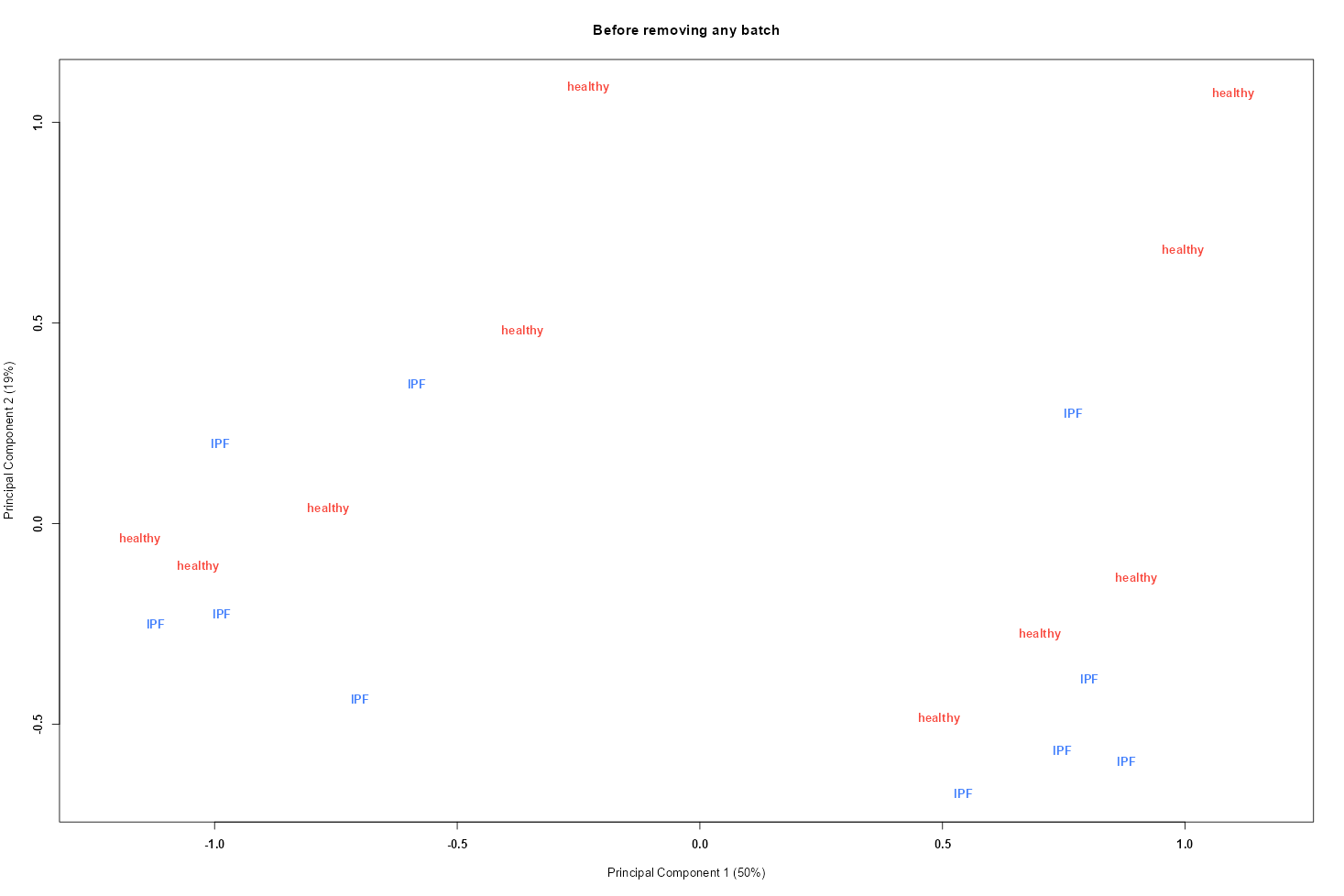

### GSE129164_MDS_before_correction_treatment.png

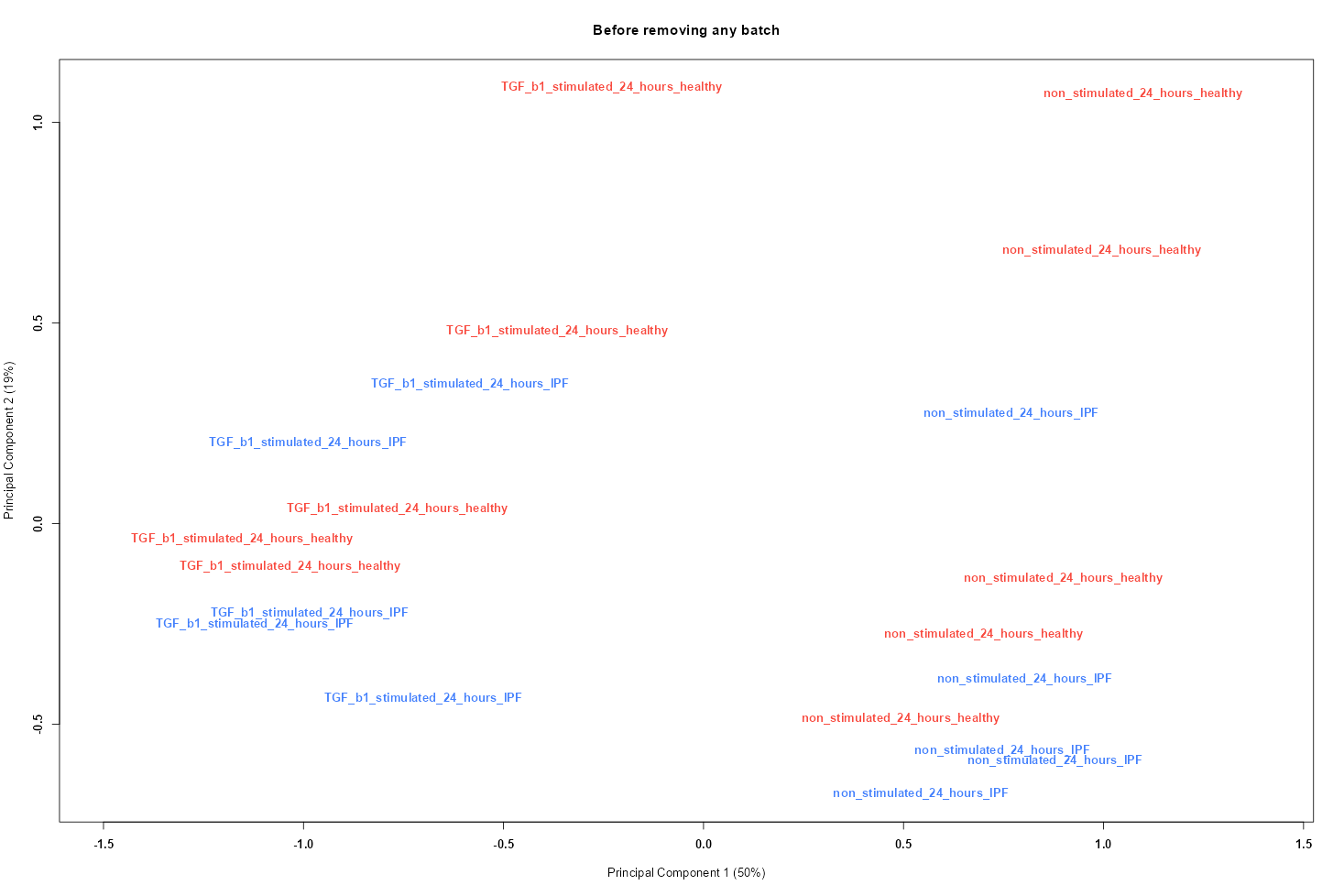

### GSE144338_boxplot_after_normalization.png

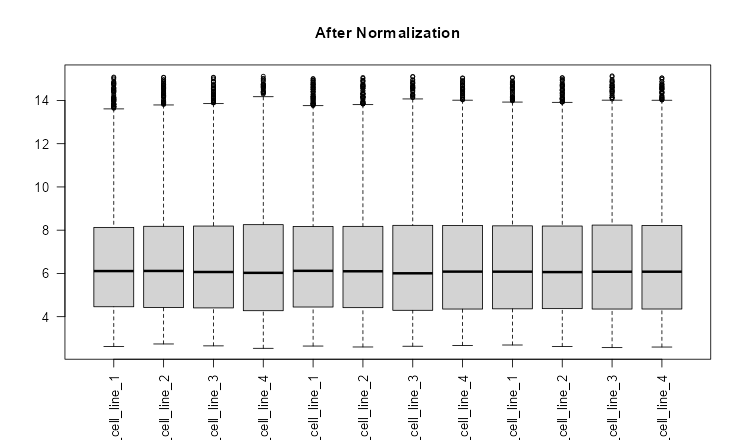

### GSE144338_MDS_before_correction.png

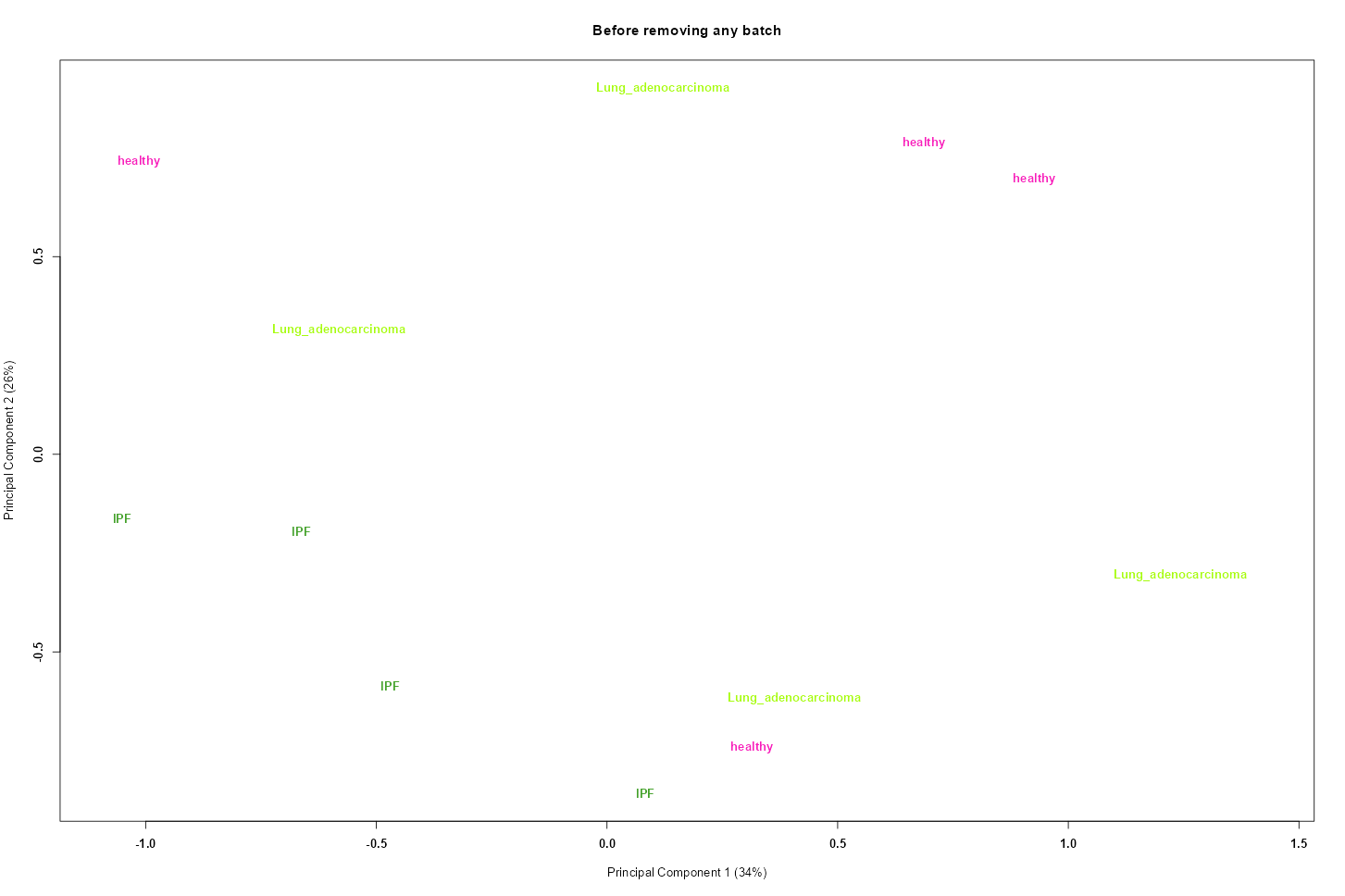

### macrophage_disease_microarray.png

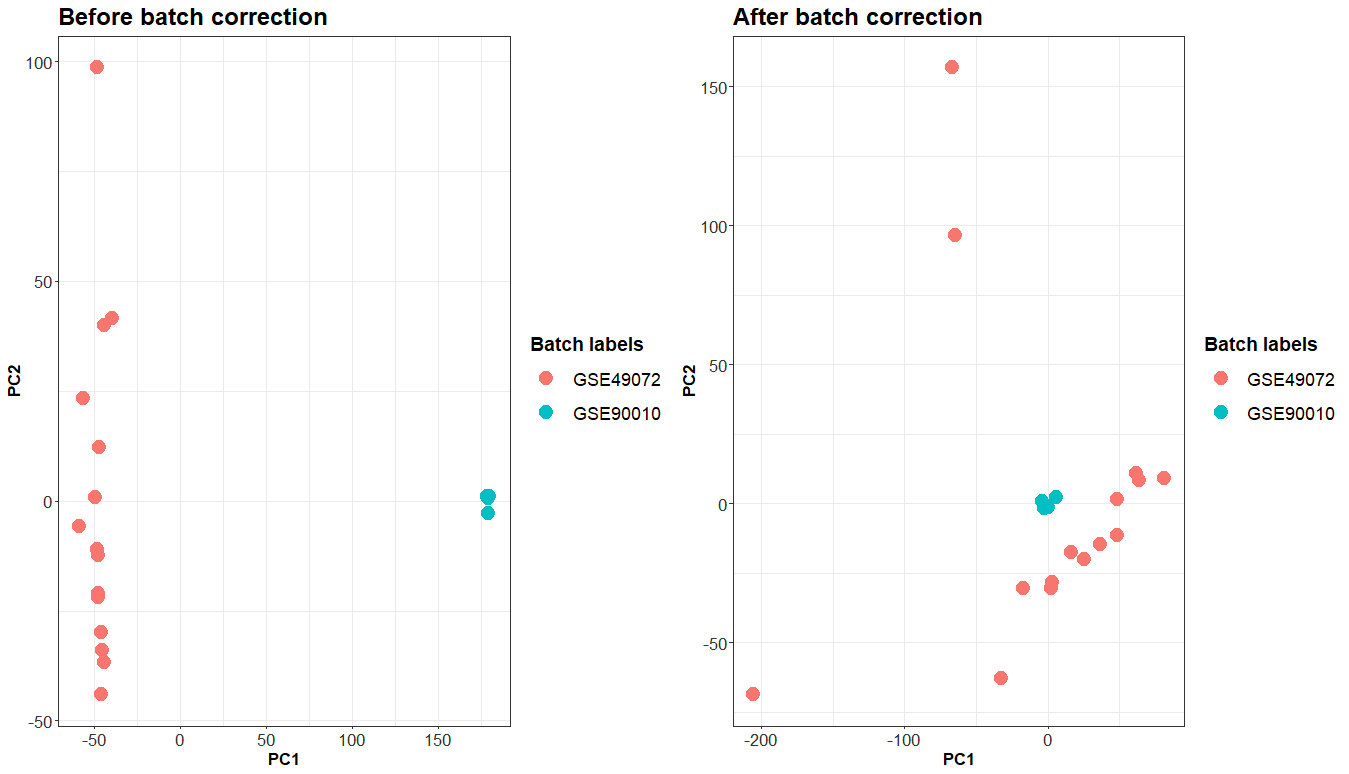

### macrophage_healthy_microarray.png

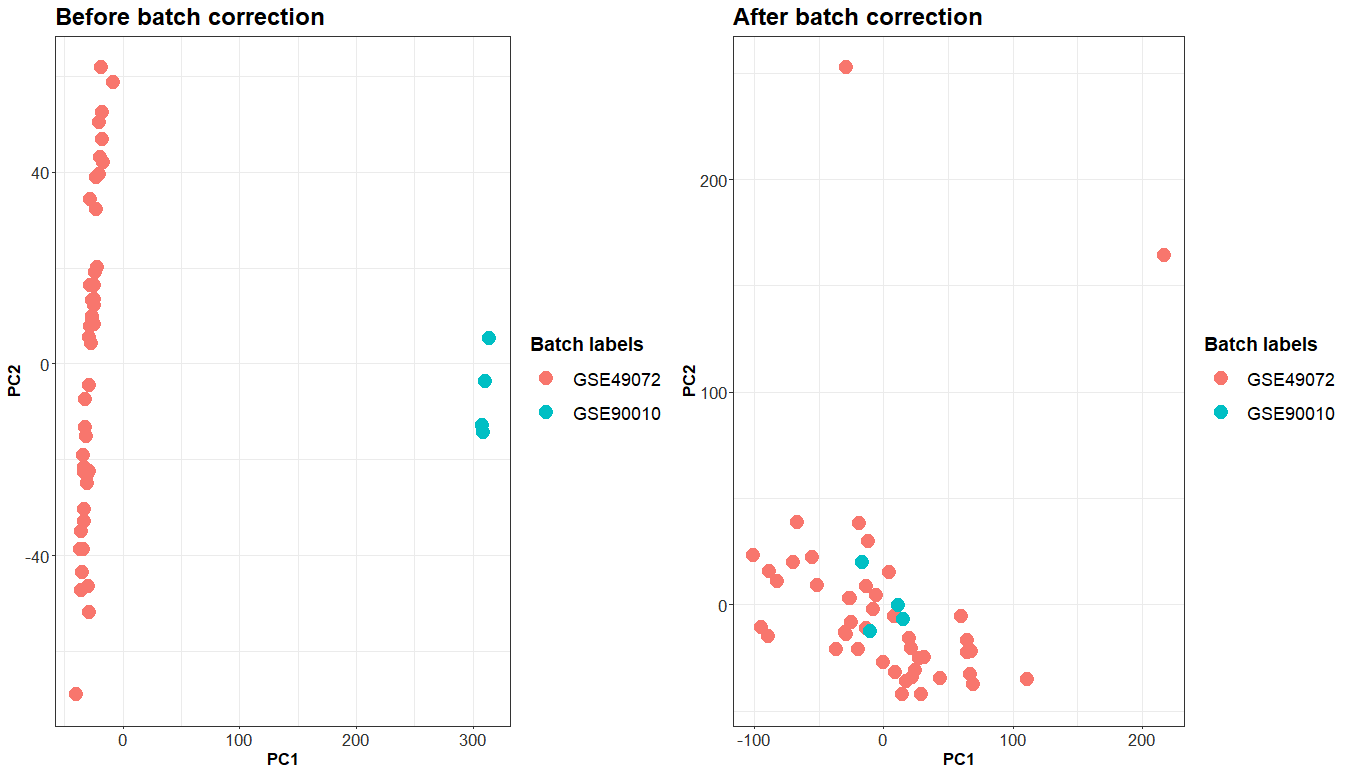

### seq_tech.png

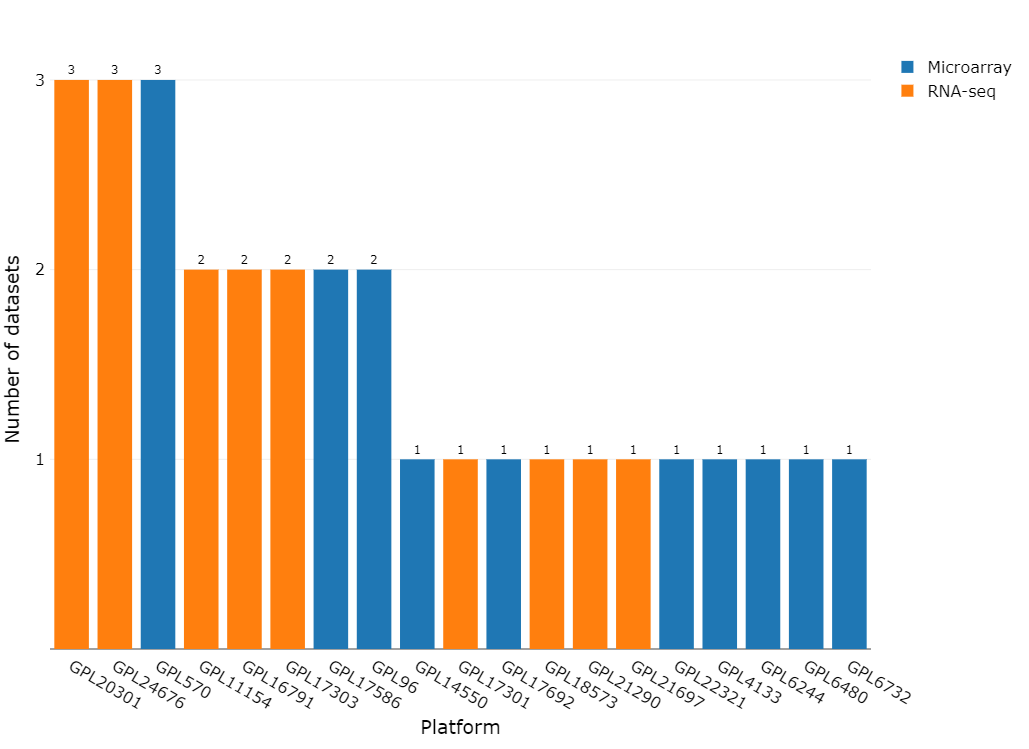

### seq_tech_samples.png

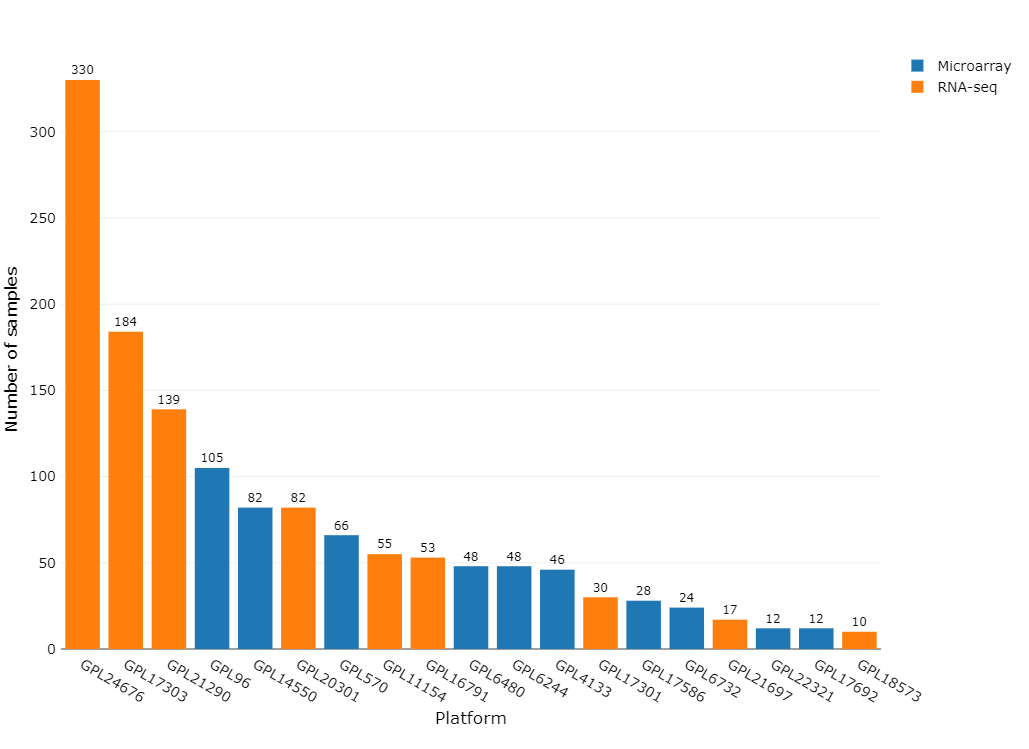

### seq_type.png

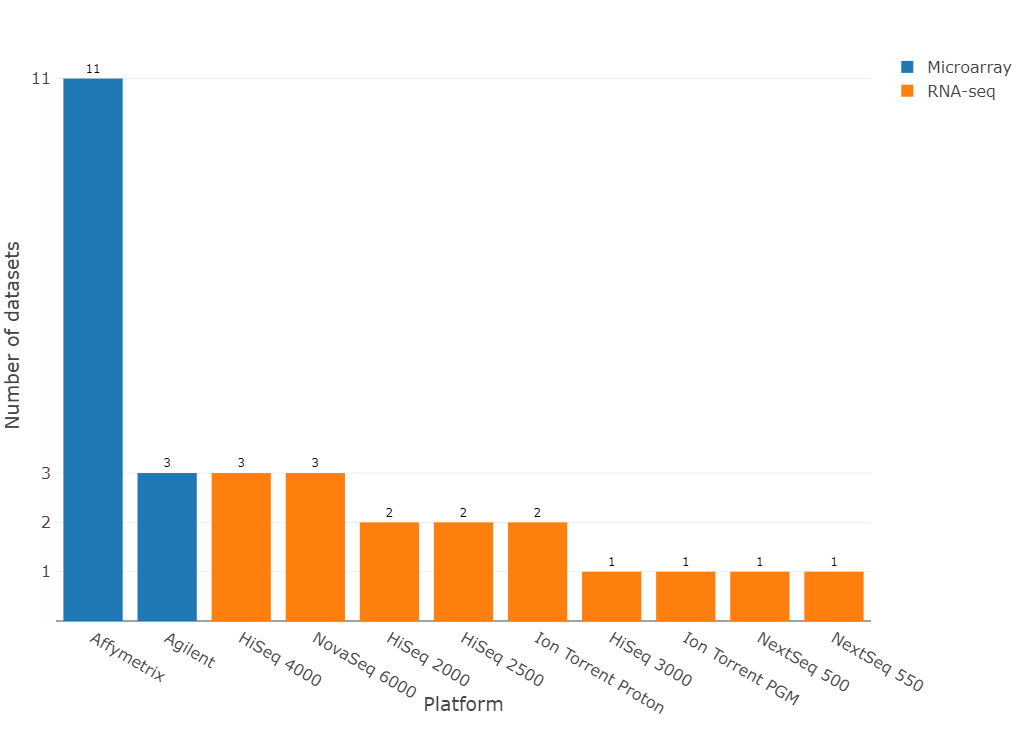

### seq_type_samples.png

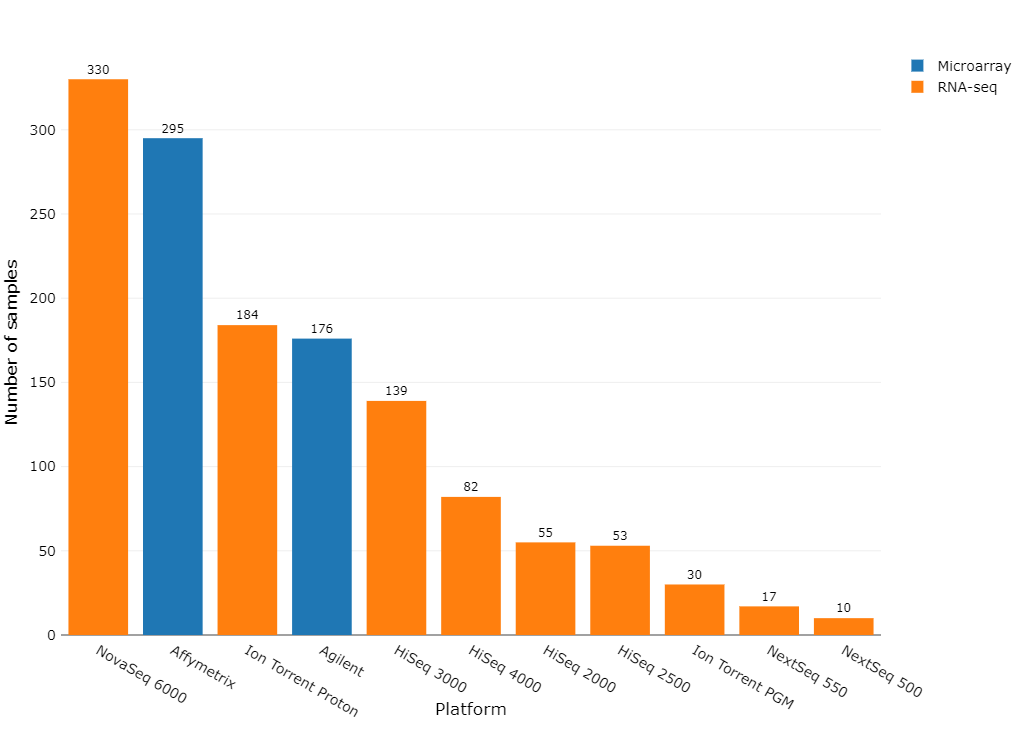
